# The peroxiredoxin Tsa1 promotes stationary phase entry by suppressing PKA activity

**DOI:** 10.1101/2025.11.26.690464

**Authors:** Víctor Garrigós, Lisa Dengler, David Henriques, Javier Buceta, Emilia Matallana, Cecilia Picazo, Jennifer C. Ewald, Agustín Aranda

## Abstract

Cells must continuously adapt their metabolism, growth, and division to fluctuating nutrient availability. In *Saccharomyces cerevisiae*, this coordination is largely governed by the Ras/cAMP/PKA pathway, which promotes fermentative growth under glucose-rich conditions and is rapidly downregulated upon glucose depletion. This downregulation enables stress responses, global metabolic rewiring, and entry into quiescence. While PKA is known to integrate multiple environmental cues with glucose availability, the crosstalk to other stress-responsive signaling mechanisms remains incompletely understood. Here, we identify the peroxiredoxin Tsa1, a central player in oxidative stress protection and redox signaling, as a key modulator of PKA activity during glucose depletion and stationary phase entry. Combining live-cell imaging of Msn2, analysis of STRE-dependent gene expression, Nth1 phosphorylation assays, and metabolic flux modelling, we show that loss of Tsa1 leads to incomplete downregulation of PKA signaling. Consequently, *tsa1*Δ cells display delayed transition to respiratory growth, reduced gluconeogenic capacity, and impaired accumulation of storage carbohydrates. Failure to properly attenuate PKA activity also disrupts accurate cell cycle control upon glucose depletion, resulting in abnormally small cells that contribute to the reduced chronological lifespan and delayed recovery upon nutrient repletion. Together, our findings establish Tsa1 as a critical link between redox regulation and nutrient signaling, ensuring proper metabolic adaptation and cellular fitness during glucose exhaustion.

## 1. Introduction

All living organisms must dynamically sense and respond to nutrient availability to maintain homeostasis, sustain growth, and ensure survival. This metabolic plasticity is particularly critical for unicellular organisms like *Saccharomyces cerevisiae*, which have evolved sophisticated signaling networks to coordinate cellular physiology with environmental cues. The Target of Rapamycin (TOR) and Protein Kinase A (PKA) nutrient signaling pathways are central to this coordination [1,2]. PKA acts as the main effector of the Ras/cAMP/PKA pathway, which senses glucose availability [3]. Under glucose-rich conditions, PKA is activated by increased intracellular cAMP levels, leading to phosphorylation of multiple downstream targets that ultimately promote fermentative metabolism, ribosomal biogenesis and cell proliferation. Conversely, glucose depletion triggers an acute decline in PKA activity, relieving repression of genes required for utilizing alternative carbon sources, accumulating storage carbohydrates, and initiating the transition to quiescence [2,4].

Beyond nutrient sensing, the PKA pathway also integrates signals from multiple stress-response modules. Under high PKA activity, stress-responsive transcription factors like Msn2 are sequestered in the cytosol, preventing the expression of the stress-responsive element (STRE)-controlled genes [5,6]. Consequently, adapting to environmental challenges, such as heat shock, osmotic, or oxidative stress, requires the precise downregulation of PKA signaling [3,7–9]. Emerging evidence suggests that multifunctional proteins, including members of the DJ-1 superfamily and peroxiredoxins, act as critical nodes in this regulatory interface, coupling stress defense with nutrient signaling [10,11]. Among these, Tsa1 is the major cytosolic 2-Cys peroxiredoxin in *S. cerevisiae*. While its canonical role involves the NADPH-dependent reduction of hydrogen peroxide, Tsa1 also functions as a potent redox relay [12–14]. Through a mechanism termed peroxiredoxinylation, Tsa1 forms transient or covalent interactions with hundreds of proteins involved in central biological processes, potentially modulating their activity and protecting the proteome under adverse conditions [15]. This dual role enables Tsa1 to play a critical role in signal integration and metabolic reprogramming [16].

One important target of peroxiredoxinylation is Tpk1, one of the catalytic subunits of PKA [15]. Previous research has established that Tsa1 can attenuate PKA activity through a direct physical interaction with Tpk1, even in the absence of exogenous oxidative stress, thereby extending yeast chronological lifespan under caloric restriction [17,18]. Furthermore, it has been reported that Tsa1 also modulates the PKA activation state by interacting and controlling the redox status of Bcy1, the regulatory subunit of the PKA complex [19]. Despite these insights, the complex interplay between Tsa1 and PKA signaling remains incompletely understood. Specifically, it is not yet clear under which nutritional conditions Tsa1-mediated inhibition of PKA is physiologically relevant, how this regulation impacts global metabolism, and whether it plays a decisive role during nutrient transitions.

Here, we investigated whether Tsa1 exerts control over PKA and cellular physiology during fermentative growth on glucose, during the diauxic shift, and upon entry into stationary phase. Deletion of *TSA1* led to a delay in the switch to respiratory growth, decreased gluconeogenic flux during growth on ethanol, and reduced accumulation of the storage carbohydrate trehalose. Live-cell imaging of Msn2, expression analysis of STRE-regulated genes, and analysis of Nth1 phosphorylation demonstrate that these phenotypes are a direct consequence of the incomplete reduction of PKA activity. Furthermore, the *tsa1*Δ mutant failed to arrest cell division upon stationary phase entry, resulting in a subpopulation of extremely small cells characterized by a decreased chronological lifespan and delayed recovery after glucose replenishment. These findings underscore a previously unappreciated role for Tsa1 in coordinating metabolic and proliferative responses upon glucose depletion through PKA regulation.

## 2. Materials and methods

### 2.1. Strain construction and cultivation conditions

All yeast strains used in this study were W303 derivatives. Most strains were prototrophic, except for several mutants that were uracil auxotrophic, as indicated in **S1 Table**. Yeast transformations were carried out using the lithium acetate method [20], and mutants were generated via standard PCR-based homologous recombination or integration of linearized plasmids. Plasmid containing *NTH1*-reporter were linearized with StuI in the *URA3* region and integrated into the *URA3* locus of the uracil auxotrophic W303 strain. All constructs were Sanger-sequenced for verification. See Supplementary **S2 Table** for details of plasmids used in this study.

For all reported experiments, yeast cells were grown in 100 ml of glucose minimal medium (GMM) (10 g/l glucose, 1.7 g/l yeast nitrogen base without amino acids (US Biological), 5 g/l ammonium sulfate, 50 mM potassium phthalate, pH adjusted to 5 with KOH). Cultures were incubated at 30°C with continuous shaking (200 rpm) and inoculated at an initial optical density (OD_600_) of 0.2 from exponentially growing cell pre-cultures. For starvation experiments, after 72 h of growth, cells were released from the stationary phase by 1:5 dilution into fresh GMM. For growth in solid media, YPD (20 g/L glucose, 20 g/L bacteriological peptone, 10 g/L yeast extract), SC (20 g/L glucose, 1.7 g/L yeast nitrogen base (w/o amino acids and ammonium sulfate), 5 g/L (NH_4_)_2_SO_4_, 2 g/L Drop-Out complete (Formedium)), and SCEtOH (same composition as SC but replacing glucose with 2% (v/v) ethanol) were prepared by supplementing with 2% (w/v) agar.

Chronological life span (CLS) experiments were assessed following established protocols [21]. Briefly, overnight precultures in GMM of selected strains were inoculated in 25 mL of GMM medium at an initial OD_600_ of 0.1 and maintained at 30°C with continuous shaking (180 rpm). Starting from day 2 of growth (defined as 100% survival), aliquots were periodically sampled, serially diluted, and plated onto YPD agar. Following 48 hours of incubation at 30°C, colony-forming units (CFUs) were quantified to determine the percentage of survival relative to the day 2 baseline.

### 2.2. Extracellular metabolite quantification

High-performance liquid chromatography (HPLC) was used to quantify glucose, ethanol, acetate, pyruvate, and succinate in the supernatants of the GMM samples. HPLC analysis was performed on a Shimadzu HPLC system equipped with a Repromer H column from Maisch GmbH, Germany. A sample volume of 10 μL was injected onto the column using an autosampler cooled to 4 °C. Metabolites were eluted isocratically with 5 mM H_2_SO_4_ (sulfuric acid solution, 49 to 51%, for HPLC, Honeywell Fluka), degassed at −95 kPa with an in-line degassing unit (DGU-20A 5R). Samples were run with a flow rate of 1.0 mL/min for 25 min at 40 °C. The analytes were monitored using a refractive index detector (Shimadzu RID-20A). Analysis was performed with the Shimadzu Labsolution software. Metabolite concentrations were calculated by interpolation from the standard curve for each assayed metabolite. Glycerol concentration was determined spectrophotometrically using coupled enzymatic reactions linked to NAD^+^/NADH redox pairs, using the commercial kit from Megazyme Ltd. (Bray, Ireland; K-GCROL kit) and following the manufacturer’s protocol.

### 2.3. Intracellular metabolite quantification

The endometabolomic analysis of central carbon metabolism was performed by the MetaToul-MetaboHUB platform (Toulouse Biotechnology Institute, France). Intracellular metabolites were extracted and quantified using an isotope dilution mass spectrometry (IDMS) approach. Briefly, culture samples at 24 h of growth were rapidly centrifuged at 4°C, and the resulting pellets were resuspended in 1 mL of a cold extraction solution (Acetonitrile/Methanol/H_2_O, 4:4:2, containing 125 mM formic acid). For absolute quantification, 50 µL of a U-^13^C-labeled yeast internal standard, derived from cells grown on U-^13^C-glucose, was added to each sample [22,23]. Following a 1-hour extraction at −20°C, samples were centrifuged (5 min, 2000 g), and the supernatants were evaporated in a SpeedVac before resuspension in ultrapure water. Targeted analysis of central carbon metabolites was performed using an Ion Chromatography (IC) system coupled to High-Resolution Mass Spectrometry (HRMS). The platform consisted of a Dionex ICS-5000+ Reagent-Free HPIC system (Thermo Fisher Scientific) equipped with an electrolytic eluent generator for automatic KOH production and an IonPac AS11-HC column (250 x 2 mm) with an AG11-HC guard column. Analytes were separated at 0.38 mL/min using a 50-min linear KOH gradient (7 to 100 mM). Mass detection was conducted on a QExactive+ mass spectrometer (Thermo Fisher Scientific) in negative electrospray ionization (ESI) mode at a resolution of 70000 (at 400 m/z). The source parameters included a capillary temperature of 350°C, a sheath gas flow of 50 a.u., and a source voltage of 2.75 kV. Data acquisition was performed via Xcalibur software, and metabolites were identified by extracting the exact mass with a tolerance of 5-10 ppm. Data processing and quantification were carried out using Skyline, MSReader, and GraphStatsR.

For intracellular trehalose quantification, 1.5 ml of yeast cell culture was sampled and chilled on ice for 1 min. Cells were harvested by centrifugation at 13300 rpm for 1 min, washed once with 750 µl minimal medium without glucose, frozen in liquid nitrogen and stored at − 80 °C. Trehalose was extracted by adding 200 µl of boiling water (90 °C) to the frozen cell pellets, followed by incubation in a boiling water bath (90 °C) for 5 minutes. During incubation, samples were vortexed every 2 minutes. After centrifugation at 13300 rpm for 5 minutes, the supernatants were transferred to fresh tubes. A second extraction was performed under the same conditions, and the resulting supernatants were combined [24]. Extracted trehalose samples were stored at −20 °C. Trehalose concentration was determined enzymatically using a commercial kit (K-TREH, Megazyme).

### 2.4. Dynamic flux balance analysis

To estimate intracellular fluxes during the diauxic shift, a system of ordinary differential equations (ODEs) was combined with the Yeast 9 genome-scale metabolic network for Flux Balance Analysis (FBA) [25]. The model presented here extends the work of Moimenta et al. [26]. Briefly, it consists of a set of differential equations describing biomass formation and the main extracellular metabolites measured in this study. However, it differs substantially from the previous work due to the inclusion of oxygen and the consumption of a non-fermentable carbon source following glucose depletion. The ODEs and specific modelling assumptions for the genome-scale metabolic network are detailed in the supplementary materials. In short, the rates obtained from the dynamic model were used to constrain the exchange fluxes of the measured metabolites and the growth rate. Since metabolic models require biomass values in dry weight (DW), we simulated several theoretical scaling factors to convert OD_600_ measurements into DW, using reference yield values from aerobic glucose-grown *S. cerevisiae* [27]. A scaling factor of 0.265 gDW/OD_600_ provided the best fit to our experimental data, being adopted for subsequent calculations. Intracellular fluxes were then computed at selected time points corresponding to ethanol- and glucose-dependent growth using the parsimonious FBA (pFBA) approach. To accurately capture the experimental dynamics, we formulated a nonlinear optimization problem with the AMIGO Toolbox [28], employing a log-likelihood function based on the standard deviations of the replicates. Relevant metrics for goodness of fit and parameter confidence are also given in the Supporting Information section (**S1 Appendix**). Flux simulations were performed using the RAVEN Toolbox [29] and the Gurobi solver. For comparison purposes, average fluxes were derived by calculating the area under the curve of each flux (mmol·DW⁻¹·h⁻¹), multiplying it by the biomass (mmol), and normalizing by the total amount of glucose or ethanol consumed [30]. All supporting information and scripts necessary to reproduce the dFBA results are available in the **S2 Appendix**.

### 2.5. RNA extraction and analysis

To quantify the relative expression levels of the target genes, yeast cells were harvested at the specified time points, and total RNA was extracted as described previously [31]. Extracted RNA was reverse transcribed using the NZY First-Strand cDNA Synthesis Kit (NZYtech, Portugal). Gene-specific primers were used to amplify *ALD4*, *CIT2*, *TPS1, TPS2, NTH1 and ATH1*, using the *ACT1* gene as an internal control [32–34]. Quantitative PCR (qPCR) was performed using the NZYSpeedy qPCR Green Master Mix (NZYtech, Portugal) on a QuantStudio 3 instrument (Thermo Fisher Scientific), following the manufacturer’s instructions. Each reaction was carried out in triplicate, and the average cycle threshold (Ct) value from the triplicates was used for analysis. Relative transcript levels were calculated using the 2^−ΔΔCt^ method [35].

### 2.6. NEM-mPEG assay and SDS-PAGE Western blot

The oxidation state of cysteine residues in the Nth1 protein was analyzed in cells expressing Myc-tagged versions of Nth1 using the *N*-ethylmaleimide (NEM)-methoxy polyethylene glycol 5000 (mPEG) assay, following the protocol described [36]. To monitor Tps1, Tps2, and Nth1 protein levels, cells expressing Myc-tagged versions of these proteins were lysed using glass beads, and whole-cell extracts were prepared in lysis buffer containing 1 M Tris-HCl (pH 7.5), 5 M NaCl, 1 M MgCl_2_, 10% (v/v) NP-40, 0.1 M PMSF, and a commercial protease inhibitor tablet (COMPLETE Mini, EDTA-free; Roche). In both assays, protein concentration was determined using the DC Protein Assay (Bio-Rad), according to the manufacturer’s instructions. Samples were separated by SDS-PAGE using an Invitrogen mini-gel system and transferred to PVDF membranes using a Novex semi-dry transfer system (Invitrogen, Carlsbad, CA, USA). Immunodetection was performed using an anti-Myc antibody (Santa Cruz Biotechnology). Anti-Pgk1 (Invitrogen) and Ponceau S staining were used as loading controls. Signal detection was carried out using the ECL Western Blotting Detection System (GE Healthcare).

### 2.7. Phos-tag SDS-PAGE Western blot

Cells growing in GMM were directly sampled at specific time points into two volumes of 75% methanol, 15 mM Tris–HCl (pH 7.5), pre-chilled to −20 °C, and centrifuged at 4000 rpm at 4 °C. Pellets were frozen in liquid nitrogen and stored at −80 °C. Lysates were prepared by bead beating in urea buffer (20 mM Tris–HCl, pH 7.5; 2 M thiourea; 7 M urea; 65 mM CHAPS; 65 mM DTT), supplemented with 1× EDTA-free protease and phosphatase inhibitor cocktails (GoldBio).

To resolve phosphorylated species of the Nth1-reporter, 10% SDS-polyacrylamide gels (29:1 Bio-Rad) were used, supplemented with 100 μM Phos-tag™ Acrylamide AAL-107 (FUJIFILM Wako) and 200 μM MnCl_2_. Gels were run at 15 mA/gel using pre-chilled running buffer, with the electrophoresis chamber maintained on ice, until the dye front reached the bottom of the gel. For Western blotting, proteins were transferred using a dry blotting system (iBlot, Invitrogen), and the Nth1-reporter was detected using an anti-FLAG M2 antibody (Sigma, product number: F1804) and an anti-mouse HRP-conjugated secondary antibody (Promega, W4021). Luminescence was imaged using a Licor Odyssey FC system.

### 2.8. Enzymatic activities

Cell-free extracts for enzymatic analysis were prepared from cells harvested at specific growth time points. Cells were collected by centrifugation, washed with sterile water and subsequently resuspended in the appropriate lysis buffer for each enzymatic activity. Cell disruption was performed using glass beads in a Precellys Evolution homogeniser (Bertin Technologies, France). Protein concentration was determined using the DC Protein Assay (Bio-Rad), following the manufacturer’s instructions. For trehalose-6-phosphate synthase (TPS) activity, cell-free extracts were prepared using a cold lysis buffer (25 mM MES, pH 7.1). TPS activity was assessed using a discontinuous method, as previously described [37]. One unit of TPS activity was defined as the amount of enzyme that catalyzes the formation of 1 nmol of UDP per min at 37 °C. For the determination of neutral and acid trehalase activities, cell-free extracts were prepared using 10 mM MES (pH 6.0) as lysis buffer. Both enzymatic activities were assayed as described above [38,39], with the pH of the incubation buffer adjusted to selectively favour either neutral or acid trehalase activity. The resulting glucose concentration in the supernatants was determined using the glucose oxidase/peroxidase assay and expressed as nmoles of glucose released per min and milligram of total protein.

### 2.9. Microscopy and image analysis

#### 2.9.1. Live-cell imaging of Msn2 subcellular localization

To monitor the real-time nucleocytoplasmic shuttling of Msn2-NeonGreen, cells were grown on GMM for 72 h until stationary phase. Cells were sonicated at a low power for 4 s and loaded onto a commercial microfluidics system (Y04C-02 plates, CellASIC ONIX2 system, Merck). During growth, the glucose medium was supplied with a pressure of 3 psi. The temperature was kept constant at 30 °C using an incubator chamber surrounding the imaging system (Okolab Cage Incubator, Okolab USA INC, San Bruno, CA). Cells were grown for 7 h in GMM. Then, the medium flow was stopped, resulting in a fast consumption of glucose, mimicking the diauxic shift.

Live-cell imaging experiments were performed on a Nikon Ti2 inverted epifluorescence microscope (Nikon Instruments, Japan) with a Lumencor SPECTRA X light engine (Lumencor, Beaverton, USA), a Photometrics Prime 95 (Teledyne Photometrics, USA) backilluminated sCMOS camera. The system was programmed and controlled by the Nikon software NIS Elements. Focus was maintained using the Nikon “Perfect Focus System”. The time-lapse images were taken using a Nikon PlanApo oil-immersion 40x objective with a frequency of 3 min. Htb2-mCherry was imaged at 10 % intensity and 100 ms exposure (excitation filter: ET585/29x, emission filter: ET650/60), and Msn2-Neongreen was imaged at 30 % intensity and 300 ms exposure (excitation filter: ET500/20x, emission filter: ET535/30m).

When the flow was stopped, comparable amounts of cells were obtained for each strain and images were recorded at 12-bit grey scale and then converted to tiffs in Cell-ACDC [40]. Cell segmentation was performed in Cell-ACDC using YeaZ (minimum area: 10 pixels, minimum solidity 0.5, maximal elongation 3.0). For each image, the median background fluorescence (cell-free area) was determined and subtracted from the signal at each time point. Htb2 was segmented using automatic Otsu thresholding with a Gaussian filter of 1.0. Using the cell and nuclear segmentation mask, the volume of the cytosol and nucleus was calculated as described [40]. Msn2 concentrations were determined by normalizing the amount (calculated as follows: amount = (mean_obj - median_background)·area_obj) to the volume of the nucleus and cytosol, respectively.

#### 2.9.2. Cell viability and morphological analysis

For propidium iodide staining (PI) (Sigma-Aldrich 537059) to evaluate cell survival, 500 μl of cells were washed twice in PBS buffer (137 mM NaCl, 2.7 mM KCl, 4.4 mM Na_2_HPO_4_·7H_2_O, 1.4 mM KH_2_PO_4_, pH 7.3) and 5 μL of a 1 mg/mL stock solution of the dye were added to be then incubated in darkness for 30 min. Cells were washed in PBS and visualized. Microscopy images were acquired using a Leica Thunder inverted microscope equipped with a CMOS camera (Leica DFC9000 GTC). Images used for quantification were captured with a FLUOTAR L 20×/0.4 dry objective, while representative zoomed images were acquired using a HC PL APO 40×/0.95 dry objective. In all cases, Köhler illumination was applied to optimize contrast in transmitted light mode. Dye excitation was performed using a Spectra-X Light Engine LED lamp (8 solid-state LEDs). For the propidium iodide observation, excitation was carried out at 555 nm (30% power) with an exposure time of 150 ms, using a GFP/mCherry filter setting (Excitation (nm): 470/40+573/35; Emission (nm): 520/40+635/65) combined with an external filter wheel at 642/80 nm. Phase contrast images were taken with a 50 ms exposure, and Differential Interference Contrast (DIC) images with a 100 ms exposure. Microscope control and image acquisition were managed using Leica Application Suite X (LAS X) software.

To quantify fluorescence per pixel as a function of cell size, large fields of view containing approximately 10^4^ cells were acquired. Cells were segmented from the phase contrast channel using a machine learning algorithm [41], producing a binary mask. A custom script written in Mathematica (Wolfram Research Inc, Champaign, IL) was then used to filter individual cells with radii between 1-3.5 μm and to compute the average fluorescence intensity per pixel for each cell. The cell radius was estimated assuming a circular geometry such that R=Sqrt(n_p_/pi), where n_p_ indicates the number of pixels occupied by a cell mask. Within this filtered dataset, the maximum mean fluorescence intensity per pixel was identified, and all intensity values were normalized to this maximum, yielding relative fluorescence per pixel values ranging from 0 to 1.

For statistical analysis, two correlation measures were computed between the equivalent cell radius (in pixels) and the normalized mean fluorescence per pixel: Pearson correlation and the Spearman rank correlation. In both tests we computed the corresponding p-value. Statistical significance was defined at the conventional threshold p < 0.05. To visualize the tendency and variability between the normalized mean intensity per pixel and the cell size (pixels) independent of uneven point density, the data were divided into bins of equal cell count (using quantiles of the radius distribution). For each bin, the mean radius, mean fluorescence, and standard error of the mean (SEM) were computed. The code necessary to reproduce the microscopy results and analysis is available in the **S2 Appendix**.

### 2.10. Cell size determination

Cell size measurement and counting were performed using the CASY TT Cell Counter & Analyzer (OMNI Life Science, Germany). Culture samples were collected and immediately placed on ice. To avoid cell aggregation, samples were briefly sonicated. Cells were subsequently diluted in PBS buffer based on their OD_600_ and measured with a 60 μm capillary. Only cells with diameters ranging from 1.99 to 9.00 μm were considered for analyses. Data were normalized, and cumulative distribution functions were calculated for each data point.

## 3. Results

### 3.1. *TSA1* deletion affects growth and division after glucose depletion

To investigate the impact of *TSA1* deletion on yeast growth during fermentative and post-diauxic phases, cells were cultured for 72 h in glucose minimal media (GMM). During the initial 8 h of fermentative growth, no significant differences were observed between the wild-type and the *tsa1*Δ mutant (**Fig 1A**). However, as glucose levels declined and cells approached the diauxic shift, the mutant exhibited a slight reduction in OD_600_ that persisted during respiratory growth at 24h (**Fig 1B**). Consistently, while *TSA1* deletion did not impair glucose consumption, ethanol uptake appeared delayed, as evidenced by higher extracellular ethanol levels in the mutant at 24 h. By 48 h, both glucose and ethanol were fully consumed in both cultures. These findings suggest that the major cytosolic peroxiredoxin does not significantly affect fermentative metabolism but does influence the transition to respiratory growth under these conditions.

**Figure 1.**
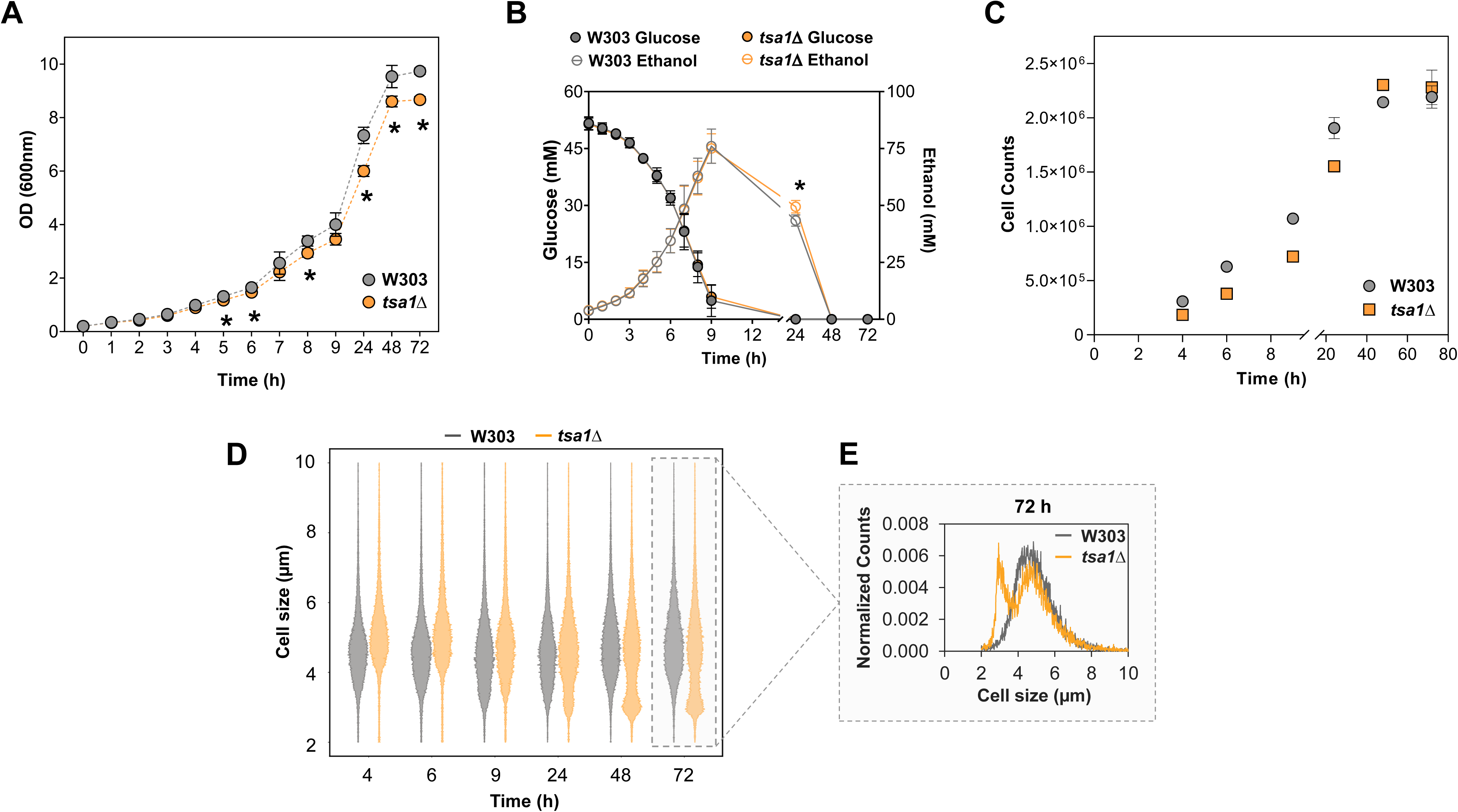
Deletion of *TSA1* impairs metabolic transitions and alters cell size distribution during long-term cultivation. **A)** Growth kinetics of wild-type and *tsa1*Δ strains during batch cultivation in GMM over 72 h, monitored by optical density (OD) at 600 nm. **B)** Extracellular glucose and ethanol concentration profiles in those conditions. **C)** Cell proliferation dynamics of wild-type and *tsa1Δ* strains; data represent two independent biological replicates. **D)** Evolution of cell size distribution in wild-type and *tsa1*Δ populations across different growth phases. **E)** Normalized cell size distribution at the 72 h time point, highlighting the emergence of a small-cell subpopulation in the *tsa1Δ* mutant. Cell size measurements were performed in triplicate; one representative distribution is shown. Unless otherwise stated, all data represent the mean and standard deviation of at least three biological replicates. Significant differences (*p < 0.05, Student’s t-test) between the *tsa1*Δ mutant and the parental strain at each specific time point are shown.

Interestingly, following ethanol depletion at 48 h, the *tsa1*Δ strain maintained a reduced OD_600_ despite reaching higher cell counts (**Fig 1C**), suggesting a shift in the population’s size distribution. Indeed, during the slow-growth phase associated with ethanol respiration and the transition to stationary phase (24–72 h), the cell size distribution of the *tsa1*Δ strain became bimodal (**Fig 1D**). The mutant developed a subpopulation of smaller cells, suggesting that it undergoes additional replication cycles, producing daughter cells that fail to achieve normal size (**Fig 1E**). These results suggest that Tsa1 contributes to halting proliferation in response to nutrient depletion, thereby facilitating the proper entry into stationary phase.

### 3.2. Tsa1 is required for proper storage carbohydrate accumulation and long-term viability

Carbon storage metabolism constitutes a critical pathway for the establishment of the stationary phase, where the accumulation of glycogen and trehalose serves as a hallmark of the transition into quiescence [42,43]. Given that our previous studies implicated Tsa1 in trehalose metabolism during the industrial propagation of wine yeast [44,45], we sought to determine whether this role represents a conserved regulatory feature under defined laboratory conditions. We found that while trehalose accumulation occurred following the diauxic shift in both wild-type and *tsa1*Δ strains, levels were significantly reduced in the mutant during both the diauxic shift and the transition to stationary phase (**Fig 2A**). Complementary iodine vapor staining confirmed a consistent glycogen storage defect in *tsa1*Δ cells across diverse growth conditions, including rich (YPD) and synthetic complete media containing either glucose (SC) or ethanol (SCEtOH) as the primary carbon source (**Fig 2B**). Together, these results demonstrate that Tsa1 is required for orchestrating storage carbohydrate homeostasis in response to nutrient depletion.

**Figure 2.**
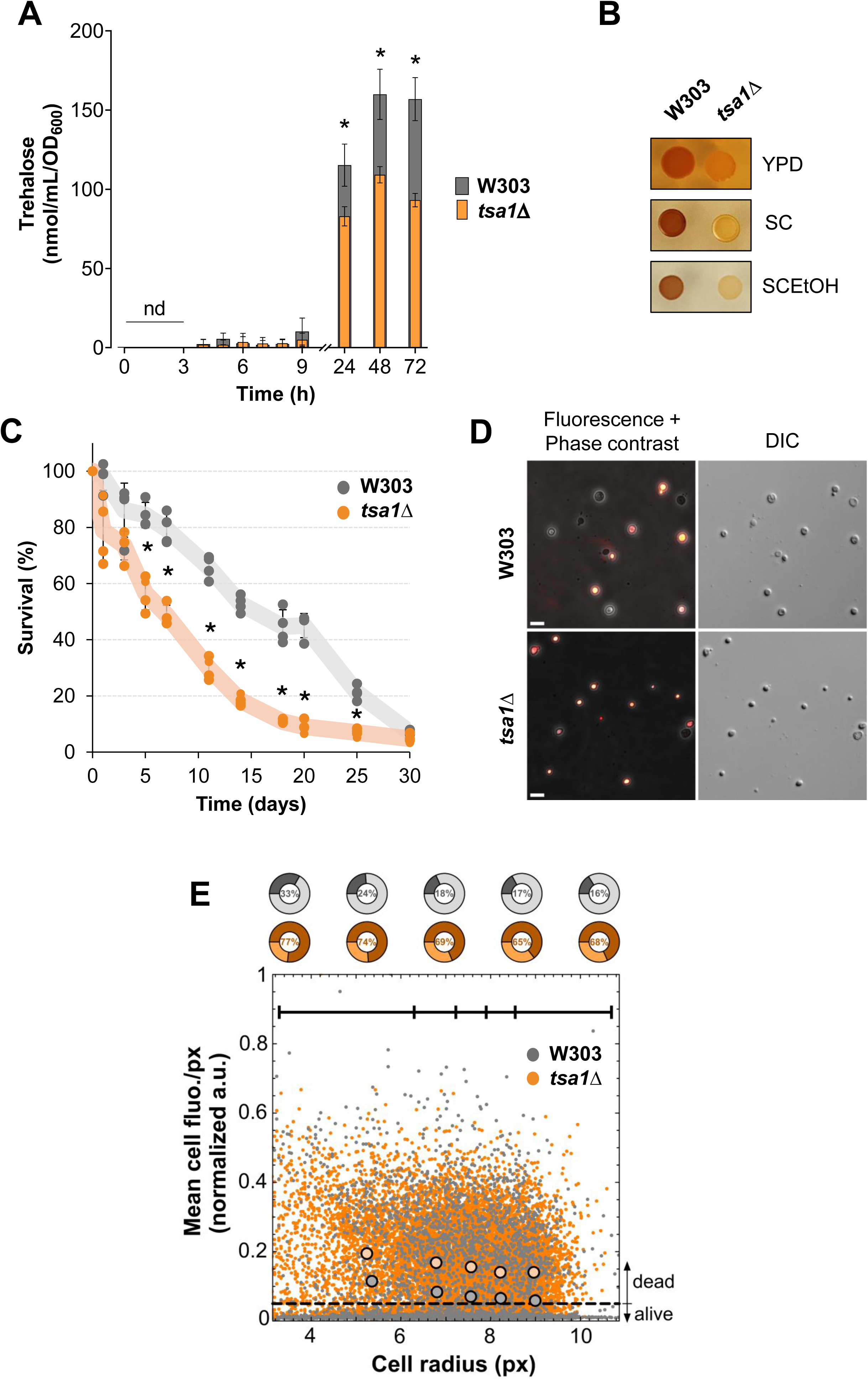
Tsa1 is required for the coordination of storage carbohydrate accumulation and maintains viability during chronological aging. **A)** Intracellular trehalose concentrations normalized to cell density at each time point throughout batch cultivation in GMM. The mean and standard deviation are shown. Significant differences (*p < 0.05, Student’s t-test) between the *tsa1*Δ mutant and the parental strain at each specific time point are shown. **B)** Intracellular glycogen levels assessment via iodine staining. 5-μL aliquots of serially diluted GMM cultures (from 10^−1^ to 10^−4^) in GMM were spotted onto solid YPD, SC, or SC-Ethanol (SCEtOH) media. After 2 days of incubation at 30°C, plates were exposed to iodine vapor for 2.5 minutes and imaged immediately. **C)** Chronological life span (CLS) of wild-type and *tsa1*Δ strains in GMM. Cell viability was monitored over time by colony-forming units (CFUs) to determine survival kinetics in four independent biological replicates. The mean and standard deviation are shown. Significant differences (*p < 0.05, Student’s t-test) between the *tsa1*Δ mutants and their parental strains at each time point are shown. **D)** Representative microscopy images of propidium iodide-stained W303 and *tsa1*Δ cells. Left: merged fluorescence and phase contrast; Right: differential interference contrast (DIC) images. Scale bar = 10 μm. **E)** Quantitative microscopy measurements of mean cell propidium iodide fluorescence (per pixel) as a function of apparent cell radius (in pixels, scale: 1 pixel = 0.32 μm) for W303 and *tsa1*Δ cells. Each individual point corresponds to a single cell. The horizontal dashed line indicates the fluorescence threshold (0.05) used to classify cells as dead (>0.05) or alive (≤0.05) (see S1 Fig A). Solid circles represent the average size and fluorescence within five size bins, delineated by the solid line segments. Bins were chosen to contain the same number of cells. The standard error of the mean is smaller than the size of the circles in all cases. At the top of the plot, pie charts indicate the percentage of dead (dark color) and alive (light color) cells in each bin, with the dead percentage also shown numerically.

The accumulation of trehalose and glycogen is critical for the establishment of quiescence and ensures long-term survival during starvation [43,46]. Moreover, Tsa1 is a known modulator of longevity, particularly under caloric restriction [17]. To determine if the metabolic defects observed in the *tsa1*Δ mutant translate into reduced fitness, we assayed the chronological lifespan (CLS) of the wild-type and *tsa1*Δ strains under our experimental conditions. Consistent with the impaired transition to stationary phase and reduced storage carbohydrate levels, the viability of the *tsa1*Δ mutant began to decline markedly since day 5 of the longevity assay (**Fig 2C**). By day 18, the mutant population exhibited less than 10% viability, whereas the wild-type strain maintained a higher survival until day 30.

Based on these results, we investigated whether this premature mortality in the *tsa1*Δ mutant was driven by a systemic failure in cellular biochemistry or by the compromised viability of the small-cell subpopulation that emerges following glucose exhaustion (**Fig 1D**). To address this, we analyzed cell viability at the single-cell level using propidium iodide (PI) staining on day 13, a time point when both wild-type and *tsa1*Δ strains began to show compromised survival (**Fig 2C and 2D**). Microscopy data confirmed the accelerated cell death of the mutant strain (**Fig 2E**). Next, we coupled fluorescence with morphological parameters using a machine-learning image-analysis pipeline. We observed a slight tendency for smaller cells to exhibit higher mortality in both strains. However, the *tsa1*Δ mutant displayed a 2- to 3-fold increase in mortality compared to the wild-type strain across all cell sizes (**Fig 2E and S1 Fig**). These data suggest that the premature collapse of the *tsa1*Δ population is driven by a multifactorial combination of events. This includes an aberrant entry into stationary phase with insufficient storage of protective carbohydrates, the emergence of a fragile subpopulation of small cells with increased mortality, and the well-described accumulation of oxidative damage in the absence of the major cytosolic peroxiredoxin [14]. Thus, Tsa1 integrates metabolic reprogramming during the diauxic shift with the maintenance of cellular integrity to promote survival during prolonged nutrient deprivation.

### 3.3. Tsa1 modulates the activation state of PKA in response to glucose exhaustion

The molecular transitions required for diauxic shift and stationary phase entry are governed by multiple regulators, many of them orchestrated by the attenuation of Protein Kinase A (PKA) signaling [47–49]. Given that Tsa1 physically interacts with Bcy1 (the regulatory subunit) and Tpk1 (a catalytic subunit) of PKA [18,19], we asked if the defects observed in the *tsa1*Δ mutant are mediated by PKA. We thus investigated whether Tsa1 is required for the proper modulation of PKA activity during glucose depletion.

PKA inhibition upon glucose exhaustion triggers the dephosphorylation and nuclear translocation of the stress-responsive transcription factor Msn2. To analyze the nucleocytoplasmic shuttling of Msn2, we monitored wild-type and *tsa1*Δ cells harboring the Msn2-NeonGreen reporter using live-cell microscopy. Cells were cultured in a microfluidic system with a continuous flow of fresh glucose minimal media (GMM) for 7 h, followed by flow cessation to allow for natural glucose depletion, thereby simulating the diauxic shift in batch cultures. During glucose abundance, Msn2 remained cytoplasmic in both the wild-type and *tsa1*Δ strains, consistent with active PKA signaling (**Fig 3A**). However, following glucose depletion (2 h 40 min post-flow stops), the wild-type cells exhibited near-absolute Msn2 nuclear accumulation, whereas this translocation was significantly reduced in the *tsa1*Δ mutant (**Fig 3A and 3B**).

**Figure 3.**
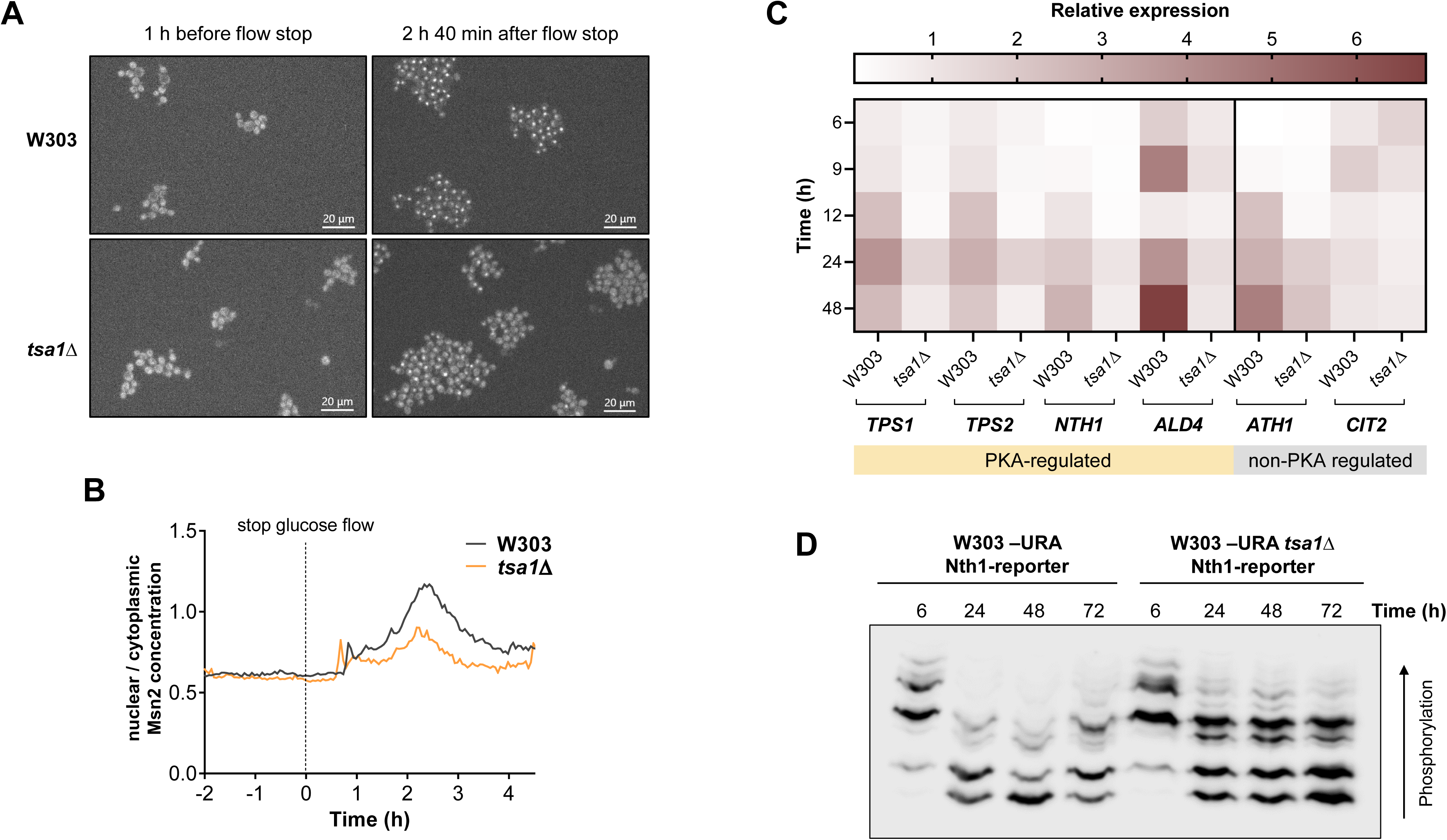
Loss of *TSA1* results in aberrant PKA activation during the fermentative-to-respiratory transition. **A)** Real-time monitoring of Msn2-NeonGreen subcellular localization via live-cell imaging. Wild-type and *tsa1*Δ mutant strains were cultured in a microfluidic system in GMM, where glucose exhaustion was induced by arresting medium flow to simulate the diauxic shift. Representative images show Msn2-NeonGreen distribution under steady-state glucose flow (1h before flow stop) and 2 h 40 min post-flow arrest. **B)** Quantitative analysis of the nuclear-to-cytoplasmic Msn2-NeonGreen fluorescence ratio. **C)** Time-course expression analysis of *TPS1*, *TPS2*, *NTH1*, *ALD4*, *ATH1* and *CIT2* genes by RT-qPCR. **D)** Phosphorylation profile of a 3xFLAG-tagged Nth1 reporter via Phos-tag SDS-PAGE and Western blot. Strains expressing the reporter integrated at the *URA3* locus were grown in GMM for 72 h, and the phosphorylation status was analyzed at the indicated time points. The experiments were carried out in triplicate. The mean and standard deviation are shown. Significant differences (*p < 0.05, Student’s t-test) between the *tsa1*Δ mutants and their parental strains at each time point are shown.

To confirm that Tsa1 is implicated in the coordination of the Msn2-dependent transcriptional program, we analyzed the expression of several Msn2 target genes containing Stress Response Elements (STREs). We primarily focused on genes involved in trehalose metabolism, such as *TPS1* and *TPS2* (encoding the trehalose synthase complex) and the neutral trehalase *NTH1*, but also including the aldehyde dehydrogenase *ALD4* as a STRE-regulated control not involved in trehalose biosynthesis. As negative controls (genes lacking STREs), we monitored the acid trehalase *ATH1* and the citrate synthase *CIT2*. Gene expression was assessed by qPCR across key growth phases: fermentative growth (6 h), the diauxic shift (9–12 h), ethanol respiration (24 h), and entry into stationary phase (48–72 h) (**Fig 3C**). As expected, all STRE-containing genes were robustly induced in the wild-type upon glucose depletion [5,6,50]. However, this induction was significantly attenuated in the *tsa1*Δ mutant, consistent with incomplete Msn2 activation (**Fig 3A and 3B**). Interestingly, the expression of *ATH1* (lacking a promoter STRE) was also reduced in the mutant starting at 12 h, suggesting that Tsa1 might exert an additional, more direct influence on trehalose metabolism regulation. In contrast, *CIT2* expression remained comparable between strains, confirming that Tsa1 does not unspecifically impact global transcription. Collectively, these results demonstrate that Tsa1 is required for the proper activation of the Msn2/4-dependent transcriptional program during the metabolic transition.

To further investigate the Tsa1-dependent PKA regulation during glucose depletion, we studied the phosphorylation of the neutral trehalase Nth1, a well-characterized direct target of PKA [51–53]. For this approach, we used a recently described Nth1-reporter comprising the N-terminal regulatory tail of Nth1 (the first 95 amino acids), which contains all known PKA phosphosites [24]. This peptide was fused to a 3xFLAG tag for detection and expressed under the control of the endogenous *NTH1* promoter. The reduced size of this reporter enables superior resolution of distinct phosphoisoforms via Phos-tag SDS-PAGE compared to the full-length protein. The phosphorylation dynamics of the Nth1-reporter were assessed across different growth phases (**Fig 3D**). During glucose abundance (6 h), both wild-type and *tsa1*Δ strains displayed the slowest-migrating phosphoisoforms of the Nth1-reporter, reflecting robust PKA-mediated phosphorylation. However, following glucose depletion, the *tsa1*Δ mutant maintained these highly phosphorylated isoforms, albeit in a lower proportion than during fermentative growth; in contrast, these isoforms were markedly reduced or absent in the wild-type. These findings demonstrate that Tsa1-dependent inhibition of PKA also affects the phosphorylation status of downstream metabolic enzymes. Collectively, these results establish that Tsa1 contributes to proper PKA inhibition during the diauxic shift and the transition into stationary phase.

### 3.4. Impact of Tsa1 on trehalose metabolism-related enzymes

Given the importance of trehalose accumulation for the transition into stationary phase, we investigated whether Tsa1 influences the abundance or activity of the enzymes responsible for its metabolism. Our analysis focused on Tps1, Tps2, and Nth1, as their transcription was impaired in the *tsa1*Δ mutant via the PKA/Msn2 axis.

Recent proteomic studies identified a physical interaction between Tsa1 and several components of the trehalose synthase complex, including Tps1 and Tps2 [15,16], underscoring the need to determine whether Tsa1 also modulates these biosynthetic enzymes post-translationally. For this purpose, wild-type and *tsa1*Δ mutant cells were grown in GMM under shake-flask conditions, and Tps1 and Tps2 protein levels and enzymatic activity were monitored across growth phases. As shown in **Fig 4A and 4B**, Tps1/2 protein levels increased following glucose depletion at 12 h. Although *TSA1* deletion resulted in a slight increase in the abundance of both proteins, which was most notable in the case of Tps2, this did not translate into higher enzymatic activity. After measuring trehalose-6-phosphate synthase (TPS) activity, we observed an increase after 12 h, coinciding with the diauxic shift and the onset of respiratory growth, before decreasing at the end of the stationary phase (**Fig 4D**). However, no significant differences in TPS activity were detected between the wild-type and *tsa1*Δ mutant. These results indicate that Tsa1 primarily influences the transcriptional regulation of trehalose synthesis rather than modulating the stability or catalytic activity of the Tps1/2 complex.

**Figure 4.**
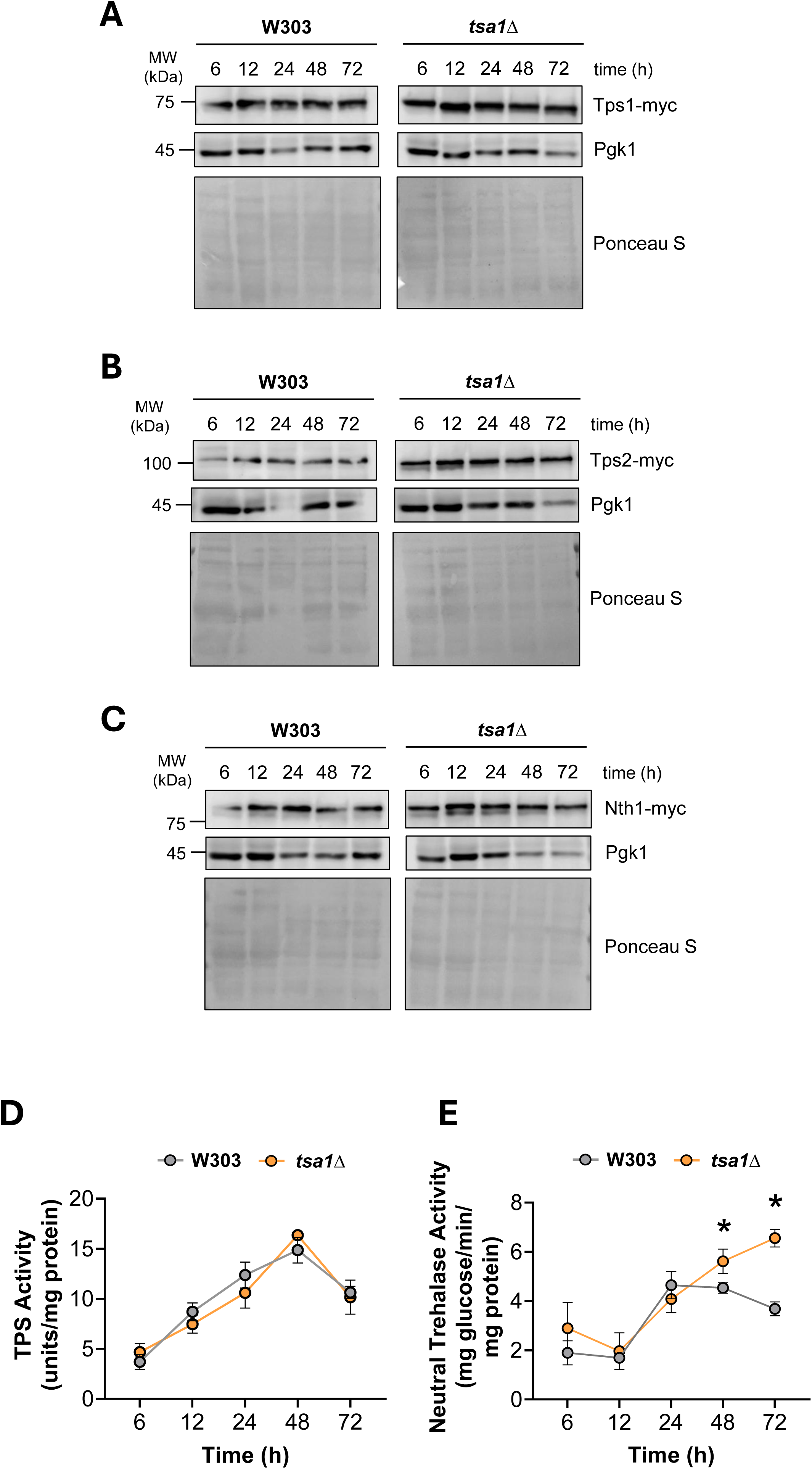
Influence of Tsa1 on protein levels and enzymatic activities of trehalose metabolism-related enzymes. Wild-type and *tsa1*Δ mutant cells expressing 13myc-tagged Tps1, Tps2 and Nth1 were used to monitor protein levels of **A)** Tps1, **B)** Tps2 and **C)** Nth1 throughout growth. **D)** Trehalose-6-phosphate synthase (TPS) and **E)** neutral trehalase activities were measured in untagged W303 and W303 *tsa1*Δ cultures. All strains were grown in GMM for 72 h under shake-flask conditions. The experiments were carried out in triplicate, and the mean and standard deviation are provided. Significant differences (*p < 0.05, Student’s t-test) between the *tsa1*Δ mutant and the wild-type strains at each time point are shown.

These observations prompted us to investigate whether Tsa1 influences trehalose degradation by modulating trehalase activity. Although both neutral (Nth1) and acid (Ath1) trehalases contribute to this process, only Nth1 is regulated by PKA at both transcriptional and post-translational levels. Therefore, we focused our analysis on Nth1 to better understand how Tsa1 might influence trehalose turnover through PKA-dependent regulation. Our temporal analysis revealed that *TSA1* deletion led to a slight increase in Nth1 protein levels, which was particularly evident during the stationary phase (48–72 h) (**Fig 4C**). Nth1 enzymatic activity increased progressively in both strains, peaking at 24 h of growth (**Fig 4E**). At this time point, in the wild-type strain, the increased protein abundance may compensate for the declining proportion of phosphorylated Nth1 (**Fig 3D**), maintaining overall activity. However, while Nth1 activity declined in the wild-type during stationary phase due to the depletion of its active phosphoisoforms, the *tsa1*Δ mutant maintained elevated activity. This sustained enzymatic activity in the mutant correlates with the persistent PKA-mediated Nth1 phosphorylation (**Fig 3D**) and provides a mechanistic explanation for the reduced trehalose accumulation in these cells (**Fig 2A**). Parallel analysis showed that *tsa1*Δ cells exhibited decreased Ath1 activity during growth (**S2 Fig**), consistent with the reduced *ATH1* expression (**Fig 3C**). However, at the end of the stationary phase, Ath1 activity increased in the mutant, suggesting a late-stage induction of vacuolar or periplasmic trehalose mobilization (**S2 Fig**).

### 3.5. Tsa1 influences the remodeling of central carbon metabolism

Beyond its role in PKA regulation, Tsa1 has been reported to interact with several enzymes involved in carbohydrate metabolism [15,16,54]. Based on this evidence, we sought to investigate the broader role of this peroxiredoxin in the remodeling of central carbon metabolism after glucose depletion. To this end, we implemented dynamic Flux Balance Analysis (dFBA), a computational approach that estimates intracellular metabolic fluxes in time-course experiments by integrating experimental growth and exometabolite data with genome-scale metabolic models (GEMs).

First, several extracellular metabolite concentrations were measured at defined time points from GMM shake-flask cultures of wild-type and *tsa1*Δ strains (**S3 Fig**). We then adapted the model developed by Moimenta et al. (2025) [26], applying the dFBA to quantify changes in the net accumulation of extracellular metabolites per unit biomass. Overall, the dynamic model robustly explained the experimental data (**S3 Fig, solid lines**), with goodness-of-fit (R^2^) values approaching 1 for most metabolites.

To analyze intracellular metabolic flux distributions between strains during the metabolic shift from fermentation to respiration, we presented flux ratios normalized to the primary carbon source: glucose for fermentative growth (**Fig 5A**) and ethanol for respiratory growth following the diauxic shift (**Fig 5B**). This source-normalization approach enabled clear resolution of metabolic state-dependent differences between strains. The absolute normalized flux values for each reaction during growth on glucose and ethanol are detailed in **S3 Table** and **S4 Table**, respectively.

**Figure 5.**
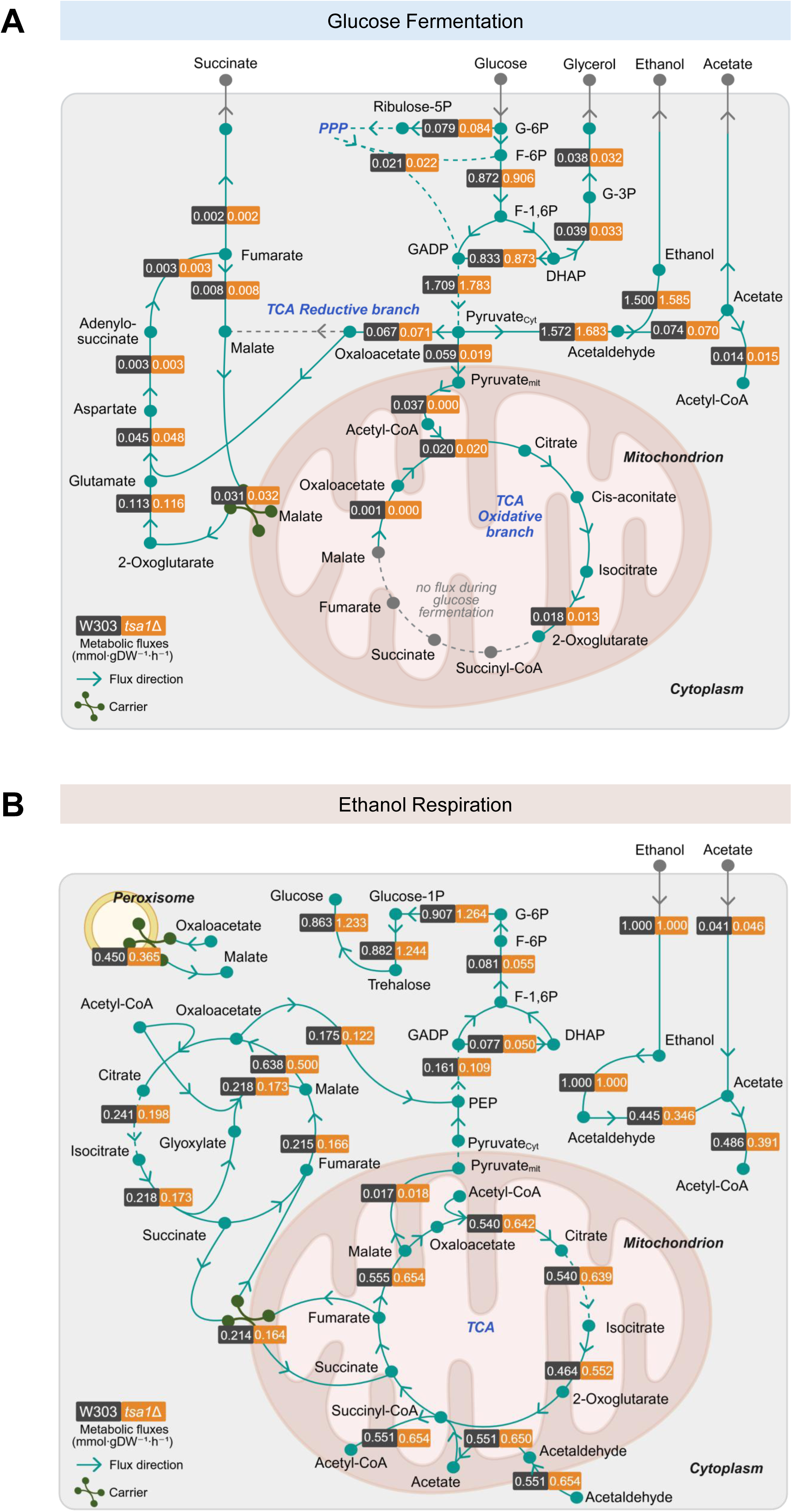
Dynamic flux balance analysis. **(A)** Predicted intracellular dynamic flux scores normalized by growth and glucose flux during the fermentative phase. **(B)** Predicted intracellular dynamic flux scores normalized by growth and ethanol flux during the post-diauxic phase. The model analyses the dynamic distribution of metabolic fluxes in the wild-type and *tsa1*Δ mutant strains during growth in GMM.

The flux analysis revealed subtle but distinct responses to *TSA1* deletion during fermentative growth (**Fig 5A**). As expected, most of the metabolic flux was directed toward ethanol production via alcoholic fermentation. Minor differences were observed in acetaldehyde oxidation, which, once oxidized, did not substantially affect extracellular acetate levels (**S3 Fig**). Notably, the wild-type strain exhibited some pyruvate transport into the mitochondria, where pyruvate dehydrogenase (PDH) converts it into acetyl-CoA to feed the tricarboxylic acid (TCA) cycle, even though PDH remains partially repressed under glucose-rich conditions due to glucose-repression signaling [55]. In the *tsa1*Δ mutant, PDH seemed to be more strongly repressed by glucose, resulting in negligible flux through this reaction. Interestingly, the mutant displayed increased mitochondrial transport of acetaldehyde (**S3 Table**, r_1632) and reduced repression of mitochondrial aldehyde dehydrogenase (**S3 Table**, r_0174), leading to enhanced mitochondrial acetate and acetyl-CoA synthesis (**S3 Table**, r_0113). This compensation enabled a TCA oxidative flux comparable to that of the wild-type strain. Altogether, during growth on glucose as the sole carbon source, Tsa1 appears to primarily influence pyruvate fate allocation flux.

We next examined the impact of *TSA1* deletion on intracellular metabolic fluxes following the diauxic shift, when ethanol became the main carbon source (**Fig 5B, S4 Table**). Consistent with established metabolic remodeling during respiratory growth, both strains exhibited a downregulation of alcoholic fermentation and pentose phosphate pathway, coupled with increased fluxes through TCA and glyoxylate cycle [47]. In both strains, glucose repression was relieved, demonstrated by increased flux through Ald4 for cytosolic acetate synthesis, which was subsequently converted into acetyl-CoA and succinyl-CoA to fuel the TCA cycle. Cytosolic acetyl-CoA was also redirected toward the TCA cycle, which was fully activated under these conditions, as evidenced by the flux through succinate dehydrogenase.

However, the *tsa1*Δ mutant displayed a distinct metabolic redirection, characterized by increased flux through the TCA cycle at the expense of reduced gluconeogenic and glyoxylate cycle fluxes. Given that Tsa1 contributes to the proper downregulation of PKA and pyruvate kinase activities after glucose depletion (**Fig 3**) [54,56], its absence was expected to favour glycolytic over gluconeogenic flow. Consistent with this, the *tsa1*Δ mutant exhibited reduced flux through the reaction catalyzed by the gluconeogenic enzyme phosphoenolpyruvate carboxykinase (PEPCK). In agreement with these findings, our endometabolomic analysis at 24 h of growth showed that lower glycolytic intermediates, such as 2/3-phosphoglycerate and phosphoenolpyruvate (PEP), were significantly decreased in the *tsa1*Δ mutant (**S4 Fig**).

Interestingly, despite the overall reduction in lower glycolytic intermediates, the *tsa1*Δ mutant exhibited higher levels of upper glycolytic metabolites, mainly glucose-6-phosphate (G6P) (**S4 Fig**). The increased G6P availability drives the dFBA to estimate an increased flux toward trehalose synthesis in the mutant (**Fig 5B**), which directly contradicts our experimental evidence of reduced intracellular trehalose accumulation (**Fig 2A**) or unaffected TPS activity (**Fig 4D**). This discrepancy, combined with the elevated Nth1 activity in the *tsa1*Δ mutant in stationary phase (**Fig 4E**), suggests the existence of a trehalose futile cycle, where accelerated degradation prevents the net accumulation of the disaccharide.

Overall, Tsa1 appears to exert only a minor influence during growth on glucose, when PKA activity is high, but has a pronounced influence when glucose is depleted and PKA activity must be downregulated. These flux analyses suggest that Tsa1 regulates metabolism both through the modulation of PKA signaling and via direct enzymatic interactions.

### 3.6. Effect of *TSA1* deletion on stationary phase exit

Since Tsa1 supports the cellular response to glucose depletion and proper entry into stationary phase, we investigated how *tsa1*Δ mutants respond to the exit from the stationary phase. In yeast, resuming growth upon glucose replenishment triggers rapid cAMP accumulation and the subsequent activation of PKA [57,58].

To assess this response, stationary phase cultures (72 h) were diluted 1:5 in fresh GMM. We first monitored the ability of cells to resume growth and division by scoring the budding index (**Fig 6A**). The *tsa1*Δ mutant showed delayed and incomplete budding over the two hours following glucose replenishment. Consistent with this delayed recovery, both glucose uptake and ethanol production were reduced in the *tsa1*Δ mutant during this initial period (**Fig 6B and 6C**). Furthermore, the mobilization of trehalose, which is characteristic of wild-type cells resuming growth, was significantly impaired in the mutant (**Fig 6D**).

**Figure 6.**
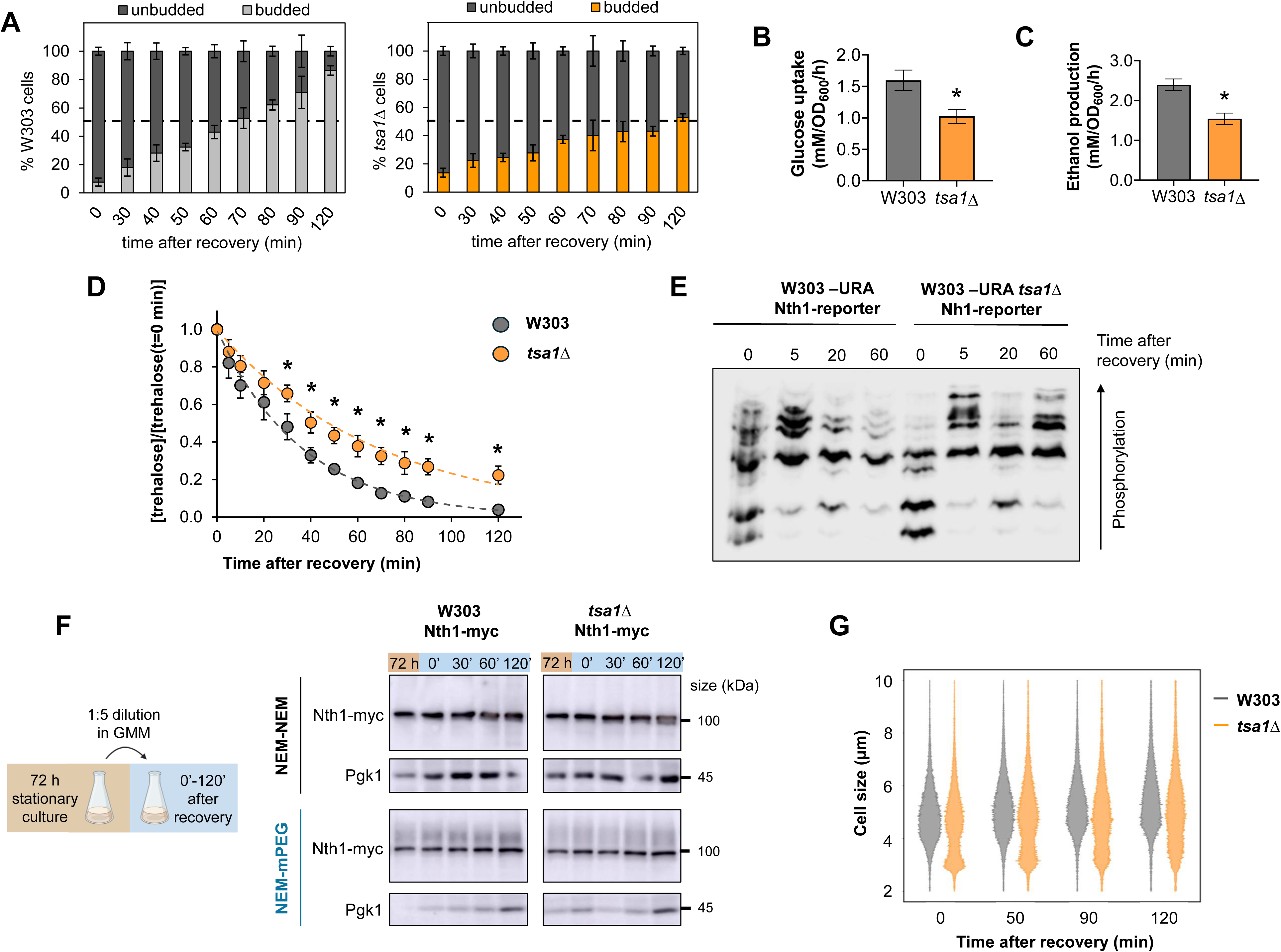
Tsa1 is required for the efficient coordination of metabolic and proliferative restart upon glucose repletion. Stationary-phase cultures (72 h) of the wild-type and the *tsa1*Δ mutant strains were diluted 1:5 into fresh GMM and monitored for 120 min in shake flasks. **A)** Proliferation dynamics of the wild-type strain and the *tsa1*Δ mutant represented by the percentage of budded and unbudded cells. Quantification was performed via microscopic examination (n ≈ 200 cells per replicate) using a manual cell counter. **B)** Glucose consumption and **C)** ethanol production profiles during the initial 120 min of metabolic recovery. **D**) Intracellular trehalose mobilization kinetics in glucose-induced cells after the stationary growth phase. **E)** Phosphorylation profiling of the 3xFLAG-tagged Nth1 reporter via Phos-tag Western blot. The construct was integrated into the *URA3* locus in W303 -URA and W303 -URA *tsa1*Δ cells. **F)** Redox status of Nth1 after glucose-induced restart, analyzed via NEM-mPEG alkylation assay in strains expressing 13myc-tagged Nth1. The oxidation status of Nth1 after glucose-metabolic restart was conducted using the NEM-mPEG assay from wild-type and *tsa1*Δ mutant cells expressing 13myc-tagged Nth1. **G)** Cell size distribution of wild-type and *tsa1*Δ populations after glucose-metabolic restart. The data represent the mean and standard deviation of three independent biological replicates. Significant differences (\**p* < 0.05, Student’s *t*-test) between the *tsa1*Δ mutant and the wild-type strain at each time point are shown.

Next, we wondered whether these delayed recovery and incomplete trehalose mobilization were a result of impaired PKA activity, compromised redox homeostasis, or a downstream consequence of the improper entry into the stationary phase. To test the first hypothesis, we monitored the phosphorylation of the trehalase Nth1 (**Fig 6E**), which is known to be rapidly phosphorylated at multiple sites upon glucose replenishment. However, no differences in Nth1 phosphorylation were observed between wild-type and *tsa1*Δ strains, indicating that PKA activation remains functional. These results suggest that the phosphorylation status of Nth1 is not the primary cause of the impaired trehalose mobilization observed in the mutant.

Given that the deletion of the main cytosolic peroxiredoxin may lead to increased oxidative damage, affecting the redox state and activity of multiple proteins, we also examined the oxidation status of Nth1. We used a differential labelling methodology for detecting reduced and oxidized cysteine (Cys) residues using N-ethylmaleimide (NEM) and methoxy polyethylene glycol (mPEG) 5000, respectively. The mPEG reagent specifically binds to oxidized Cys residues, resulting in reduced electrophoretic mobility proportional to the number of modified residues. Our analysis revealed that Nth1 protein levels and oxidation status in the *tsa1*Δ mutant strain remained unaffected following glucose replenishment (**Fig 6F**), suggesting this is also not the main reason for delayed recovery.

Consequently, we hypothesized that the delayed exit from stationary phase was a downstream consequence of the improper entry observed above, specifically focusing on the behavior of the small-cell subpopulation. The analysis of the cell size distribution in the two hours following glucose recovery revealed that the small-sized cell population gradually shifted toward larger sizes (**Fig 6G**). However, by 120 minutes, it had not yet reached the mean of the wild-type population. This suggests that the small-cell subpopulation in *tsa1*Δ mutant struggles to resume growth and requires additional time to reach the critical size necessary for division.

These results provide evidence that Tsa1 does not directly regulate the response to glucose replenishment but instead ensures an ordered entry into stationary phase. A failure to properly enter to stationary state in the absence of Tsa1 results in a subpopulation unable to recover rapidly, thereby impacting the overall fitness of the population.

## 4. Discussion

In the model yeast *S. cerevisiae*, the transition from fermentation to respiration and the subsequent entry into stationary phase represents one of the most drastic metabolic shifts in its cellular life cycle [47,59]. During this transition, nutrient signaling pathways play a pivotal role by eliciting molecular responses that adapt metabolism to the changing environment, achieving a perfectly orchestrated balance between the attenuation of growth-promoting pathways and the activation of survival programs [2,60]. Understanding the effectors that sense nutrient-derived signals and modulate these pathways is crucial for elucidating the molecular mechanisms underlying cellular adaptation. Over the past few years, the peroxiredoxin Tsa1 has emerged as a metabolic regulator that integrates redox signals with cellular physiology. Indeed, recent studies demonstrate its ability to interact with numerous metabolic proteins and modulate pathway activation or metabolite production [15,16,61]. Based on this evidence, in this work, we investigated the contribution of Tsa1 to the metabolic and physiological transition triggered by glucose exhaustion. Our findings identify this peroxiredoxin as a critical integrative node that orchestrates the shift to a quiescent state by modulating both PKA signaling and central carbon metabolism.

Deletion of the major cytosolic peroxiredoxin does not significantly affect fermentative metabolism, as the glucose consumption profile is not impaired (**Fig 1B and 5A**). However, the *tsa1*Δ mutant exhibits a delay in ethanol consumption (**Fig 1B**) and impaired trehalose accumulation (**Fig 2A**), revealing a potential role of Tsa1 in coordinating the metabolic reprogramming required for respiratory growth, post-diauxic survival, and entry into a stable quiescent state. During the metabolic shift to respiratory growth, glucose exhaustion leads to reduced cAMP levels and the subsequent attenuation of PKA activity [62]. A central finding of this work is that the peroxiredoxin Tsa1 is required for this timely downregulation of PKA signaling. We provide three distinct lines of evidence to support this regulation: first, subcellular localization analysis revealed significantly reduced nuclear accumulation of the stress-responsive transcription factor Msn2 in the *tsa1*Δ mutant following glucose depletion (**Fig 3A and 3B**); second, the mutant maintained highly phosphorylated isoforms of the Nth1-reporter into stationary phase (**Fig 3D**); and third, our transcriptional analysis showed an attenuated induction of different STRE-regulated genes (**Fig 3C**). Additionally, our dFBA model predicts reduced metabolic flux through the glyoxylate and gluconeogenic pathways in the *tsa1*Δ mutant after glucose depletion (**Fig 5B**). We hypothesize that this Tsa1-dependent PKA dysregulation negatively impacts the Snf1 kinase, which orchestrates the expression of genes required for non-fermentable carbon utilization, comprising those involved in the glyoxylate cycle and gluconeogenesis [63–65]. While a direct connection between Tsa1 and Snf1 remains to be fully elucidated, increased PKA activity is well established to antagonize Snf1 functionality, potentially through the phosphorylation and inhibition of the Snf1-activating kinase Sak1 [66].

However, a more direct role cannot be ruled out and may work in parallel. Key metabolic enzymes may be oxidized when Tsa1 is absent, although that was not seen in the only enzyme that was tested, Nth1 (**Fig 6F**). Beyond PKA modulation, Tsa1 physically interacts with several glycolytic enzymes [15,16]. For instance, Tsa1 contributes to the inhibition of pyruvate kinase (Cdc19) activity through direct interaction [54,56]. So, in the *tsa1*Δ mutant, the absence of this regulation favours glycolytic over gluconeogenic fluxes after the diauxic shift. This is consistent with the significant decrease we observed in lower glycolytic intermediates, such as PEP and 2/3-PG (**S4 Fig**), and the reduced flux through the key gluconeogenic enzyme PEPCK (**Fig 5B**). This interference in signaling and enzymatic control could lead to a state of metabolic jamming following glucose depletion, as evidenced by our endometabolomic analysis. While the *tsa1*Δ mutant exhibits a decrease in lower glycolytic intermediates, it simultaneously presents a significant accumulation of the upper intermediate G6P (**S4 Fig**). This metabolic profile generates a striking paradox: while the dFBA estimates an increased flux toward trehalose synthesis driven by high precursor availability (**Fig 5B**), our experimental data show drastically reduced levels of the disaccharide (**Fig 2A**). As the mutant maintains elevated Nth1 trehalase activity during stationary phase due to persistent PKA activation (**Fig 4E**), we propose the existence of an operative, ATP-consuming trehalose futile cycle [67]. Such futile cycling has been previously described as a strategy to maintain metabolic stability and prevent glycolytic collapse in trehalose-defective cells [68], and here may be a “pathological” condition due to incomplete induction of the quiescence program. Additionally, these high levels of G6P may also reflect a lower ability of the mutant to channel it into glycogen synthesis (**Fig 2B**).

The broader consequences of this metabolic and physiological failure during entry into stationary phase also affect long-term viability. Both Tsa1 and trehalose are critical for maintaining cellular integrity during prolonged cultivation. Tsa1 is considered a gerontogene, contributing to the extension of chronological lifespan under caloric restriction by attenuating the Ras/cAMP/PKA pathway [18,69]. Concurrently, trehalose acts as a protectant against oxidative stress and ensures proper protein-folding dynamics throughout the lifespan [70,71]. Therefore, the deletion of *TSA1* and the inability to accumulate proper trehalose levels deprive the cell of critical mechanisms to cope with oxidative stress and the accumulation of protein aggregates in chronologically aged cells during stationary phase. These defects lead to the premature collapse of the population and a significant loss of chronological viability (**Fig 2C**).

Additionally, the disordered entry into stationary phase experienced by the *tsa1*Δ mutant triggers the emergence of a small-cell subpopulation characterized by higher mortality (**Fig 2F and S1 Fig**). This phenotypic heterogeneity could denote the coexistence of distinct metabolic subpopulations during nutrient transitions, where a failure to properly start or stop specific pathways yields nongrowing or compromised cells. But this finding also could suggest that Tsa1 may mediate the relationship between trehalose accumulation, cell size, and cell cycle progression. While Tsa1 is known to synchronize the yeast metabolic cycle with cell division through redox regulation (51), trehalose accumulation is also intrinsically linked to the regulation of cell size at the end of the growth phase [73]. In the *tsa1*Δ mutant, diminished trehalose accumulation appears to redirect carbohydrates toward an additional, ill-timed round of division, resulting in higher cell titters (**Fig 1C**) but producing daughter cells that fail to attain normal size (**Fig 1D, 1E**). Moreover, the *tsa1*Δ mutant failed to properly resume cell cycle progression upon glucose replenishment (**Fig 6A**). This defect could stem from impaired metabolic reprogramming back to fermentative growth, as the mutant failed to complete intracellular trehalose mobilization (**Fig 6D**). Since the liquidation of stored carbohydrates is coordinated by cyclin-dependent kinase (CDK) activity, which fine-tunes the final cell division to match nutrient availability, this failure suggests a significant disruption in the coupling between metabolism and the cell cycle [42,73,74].

Overall, this study provides evidence of the intricate interplay between metabolism, stress response, and cell cycle regulation, positioning the multifunctional peroxiredoxin Tsa1 as an integrative node. While Tsa1 is a well-established component of the oxidative stress defense system in *S. cerevisiae*, our findings underscore its role as a metabolic regulator, ensuring that the transition to quiescence is a reversible, survival-oriented process.

## Author contributions

**Víctor Garrigós:** conceptualization, investigation, formal analysis, validation, visualization, methodology, writing – original draft, writing – review and editing. **Lisa Dengler:** investigation, formal analysis, validation, visualization, methodology, writing – review and editing. **David Henriques:** software, formal analysis, methodology, writing – review and editing. **Javier Buceta:** software, data curation, formal analysis, visualization, methodology, writing – review and editing. **Emilia Matallana:** supervision, funding acquisition, writing – review and editing. **Cecilia Picazo:** supervision, funding acquisition, writing – review and editing. **Jennifer C Ewald:** conceptualization, formal analysis, methodology, supervision, funding acquisition, writing – review and editing. **Agustín Aranda:** conceptualization, formal analysis, methodology, supervision, funding acquisition, writing – review and editing.

## Acknowledgements

The authors would like to acknowledge the financial support from the Spanish Ministry of Science and Innovation through grant PID2021-122370OB-I00 (co-financed by FEDER funds) to AA and EM, from the Generalitat Valenciana through Emergentes grant number CIGE/2023/32 to CP, from the Cost Action TRANSLACORE CA21154 to CP, and from the Deutsche Forschungsgemeinschaft (DFG) - GRK MOMbrane (Project 327043846) and Project 426546316 to JCE. VG was supported by a predoctoral grant from the University of Valencia (Atracció de Talent Program), and CP was supported by a Maria Zambrano postdoc contract (ZA21-068) from the Spanish Ministry of Universities.

## Supporting information

**S1 Appendix. Extended methods for estimating intracellular metabolic fluxes by dynamic flux balance analysis.**

**S2 Appendix. Code and scripts necessary to reproduce the results from dFBA and microscopy image analysis.**

**S1 Figure. Quantitative microscopy analysis of propidium iodide-stained cells on day 13 of the chronological lifespan assay. A)** Histogram of mean cell fluorescence (per pixel) for W303 and *tsa1*Δ cells. The vertical dashed line indicates the fluorescence threshold (0.05) used to classify cells as dead (>0.05) or alive (≤0.05). This criterion was set based on the minimum fluorescence level observed in both histograms before fluorescence begins to increase. **B)** Histogram of apparent cell radius (in pixels) for W303 and *tsa1*Δ cells, as obtained by quantitative microscopy. The estimated cell radius was calculated assuming a circular cell geometry (see Methods). The overlapping histograms reveal that *tsa1*Δ cells display a smaller size compared to W303 cells, in agreement with data from CASY TT Cell Counter and Analyzer and shown in Fig 1D.

**S2 Figure. Impact of Tsa1 on Ath1 activity during growth in GMM.** Acid trehalase activity was measured in W303 and W303 *tsa1*Δ cultures. Both strains were grown in GMM for 72 h under shake-flask conditions. The experiments were carried out in triplicate, and the mean and standard deviation are provided. Significant differences (*p < 0.05, Student’s t-test) between the *tsa1*Δ mutant and the wild-type strains at each time point are shown.

**S3 Figure. Dynamic flux balance model performance.** Validation of the dFBA model fit to the data. Continuous lines show model predictions. Circle symbols represent the mean of experimentally determined values, while error bars indicate the standard deviation from experimental replicates. Experimental data were obtained from triplicate cultures of each strain grown in GMM for 72 h in shake flasks.

**S4 Figure. Endometabolomic profiling of glycolytic intermediates during the post-diauxic growth phase. A)** Schematic representation of glycolysis and trehalose cycling in *S. cerevisiae*. The diagram illustrates the key enzymatic reactions in upper and lower glycolysis, trehalose biosynthesis and mobilization. Enzymatic steps are indicated in blue, central metabolites in black and energetic cofactors (ATP, ADP, NAD^+^, NADH) in grey. Encircled numbers (1–10) correspond to the specific metabolites quantified in panel B. **B)** Quantification of intracellular glycolytic intermediates in wild-type (WT) and *tsa1*Δ strains after 24 h of growth in GMM. Metabolite abundance is normalized to cell density (OD_600nm_). Data represent the mean and standard deviation of three independent biological replicates. Significant differences (*p < 0.05, Student’s t-test) between the *tsa1*Δ mutant and the wild-type strains are shown. Metabolite abbreviations: G6P, glucose-6-phosphate; UDP-Glc, UDP-glucose; T6P, trehalose-6-phosphate; TREH, trehalose; F6P, fructose-6-phosphate; FBP, fructose-1,6-bisphosphate; DHAP, dihydroxyacetone phosphate; GAP, glyceraldehyde-3-phosphate; G3P, glycerol-3-phosphate; GLY, glycerol; GLYC, glycogen; BPG, 1,3-bisphosphoglycerate; 3PGA, 3-phosphoglycerate; 2PGA, 2-phosphoglycerate; PEP, phosphoenolpyruvate; and PYR, pyruvate.

**S1 Table. List of the strains used in this study.**

**S2 Table. List of the plasmids used in this study.**

**S3 Table. Flux values during fermentative growth in glucose.**

**S4 Table. Flux values during respiratory growth in ethanol.**

