## Supplementary figures and images for "The peroxiredoxin Tsa1 promotes stationary phase entry by suppressing PKA activity"

### Supplementary Fig 1

**Supplementary Figure 1**

A

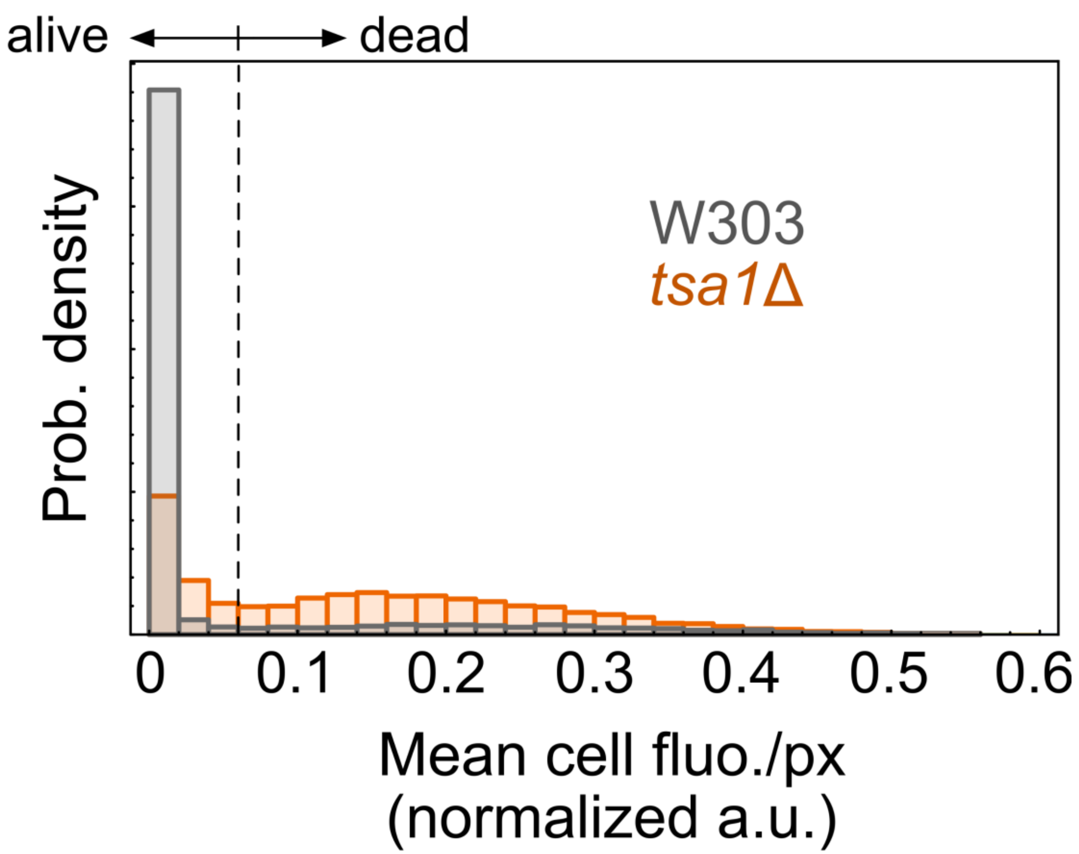


B

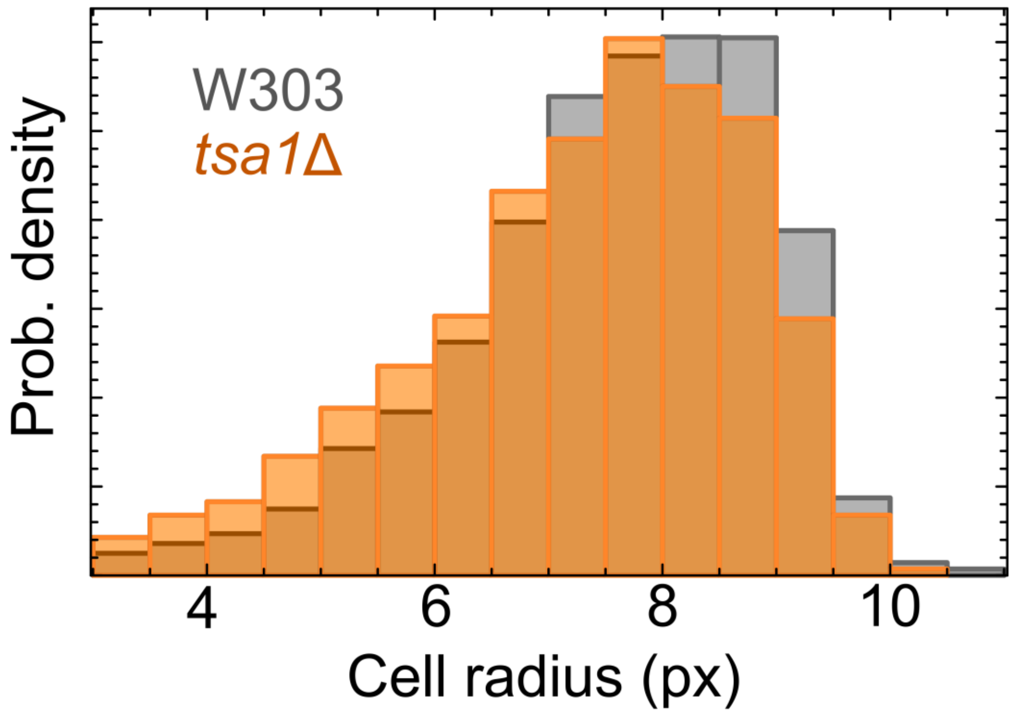

### Supplementary Fig 2

Supplementary
Figure 2

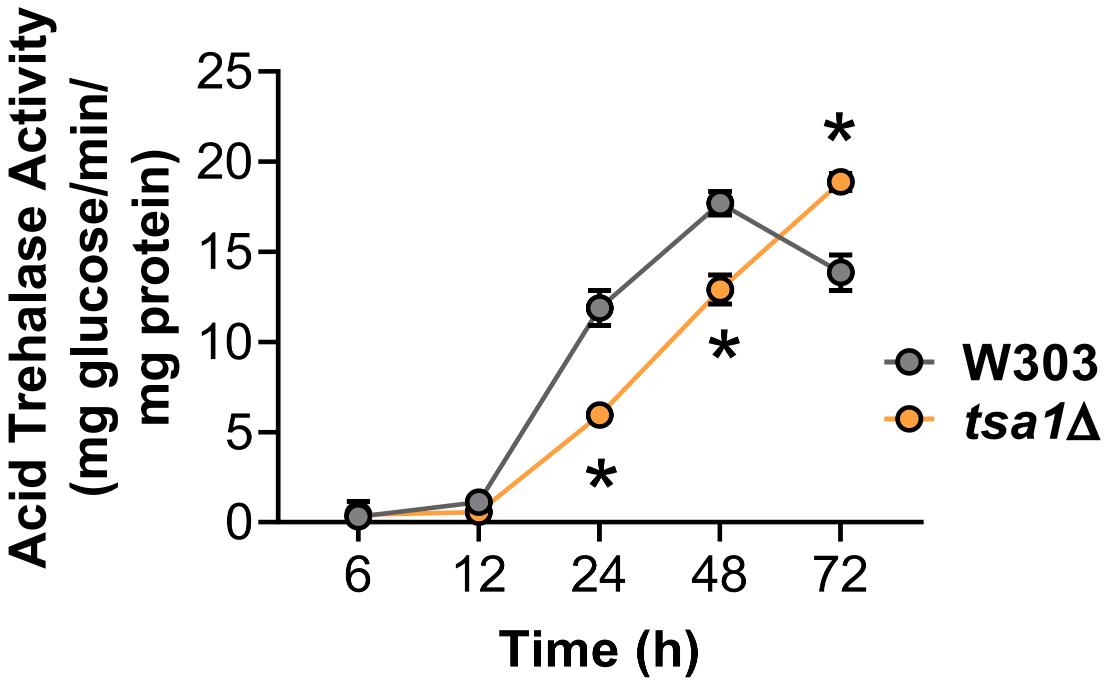

### Supplementary Fig 3

**Supplementary Figure 3**

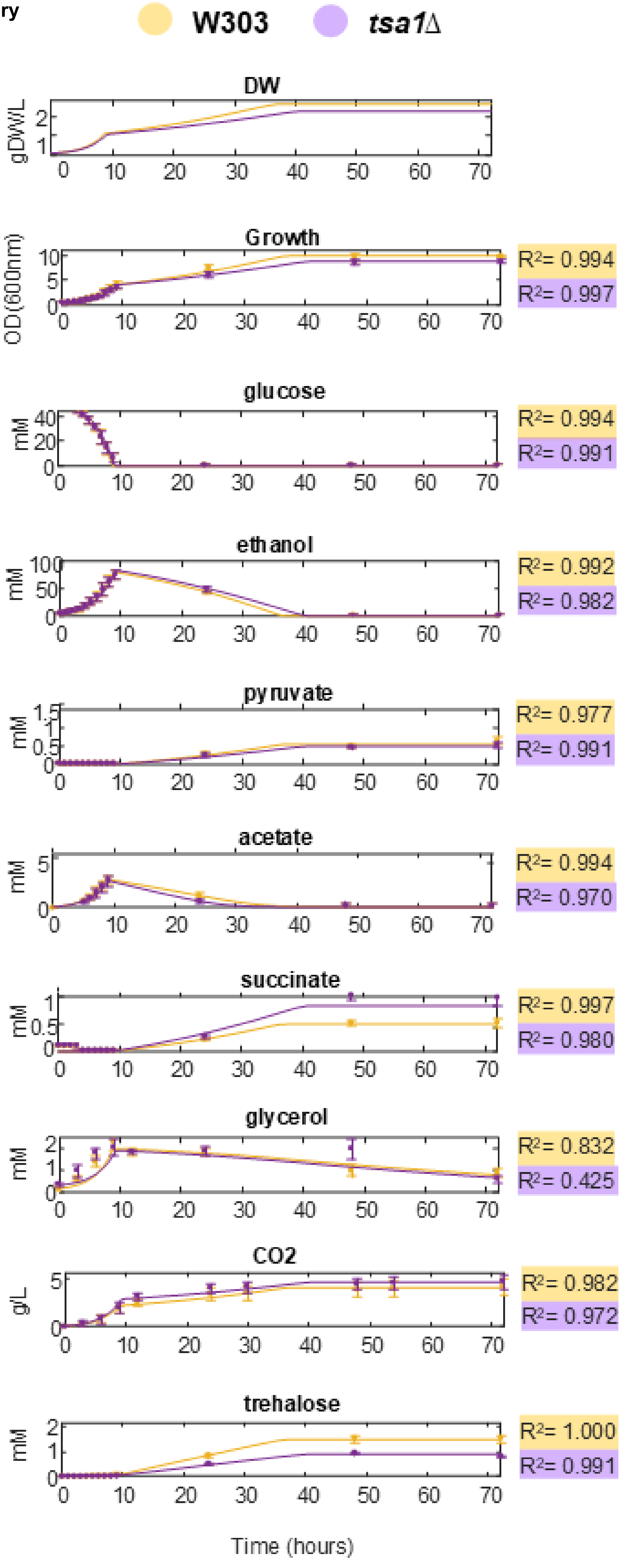

### Supplementary Fig 4

**Supplementary Figure 4**

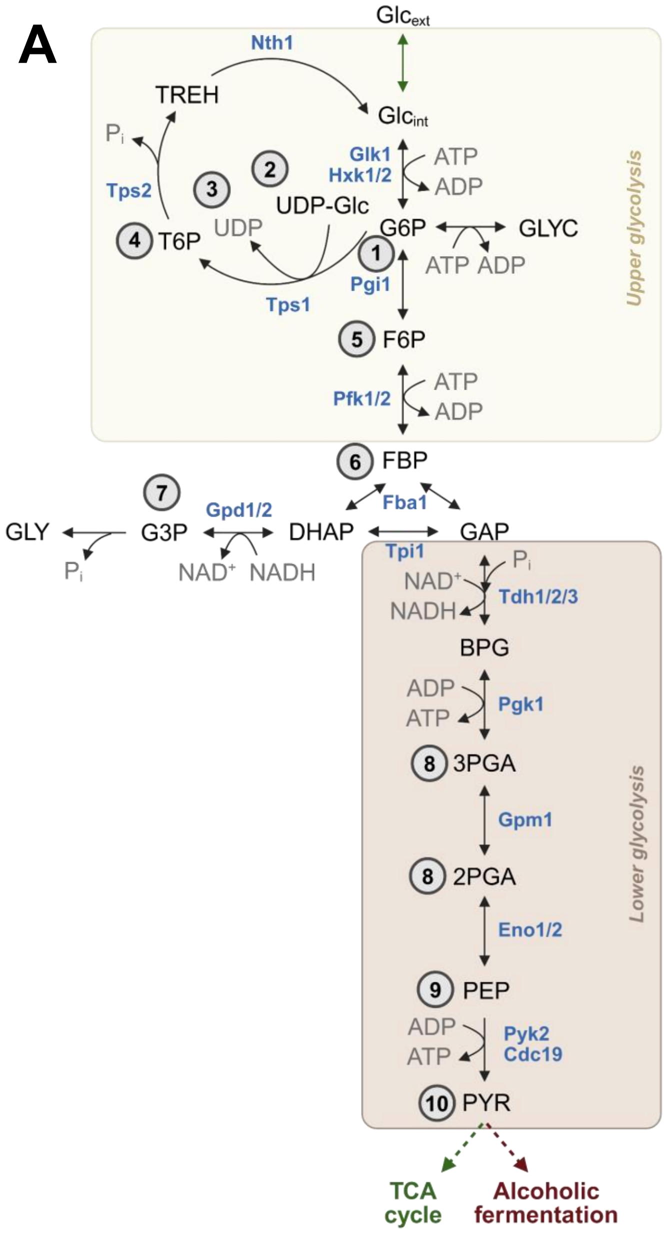


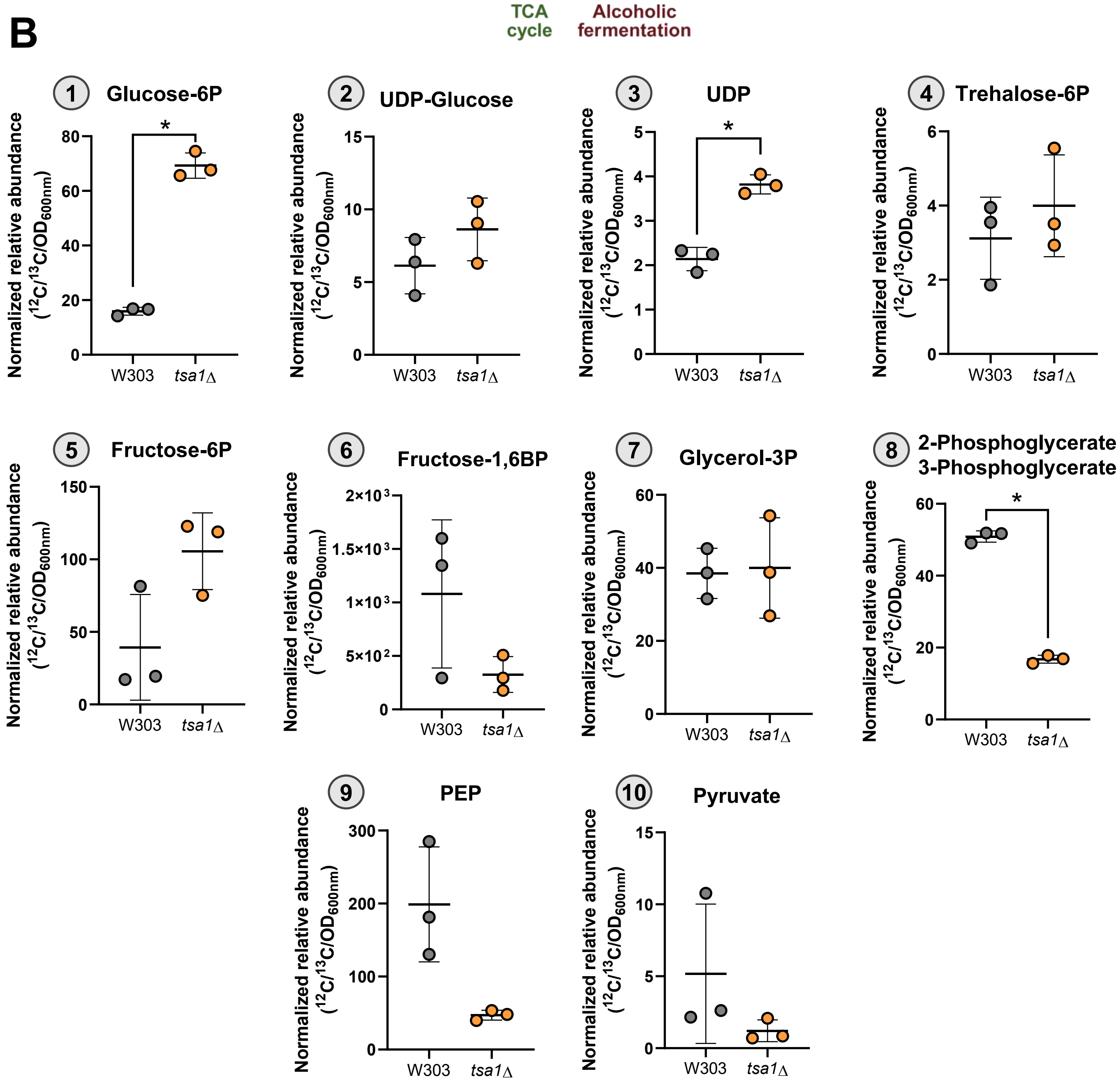
