## Supplementary Table 1 for "The peroxiredoxin Tsa1 promotes stationary phase entry by suppressing PKA activity"

**S1 Table**. List of the strains used in this study.

| Yeast Strain | Genotype | Source |
|---|---|---|
| W303 | MAT a, *ADE*, *LEU*, *HIS*, *TRP*, *URA* | Jennifer C. Ewald laboratory |
| W303 Msn2-mNeongreen Htb2-mCherry | W303 Msn2-mNeongreen-*nat*MX Htb2-mCherry-*kan*MX | This study |
| W303 Nth1-Myc | W303 *NTH1-13myc-kan*MX | This study |
| W303 Tps1-Myc | W303 *TPS1-13myc-kan*MX | This study |
| W303 Tps2-Myc | W303 *TPS2-13myc-kan*MX | This study |
| W303 -URA | MAT a, *ADE*, *LEU*, *HIS*, *TRP* | Jennifer C. Ewald laboratory |
| W303 -URA Nth1-reporter | W303 *-URA, ura3::URA3-NTH1*$^{WT}$*-reporter-3XFLAG* | [1] |
| W303 *tsa1*Δ | W303 *tsa1::loxP* | [2] |
| W303 *tsa1*Δ Msn2-mNeongreen Htb2-mCherry | W303 *tsa1::loxP* Msn2-mNeongreen-*nat*MX Htb2-mCherry-*kan*MX | This study |
| W303 *tsa1*Δ Nth1-Myc | W303 *tsa1::loxP NTH1-13myc-kan*MX | This study |
| W303 *tsa1*Δ Tps1-Myc | W303 *tsa1::loxP TPS1-13myc-kan*MX | This study |
| W303 *tsa1*Δ Tps2-Myc | W303 *tsa1::loxP TPS2-13myc-kan*MX | This study |
| W303 -URA *tsa1*Δ | W303 *-URA tsa1::loxP* | This study |
| W303 -URA *tsa1*Δ Nth1-reporter | W303 *-URA tsa1::loxP, ura3::URA3-NTH1*$^{WT}$*-reporter-3XFLAG* | This study |
