## SupplementaryTable 2 for "The peroxiredoxin Tsa1 promotes stationary phase entry by suppressing PKA activity"

**S2 Table**. List of the plasmids used in this study.

| Plasmid | Description | Source |
|---|---|---|
| pUG6 | Multi-copy plasmid with the *lox*P–*kan*MX–*lox*P disruption module and *kan*MX marker | [1] |
| pFA6a-13Myc-*kan*MX | Plasmid with a *kan*MX marker for adding a C-terminal 13xMyc tag | [2] |
| pYEp351-cre-cyh | Multi-copy plasmid with a *GAL1*-cre recombinase system and Cyh marker | [3] |
| pJE134 | pRS406 plasmid (Stratagene) containing *NTH1* promoter-*NTH1*$^{WT}$ -reporter-3XFLAG (first 95 aa) and *URA* marker | Dr. Jennifer C. Ewald laboratory |
| pYLB9 | Plasmid containing the fluorescent protein mNeongreen and *nat*MX marker | Dr. Jennifer C. Ewald laboratory |
| pCAE011 | Plasmid containing the fluorescent protein mCherry and *kan*MX marker | Dr. Jennifer C. Ewald laboratory |
