## Supplementary Appendix 1 for "The peroxiredoxin Tsa1 promotes stationary phase entry by suppressing PKA activity"

### S1 Appendix. Extended methods for estimating intracellular metabolic fluxes by dynamic flux balance analysis.

In this section, we describe the procedure used to estimate intracellular fluxes using dynamic Flux Balance Analysis (dFBA). FBA is a widely used and accepted method for simulating microbial phenotypes [1–3]. This method was initially developed to estimate intracellular fluxes and predict growth rates under steady-state conditions typically obtained in a chemostat experiment. In this approach, the rationale is that by knowing the substrates entering the cell and the structure of metabolic pathways comprising the metabolism of an organism (metabolic network), it is possible to compute reasonable estimates of how healthy wild-type cells and mutants use these substrates to grow.

When the number of reactions in a metabolic network covers most of the metabolic genes and associated enzymatic reactions, these networks are called genome-scale metabolic networks (GEMs) [4]. These networks are typically represented in a matrix format, containing as many columns as reactions and as many rows as metabolites. Each reaction includes coefficients associated with the stoichiometry of the reaction, where negative values describe substrate consumption and positive numbers describe product formation. Under the steady-state assumption, no metabolite can have net accumulation or production, meaning that the sum of all reactions entering or leaving a metabolite node must be equal to zero. Formally, this is described by the well-known equation [1]:

$$S \cdot v = 0$$

where S is the stoichiometry network and $v$ is the vector of fluxes ($mmol \cdot gDW^{-1} \cdot h^{-1}$). Algebra tells us that if there are more equations (metabolites) than variables (fluxes), a system is overdetermined and has no exact solution. Alternatively, if the system is determined, a unique solution can be determined for the fluxes [1]. Unfortunately, this is rarely the case, as stoichiometric matrices typically contain more fluxes variables than equations. This necessitates the use of additional constraints and assumptions to obtain a unique solution.

The most commonly used approach to improve the identification of compatible fluxes with the observed consumptions of substrates and product formation is to transform the linear algebra problem into an optimization problem. The objective is to determine the set of reaction fluxes that maximize or minimize one or more defined metabolic functions. This optimization framework is known as flux balance analysis (FBA).

Under the assumption that microorganisms channel their resources optimally for biomass production (growth), the most widely used objective in FBA is growth rate maximization. Since biomass production involves a complex series of steps (tRNA charging, polymerization, DNA duplication, mitosis, etc.) and therefore not a real reaction, this process is typically approximated by the pseudo reaction known as the biomass equation. Essentially, this equation describes the main cellular polymers and the energy requirements (primarily ATP) for cell growth and division:

$$q_{ATP}\, ATP + \, q_{Protein}\, \text{protein} + \, q_{Carbohydrate}\, Carbohydrates \ldots \; \rightarrow \; 1\, \text{gDW} + q_{ATP}\, ADP \ldots$$

Earlier studies have measured the biomass composition of microorganisms such as *S. cerevisiae*, and these coefficients ($q_{ATP}$, $q_{Protein}$, etc.) are typically taken from those works.

When biomass maximisation is still insufficient to identify a unique flux solution, additional strategies can be utilised to improve the situation. The strategy used here - and perhaps the most widely used in conjunction with FBA - is parsimonious FBA (pFBA) [5]. As the name indicates, the goal is to find the most parsimonious of solutions under the assumption that cells utilise the minimal amount of enzyme possible and tend to avoid futile cycles and energy/carbon waste. In practical terms, pFBA is formulated as a two-step optimisation problem. First, the model is used to compute the maximum growth rate. Then, with the biomass reaction fixed at this value, the total sum of fluxes is minimised to identify the most parsimonious solution.

In this work, a batch growth system was used to study the physiology of *TSA1* deletion in dynamic diauxic shift experiments. Importantly, FBA can also be used to study metabolism in time-course experiments. This approach, known as dynamic FBA (dFBA), assumes that changes in intracellular compartments occur at a much faster pace (seconds to minutes) than the extracellular ones (hours to days). From an algorithmic point of view, dFBA requires that exchange fluxes ($mmol \times gDW^{-1} \times h^{-1}$) are computed at specific time points. As recently described [6], this variation can be computed with the help of a dynamic model.

#### 1.1. Dynamic model equations for growth

To estimate substrate uptake and product secretion rates in the dFBA model, as well as the intracellular fluxes affected by the *TSA1* deletion, we used a modified version of the model developed by Moimenta et al. (2025) [6] for simulating the diauxic shift from glucose to ethanol. To infer exchange rates for the methodology in the latter work, we first wrote a set of ordinary differential equations describing the variation in the concentration of biomass and different extracellular metabolites, represented by the following variables:

- $X(t)[gDW \cdot L^{-1}]$: dry-weight biomass concentration
- $S_{glc}(t)$, $S_{eth}(t)$, $S_{ac}(t)$, $S_{gly}(t)$, $S_{tre}(t)$ [mM]: extracellular concentrations of glucose, ethanol, acetate, and glycerol
- E(t) [Arbitrary Units]: relative activity of the non-fermentable-carbon gene module (*ADH2*, *ADY2*, etc.)

The production of biomass (g/L) was described as a function of three main components that depended on glucose, ethanol, and acetate concentrations (mM):

$$\frac{dW}{dt} = X \cdot \left(\mu_{glc} + \mu_{eth} + \mu_{ac}\right)$$

In turn, to describe $OD_{600}$ the following equation was used:

$$\frac{dOD600}{dt} = \frac{dX}{dt} \cdot \frac{1}{\lambda_{OD600}}$$

where the variation of dry-weight biomass was scaled by the factor $\lambda_{OD600}$. This value was fixed and chosen as 0.265 for all strains, based on preliminary tests. Because experimental data was available only for $OD600$, the initial condition for dry-weight biomass was set to $OD600_0 \cdot \lambda_{OD600}$, where the $OD600_0$ corresponds to the initial condition of optical density.

Growth rates for the three biomass-forming components were described using the Monod equation [7]. For example, in the case of glucose this was described by the following equation:

$$\mu_{Glucose} = \mu_{max,glc} \cdot \frac{S_{glc}}{S_{glc} + k_{s,glc}}$$

Where $\mu_{max,glc}$ is the maximal growth rate and $k_{s,glc}$ is the concentration (mM) at which growth is half-maximal.

In the case of ethanol- and acetate-dependent growth, we assumed that consumption of non-fermentable carbon sources was repressed by the absence of ethanol dehydrogenase *ADH2* and the transporter *ADY2*. Although the expression of *ADH2* was not measured in this work, this simplification is justified by the earlier proteomic measurements in diauxic shift experiments [8,9].

$$\mu_{Ethanol} = \mu_{max,Ethanol} \cdot E \cdot \frac{S_{eth}}{S_{eth} + k_{s,eth}}$$

$$\mu_{Acetate} = \mu_{max,Acetate} \cdot E \cdot \frac{S_{\mathrm{ac}}}{S_{\mathrm{ac}} + k_{s,ac}}$$

where $\mu_{max,Ethanol}$, $\mu_{max,ac}$, $k_{s,Acetate}$ and $k_{s,eth}$ are equivalent to those used in the glucose dependent Monod equation described above and have the same units.

Because the expression of genes associated with the consumption of non-fermentable carbon sources is generally correlated [8,9] to avoid adding unnecessary model complexity in the absence of expression data, here, we assumed that *ADY2* and *ADH2* can be approximately described by the same variable $E$. The expression of *ADH2*/*ADY2* was implemented with the help of logic-based ODEs [10–12]:

$$\frac{dE}{dt} = (\phi_{Glucose} - E) \cdot \mathrm{Tau}_E$$

where $\phi_{Glucose}$ is an inhibitory function describing glucose repression while $\tau_E$ is a parameter ($h^{-1}$) that defines how quickly $E$ converges to the set-point determined by $\phi_{Glucose}$, which is described by the following equation:

$$\phi_{Glucose} = 1 - \frac{S_{\mathrm{glc}}/X}{(S_{\mathrm{glc}}/X) + k_\phi}$$

where $k_\phi$ (mM) describes the value of $S_{\mathrm{glc}}/X$ (mM/gDW) at which the inhibition is half-maximal.

**1.2. Dynamic model equations for extracellular metabolites**

The consumption of all metabolites related to growth was described by the following equation:

$$v_{metabolite,in} = \frac{\mu_{metabolite}}{Y_{metabolite} \rightarrow X} \cdot X$$

where the metabolite is glucose (glc), ethanol (eth) or acetate (ac), and $\mu_{metabolite}$is the growth rate expression, described in the section above and $Y_{metabolite}$ (gDW· $mmol^{-1}$).

Glycerol was also consumed after the depletion of glucose. However, because its concentration was much lower than ethanol and acetate, we decided not to include it as a contributor to biomass production since this would introduce an additional parameter. Accordingly, glycerol consumption was described by the following Michaelis-Menten type equation to represent glycerol transport:

$$v_{Glycerol,in} = X \cdot v_{max,Glycerol} \cdot E \cdot \frac{S_{gly}}{S_{gly} + S_{gly}}$$

where $v_{max,Glycerol}$ is the maximal specific glycerol transport rate, $k_{s,Glycerol}$ (mM) is the concentration at which transport is half-maximal, and $E$ represents the expression of genes associated with the consumption of non-fermentable carbon sources.

In the model, we assumed that the metabolite production was associated either with the consumption of glucose during fermentation or ethanol consumption in the post-diauxic shift phase:

$$v_{metabolite,out} = v_{metabolite,in} \cdot Y_{metabolite,in \rightarrow metabolite,out}$$

where $Y_{metabolite,in \rightarrow metabolite,out}$ (mmol · mmol$^{-1}$) is the conversion yield of the consumed metabolite to the produced metabolite.

The net accumulation of extracellular metabolites was described as the difference between $v_{metabolite,out}$ and $v_{metabolite,in}$:

$$\frac{dmetabolite}{dt} = v_{metabolite,out} - v_{metabolite,in}$$

**Table 1** summarises the metabolite interactions that define the ordinary differential equations for each extracellular metabolite.

**Table 1. Summary of metabolite interactions in the dynamic model.**

| | Is consumed | Is produced from |
|---|---|---|
| Glucose | Yes, for biomass | - |
| Ethanol | Yes, for biomass | Glucose |
| Acetate | Yes, for biomass | Glucose |
| Glycerol | Yes, no sink | Glucose |
| Succinate | - | Ethanol |
| Pyruvate | - | Ethanol and Glucose |
| Succinate | | Ethanol |
| Trehalose (biomass; expressed in mM) | - | Ethanol |

#### 1.3. Estimation of $\mu_{max,glc}$

Growth during the exponential phase can be described with a linear model, as shown in the following equation:

$$log\ (y^m(\mathrm{t})) = log\left(\mathrm{OD}_{600,\mathrm{t}=0}\right) + \mu_{max,Glcucose} \cdot t$$

where $y^m$ is the linear model prediction, $X_0$ the initial biomass concentration and $\mu_{max,glc}$ is the maximal growth rate [7]. Therefore, to reduce potential identifiability problems with the rest of the parameters, we estimated $\boldsymbol{\mu_{max,glc}}$ from the experimental data. The model was fitted to the biomass data using the fitlm function in MATLAB.

As shown in **Figure 1**, the model shows almost perfect adjustment to the data. Interestingly, for both strains, the wild-type variant grows faster than the mutants and no substantial lag is observed in either case.

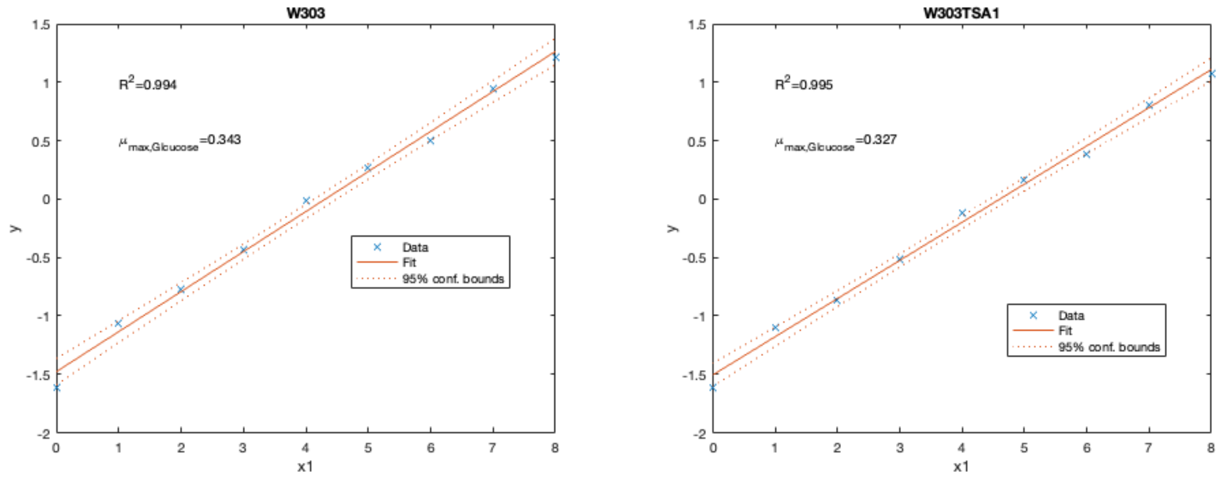


**Figure 1. Estimation of $\mu_{max,glc}$.** The adjustment of the linear model shows a near-perfect adjustment of the exponential growth model as indicated by the $R^2$ values close to 1.

#### 1.4. Choice the $Y_{glc \to X}$ parameter

To track biomass changes we used $OD_{600}$. However, the genome-scale metabolic model requires biomass information to scale the uptake rates. To convert the $OD_{600}$ observations to dry-weight during the glucose phase, we used glucose to biomass yield parameters taken from the literature and the previously mentioned scaling factor of $\lambda_{OD600}$.

Glucose-to-biomass conversion yield was chosen as 0.11 (gDW·$g^{-1}$; 0.019 gDW·$mmol^{-1}$) for W303. This yield value has been reported by van Dijken et al. [13] in different laboratory *S. cerevisiae* strains for growth during batch aerobic conditions using glucose as a carbon source.

This contrasts with fully respiratory growth, where the glucose-to-biomass conversion yield is roughly five times higher [13]. This choice is supported by the ethanol production yields since that, during exponential growth, the yield of ethanol was close to 1.5 mmol ethanol per mmol glucose. In comparison with earlier works [14], this yield suggests that growth is predominantly fermentative.

Finally, we tested different scaling factors between $OD_{600}$ and dry weight and found that a value of 0.265 g/OD provided a good fit to the data and was consistent across strains.

#### 1.5. Parameter estimation and calculation of confidence intervals

In total, 7 parameters were fixed and 13 were estimated. The fixed parameters and the rationale are given below:

- $\lambda_{OD600}$. Rationale: A value common to all strains was needed. In preliminary tests we found a value that was compatible with the experimental data and the glucose to biomass yields reported in the literature.
- $Y_{glc} \rightarrow X$. Rationale: Biomass was not measured. To convert $OD_{600}$ to dry weight, the values from van Dijken et al. (2000) [13] were used.
- $k_{s,eth}$. Rationale: Scarce data towards the end of the fermentation precluded the identification of this parameter. We focused in recovering $\mu_{max,eth}$ and fixed $k_{s,eth}$ to 1 mM.
- $k_{s,ac}$. Rationale: Scarce data towards the of the fermentation precluded the identification of this parameter. We focused in recovering $\mu_{max,ac}$ and $\mu_{max,eth}$ and $k_{s,ac}$ to 1 mM.
- $k_{s,gly}$. Rationale: Scarce data towards the of the fermentation precluded the identification of this parameter. We focused in recovering $v_{max,gly}$ and fixed $k_{s,gly}$ to 1 mM.
- $Tau_E$. Rationale: Since no expression data for glucose-repressed genes were available, we chose a relatively high value $Tau_E$.

The remaining parameters are taken as unknown and must be inferred from experimental measurements. This is formulated as a model calibration problem, in which one seeks the parameter values that minimise the discrepancy between model predictions and data, subject to the system's dynamic equations and any prescribed parameter bounds [15]. Equivalently, the optimal parameters maximise the (log-) likelihood of the observed data given the model:

$$\mathcal{J} = \log\left(\Pi(\hat{y}|\theta)\right)$$

Under the assumptions of independently identically distributed measurements according to a Gaussian law, the maximization of $\mathcal{J}$ is equivalent to minimizing:

$$J_{llk}(\theta) = \sum_{i=1}^{n_{exp}} \sum_{d=1}^{n_d} \frac{(y^d - y^m)^2}{\sigma_d^2}$$

where $n_{exp}$ corresponds to the number of experiments $n_d$ corresponds to the number of data points per experiment; $y^m$ and $y^d$ regard the model predictions and the measured data, respectively, and $\sigma_d$ is the standard deviation associated to the experimental data as computed from the experimental replicates.

Lack of sufficiently informative experimental data and a convoluted model structure can lead to poor identifiability of the parameters. Because of this, it is important to estimate confidence intervals for the parameters to ensure model robustness. To automate the model calibration and confidence interval estimation, we used the AMIGO2 toolbox [16]. This toolbox is a MATLAB-based software focused on dynamic model identification and optimisation that automatically reports the parametric confidence intervals as obtained by means of the Crammer-Rao inequality [15]. Here, CVODES [17] was used to obtain time-course trajectories and sensitivities, while the *enhanced Scatter Search* method [18] was used to optimise parameters.

The recovered parameters are indicated in **Table 2** along with the bounds used in the formulation of the parameter estimation. Parameters that are fixed to reduce estimation identifiability issues and their values are also indicated with ✓ in the fifth column.

**Table 2. Parameters of the dynamic model.** The “estimated” column indicates if parameters were estimated (✓) or fixed (X). Upper and lower bounds are also given, along with the recovered values for each strain.

| Parameter | Units | LB | UB | Estimated | W303 | W303 *tsa1*Δ |
|---|---|---|---|---|---|---|
| $k_{s,Glucose}$ | mM/gDW | 0.01 | 4 | ✓ | 0.01 | 0.033 |
| $k_{\phi}$ | mM | 0.01 | 100 | ✓ | 1.581 | 3.233 |
| $Y_{glc\rightarrow eth}$ | $mmol\cdot mmol^{-1}$ | 1 | 1.8 | ✓ | 1.516 | 1.604 |
| $Y_{\mathrm{eth/ac}\rightarrow X}$ | $gDW\cdot g^{-1}$ | 0.001 | 0.15 | ✓ | 0.018 | 0.014 |
| $Y_{eth\rightarrow \mathrm{pyr}}$ | $mmol\cdot mmol^{-1}$ | 0 | 2 | ✓ | 0.006 | 0.005 |
| $\mu_{max,\mathrm{eth}}$ | $h^{-1}$ | 0.01 | 1 | ✓ | 0.031 | 0.024 |
| $Y_{\mathrm{glc}\rightarrow ac}$ | $mmol\cdot mmol^{-1}$ | 0.001 | 1 | ✓ | 0.061 | 0.057 |
| $\mu_{max,\mathrm{ac}}$ | $h^{-1}$ | 0.0001 | 10 | ✓ | 0.002 | 0.002 |
| $Y_{Ethanol\rightarrow \mathrm{succ}}$ | $mmol\cdot mmol^{-1}$ | 0.001 | 0.03 | ✓ | 0.007 | 0.011 |
| $Y_{glc\rightarrow \mathrm{gly}}$ | $mmol\cdot mmol^{-1}$ | 0.01 | 0.2 | ✓ | 0.037 | 0.031 |
| $v_{max,gly}$ | $h^{-1}$ | 0 | 2 | ✓ | 0.0158 | 0.019 |
| $Y_{glc\rightarrow pyr}$ | $mmol\cdot mmol^{-1}$ | 0 | 10 | ✓ | 5.29E-10 | 2 E-09 |
| $Y_{\mathrm{eth}\rightarrow tre}$ | $mmol\cdot mmol^{-1}$ | 0.01 | 1 | ✓ | 0.032 | 0.016 |
| $\lambda_{OD600}$ | $OD\cdot DW^{-1}$ | 0.1 | 1 | X | 0.265 | 0.265 |
| $\mu_{max,glc}$ | $h^{-1}$ | 0.3 | 0.45 | X | 0.342 | 0.3269 |
| $Y_{glc} \rightarrow X$ | $gDW\cdot mmol^{-1}$ | 0.012 | 0.18 | X | 0.019 | 0.019 |
| $k_{s,eth}$ | mM | 0.02 | 100 | X | 1 | 1 |
| $k_{s,\mathrm{ac}}$ | mM | 0 | 100 | X | 1 | 1 |
| $k_{s,\mathrm{gly}}$ | mM | 0.001 | 1 | X | 1 | 1 |
| $Tau_E$ | $h^{-1}$ | 0.01 | 10 | X | 10 | 10 |

As shown in **Table 3**, the parameters exhibit reasonable confidence intervals, with few exceptions. One example is $k_{,s,Glucose}$ , which tended to approximate the lower bound and whose estimate is somewhat below the typical reported values for the high-affinity hexose transporters (around 1 mM).

**Table 3. Confidence intervals for model parameters.**

| | W303 | | | W303 *tsa1*Δ | | |
|---|---|---|---|---|---|---|
| **Parameter** | **Value** | **± Std Dev** | **Relative Error (%)** | **Value** | **± Std Dev** | **Relative Error (%)** |
| $k_{s,Glucose}$ | 1.00E-02 | 1.65E+00 | 16500% | 3.24E-02 | 9.49E-01 | 2930% |
| $k_{\phi}$ | 1.58E+00 | 1.15E+01 | 729% | 3.23E+00 | 1.73E+01 | 535% |
| $Y_{glc \to eth}$ | 1.52E+00 | 2.13E-01 | 14% | 1.60E+00 | 2.20E-01 | 13.70% |
| $Y_{eth \to X}$ | 1.82E-02 | 3.08E-03 | 16.90% | 1.40E-02 | 2.38E-03 | 17% |
| $Y_{eth \to pyr}$ | 6.48E-03 | 1.33E-03 | 20.60% | 5.46E-03 | 1.04E-03 | 19.10% |
| $\mu_{max,eth}$ | 3.17E-02 | 4.78E-03 | 15.10% | 2.45E-02 | 4.31E-03 | 17.60% |
| $Y_{glc \to ac}$ | 6.18E-02 | 1.27E-02 | 20.60% | 5.72E-02 | 1.09E-02 | 19.10% |
| $\mu_{max,ac}$ | 2.35E-03 | 1.25E-03 | 53.30% | 2.70E-03 | 8.59E-04 | 31.80% |
| $Y_{Ethanol \to succ}$ | 7.32E-03 | 1.41E-03 | 19.30% | 1.18E-02 | 2.42E-03 | 20.50% |
| $Y_{Glucose \to gly}$ | 3.71E-02 | 4.29E-03 | 11.60% | 3.15E-02 | 2.87E-03 | 9.08% |
| $v_{max,gly}$ | 1.59E-02 | 6.98E-03 | 43.90% | 1.90E-02 | 7.30E-03 | 38.40% |
| $Y_{glc \to pyr}$ | 3.63E-10 | 4.54E-04 | 1.25E+06 | 3.11E-09 | 4.08E-04 | 1.31E+05 |
| $Y_{eth \to CO2}$ | 5.36E-01 | 3.01E-01 | 56.10% | 4.75E-01 | 2.74E-01 | 57.80% |
| $Y_{glc \to CO2}$ | 9.40E-01 | 2.20E-01 | 23.40% | 1.24E+00 | 2.74E-01 | 22.10% |
| $Y_{eth \to tre}$ | 3.22E-02 | 6.30E-03 | 19.60% | 1.66E-02 | 2.82E-03 | 17% |

### 1.6. Model goodness of fit

As shown in **Figure 2**, the model fit was excellent, with $R^2$ values close to 1 for most observed variables. An exception to this was glycerol. This can likely be explained by the fact that glycerol was measured a posteriori using an enzymatic test kit, and the experiments were performed in a slightly different fermenter vessel.

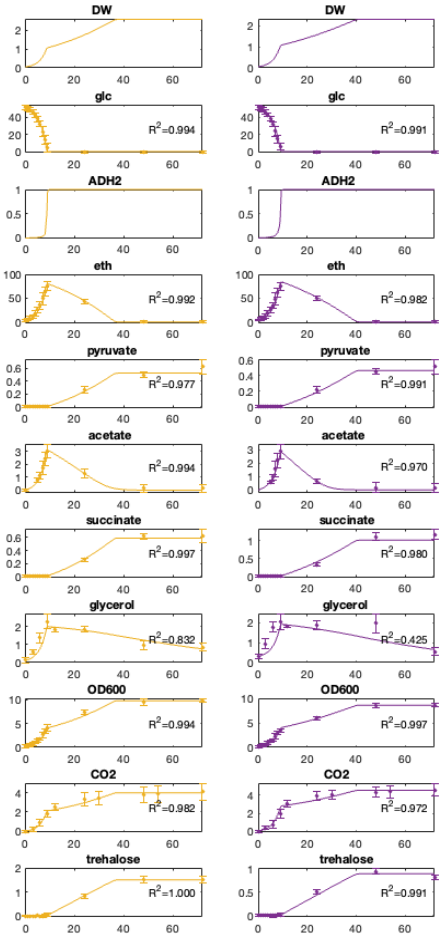


**Figure 2. Trajectories for the simulated data.** Points and error bars are the measured experimental data (mean of triplicates) and standard deviation estimated from the replicates. The high $R^2$ values (close to 1) for most observed variables indicate that the model explains the data well. W303 (yellow), W303 *tsa1*Δ (purple).

### 1.7. Dynamic FBA simulation.

Similarly to what was done in [6], we used the continuous model equations to constrain the exchange reactions of the genome-scale model. At specified time points ($t_i$) we used the dynamic model to evaluate the variation in net accumulation of extracellular metabolites per unit of biomass ($v_{Ex,ti}$):

$$v_{Ex,ti} = \frac{\frac{dmetabolite_{t_i}}{dt}}{X_{t_i}}$$

where $\frac{dmetabolite}{dt}$ represents the net accumulation or depletion of the compound (mmol·gDW$^{-1}$·h$^{-1}$) and $X_{t_i}$ is the simulated value of biomass.

The estimation of the fluxes was performed by solving the following linear optimization problem:

$$\max_{v_{ti}} c^T v_{t_i}$$

subject to

$$S \cdot v_{t_i} = 0$$

$$v_\mu = \frac{dX_{t_i}}{dt} / X_{t_i}$$

$$v_{min} \leq v_{t_i} \leq v_{max}$$

$$v_{Ex,t_i} \cdot (1 - tol) \leq v_{Ex_{t_i}} \leq v_{Ex,t_i} \cdot (1 + \mathrm{tol})$$

where, c is the objective function, S is stoichiometric matrix, $v_{min}$ and $v_{max}$ are the vector of upper and lower bounds for the fluxes (non-dynamic), $v_{Ex_{ti}}$ (dynamically changing) are the exchange fluxes for the extracellular metabolites for which we have data and *tol* is a tolerance factor.

The solutions for each time point within specific intervals were compiled so that the total flux was approximated using the following formula:

$$V_j = 100 \, \mathrm{x} \frac{\int_{t_A}^{t_B} v_j(t_i) \cdot DW((t_i)}{\int_{t_A}^{t_B} v_{Glx}(t_i) \cdot DW(t_i)}$$

where $t_A$ is lower bound and $t_B$ the upper bound of the time interval under analysis, and $V_j$ is the total flux (mmol; equivalent to concentration). This calculation was approximated with trapezoidal numerical integration in MATLAB.

Finally, the reported flux values are flux ratios, with respect to the main carbon source:

$$V_{Nj} = V_j / V_{CS}$$

where $V_{CS,t_i}$ (mmol; equivalent to concentration) is the consumed amount of glucose or ethanol during the analyzed period.

Here, because the growth rate was known, we set this value as a constraint and used other objective functions specific to each growth phase (see below). In addition, after optimizing the solutions for the objective function, we used parsimonious flux balance analysis to obtain the simplest explanation for the experimental data.

For the FBA problem, we used the Yeast9 (version 9.0.1) genome-scale metabolic model taken from: https://github.com/SysBioChalmers/yeast-GEM. A few curation steps were added to the model as part of an ongoing joint curation effort. These curation steps are currently available at: https://github.com/SysBioChalmers/yeast-GEM/tree/feat/anaerobic.

To solve the dFBA problem, we have coupled AMIGO initial value problem solver (CVODES) with COBRA [19]. We used a variable-step, variable-order We used a variable-step, variable-order Adams–Bashforth–Moulton method [20] to solve the initial value problem defined by the system of ordinary differential equations that describe the dynamics of the extracellular metabolites.

Two different intervals were analysed in this work with the help of the dFBA model. These used different constraints whose rationale will be described below.

**1.8. Parameters used for the dFBA during glucose-dependent growth**

The main parameters in the flux estimation problem during growth on glucose were:

- Period between 1 ($t_A$) and 8 ($t_B$) hours.
- Tolerance of 0.01 (1%) was assumed to avoid occasional infeasibility generated by the constraints.
- *GCY1* (glycerol dehydrogenase) does not carry flux due to unfavourable thermodynamics.
- *MDH2* is strongly repressed by glucose, and flux is constrained to 0.
- *IDP2* is strongly repressed by glucose and flux is constrained to 0.
- The growth rate is used as an equality constraint rather than as the objective. Data for the constraints is taken from the dynamic model.
- Acetate export was assumed to be carried out with the help of ATO channel transporters without proton co-transport.
- The objective function (c) was taken as the default biomass maximization.
- Growth-Associated-Maintenance (GAM) ATP = 55 mmol ATP/gDW.
- Non-Growth-Growth-Associated (NGAM) ATP was set to 1 (mmol·$gDW^{-1}$·$h^{-1}$).
- Flux ratios $V_{Nj}$ ratios were obtained by dividing $V_j$ by $V_{Glucose}$

#### 1.9. dFBA simulation after the diauxic shift.

The main parameters in the flux estimation problem during growth on ethanol were:

- Analyzed period between 10 ($t_A$) and 34 ($t_B$) hours of all strains.
- No tolerance (tol=0) was needed to obtain a solution feasible for all species
- The mitochondrial Acetyl-CoA encoded in the model was blocked because neither *ACS1* or *ACS2* are mitochondrial enzymes. Curation for future versions of the model is recommended.
- The objective function used was the cytosolic trehalase (r_0194) maximisation (c). In general we found an important decoupling between energy production and biomass formation during growth on ethanol for all strains.
- Acetate uptake was assumed to be carried out with the help of ADY or JEN1 transporters and therefore co-transported with a proton.
- Growth Associated Maintenance ATP = 55 mmol ATP/gDW
- Non-growth Growth associate ATP is set to 1 (mmol·gDW$^{-1}$·h$^{-1}$).
- Flux ratios $V_{Nj}$ ratios were obtained by diving $V_j$ by diving $V_{Ethanol}$.
- The biomass coefficient for trehalose ($q_{Trehalose}$) was set as $\frac{dtrehalose}{dt}/\frac{dDW}{dt}$ (mmol·gDW$^{-1}$). The interpretation of this is that for each gram of formed biomass ($\frac{dX}{dt}$) ,$\frac{dtrehalose}{dt}$ mmol trehalose is produced. In the model, the increase in trehalose was compensated by the reduction in protein content.
