## Supplementary Appendix 2 for "The peroxiredoxin Tsa1 promotes stationary phase entry by suppressing PKA activity": changeGAM.m

```
function model = changeGAM(model,GAM,NGAM)
bioPos = strcmp(model.rxnNames,'biomass pseudoreaction');
for i = 1:length(model.mets)
    S_ix  = model.S(i,bioPos);
    isGAM = sum(strcmp({'ATP','ADP','H2O','H+','phosphate'},model.metNames{i})) == 1;
    if S_ix ~= 0 && isGAM
        model.S(i,bioPos) = sign(S_ix)*GAM;
    end
end

if nargin >2
    pos = strcmp(model.rxnNames,'non-growth associated maintenance reaction');%NGAM
    model = setParam(model,'eq',model.rxns(pos),NGAM);% set both lb and ub
end

end
```
