## Supplementary Appendix 2 for "The peroxiredoxin Tsa1 promotes stationary phase entry by suppressing PKA activity": computeR2.m

```
load_zampar;
res=AMIGO_SData(inputs);

SSres=sum((res.sim.sim_data{1}-res.sim.exp_data{1}).^2,'omitnan');
SStot=sum((res.sim.exp_data{1}-mean(res.sim.exp_data{1},'omitnan')).^2,'omitnan');
R2_Z=1-SSres./SStot;

load_murphy;
res=AMIGO_SData(inputs);
SSres=sum((res.sim.sim_data{1}-res.sim.exp_data{1}).^2,'omitnan');
SStot=sum((res.sim.exp_data{1}-mean(res.sim.exp_data{1},'omitnan')).^2,'omitnan');
R2_M=1-SSres./SStot;

load_DiBartolomeo2020;
res=AMIGO_SData(inputs);
SSres=sum((res.sim.sim_data{1}-res.sim.exp_data{1}).^2,'omitnan');
SStot=sum((res.sim.exp_data{1}-mean(res.sim.exp_data{1},'omitnan')).^2,'omitnan');
R2_B=1-SSres./SStot;

load_diderich1999;
res=AMIGO_SData(inputs);
SSres=sum((res.sim.sim_data{1}-res.sim.exp_data{1}).^2,'omitnan');
SStot=sum((res.sim.exp_data{1}-mean(res.sim.exp_data{1},'omitnan')).^2,'omitnan');
R2_D=1-SSres./SStot;
```
