## Supplementary Appendix 2 for "The peroxiredoxin Tsa1 promotes stationary phase entry by suppressing PKA activity": eval_pars.m

```
function [f,h,g,regobj,regres] = eval_pars(theta,inputs,results,privstruct,ALPHA)

global n_amigo_sim_success;
global n_amigo_sim_failed;

theta = theta(:);
f=0.0;
g1=0.0;
g2=0.0;
g=[];
regobj = 0;
regres = [];
ncons=0;

h=zeros(1,privstruct.ntotal_constraints);
theta = AMIGO_scale_theta_back(theta,inputs);
privstruct.theta=theta;
privstruct=AMIGO_transform_theta(inputs,results,privstruct);

ndata=0;
nexpdata = zeros(1,inputs.exps.n_exp);
% number of data: total and for each experiment
for iexp=1:inputs.exps.n_exp
    % count the numerical element of exp_data. (excluding the nan-s)
    exp_data = inputs.exps.exp_data{iexp};
    
    if isempty(inputs.exps.nanfilter{iexp})
        nexpdata(iexp) = inputs.exps.n_s{iexp}*inputs.exps.n_obs{iexp};
        inputs.exps.nanfilter{iexp} = true(inputs.exps.n_s{iexp},inputs.exps.n_obs{iexp});
    else
        nexpdata(iexp) = numel(exp_data(inputs.exps.nanfilter{iexp}));
    end
    
    ndata=ndata+nexpdata(iexp);
end
g = zeros(1,ndata);

% USER PROVIDED COST FUNCTION
if strcmpi(inputs.PEsol.PEcost_type, 'user_PEcost')
    if ~isempty(inputs.PEsol.PEcost_file)
        [f, h ,g]=feval(strcat(inputs.PEsol.PEcost_file),theta);
    elseif isa(inputs.PEsol.PEcost_fun,'function_handle')
        [f, h ,g]=feval(inputs.PEsol.PEcost_fun,theta);
    else
        error('User defined PEcost must be either a function handler in inputs.PEsol.PEcost_fun or a name of a function in inputs.PEsol.PEcost_file')
    end
    return;
end

nprocessedData = 0;

PARS=[];

for iexp=1:inputs.exps.n_exp
    
    error_matrix=[];
    
    PAR=theta((((iexp-1)*inputs.model.n_par+1)):(iexp*inputs.model.n_par));
    PARS=[PARS PAR];
    privstruct.par=PAR;
    
    [yteor,privstruct,results]=...
        AMIGO_ivpsol(inputs,...
        privstruct,...
        inputs.exps.exp_y0{iexp},...
        PAR,...
        iexp,...
        results ...
        );
    
    privstruct.yteor=yteor;
    obsfunc=inputs.pathd.obs_function;
    
    ms=feval(...
        obsfunc,...
        privstruct.yteor,...
        inputs,...
        PAR,...
        iexp, ...
        privstruct.t_int{iexp}' ...
        );
    
    
    if(privstruct.ivpsol.ivp_fail | any(isnan(ms)))
        
        f=1e20;
        g(1:ndata) = Inf;
        h(1:privstruct.ntotal_constraints) =Inf(1,privstruct.ntotal_constraints);   
        
        %% EBC this I have to added for the case of constrained problems
        
        return;
        
    end
    
    % OBJECTIVE FUNCTION CALCULATION.
    switch inputs.PEsol.PEcost_type
        
        %LEAST SQUARES FUNCTION
        case 'lsq'
            
            % residuals in matrix form without weighting:
            resM = ms(inputs.exps.nanfilter{iexp})-inputs.exps.exp_data{iexp}(inputs.exps.nanfilter{iexp});
            
            switch inputs.PEsol.lsq_type
                
                case 'Q_mat'
                    % weighting matrix:
                    Q = inputs.PEsol.lsq_Qmat{iexp};
                    
                    scaledResM = resM.*Q;
                    % reshape to a columnvector:
                    g(nprocessedData+1 : nprocessedData + nexpdata(iexp)) = scaledResM(:);
                    f=g*g';
                    if isreal(f)==0 || isnan(f)==1
                        f=1e20; end
                    
                case 'Q_mat_obs' % for the cases where the observables have different matrices
                    % weighting matrix:
                    Q = inputs.PEsol.lsq_Qmat{iexp};
                    scaledResM = reshape(resM,1,ndata)*Q;
                    % reshape to a columnvector:
                    g = scaledResM;
                    f=g*g';
                    if isreal(f)==0 || isnan(f)==1
                        f=1e20; end
                    
                case 'Q_I'
                    g(nprocessedData+1 : nprocessedData + nexpdata(iexp)) = resM(:);
                    f=g*g';
                    
                    if isreal(f)==0 || isnan(f)==1
                        f=1e20; end
                    
                case 'Q_exp'
                    % weighting matrix is the measurements.
                    Q = 1./inputs.exps.exp_data{iexp};
                    tmp1 = or(isinf(Q),isinf(Q));
                    % if the inverse is inf or too huge (data near zero), than the
                    Q(tmp1) = 1;
                    scaledResM = resM.*Q;
                    % reshape to a columnvector:
                    g(nprocessedData+1 : nprocessedData + nexpdata(iexp)) = scaledResM(:);
                    f=g*g';
                    if isreal(f)==0 || isnn(f)==1
                        f=1e20; end
                    
                case 'Q_expmax'
                    expDataMax = max(inputs.exps.exp_data{iexp});
                    % every timepoint of each observable have the same
                    % weighting factor:
                    QQ = repmat(1./expDataMax, inputs.exps.n_s{iexp},1);
                    Q=QQ(inputs.exps.nanfilter{iexp});
                    % handles close infinity weights / clsoe zero experiments:
                    tmp1 = or(isnan(Q),isinf(Q));
                    Q(tmp1) = 1;
                    scaledResM = resM.*Q(:);
                    g(nprocessedData+1 : nprocessedData + nexpdata(iexp)) = scaledResM(:);
                    f=g*g';
                    if isreal(f)==0 || isnan(f)==1
                        f=1e20; end
                    
                case 'Q_expmean'
                    % weighting matrix is the mean of the trajectories
                    expDataMean = mean(inputs.exps.exp_data{iexp});
                    QQ = repmat(1./expDataMean, inputs.exps.n_s{iexp},1);
                    Q=QQ(inputs.exps.nanfilter{iexp});
                    % handles close infinity weights:
                    tmp1 = or(isnan(Q),isinf(Q));
                    Q(tmp1) = 1;
                    
                    scaledResM = resM.*Q(:);
                    % reshape to a columnvector:
                    g(nprocessedData+1 : nprocessedData + nexpdata(iexp)) = scaledResM(:);
                    f=g*g';
                    if isreal(f)==0 || isnan(f)==1
                        f=1e20;
                    end
                    
                otherwise
                    error('There is no such kind of lsq_type: %s', inputs.PEsol.lsq_type);
                    
                    
            end %switch inputs.lsq_type
            
        case 'llk'
            % EBC to make it work for homo_var--- REMOVE
            %inputs.PEsol.llk_type='homo_var';
            
            switch inputs.PEsol.llk_type
                
                case {'homo','homo_var'}
                    
                    error_data =  repmat(max(inputs.exps.exp_data{iexp}).*inputs.exps.std_dev{iexp},[inputs.exps.n_s{iexp},1]);
                    
                    error_data(error_data <= 1e-12) = inputs.ivpsol.atol;    %To avoid /0
                    
                    
                    for i_obs=1:inputs.exps.n_obs{iexp}
                        error_matrix=...
                            [error_matrix; (ms(:,i_obs)-inputs.exps.exp_data{iexp}(:,i_obs))./(error_data(:,i_obs))];
                    end
                    error_matrix = error_matrix(inputs.exps.nanfilter{iexp});
                    g(nprocessedData+1 : nprocessedData + nexpdata(iexp)) = error_matrix';
                    %                             g=[g error_matrix'];
                    %f = 0.5*sum(g.^2); if the 0.5 is needed here,
                    %be sure that the Least Squares local algorithm
                    %also takes the half. Otherwise the value
                    %computed by the eSS and the value computed by
                    %the NLS solver are different.
                    f = sum(g.^2);
                    
                    if isreal(f)==0 || isnan(f)==1
                        f=1e20;
                    end
                    
                    
                case 'hetero'
                    % changed by AG
                    % heteroscedastic noise with known  standard
                    % deviation (sigma and not sigma^2!)
                    % provided in inputs.exps.error_data{iexp}.
                    % weighted least squares case.
                    
                    
                    error_data = abs(inputs.exps.error_data{iexp});
                    error_data(abs(error_data) <= inputs.ivpsol.atol) = inputs.ivpsol.atol;    %To avoid /0
                    
                    try
                        error_matrix = (ms - inputs.exps.exp_data{iexp})./error_data;
                    catch
                        fprintf('The observation function is probably corrupted.\n')
                        keyboard
                    end
                    
                    error_matrix = error_matrix(inputs.exps.nanfilter{iexp});
                    
                    g(nprocessedData+1 : nprocessedData + nexpdata(iexp)) = error_matrix(:)';
                    
                    f=sum(g.^2);
                    
                    if isreal(f)==0 || isnan(f)==1
                        f=1e20;
                    end
                    
                case 'hetero_proportional'
                    %Standard deviation is assumed to be
                    %proportional to ms, noise = std_dev*y
                    
                    ms(ms<=1e-12)=inputs.ivpsol.atol;   % To avoid log(0) and /0
                    
                    for i_obs=1:inputs.exps.n_obs{iexp}
                        
                        g1=g1+sum(log(abs(ms(:,i_obs))));
                        
                        % dREAM g1=g1+sum(log(0.01*ones(size(ms(:,i_obs)))+0.04*ms(:,i_obs).^2));
                        
                        error_matrix=...
                            [error_matrix; (ms(:,i_obs)-inputs.exps.exp_data{iexp}(:,i_obs))...
                            ./(inputs.exps.std_dev{iexp}(i_obs).*ms(:,i_obs))];
                        
                    end %i_obs=1:inputs.exps.n_obs{iexp}
                    
                    %g is only approximated in this case, better
                    %not to use n2fb or dn2fb or NL2SOL
                    % DO NOT USE NL2SOL, THIS IS NOT A NONLINEAR
                    % LEAST SQUARES PROBLEM!!!
                    g(nprocessedData+1 : nprocessedData + nexpdata(iexp)) = error_matrix';
                    %g=[g error_matrix'];
                    
                    f=(2/ndata)*g1+sum(g.^2);
                    
                    if isreal(f)==0 || isnan(f)==1
                        f=1e20;
                    end
                    
                case 'hetero_lin'
                    
                    for i_obs=1:inputs.exps.n_obs{iexp}
                        
                        for i_s=1:inputs.exps.n_s{iexp}
                            if(abs(ms(i_s,i_obs))<=1e-12) % To avoid log(0) and /0
                                ms(i_s,i_obs)=inputs.ivpsol.atol;
                            end
                        end
                        
                        g1=g1+sum(log((inputs.PEsol.llk.stddeva{iexp}(i_obs)^2)*ones(size(ms(:,i_obs)))+(inputs.PEsol.llk.stddevb{iexp}(i_obs)^2)*ms(:,i_obs).^2));
                        % dREAM g1=g1+sum(log(0.01*ones(size(ms(:,i_obs)))+0.04*ms(:,i_obs).^2));
                        error_matrix=[error_matrix; (ms(:,i_obs)-inputs.exps.exp_data{iexp}(:,i_obs))./(inputs.PEsol.llk.stddeva{iexp}(i_obs)+inputs.PEsol.llk.stddevb{iexp}(i_obs).*ms(:,i_obs))];
                    end %i_obs=1:inputs.exps.n_obs{iexp}
                    
                    g(nprocessedData+1 : nprocessedData + nexpdata(iexp)) = error_matrix';
                    %g=[g error_matrix'];  % g is only approximated in this case, better not to use n2fb or dn2fb
                    
                    f=(2/ndata)*g1+sum(g.^2);
                    if ~isreal(f) || isnan(f)
                        f=1e20;
                    end

                otherwise
                    error('There is no such kind of llk_type: %s', inputs.PEsol.llk_type);
                    
            end %switch inputs.llk_type
            
        otherwise
            error('There is no such kind of cost-function type: %s', inputs.PEsol.PEcost_type);
    end %   switch inputs.PEsol.PEcost_type
    
    nprocessedData = nprocessedData + nexpdata(iexp);
    
    %%%%%%%%%%%%%%%%%%%%%%%%%%%%%
    % HANDLING CONSTRAINTS
    %%%%%%%%%%%%%%%%%%%%%%%%%%%%%
    if privstruct.n_const_ineq_tf>0 || privstruct.n_const_eq_tf>0
        
        if(numel(inputs.model.st_names)>0)
            for i=1:inputs.model.n_st
                eval(sprintf('%s=privstruct.yteor(end,%u);\n',inputs.model.st_names(i,:),i));
            end
            
        end
    end
    
    %%%%%%%%%%%%%%%%%%%%%%%%%%%%%%%%%%%
    % CONSTRAINTS: END-POINT EQUALITY
    %%%%%%%%%%%%%%%%%%%%%%%%%%%%%%%%%%%%
    % MOST METHODS HANDLE INEQUALITY CONSTRAINTS THUS WE WILL TRANSFORM
    % EQUALITY CONSTRAINTS INTO INEQUALITIES WITH A TOLERANCE INTRODUCED
    % BY USER.
    
    ncons=0;
    
    if sum(cell2mat(inputs.exps.n_const_eq_tf))>0
        
        if inputs.exps.n_const_eq_tf{iexp}>0
            for icons=1:inputs.exps.n_const_eq_tf{iexp}
                eval(sprintf('hh{iexp}(1,%u)=abs(%s);\n',icons,inputs.exps.const_eq_tf(icons,:)));
            end
            
            % FOR DE WE NEED TO IMPLEMENT THE DEATH PENALTY
            
            switch inputs.nlpsol.global_solver
                case 'de'
                    
                    for icons=1:inputs.exps.n_const_eq_tf{iexp}
                        if hh{iexp}(1,icons) > inputs.exps.eq_const_max_viol
                            f=inf;
                        end
                        
                    end
            end
        end  %if inputs.exps.n_const_eq_tf{iexp}>0
        
        ncons=ncons+inputs.exps.n_const_eq_tf{iexp};
    end  % sum(cell2mat(inputs.exps.n_const_eq_tf))>0
    
    
    %%%%%%%%%%%%%%%%%%%%%%%%%%%%%%%%%%%
    % CONSTRAINTS: END-POINT INEQUALITY
    %%%%%%%%%%%%%%%%%%%%%%%%%%%%%%%%%%%%
    niqcons=ncons;
    
    if sum(cell2mat(inputs.exps.n_const_ineq_tf))>0
        
        if inputs.exps.n_const_ineq_tf{iexp}>0
            
            for icons=1:inputs.exps.n_const_ineq_tf{iexp}
                eval(sprintf('hh{iexp}(1,%u) = %s;\n',icons,inputs.exps.const_ineq_tf{iexp}(icons,:)));
            end
            
            switch inputs.nlpsol.nlpsolver
                
                case 'de'
                    
                    for icons=1:inputs.exps.n_const_ineq_tf{iexp}
                        if hh{iexp}(1,icons) > inputs.exps.ineq_const_max_viol
                            f=inf;
                        end
                    end
            end
            
            if isempty(inputs.nlpsol.global_solver)
                inputs.nlpsol.global_solver=inputs.nlpsol.nlpsolver;
            end
            
            switch inputs.nlpsol.global_solver
                case 'de'
                    for icons=1:inputs.exps.n_const_ineq_tf{iexp}
                        if hh{iexp}(1,icons) > inputs.exps.ineq_const_max_viol
                            f=inf;
                        end
                    end
            end
            
            
        end
        
        niqcons=niqcons+inputs.exps.n_const_ineq_tf{iexp};
        
    end %if sum(cell2mat(inputs.exps.n_const_ineq_tf))>0
    
end

%%%%%%%%%%%%%%%%%%%%%%%%%%%%%%%
%%% HANDLING REGULARIZATION
%%%%%%%%%%%%%%%%%%%%%%%%%%%%%%%%
if inputs.nlpsol.regularization.ison
    %             fprintf('is on,\t')
    alpha = inputs.nlpsol.regularization.alpha;
    sqrt_alpha = realsqrt(alpha);
    
    [objReg, resReg] = AMIGO_PEcostreg(privstruct.theta,inputs,privstruct);
    
    %fprintf(1,'\treg.: Q_LS: %3.3g\t Pen: (%3.3g)*%3.3g = %3.3g \n',f,alpha,objReg,alpha*objReg);
    f = f + alpha*objReg;
    %             fprintf(2,'g size: %d\t',length(g));
    %             fprintf(2,'reg size: %d\t',length(sqrt_alpha*resReg'));
    g = [g sqrt_alpha*resReg'];
    %             fprintf(2,'tot size: %d\t',length(g));
    regobj = regobj + objReg;
    regres = [regres; resReg];
end

if privstruct.ntotal_constraints >0
    eval(sprintf('%s',inputs.pathd.ssconstraints));
    h=cell2mat(hh);
else
    h(1)=0;
end

inputs.exps.ndata=size(g,2);

DIFF=cell(3,3);
mean_P=mean(PARS');
cost_reg=0;
index_reg=ones(inputs.model.n_par,1);
index_reg(1)=0;
index_reg=logical(index_reg);

for i=1:inputs.exps.n_exp
    for j=1:inputs.exps.n_exp
        cost_reg= cost_reg+ sum(sum((ALPHA.*(PARS(index_reg,i)-PARS(index_reg,j))./mean_P(index_reg)).^2));
    end
end

f=f+cost_reg;
%f

return
```
