## Supplementary Appendix 2 for "The peroxiredoxin Tsa1 promotes stationary phase entry by suppressing PKA activity": fit_all.m

```
clear;
close all;
STRAINS{1}='zampar2013';
STRAINS{2}='murphy2015';
STRAINS{3}='DiBartolomeo2020';
STRAINS{4}='diderich1999';

data_set=1:4;

load_data;
LB=repmat(inputs.PEsol.global_theta_min,1,inputs.exps.n_exp);
UB=repmat(inputs.PEsol.global_theta_max,1,inputs.exps.n_exp);
X0=[];

for i=1:length(STRAINS)
    X0=[X0 PARS(:,i)'];
end

ALPHA=1e-5:1e-5:1e-4;
BETA=1e-4:1e-4:1e-3;
%% obtained with 2e-5
%XM=[0.036634	0.641223	0.364149	0.21353     26.1595	0.133592	0.635983	1.20882	5.29473	49.9977	386.03	944.274	0.0225613	0.0394144	0.0241259	1.82689	0.891884	1.94284	1.00123	0.131361	1.45877	0.0441828	1.31942	79.1991	1.58228	0.00892384	0.123387	0.285284	0.708559	0.246433	0.302062	0.147094	3.09429	5.87122	2.37632	3.23796	0.66847     20.4583	0.0404305	29.9704	0.375469	29.7433	1.43108	1.00189	1.14843	3.90126	0.21353	2.66966	0.957689	3.92878	0.057347	2.5785	0.394218	2.73727	0.517846	3.97515	0.764202	0.705066	0.101725	2.98223	0.902364	1.74217	0.0551153	2.11091	0.0320746	0.166389	0.796833	0.0060679	0.00249079	0.0968602	0.444239	4.69961	9.99981	0.0820012	0.755213	0.200074	0.298156	31.4399	0.000113725	1.54848	0.831011	6.35108	0.176598	381.818	946.036	0.0230455	0.0742566	0.0705741	0.780281	4.71782	1.77466	1.06182	0.0540695	1.51453	0.0435651	0.0906935	6.88891	0.456551	0.117034	0.0222863	0.266608	0.528037	0.137919	0.244884	3.76409	5.45561	1.17637	3.91553	3.94472	1.94924	0.26068	9.8142	29.9274	0.0131316	22.352	0.687861	1.01243	1.12143	1.80539	0.0502863	2.60795	0.80942	3.38516	0.116774	2.96189	0.301435	2.93137	0.405623	3.49697	0.0711051	0.664096	0.168472	2.9137	0.491085	0.210381	0.298685	9.99988	0.906941	0.00335008	0.0132459	1.87621	0.243195	0.928262	0.256575	15.2517	0.342837	0.191976	0.714886	0.810145	0.166616	8.12492	0.0952748	0.271456	1.99098	29.2596	17.0713	384.384	947.779	0.035276	0.0815774	0.098778	0.890747	1.12238	1.71162	1.81959	0.108939	1.99126	2.26494	1.08287	4.57293	0.931833	0.3083	7.74E-06	0.468823	0.0168828	0.029722	8.75051	0.00521927	0.0335044	2.12795	3.39147	2.40319	1.99962	15.2356	9.99729	0.301084	1.00078	22.0879	2.5212	3.84175	0.295756	1.26542	0.0589682	2.58804	1.53728	3.66801	0.0105691	3.55515	0.284315	1.00534	0.40364	3.71246	0.351776	0.637118	0.623567	2.90806	0.899525	1.02232	0.0846808	7.73206	0.592006	0.0287987	0.0913408	0.00030024	0.00505638	0.230699	0.021654	2.06103	2.84498	0.790151	0.447427	0.410139	0.216756	7.324	1.95416	14.9979	0.897866	1.55502	0.0940381	389.513	944.343	0.0199148	0.107968	0.093074	0.403186	3.52621	1.50427	1.58908	0.94726	1.62753	0.123999	0.799876	0.375409	8.92766	0.253582	0.13403	0.566358	0.864774	0.00454742	7.98889	0.423984	9.96825	9.64164	1.29755	3.67874	1.64025	11.5007	2.85617	29.7017	22.8869	21.8815	21.5043	3.87768	0.0208971	2.70711	0.789526	3.63428	3.99223	3.1396	2.53265	2.28343	3.99969	1.1643	3.46373	1.75131	2.86155	1.59136	0.00953197	0.000431281	2.55433	1.47095	0.0391777	2.8297	0.1067	0.177277	0.984998	0.0262908	0.0103167	0.200104	0.326432	4.99159	8.17423];
% eval_pars(XO,inputs,results,privstruct,ALPHA(8));


%% ALPHA(5)
problem.f   = 'eval_pars';
problem.x_L = LB;
problem.x_U = UB;
problem.x_0 = X0;
%       problem.neq     = Number of equality constraints (Integer; do not define it
%       problem.c_L     = Lower bounds of nonlinear inequality constraints
%       problem.c_U     = Upper bounds of nonlinear inequality constraints
%       problem.int_var = Number of integer variables (Integer)
%       problem.bin_var = Number of binary variables  (Integer)
%       problem.vtr     = Objective function value to be reached (optional)
opts.maxeval    = 10000;
%           opts.maxtime    = Maximum CPU time in seconds (Default 60)
%           opts.iterprint  = Print each iteration on screen: 0-Deactivated
%           opts.plot       = Plots convergence curves: 0-Deactivated,
%           opts.weight     = Weight that multiplies the penalty term added
%           opts.log_var    = Indexes of the variables which will be used
%           opts.tolc       = Maximum absolute violation of the constraints
%           opts.prob_bound = Probability (0-1) of biasing the search towards
%           opts.n_stuck    = Number of consecutive iterations without
opts.dim_refset         = 20;
opts.ndiverse           = 100;
%           opts.combination        = Type of combination of Refset
opts.local.solver = 'fminsearch';
%           opts.local.tol                  = Level of tolerance in local
opts.local.iterprint = 1;
opts.local.n1 = 1;
opts.local.n2= 2;
opts.local.finish = 0;

Results=ess_kernel(problem,opts,inputs,results,privstruct,BETA(4));
% plot_data(XM,inputs,results,privstruct,[0 0 0],1:3);   

plot_data(Results.xbest,inputs,results,privstruct,[0 0 0],1:3);   
% plot_data(X0,inputs,results,privstruct,[0 0 0],1:3);   
```
