## Supplementary Appendix 2 for "The peroxiredoxin Tsa1 promotes stationary phase entry by suppressing PKA activity": load_data.m

```
STRAINS{1}='zampar2013';
STRAINS{2}='murphy2015';
STRAINS{3}='DiBartolomeo2020';
STRAINS{4}='diderich1999';

STRAINS=STRAINS(data_set);

%% Paths folder
inputs.pathd.results_folder='diauxic';
inputs.pathd.short_name='diauxic_shift';

%% States and EQUATIONS
[vals_model eqns_model]=xlsread('modelA.xlsx','model');
type=char({eqns_model{2:end,3}});
ode_indices = find(~cellfun(@isempty,regexp({eqns_model{2:end,3}},'ode$')));
other_indices= find(cellfun(@isempty,regexp({eqns_model{2:end,3}},'ode$')));
flux_indices= find(~cellfun(@isempty,regexp({eqns_model{2:end,3}},'flux$')));
state_names={eqns_model{2:end,1}};
state_names=state_names(ode_indices);
Y0=vals_model(ode_indices,:);
N=length({eqns_model{2:end,1}});
d=repmat('d',N,1);
d(other_indices)=' ';
eqns=[d char({eqns_model{2:end,1}}') repmat('=',N,1) char({eqns_model{2:end,2}}')];

%% ODE model
inputs.model.n_st=length(state_names);
inputs.model.n_stimulus=0;
inputs.model.st_names=char(state_names);
inputs.model.eqns=eqns;

%% Pars
[vals_pars text_pars]=xlsread('modelA.xlsx','pars');
inputs.model.par_names=char(text_pars(2:end,1));
isOpt=vals_pars(:,3);
NSTRAINS=length(STRAINS);
PARS=vals_pars(:,3+data_set);

inputs.model.par=PARS(:,1);
inputs.model.n_par=length(inputs.model.par);

%% Parameter estimation
inputs.PEsol.id_global_theta=inputs.model.par_names(find(isOpt),:);
inputs.PEsol.global_theta_guess=inputs.model.par(find(isOpt))';
inputs.PEsol.global_theta_max=vals_pars(find(isOpt),2)';
inputs.PEsol.global_theta_min=vals_pars(find(isOpt),1)';

%% Error model
inputs.exps.data_type='real';
inputs.exps.noise_type='homo_var';
inputs.PEsol.PEcost_type='lsq'; %'llk';
inputs.PEsol.lsq_type='Q_expmax';
inputs.model.input_model_type='charmodelC';
inputs.ivpsol.max_step_size=10;
inputs.ivpsol.ivp_maxnumsteps=10000;
inputs.ivpsol.atol=1e-7;
inputs.ivpsol.rtol=1e-7;

%% STIMIULI
inputs.model.n_stimulus=1;
inputs.model.stimulus_names=char('D');

%% Experiments
inputs.exps.n_exp=length(STRAINS);

for iexp=1:inputs.exps.n_exp

    [vals_obs text_obs]=xlsread('modelA.xlsx',['obs_' STRAINS{iexp}]);
    
    index_col=find(~cellfun(@isempty,regexp({text_obs{1,:}},'obs_name')));
    
    inputs.exps.obs_names{iexp}=char(text_obs(1,index_col+2:end)');
    inputs.exps.n_obs{iexp}=size(inputs.exps.obs_names{iexp},1);
    inputs.exps.obs{iexp}=[char(text_obs(2,index_col+2:end')) repmat(';',inputs.exps.n_obs{iexp},1)];
    inputs.exps.exp_y0{iexp}=Y0(:,data_set(:,iexp));
    inputs.exps.exp_data{iexp}=vals_obs(1:end,(index_col+1):end);
    inputs.exps.t_s{iexp}=vals_obs(1:end,index_col)';
    inputs.exps.n_s{iexp}=length(inputs.exps.t_s{iexp});
    inputs.exps.t_f{iexp}=inputs.exps.t_s{iexp}(end);
    inputs.exps.error_data{1}=inputs.exps.exp_data{iexp}.*0.05;
    index_stim=find(~cellfun(@isempty,regexp({text_obs{1,:}},'stim_name')));
    index_obs=find(~cellfun(@isempty,regexp({text_obs{1,:}},'obs_name')));
    t_con=vals_obs(1:end,index_stim);
    inputs.exps.u_interp{iexp}='step';             
    inputs.exps.n_steps{iexp}=length(t_con)-1;
    inputs.exps.t_con{iexp}=t_con';             
    inputs.exps.u{iexp}=vals_obs(2:end,(index_stim+1):(index_obs-2))';
    
end

% inputs=AMIGO_Prep(inputs);
[results, inputs, privstruct]=prepare_PE(inputs);
```
