## Supplementary Appendix 2 for "The peroxiredoxin Tsa1 promotes stationary phase entry by suppressing PKA activity": plot_all.m

```
clear;
close all;
STRAINS{1}='zampar2013';
STRAINS{2}='murphy2015';
STRAINS{3}='DiBartolomeo2020';
data_set=1:3;

load_data;
LB=repmat(inputs.PEsol.global_theta_min,1,inputs.exps.n_exp);
UB=repmat(inputs.PEsol.global_theta_max,1,inputs.exps.n_exp);
X0=[];

for i=1:length(STRAINS)
    X0=[X0 PARS(:,i)'];
end

plot_data(X0,inputs,results,privstruct,[0 0 0],1:3);   
```
