## Supplementary Appendix 2 for "The peroxiredoxin Tsa1 promotes stationary phase entry by suppressing PKA activity": prepare_PE.m

```
function [results, inputs, privstruct]=prepare_PE(input_file)

%Reads defaults
[inputs_def]= AMIGO_private_defaults;

[inputs_def]= AMIGO_public_defaults(inputs_def);
% [inputs_def, results_def]= AMIGO_public_defaults(inputs_def);

%Checks for optional arguments
if nargin>1
    inputs_def.pathd.runident_cl=run_ident;
    inputs_def.pathd.runident=run_ident;
else
    %results_def.pathd.runident_cl=results_def.pathd.runident;
    inputs_def.pathd.runident=inputs_def.pathd.runident;
end


%Reads inputs

[inputs]=AMIGO_check_model(input_file,inputs_def);


[inputs]=AMIGO_check_exps(inputs);
[inputs]=AMIGO_check_obs(inputs);
[inputs]=AMIGO_check_data(inputs);

inputs = AMIGO_check_Q(inputs);
[inputs]= AMIGO_check_theta(inputs);
[inputs]= AMIGO_check_theta_bounds(inputs);
[inputs]=AMIGO_check_nlp_options(inputs);

[inputs]= AMIGO_scale_model_parameters(inputs);

privstruct=inputs.exps;
AMIGO_init_times
[inputs,privstruct]=AMIGO_check_NLPsolver(inputs,inputs.nlpsol.nlpsolver,privstruct);

%DETECTS PATH
AMIGO_path

%Creates necessary paths
AMIGO_paths_PE
AMIGO_init_report(inputs.pathd.report,inputs.pathd.problem_folder_path,inputs.pathd.task_folder)

AMIGO_gen_obs(inputs);

%Memory allocation and some necesary assignements
AMIGO_init_PE_guess_bounds
% Regularization needs the indexis of the theta
inputs = AMIGO_check_regularization(inputs);

%Generates matlab files for constraints
privstruct.n_const_ineq_tf=sum(cell2mat(inputs.exps.n_const_ineq_tf));
privstruct.n_const_eq_tf=sum(cell2mat(inputs.exps.n_const_eq_tf));
privstruct.n_control_const=sum(cell2mat(inputs.exps.n_control_const));
privstruct.ntotal_constraints= privstruct.n_const_ineq_tf+privstruct.n_const_eq_tf+privstruct.n_control_const;
privstruct.ntotal_obsconstraints=0;
privstruct.ntotal_tsconstraints=0;

for iexp=1:inputs.exps.n_exp
    privstruct.w_obs{iexp}=ones(1,inputs.exps.n_obs{iexp});
end

if privstruct.ntotal_constraints >0
    [results,inputs]=AMIGO_gen_PEconstraints(inputs,privstruct);
end


% creating indexing for the decision vector:
privstruct.index_theta = AMIGO_index_theta_vector(inputs);


switch inputs.exps.data_type
    
    case {'pseudo','pseudo_pos'}
        % generate pseudo experimental data.
        [inputs.exps.exp_data,inputs.exps.error_data,results.fit.residuals,results.fit.norm_residuals]=...
            AMIGO_pseudo_data(inputs,results,privstruct);
        results.sim.exp_data=inputs.exps.exp_data;
        results.sim.error_data=inputs.exps.error_data;        
        
    case {'real'}
        % transfer the data to the results.
        results.sim.exp_data=inputs.exps.exp_data;
        results.sim.error_data=inputs.exps.error_data;        
end
```
