## Supplementary Appendix 2 for "The peroxiredoxin Tsa1 promotes stationary phase entry by suppressing PKA activity": rescalePseudoReaction.m

```
function model = rescalePseudoReaction(model,metName,f)
  % rescalePseudoReaction
  %   Rescales a specific pseudoreaction by a given factor
  %
  %   model      (struct) the yeast GEM
  %   metName    (str) name of the component to rescale (e.g. "protein")
  %   f          (float) fraction to use for rescaling
  %
  %   model    (struct) the (rescaled) yeast GEM
  %
  %   Usage: model = rescalePseudoReaction(model,metName,f)
  %

if strcmp(metName,'lipid')
    model = rescalePseudoReaction(model,'lipid backbone',f);
    model = rescalePseudoReaction(model,'lipid chain',f);
else
    rxnName = [metName ' pseudoreaction'];
    rxnPos  = strcmp(model.rxnNames,rxnName);
    for i = 1:length(model.mets)
        S_ir   = model.S(i,rxnPos);
        isProd = strcmp(model.metNames{i},metName);
        if S_ir ~= 0 && ~isProd
            model.S(i,rxnPos) = f*S_ir;
        end
    end
end

end
```
