## Supplementary Appendix 2 for "The peroxiredoxin Tsa1 promotes stationary phase entry by suppressing PKA activity": scaleBioMass.m

```
function model = scaleBioMass(model,component,new_value,balance_out,dispOutput)
  % scaleBioMass
  %   Scales the biomass composition
  %
  %   model          (struct) metabolic model in COBRA format
  %   component      (str) name of the component to rescale (e.g. "protein")
  %   new_value      (float) new total fraction for said component
  %   balance_out    (str, opt) if chosen, the name of another component with which
  %                  the model will be balanced out so that the total mass remains = 1 g/gDW
  %   dispOutput     (bool, opt) if output from sumBioMass should be displayed (default = true)
  %
  %   model          (struct) modified model
  %
  %   Usage: model = scaleBioMass(model,component,new_value,balance_out,dispOutput)
  %

if nargin < 5
    dispOutput = true;
end
  
%Measure current composition and rescale:
[~,P,C,R,D,L,I,F] = sumBioMass(model,dispOutput);
content_all = {'carbohydrate','protein','lipid','RNA','DNA','ion','cofactor'};
content_Cap = {'C','P','L','R','D','I','F'};
pos         = strcmp(content_all,component);
old_value   = eval(content_Cap{pos});
f           = new_value / old_value;
model       = rescalePseudoReaction(model,component,f);

%Balance out (if desired):
if nargin > 3
    pos           = strcmp(content_all,balance_out);
    balance_value = eval(content_Cap{pos});
    f             = (balance_value - (new_value - old_value)) / balance_value;
    model         = rescalePseudoReaction(model,balance_out,f);
end

end
```
