## Supplementary Appendix 2 for "The peroxiredoxin Tsa1 promotes stationary phase entry by suppressing PKA activity": sim_obs.m

```
function obs = sim_obs(x,inputs,privstruct,results)

obs=cell(1,inputs.exps.n_exp);

for iexp=1:inputs.exps.n_exp
    
    PAR=x((((iexp-1)*inputs.model.n_par+1)):(iexp*inputs.model.n_par));
    privstruct.par=PAR;
    
    [yteor,privstruct,results]=...
        AMIGO_ivpsol(inputs,...
        privstruct,...
        inputs.exps.exp_y0{iexp},...
        PAR,...
        iexp,...
        results ...
        );
    
    privstruct.yteor=yteor;
    obsfunc=inputs.pathd.obs_function;
    
    obs{iexp}=feval(...
        obsfunc,...
        privstruct.yteor,...
        inputs,...
        PAR,...
        iexp,...
        privstruct.t_int{iexp}'...
        );
    
    
end
```
