## Supplementary Appendix 2 for "The peroxiredoxin Tsa1 promotes stationary phase entry by suppressing PKA activity": sumBioMass.m

```
function [X,P,C,R,D,L,I,F] = sumBioMass(model,dispOutput)
  % sumBioMass
  %   Calculates breakdown of biomass
  %
  %   model         (struct) Metabolic model in COBRA format
  %   dispOutput    (bool, opt) If output should be displayed (default = true)
  %
  %   X             (float) Total biomass fraction [gDW/gDW]
  %   P             (float) Protein fraction [g/gDW]
  %   C             (float) Carbohydrate fraction [g/gDW]
  %   R             (float) RNA fraction [g/gDW]
  %   D             (float) DNA fraction [g/gDW]
  %   L             (float) Lipid fraction [g/gDW]
  %   F             (float) cofactor [g/gDW]
  %   I             (float) ion [g/gDW]
  %
  %   Usage: [X,P,C,R,D,L,I,F] = sumBioMass(model,dispOutput)
  %
  %   Function adapted from SLIMEr: https://github.com/SysBioChalmers/SLIMEr
  %

if nargin < 2
    dispOutput = true;
end

%Load original biomass component MWs:
%TODO: compute MW automatically from chemical formulas (check that all components have them first)
fid = fopen('biomassComposition_Forster2003.tsv');
Forster2003 = textscan(fid,'%s %s %f32 %f32 %s','Delimiter','\t','HeaderLines',1);
data.mets   = Forster2003{1};
data.MWs    = double(Forster2003{4});
fclose(fid);

%load additional cofactor/ion MWs:
fid = fopen('biomassComposition_Cofactor_Ion.tsv');
CofactorsIons = textscan(fid,'%s %s %f32 %f32 %s %s','Delimiter','\t','HeaderLines',1);
data_new.mets = CofactorsIons{1};
data_new.MWs  = double(CofactorsIons{4});
fclose(fid);
for i = 1:length(data_new.mets)
    if ~ismember(data_new.mets(i),data.mets)
        data.mets = [data.mets; data_new.mets(i)];
        data.MWs  = [data.MWs; data_new.MWs(i)];
    end
end

%Get main fractions:
[P,X] = getFraction(model,data,'P',0,dispOutput);
[C,X] = getFraction(model,data,'C',X,dispOutput);
[R,X] = getFraction(model,data,'R',X,dispOutput);
[D,X] = getFraction(model,data,'D',X,dispOutput);
[L,X] = getFraction(model,data,'L',X,dispOutput);
[I,X] = getFraction(model,data,'I',X,dispOutput);
[F,X] = getFraction(model,data,'F',X,dispOutput);

if dispOutput
    disp(['X -> ' num2str(X) ' gDW/gDW'])
    % Simulate growth:
    sol = optimizeCbModel(model);
    disp(['Growth = ' num2str(sol.f) ' 1/h'])
    disp(' ')
end

end

%%

function [F,X] = getFraction(model,data,compType,X,dispOutput)

%Define pseudoreaction name:
rxnName = [compType ' pseudoreaction'];
rxnName = strrep(rxnName,'P','protein');
rxnName = strrep(rxnName,'C','carbohydrate');
rxnName = strrep(rxnName,'N','biomass');
rxnName = strrep(rxnName,'L','lipid backbone');
rxnName = strrep(rxnName,'R','RNA');
rxnName = strrep(rxnName,'D','DNA');
rxnName = strrep(rxnName,'I','ion');
rxnName = strrep(rxnName,'F','cofactor');

%Add up fraction:
rxnPos = strcmp(model.rxnNames,rxnName);
if ~all(rxnPos==0)
    isSub   = model.S(:,rxnPos) < 0;        %substrates in pseudo-rxn
    if strcmp(compType,'L')
        F = -sum(model.S(isSub,rxnPos));   %g/gDW
    else
        F = 0;
        %Add up all components:
        for i = 1:length(model.mets)
            pos = strcmp(data.mets,model.mets{i});
            if isSub(i) && sum(pos) == 1
                if strcmp(compType,'I') || strcmp(compType,'F')
                    MW = data.MWs(pos);
                else
                    MW = data.MWs(pos)-18;
                end
                abundance = -model.S(i,rxnPos)*MW/1000;
                F         = F + abundance;
            end
        end
    end
    X = X + F;
    
    if dispOutput
        disp([compType ' -> ' num2str(F) ' g/gDW'])
    end
else
    if dispOutput
        disp([compType ' does not exist '])
    end
    F = 0;
    X = X + F;
end

end
```
