## Supplementary Appendix 2 for "The peroxiredoxin Tsa1 promotes stationary phase entry by suppressing PKA activity": eth_cons.m

```
function model = eth_cons(model)

RD=0;
%
% Updated the adjustment of degree of reduciton /Gustav
% Try NADPH for DR balance instead. Also, try to convert NADH to NADPH at an adjustable ratio
% is not required. But we can consider if NADH or NADPH should be used to
% balance the degree of reduction of the biomass
NADH_NADPH=0;

% NADPH
model.S(find(strcmp(model.mets,'s_1212')),strcmp(model.rxns,'r_4041'))=model.S(find(strcmp(model.mets,'s_1212')),strcmp(model.rxns,'r_4041'))-RD-NADH_NADPH;
% NADP
model.S(find(strcmp(model.mets,'s_1207')),strcmp(model.rxns,'r_4041'))=model.S(find(strcmp(model.mets,'s_1207')),strcmp(model.rxns,'r_4041'))+RD+NADH_NADPH;
% NADH
model.S(find(strcmp(model.mets,'s_1203')),strcmp(model.rxns,'r_4041'))=model.S(find(strcmp(model.mets,'s_1203')),strcmp(model.rxns,'r_4041'))+NADH_NADPH;
% NAD
model.S(find(strcmp(model.mets,'s_1198')),strcmp(model.rxns,'r_4041'))=model.S(find(strcmp(model.mets,'s_1198')),strcmp(model.rxns,'r_4041'))-NADH_NADPH;
% H+
model.S(find(strcmp(model.mets,'s_0794')),strcmp(model.rxns,'r_4041'))=model.S(find(strcmp(model.mets,'s_0794')),strcmp(model.rxns,'r_4041'))-RD;


%% Fumarate reductase is required for to recycle FADH2 derived from
% disulphide bound formation in anaerobic conditions through Ero1 (Camarasa
% et al (2007) 10.1002/yea.1467; Kim et al (2018) 10.1038/s41467-018-07285-9). 

FADH2_prod=0.08;
metIdx = getIndexes(model,{'s_0689','s_0687','s_0794'},'mets'); % FADH2[c], FAD[c], H+[c]
bioIdx = getIndexes(model,'r_4041','rxns');

currCoeff = full(model.S(metIdx,bioIdx)); % Gather the current coefficients
model.S(metIdx,bioIdx) = currCoeff + [FADH2_prod; -FADH2_prod; -2*FADH2_prod];

end
```
