## Supplementary Appendix 2 for "The peroxiredoxin Tsa1 promotes stationary phase entry by suppressing PKA activity": fit_mu_data.m

```
clear;
close all;

STRAINS={'T73','T73TSA1','W303','W303TSA1'};
for data_set=1:4

    subplot(2,2,data_set);

    STRAIN=STRAINS{data_set};
    load_data;

    X = inputs.exps.exp_data{1}(1:9,1); % Biomass measurements (e.g., OD600)
    t=0:8;
    % Take the natural logarithm of biomass
    lnX = log(X);
    t=0:8;
    % Fit a linear regression model (Y = mx + b)
    A = [t(:), ones(length(t),1)]; % Design matrix
    b = lnX(:); % Response variable (lnX)
    mdl = fitlm(t,b);
    plot(mdl);
    text(1,1,['R^2=' sprintf('%.3f',mdl.Rsquared.Ordinary)]);
    text(1,0.5,['\mu_{max,Glcucose}=' sprintf('%.3f',mdl.Coefficients.Estimate(2))]);
    title(STRAINS(data_set));

end

% Solve using leas
% t squares (A \ b is the equivalent of inv(A'*A) * A' * b)
% params = A \ b;
%
% % Extract growth rate (slope of lnX vs. t)
% mu = params(1);  % The slope is the growth rate
%
% % Display results
% fprintf('Estimated Growth Rate (mu): %.4f per hour\n', mu);
%
% % Optional: Plot the data and fitted line
% figure;
% plot(t, lnX, 'bo', 'MarkerFaceColor', 'b', 'DisplayName', 'Log Biomass Data');
% hold on;
% plot(t, A*params, 'r-', 'LineWidth', 2, 'DisplayName', 'Linear Fit');
% xlabel('Time (hours)');
% ylabel('ln(Biomass)');
% title('Growth Rate Estimation');
% legend;
% grid on;
```
