## Supplementary Appendix 2 for "The peroxiredoxin Tsa1 promotes stationary phase entry by suppressing PKA activity": load_data.m

```
STRAINS={'T73','T73TSA1','W303','W303TSA1'};
STRAIN=STRAINS{data_set};

[DATA BB]= xlsread('data_ferm.xlsx',['data_' STRAIN]);

t=DATA(:,1);
TT=unique(t);
t=t(1:end);
exp_data=[];
error_data=[];

for i=1:length(TT)
    row= find(t==TT(i));
    exp_data=[exp_data; mean(DATA(row,2:end),'omitnan')];    
    error_data=[error_data; std(DATA(row,2:end),'omitnan')];
end

%% Paths folder
inputs.pathd.results_folder='diauxic';
inputs.pathd.short_name='diauxic_shift';

%% States and EQUATIONS
[vals_model eqns_model]=xlsread('data_ferm.xlsx','model');
type=char({eqns_model{2:end,3}});
ode_indices = find(~cellfun(@isempty,regexp({eqns_model{2:end,3}},'ode$')));
other_indices= find(cellfun(@isempty,regexp({eqns_model{2:end,3}},'ode$')));
flux_indices= find(~cellfun(@isempty,regexp({eqns_model{2:end,3}},'flux$')));
state_names={eqns_model{2:end,1}};
state_names=state_names(ode_indices);
Y0=vals_model(ode_indices,data_set);
N=length({eqns_model{2:end,1}});
d=repmat('d',N,1);
d(other_indices)=' ';
eqns=[d char({eqns_model{2:end,1}}') repmat('=',N,1) char({eqns_model{2:end,2}}')];

%% ODE model
inputs.model.n_st=length(state_names);
inputs.model.st_names=char(state_names);
inputs.model.eqns=eqns;

%% Pars
[vals_pars text_pars]=xlsread('data_ferm.xlsx','pars');
inputs.model.par_names=char(text_pars(2:end,1));
isOpt=vals_pars(:,3);

NSTRAINS=1;
PARS=vals_pars(:,3+data_set);

inputs.model.par=PARS(:,1);
inputs.model.n_par=length(inputs.model.par);

%% Parameter estimation
inputs.PEsol.id_global_theta=inputs.model.par_names(find(isOpt),:);
inputs.PEsol.global_theta_guess=inputs.model.par(find(isOpt))';
inputs.PEsol.global_theta_max=vals_pars(find(isOpt),2)';
inputs.PEsol.global_theta_min=vals_pars(find(isOpt),1)';

%% Error model
inputs.exps.data_type='real';
inputs.exps.noise_type='homo_var';
inputs.PEsol.PEcost_type='llk'; %'llk';
inputs.PEsol.lsq_type='Q_expmax';
inputs.model.input_model_type='charmodelC';
inputs.ivpsol.max_step_size=10;
inputs.ivpsol.ivp_maxnumsteps=10000;
inputs.ivpsol.atol=1e-7;
inputs.ivpsol.rtol=1e-7;

inputs.nlpsol.eSS.maxeval=1000;

inputs.nlpsol.eSS.maxtime=inf;
inputs.nlpsol.eSS.ndiverse           = 200;

inputs.nlpsol.eSS.dim_refset         = 10;
inputs.nlpsol.eSS.local.n1=5;
inputs.nlpsol.eSS.local.n2=5;
inputs.nlpsol.eSS.local.n1=0;
inputs.nlpsol.eSS.local.n2=0;


%% Experiments
inputs.exps.n_exp=1;

for iexp=1:inputs.exps.n_exp
    inputs.exps.obs_names{iexp}=  char(BB(2:end)');
    inputs.exps.n_obs{iexp}=length(BB)-1;
    inputs.exps.obs{iexp}=char(strcat(BB(2:end)' ,cellstr(repmat('_o=',length(BB)-1,1)),BB(2:end)'));
    inputs.exps.exp_y0{iexp}=Y0;
    inputs.exps.exp_data{iexp}=exp_data;
    inputs.exps.error_data{iexp}=error_data+repmat(max(exp_data).*0.025,length(TT),1);
    inputs.exps.t_s{iexp}=TT';
    inputs.exps.n_s{iexp}=length(inputs.exps.t_s{iexp});
    inputs.exps.t_f{iexp}=inputs.exps.t_s{iexp}(end);
end
```
