## Supplementary Appendix 2 for "The peroxiredoxin Tsa1 promotes stationary phase entry by suppressing PKA activity": model_curation.m

```
%% ========================================================================
% We blocked MDH2 in anaerobic conditions (see details in the anerobicModel
% script) Experiments suggest that AKG needs to produced inside the
% mitochondria and exported with ODC1/2 the help or YHM2. After inspecting
% the kinetics parameters for OAC1, DIC1, YHM2 and ODC1/ODC2 (SFC1 is
% strongly repressed by glucose) we added sulphate and malate as substrate
% for the OAC1 transporter.

% Add sulphate[m]
newMet = struct('metNames', {{'sulphate'}}, ...
    'compartments', {{'m'}});
model = addMets(model,newMet,true,'s_');
fprintf('Identifier of new metabolite "%s[%s]": %s\n', newMet.metNames{1}, newMet.compartments{1}, model.mets{end});

% Add oxaloacetate/sulphate antiporter
newRxn = struct('rxns', {generateNewIds(model,'rxns','r_',1)}, ...
    'equations', {{'oxaloacetate[c] + sulphate[m] <=> oxaloacetate[m] + sulphate[c]'}}, ...
    'rxnNames', {{'oxaloacetate/sulphate antiport, mitochondrial'}}, ...
    'subSystems', {{'Transport [c, m]'}}, ...
    'grRules', {{'YKL120W'}},...
    'rxnReferences', {{'10.1074/jbc.274.32.22184'}}, ...
    'rxnConfidenceScores', {3});
model = addRxns(model,newRxn,3);
fprintf('Identifier of new reaction "%s": %s\n', newRxn.rxnNames{1}, model.rxns{end});

% Add malate/sulphate antiporter
newRxn = struct('rxns', {generateNewIds(model,'rxns','r_',1)}, ...
    'equations', {{'(S)-malate[c] + sulphate[m] <=> (S)-malate[m] + sulphate[c]'}}, ...
    'rxnNames', {{'malate/sulphate antiport, mitochondrial'}}, ...
    'subSystems', {{'Transport [c, m]'}}, ...
    'grRules', {{'YKL120W'}},...
    'rxnReferences', {{'10.1074/jbc.274.32.22184'}}, ...
    'rxnConfidenceScores', {3});
model = addRxns(model,newRxn,3);
fprintf('Identifier of new reaction "%s": %s\n', newRxn.rxnNames{1}, model.rxns{end});

%% ========================================================================
% Look for all proton symport/antiport reactions and make sure that they
% only enter the cell.
% HcytIdx = getIndexes(model,'s_0794','mets'); % H+[c]
% HextIdx = getIndexes(model,'s_0796','mets'); % H+[e]
%
% symporterIDs = find(model.S(HcytIdx,:) & model.S(HextIdx,:));
% for i = 1:length(symporterIDs)
%     if strcmp(model.rxns(symporterIDs(i)), 'r_1258')
%         % Ignore the sodium transporter, without it, the model does not work
%         continue
%     end
%     if model.S(HextIdx,symporterIDs(i))<0 % If defined H+[e] => H+[c]
%         model.lb(symporterIDs(i))=0;
%     else % If defined H+[c] => H+[e]
%         model.ub(symporterIDs(i))=0;
%     end
% end

%% ========================================================================
% This section balances reactions and ensures that a correct molecular
% weight can be calculated for the biomass

% Set the charge of all biomass components to 0
model.metCharges(strcmp(model.mets,'s_3717'))=0; % Protein
model.metCharges(strcmp(model.mets,'s_3718'))=0; % Carbohydrate
model.metCharges(strcmp(model.mets,'s_3719'))=0; % RNA
model.metCharges(strcmp(model.mets,'s_3720'))=0; % DNA
model.metCharges(strcmp(model.mets,'s_3746'))=0; % Lipid backbone
model.metCharges(strcmp(model.mets,'s_3747'))=0; % Lipid chain
model.metCharges(strcmp(model.mets,'s_4205'))=0; % Cofactor
model.metCharges(strcmp(model.mets,'s_4206'))=0; % Ion

% Make the charge of K and Na 1+
model.metCharges(strcmp(model.mets,'s_1373'))=1;
model.metCharges(strcmp(model.mets,'s_1374'))=1;
model.metCharges(strcmp(model.mets,'s_3776'))=1;
model.metCharges(strcmp(model.mets,'s_1437'))=1;
model.metCharges(strcmp(model.mets,'s_1438'))=1;
model.metCharges(strcmp(model.mets,'s_3775'))=1;


% Make the charge of HS (hydrogen sulfide) -1
model.metCharges(strcmp(model.mets,'s_0841'))=-1;
model.metCharges(strcmp(model.mets,'s_3906'))=-1;
model.metCharges(strcmp(model.mets,'s_4263'))=-1;

% Balance the reaction r_4702, 'L-cysteine:2-oxoglutarate aminotransferase'
% by adding a proton as reactant
model.S(find(strcmp(model.mets,'s_0794')),strcmp(model.rxns,'r_4702'))=-1;

% Balance the reaction r_4703, 'L-cysteine:2-oxoglutarate aminotransferase'
% by adding a proton as product
model.S(find(strcmp(model.mets,'s_0794')),strcmp(model.rxns,'r_4703'))=1;

% Balance the reactions 'r_0774' and 'r_0775', 'NAPRtase' by removing H+
% consumption and adding a H2O as a reactant
model.S(find(strcmp(model.mets,'s_0794')),strcmp(model.rxns,'r_0774'))=0;
model.S(find(strcmp(model.mets,'s_0803')),strcmp(model.rxns,'r_0774'))=-1;
model.S(find(strcmp(model.mets,'s_0794')),strcmp(model.rxns,'r_0775'))=0;
model.S(find(strcmp(model.mets,'s_0803')),strcmp(model.rxns,'r_0775'))=-1;

%% ========================================================================
% This section focuses on individual reactions that have the wrong
% reversibility/direction/cofactor or should be completley removed

% Make GCY1 irreversible. Has a positive DeltaGo' (+20.9) and is part of a
% transhydrogenase cycle (NADH -> NADPH) at the cost of one ATP. High
% cytosolic NADPH/NADP ratio makes it thermodynamically infeasible that it
% runs in reverse direction.
model = setParam(model,'ub','r_0487',0);

% The mitochondrial ATP synthase is able to run in reverse, which occurs
% in anaerobic conditions
model = setParam(model,'lb','r_0226',-1000);

% Rename r_0227, it is the plasma membrane ATPase, not a cytosolic ATPase
model.rxnNames(strcmp(model.rxns,'r_0227')) = {'ATPase, plasma membrane'};

% Make sure both formate-THF ligases are reversible (r_0447 already is).
model = setParam(model,'lb','r_0446',-1000);

% Make methylenetetrahydrofolate dehydrogenases ADE3 and MIS1 irreversible
model = setParam(model,'ub',{'r_0732','r_0733'},0);

% There is no evidence for this PFK1 side reaction in yeast. Consider
% removing the reaction completley
model = setParam(model,'eq','r_0887',0);

% TYR1 incorrectly annotated as using NAD, should be NADP
model.S(find(strcmp(model.mets,'s_1212')),strcmp(model.rxns,'r_0939'))=0; %NADPH
model.S(find(strcmp(model.mets,'s_1207')),strcmp(model.rxns,'r_0939'))=0; %NADP
model.S(find(strcmp(model.mets,'s_1203')),strcmp(model.rxns,'r_0939'))=1; %NADH
model.S(find(strcmp(model.mets,'s_1198')),strcmp(model.rxns,'r_0939'))=-1;%NAD

% Make esterification reactions irreversible. positive deltaG
model.ub(strcmp(model.rxns,'r_4713')) = 0; %diethyl succinate
model.ub(strcmp(model.rxns,'r_4714')) = 0; %monoethyl succinate

% Make polyphosphate hydrolase and diphosphate transport over cell membrane
% both irreversible
model = setParam(model,'lb',{'r_4723','r_4724','r_4725'},0);
model = setParam(model,'lb','r_4460',0);

% While r_0013 was elementary balanced, it was not charged balanced. The
% reaction mechanism was incorrect. Corrected to mimic a combination of
% MetaCyc rxns: R83-RXN and R147-RXN; or KEGG rxns: R07364 and R07395.
model = changeRxns(model,'r_0013','5-(methylsulfanyl)-2,3-dioxopentyl phosphate[c] + H2O[c] + oxygen[c] => 4-methylthio-2-oxobutanoate[c] + formate[c] + 2 H+[c] + phosphate[c]',3);

% Represent ACP with formula "RHS"
model.metFormulas(getIndexes(model,'s_1845','mets')) = {'RHS'};

% Set formula of ferr(i/o)cytochrome b3
idx_mit = getIndexes(model,{'s_3826','s_3827'},'mets');
idx_erm = getIndexes(model,{'s_4210','s_4209'},'mets');
model.metFormulas(idx_erm) = model.metFormulas(idx_mit);
model.metCharges(idx_erm) = model.metCharges(idx_mit);
model.metMiriams(idx_erm) = model.metMiriams(idx_mit);
model.metNames([idx_mit; idx_erm]) = regexprep(model.metNames([idx_mit; idx_erm]),'F','f');

% Correct metFormula and metCharge
idx = getIndexes(model,{'s_4265','s_4266'},'mets');
model.metFormulas(idx) = {'CH2O3S'};
model.metCharges(idx) = -1;


% [~,metFormulae] = computeMetFormulae(model,'metMwRange','s_0338','fillMets','none')
% model.metFormulas(getIndexes(model,'s_0329','mets')) = {'C17H28NO17P'};
% model.metFormulas(getIndexes(model,'s_0330','mets')) = {'C33H58NO18P'};
% model.metFormulas(getIndexes(model,'s_0331','mets')) = {'C19H30NO18P'};
% model.metFormulas(getIndexes(model,'s_0334','mets')) = {'C39H68NO23P'};
% model.metFormulas(getIndexes(model,'s_0337','mets')) = {'C44H78N2O27P2'};
% model.metFormulas(getIndexes(model,'s_0338','mets')) = {'C41H38N2O41P3'};
% model.metFormulas(getIndexes(model,'s_0339','mets')) = {'C50H88N2O32P2'};

% Copy annotations between same metabolites in separate compartments
model.metFormulas(getIndexes(model,'s_4211','mets')) = model.metFormulas(getIndexes(model,'s_2885','mets'));
model.metCharges(getIndexes(model,'s_4211','mets')) = model.metCharges(getIndexes(model,'s_2885','mets'));
model.metFormulas(getIndexes(model,'s_4209','mets')) = model.metFormulas(getIndexes(model,'s_3826','mets'));
model.metCharges(getIndexes(model,'s_4209','mets')) = model.metCharges(getIndexes(model,'s_3826','mets'));
model.metMiriams(getIndexes(model,'s_4209','mets')) = model.metMiriams(getIndexes(model,'s_3826','mets'));

% r_4323 is an less precise half-reaction of r_4324 and will be removed
% model = removeReactions(model,'r_4323',true,true,true);

% r_4325 represents scaffolding during [Fe-S]-cluster synthesis, not a
% metabolic process, and will therefore be removed
% model = removeReactions(model,'r_4325',true,true,true);

% r_0229
%model = changeRxns(model,'r_0229','dethiobiotin[c] + polysulphur[c]  <=> biotin[c] + 2 H+[c]',2)
%dethiobiotin[c] + hydrogen sulfide[c] + 2 S-adenosyl-L-methionine[c] + 2 H+[c] <=> biotin[c] + 2 L-methionine[c] + 2 5'-Deoxyadenosine


%% ========================================================================
% Condition-specific gene expression. These can be enabled with scripts
% Glycine cleavage only active when glycine is used as nitrogen source
model = setParam(model,'eq','r_0501',0); %glycine cleavage, mitochondrion
model = setParam(model,'eq','r_0507',0); %glycine cleavage complex (lipoylprotein), mitochondrion
model = setParam(model,'eq','r_0509',0); %glycine cleavage complex (lipoamide), mitochondrion
model.rxnNotes(ismember(model.rxns,{'r_0501','r_0507','r_0509'})) = {'Only active if glycine is nitrogen source'};

% Glutamate synthase repressed in excess nitrogen
model = setParam(model,'eq','r_0472',0);
model.rxnNotes(ismember(model.rxns,{'r_0472'})) = {'Only active during nitrogen limitation'};

% The carnitine shuttle requires exogeneous carnitine, which is absent from
% defined medium.
model = setParam(model,'eq',{'r_0252'},0);
model.rxnNotes(ismember(model.rxns,{'r_0252'})) = {'Only active if growth medium contains carnitine'};

% %% Rescale protein fraction so that biomass sums up to 1 g/gDCW
% % Protein is the largest fraction, so increasing
% [X,P]  = sumBioMass(model, false);
% fprintf('Current biomass adds up to %.4f g/g. Protein fraction is scaled from %.4f to %.4f g/g to reach 1 g/g total biomass.\n', X, P, (1-X)+P)
% model = scaleBioMass(model,'biomass',1,'protein');

% Balance the charge of all biomass component pseudo reactions by adding the required amount of H+
model.S(find(strcmp(model.mets,'s_0794')),strcmp(model.rxns,'r_4047')) = -sum(model.S(:,strcmp(model.rxns,'r_4047')).*model.metCharges,'omitnan'); % Protein
model.S(find(strcmp(model.mets,'s_0794')),strcmp(model.rxns,'r_4049')) = -sum(model.S(:,strcmp(model.rxns,'r_4049')).*model.metCharges,'omitnan'); % RNA
model.S(find(strcmp(model.mets,'s_0794')),strcmp(model.rxns,'r_4050')) = -sum(model.S(:,strcmp(model.rxns,'r_4050')).*model.metCharges,'omitnan'); % DNA
model.S(find(strcmp(model.mets,'s_0794')),strcmp(model.rxns,'r_4598')) = -sum(model.S(:,strcmp(model.rxns,'r_4598')).*model.metCharges,'omitnan'); % Cofactor
model.S(find(strcmp(model.mets,'s_0794')),strcmp(model.rxns,'r_4599')) = -sum(model.S(:,strcmp(model.rxns,'r_4599')).*model.metCharges,'omitnan'); % Ion

% Special case for SLIME rxns
model.metCharges(find(contains(model.metNames,'chain')+contains(model.metNames,'backbone'))) = 0;

% Now, based on the charge balance, find all the reactions that are
% imbalanced, add or remove hydrogen as necessary

% Balance the charge of all imbalanced SLIME reactions by adding the
% required amount of H+,
model.S(find(strcmp(model.mets,'s_0794')),strcmp(model.rxns,'r_3975')) = -sum(model.S(:,strcmp(model.rxns,'r_3975')).*model.metCharges,'omitnan'); % Protein
model.S(find(strcmp(model.mets,'s_0794')),strcmp(model.rxns,'r_3976')) = -sum(model.S(:,strcmp(model.rxns,'r_3976')).*model.metCharges,'omitnan'); % Protein
model.S(find(strcmp(model.mets,'s_0794')),strcmp(model.rxns,'r_3977')) = -sum(model.S(:,strcmp(model.rxns,'r_3977')).*model.metCharges,'omitnan'); % Protein
model.S(find(strcmp(model.mets,'s_0794')),strcmp(model.rxns,'r_3978')) = -sum(model.S(:,strcmp(model.rxns,'r_3978')).*model.metCharges,'omitnan'); % Protein
model.S(find(strcmp(model.mets,'s_0794')),strcmp(model.rxns,'r_4076')) = -sum(model.S(:,strcmp(model.rxns,'r_4076')).*model.metCharges,'omitnan'); % Protein
model.S(find(strcmp(model.mets,'s_0794')),strcmp(model.rxns,'r_4077')) = -sum(model.S(:,strcmp(model.rxns,'r_4077')).*model.metCharges,'omitnan'); % Protein
model.S(find(strcmp(model.mets,'s_0794')),strcmp(model.rxns,'r_4078')) = -sum(model.S(:,strcmp(model.rxns,'r_4078')).*model.metCharges,'omitnan'); % Protein
model.S(find(strcmp(model.mets,'s_0794')),strcmp(model.rxns,'r_4079')) = -sum(model.S(:,strcmp(model.rxns,'r_4079')).*model.metCharges,'omitnan'); % Protein
```
