## Supplementary Appendix 2 for "The peroxiredoxin Tsa1 promotes stationary phase entry by suppressing PKA activity": plot_data_pairs.m

```
clear;
close all;
STRAINS={'T73','T73TSA1','W303','W303TSA1'};

units={'gDW/L', 'mM','A.U.','mM','mM','mM','mM','mM','OD units','g/L','mM'};

YLIMMAX=[2.8 45 1.1 100 1.5 6 1.2 2.5 11 5.5 2.2]

for turn=1:1

    NPLOTi=11;
    NPLOTj=2;

    cols = [
        0.00, 0.45, 0.74;   % Deep Blue
        0.85, 0.33, 0.10;   % Red-Orange
        0.93, 0.69, 0.13;   % Mustard Yellow
        0.49, 0.18, 0.56;   % Purple
        0.47, 0.67, 0.19;   % Green
        0.30, 0.75, 0.93;   % Cyan
        0.64, 0.08, 0.18;   % Dark Red
        0.78, 0.78, 0.78;   % Light Gray
        0.99, 0.41, 0.23;   % Coral
        0.30, 0.30, 0.30;   % Dark Gray
        ];
    counter=1;

    for  data_set=1:4

        if data_set>2
            column=2;
        else
            column=1;
        end

        STRAIN=STRAINS{data_set};
        inputs.plotd.plotlevel='noplot';
        load_data;
        res=AMIGO_SData(inputs);

        st_names=cellstr(inputs.model.st_names);
        obs_names=cellstr(inputs.exps.obs_names{1});


        for k=1:inputs.model.n_st-1

            counter=counter+1;
            subplot(NPLOTi, NPLOTj,(k-1)*2+column );


            plot(res.sim.tsim{1},res.sim.states{1}(:,k),"Color",cols(data_set,:));
            ax = gca;
            title(inputs.model.st_names(k,:));
            hold on;

            if turn==1
                xlim([0 res.sim.tsim{1}(end)]);
            else
                xlim([0 11]);
            end

            hold on;
            result=strfind(obs_names,st_names(k));
            indices = find(~cellfun(@isempty, result));
            if ~isempty(indices)
                a=errorbar(inputs.exps.t_s{1},inputs.exps.exp_data{1}(:,indices),...
                    inputs.exps.error_data{1}(:,indices),'o',"Color",cols(data_set,:),...
                    "LineStyle","none","MarkerFaceColor",cols(data_set,:),'MarkerSize',3);

                ymean=mean(inputs.exps.exp_data{1}(:,indices),'omitnan');
                SSres=sum((inputs.exps.exp_data{1}(:,indices)-res.fit.ms{1}(:,indices)).^2,'omitnan');
                SStot=sum((inputs.exps.exp_data{1}(:,indices)-ymean).^2,'omitnan');
                R2=1-SSres/SStot;

                ylim([0 YLIMMAX(k)]);
                ymin = ax.YLim(1);
                ymax = ax.YLim(2);
                % if data_set==1
                %     text(40,(ymax-ymin)*0.5,sprintf('R_{WT}^2=%.3f',R2));
                % else
                %     text(60,(ymax-ymin)*0.5,sprintf('R_{\\DeltaTSA1}^2=%.3f',R2));
                % end


            end
            ylabel(units{k});
            ylim([0 YLIMMAX(k)]);
        end
        xlabel('Time (hours)')

    end


end
```
