## Supplementary Appendix 2 for "The peroxiredoxin Tsa1 promotes stationary phase entry by suppressing PKA activity": README

# README

This repository contains the code and data required to reproduce the modelling results presented in the accompanying paper. A detailed methodological explanation is provided in the Supplementary Materials.

For questions, comments, or issues related to the code, please contact David Henriques at.

The model curations used here are based on an extended curation of yeast-GEM. The curated model is available at:

https://github.com/SysBioChalmers/yeast-GEM/tree/feat/anaerobic

A manuscript describing the rationale behind these curation choices is currently in preparation. If you use these changes (SJÖBERG et al; in preparation), please cite the associated manuscript once available, as well as the yeast-GEM 9 paper.

The code is divided into several tasks corresponding to the main steps of the modelling workflow. Additional scripts and helper functions used by the main workflows are stored in the `code/` folder.

The main data file is `data_ferm.xlsx`. This Excel file contains fixed parameters, equations, experimental data, and other information required by the model. Fixed parameters can be edited in the `pars` sheet of `data_ferm.xlsx`. Data import is handled by `load_data.m`.

The modelling workflow consists of the following main steps: import experimental data and model information, estimate growth rates from exponential-phase data, estimate the remaining dynamic model parameters, simulate the dynamic model, run dFBA simulations for glucose-based and ethanol-based growth, and plot the parameter adjustment and strain comparisons.

Growth rates are estimated from exponential-phase data using log-transformed biomass data and fixed parameters. This is done with `fit_mu_data.m`.

The remaining model parameters are estimated using `fit_data.m`. This script performs the parameter adjustment of the dynamic model against the experimental data. Choose strain by modifying data_set variable.
Before parameter estimation you will need compile the model. use compile_model.m.

The dynamic model is then used to simulate dFBA and compute flux distributions associated with glucose-based growth and ethanol-based growth. To run the glucose dFBA simulation, use `run_dfba_glc.m`. To run the ethanol dFBA simulation, use `run_dfb_eth.m`.

The resulting flux distributions are stored in `glucoseflux.xlsx` and `ethanolflux.xlsx`. These files contain the dFBA results obtained with the simulated model.

To plot the results of the parameter adjustment, use `plot_data.m`. This script plots all strains in four columns. Alternatively, use `plot_data_pairs.m`, which plots each wild-type strain against the corresponding mutant.

In most scripts, the strain can be selected by changing the `data_set` variable:

STRAINS = {'T73','T73TSA1','W303','W303TSA1'};
STRAIN = STRAINS{data_set};

The available strains are T73, T73TSA1, W303, and W303TSA1. For example, setting data_set = 1 selects T73, while data_set = 2 selects T73TSA1.

The main scripts are as follows. load_data.m imports experimental data, fixed parameters, equations, and other model information from data_ferm.xlsx. fit_mu_data.m estimates growth rates from exponential-phase biomass data. fit_data.m estimates the remaining dynamic model parameters. run_dfba_glc.m runs the dFBA simulation associated with glucose-based growth. run_dfb_eth.m runs the dFBA simulation associated with ethanol-based growth. plot_data.m plots the parameter-adjustment results for all strains in four columns. plot_data_pairs.m plots each wild-type strain against the corresponding mutant.

To run the code, you will need MATLAB, the MEIGO toolbox, the COBRA Toolbox, a Gurobi solver or equivalent, and AMIGO2. The required resources are available from the following links:

MEIGO toolbox: https://github.com/gingproc-IIM-CSIC/MEIGO64
RAVEN Toolbox: https://github.com/SysBioChalmers/RAVEN
Gurobi solver or equivalent: https://www.gurobi.com/
AMIGO2: https://sites.google.com/site/amigo2toolbox

Academic licenses for Gurobi are generally available for free. Please make sure that the required toolboxes are correctly installed and added to the MATLAB path before running the scripts.

If you use this repository, please cite the accompanying paper. If you use the curated yeast-GEM changes, please also cite the associated manuscript (SJÖBERG et al; in preparation) describing the curation choices once it becomes available, as well as the yeast-GEM 9 paper.
