## Supplementary Appendix 2 for "The peroxiredoxin Tsa1 promotes stationary phase entry by suppressing PKA activity": run_dfba_glc.m

```
clear;

for data_set=3:4

    clearvars -except data_set TAB;

    close all;
    addpath('code')
    STRAINS={'T73','T73TSA1','W303','W303TSA1'};
    STRAIN=STRAINS{data_set};
    load_data;

    res=AMIGO_SModel(inputs);

    FLUX=[];
    samples=find(res.sim.tsim{1}>1 & res.sim.tsim{1}<8);
    tol=0.01;

    %% Iterate over to time-points
    for ti=samples

        load(fullfile(pwd,'yeast-GEM.mat'));
        GAM   = 55;
        P     = 0.461;  %Data from Nissen et al. 1997
        NGAM  = 1;
        model = changeGAM(model,GAM,NGAM);
        model = scaleBioMass(model,'protein',P,'carbohydrate',false);
       model_curation;                  % curate base model
        model=other_cons(model);


        t=res.sim.tsim{1}(ti);
        D=0;

        for i=1:inputs.model.n_st
            namevar=regexprep(inputs.model.st_names(i,:),' ','');
            eval([namevar '=res.sim.states{1}(ti,i);']);
        end

        for i=1:inputs.model.n_par
            namepar=regexprep(inputs.model.par_names(i,:),' ','');
            eval([namepar '=inputs.model.par(i);']);
        end

        for i=1:size(inputs.model.eqns,1)
            eqn=regexprep(inputs.model.eqns(i,:),' ','');
            eval([eqn ';']);
        end

        %% last N equations are used for addint the constraints
        for i=1:size(inputs.model.eqns,1)
            eqn=regexprep(inputs.model.eqns(i,:),' ','');
            eval([eqn ';']);
        end

        %% Glucose
        model.lb(findRxnIDs(model,'r_1714'))=dglc./DW;
        model.ub(findRxnIDs(model,'r_1714'))=dglc./DW;

        %% Ethanol
        model.lb(findRxnIDs(model,'r_1761'))=deth./DW*(1-tol);
        model.ub(findRxnIDs(model,'r_1761'))=deth./DW*(1+tol);

        %% succinate UB
        model.lb(findRxnIDs(model,'r_2056'))=dsuccinate./DW*(1-tol);
        model.ub(findRxnIDs(model,'r_2056'))=dsuccinate./DW*(1-tol);

        %% Acetic
        model.lb(findRxnIDs(model,'r_1634'))=dacetate./DW*(1-tol);
        model.ub(findRxnIDs(model,'r_1634'))=dacetate./DW*(1+tol);

        %% Pyruvate
        model.lb(findRxnIDs(model,'r_2033'))=dpyruvate./DW;
        model.ub(findRxnIDs(model,'r_2033'))=dpyruvate./DW;

	    % r_1936	methylglyoxal synthase	dihydroxyacetone phosphate[c] => methylglyoxal[c] + phosphate[c]	
        % kegg.reaction/R01016;metanetx.reaction/MNXR101551;bigg.reaction/MGSA;sbo/SBO:0000176					0
        model = setParam(model,'eq',{'r_1936'},0);
        
        %% Growth
        model.lb(findRxnIDs(model,'r_4041'))=dDW./DW*(1-tol);
        model.ub(findRxnIDs(model,'r_4041'))=dDW./DW*(1+tol);

        %% glycerol exchange
        model.lb(findRxnIDs(model,'r_1808'))=dglycerol./DW*(1-tol);
        model.ub(findRxnIDs(model,'r_1808'))=dglycerol./DW*(1+tol);

        model = setParam(model,'eq',{'r_1936'},0);

        flux=solveLP(model,1);
        if isempty(flux.x)
            error('falied sim');
        end
        FLUX=[FLUX flux.x];

    end

    %% Calculate average fluxes
    DW=res.sim.states{1}(samples,1)';
    v_glc=-trapz(res.sim.tsim{1}(samples),FLUX(findRxnIDs(model,'r_1714'),:).*DW);
    for i=1:size(FLUX,1)
        V(i)=trapz(res.sim.tsim{1}(samples),FLUX(findRxnIDs(model,model.rxns{i}),:).*DW);
        Va(i)=V(i);
        V(i)=V(i)/v_glc;
    end

    model.metNames=strcat(model.metNames, repmat('[',length(model.mets),1),model.compNames(model.metComps),repmat(']',length(model.mets),1) );
    fluxtable=table(model.rxns,model.rxnNames,printRxnFormula(model,'metNameFlag',true,'rxnAbbrList',model.rxns,'printFlag',0),V');
    tab = fluxtable;
    [FLUX flux.x]


    if ~exist('TAB') || isempty(TAB)
        TAB=[tab.Var4]
    else
        TAB=[TAB tab.Var4]
    end

end

%% Package Data
TAB=[TAB abs(mean(TAB')')];
TAB = array2table(TAB, 'VariableNames', compose('Cond%d', 1:size(TAB,2)));
TAB = addvars(TAB, model.rxns, model.rxnNames,printRxnFormula(model,'metNameFlag',true,'rxnAbbrList',model.rxns,'printFlag',0), ...
    'Before', 1, ...
    'NewVariableNames', {'RxnID','RxnName','RxnFormula'});
TAB = sortrows(TAB,'Cond3','descend');
save('RES_GLC_DFBA','TAB');
```
