## Supplementary Appendix 2 for "The peroxiredoxin Tsa1 promotes stationary phase entry by suppressing PKA activity": yeast-GEM.yml

```
---
!!omap
- metaData:
    id: "yeastGEM_develop"
    name: "The Consensus Genome-Scale Metabolic Model of Yeast"
    version: ""
    date: "2024-11-23"
    defaultLB: "-1000"
    defaultUB: "1000"
    givenName: "Eduard"
    familyName: "Kerkhoven"
    email: ""
    organization: "Chalmers University of Technology"
    taxonomy: "taxonomy/559292"
    note: "Saccharomyces cerevisiae - strain S288C"
- metabolites:
    - !!omap
      - id: "s_0001"
      - name: "(1->3)-beta-D-glucan"
      - compartment: "ce"
      - formula: "C6H10O5"
      - charge: 0
      - smiles: "C(C1C(C(C(C(O1)O)O)O)O)O"
      - annotation: !!omap
          - bigg.metabolite: "13BDglcn"
          - chebi: "CHEBI:37671"
          - kegg.compound: "C00965"
          - metanetx.chemical: "MNXM6492"
          - sbo: "SBO:0000247"
      - deltaG: 10000000
    - !!omap
      - id: "s_0002"
      - name: "(1->3)-beta-D-glucan"
      - compartment: "c"
      - formula: "C6H10O5"
      - charge: 0
      - smiles: "C(C1C(C(C(C(O1)O)O)O)O)O"
      - annotation: !!omap
          - bigg.metabolite: "13BDglcn"
          - chebi: "CHEBI:37671"
          - kegg.compound: "C00965"
          - metanetx.chemical: "MNXM6492"
          - sbo: "SBO:0000247"
      - deltaG: 10000000
    - !!omap
      - id: "s_0003"
      - name: "(1->3)-beta-D-glucan"
      - compartment: "e"
      - formula: "C6H10O5"
      - charge: 0
      - smiles: "C(C1C(C(C(C(O1)O)O)O)O)O"
      - annotation: !!omap
          - bigg.metabolite: "13BDglcn"
          - chebi: "CHEBI:37671"
          - kegg.compound: "C00965"
          - metanetx.chemical: "MNXM6492"
          - sbo: "SBO:0000247"
      - deltaG: 10000000
    - !!omap
      - id: "s_0004"
      - name: "(1->6)-beta-D-glucan"
      - compartment: "ce"
      - formula: "C6H10O5"
      - charge: 0
      - smiles: "C(C1C(C(C(C(O1)O)O)O)O)O"
      - annotation: !!omap
          - bigg.metabolite: "16BDglcn"
          - chebi: "CHEBI:27380"
          - kegg.compound: "C02493"
          - metanetx.chemical: "MNXM9375"
          - sbo: "SBO:0000247"
      - deltaG: -229.85
    - !!omap
      - id: "s_0006"
      - name: "(2-amino-4-hydroxy-7,8-dihydropteridin-6-yl)methyl trihydrogen diphosphate"
      - compartment: "m"
      - formula: "C7H8N5O8P2"
      - charge: -3
      - smiles: "C1C(=NC2=C(N1)N=C(NC2=O)N)COP(=O)(O)OP(=O)(O)O"
      - annotation: !!omap
          - bigg.metabolite: "2ahhmd"
          - chebi: "CHEBI:57602"
          - kegg.compound: "C04807"
          - metanetx.chemical: "MNXM1160"
          - sbo: "SBO:0000247"
      - deltaG: -343.18
    - !!omap
      - id: "s_0007"
      - name: "(2E)-3-(methoxycarbonyl)pent-2-enedioic acid"
      - compartment: "c"
      - formula: "C7H6O6"
      - charge: -2
      - smiles: "COC(=O)C(=CC(=O)O)CC(=O)O"
      - annotation: !!omap
          - bigg.metabolite: "aconm"
          - chebi: "CHEBI:57470"
          - kegg.compound: "C11514"
          - metanetx.chemical: "MNXM3742"
          - sbo: "SBO:0000247"
      - deltaG: -160.48
    - !!omap
      - id: "s_0008"
      - name: "(2R,3R)-2,3-dihydroxy-3-methylpentanoate"
      - compartment: "m"
      - formula: "C6H11O4"
      - charge: -1
      - smiles: "CCC(C)(C(C(=O)[O-])O)O"
      - annotation: !!omap
          - bigg.metabolite: "23dhmp"
          - chebi: "CHEBI:49258"
          - kegg.compound: "C06007"
          - metanetx.chemical: "MNXM1202"
          - sbo: "SBO:0000247"
      - deltaG: -45.17
    - !!omap
      - id: "s_0009"
      - name: "(2R,3S)-3-isopropylmalate"
      - compartment: "c"
      - formula: "C7H10O5"
      - charge: -2
      - smiles: "CC(C)C(C(C(=O)[O-])O)C(=O)[O-]"
      - annotation: !!omap
          - bigg.metabolite: "3c2hmp"
          - chebi: "CHEBI:35121"
          - kegg.compound: "C04411"
          - metanetx.chemical: "MNXM891"
          - sbo: "SBO:0000247"
      - deltaG: -103.17
    - !!omap
      - id: "s_0010"
      - name: "(2S)-2-isopropyl-3-oxosuccinate"
      - compartment: "c"
      - formula: "C7H8O5"
      - charge: -2
      - smiles: "CC(C)C(C(=O)C(=O)O)C(=O)O"
      - annotation: !!omap
          - bigg.metabolite: "3c4mop"
          - chebi: "CHEBI:17214"
          - kegg.compound: "C04236"
          - metanetx.chemical: "MNXM1602"
          - sbo: "SBO:0000247"
      - deltaG: -112.16
    - !!omap
      - id: "s_0011"
      - name: "(2S)-2-isopropyl-3-oxosuccinate"
      - compartment: "m"
      - formula: "C7H8O5"
      - charge: -2
      - smiles: "CC(C)C(C(=O)C(=O)O)C(=O)O"
      - annotation: !!omap
          - bigg.metabolite: "3c4mop"
          - chebi: "CHEBI:17214"
          - kegg.compound: "C04236"
          - metanetx.chemical: "MNXM1602"
          - sbo: "SBO:0000247"
      - deltaG: -112.16
    - !!omap
      - id: "s_0012"
      - name: "(2S,3R)-3-hydroxybutane-1,2,3-tricarboxylic acid"
      - compartment: "m"
      - formula: "C7H7O7"
      - charge: -3
      - smiles: "CC(C(CC(=O)O)C(=O)O)(C(=O)O)O"
      - annotation: !!omap
          - bigg.metabolite: "micit"
          - chebi: "CHEBI:57429"
          - kegg.compound: "C04593"
          - metanetx.chemical: "MNXM1694"
          - sbo: "SBO:0000247"
      - deltaG: -209.44
    - !!omap
      - id: "s_0013"
      - name: "(5S,6S)-di-HETE"
      - compartment: "c"
      - formula: "C20H31O4"
      - charge: -1
      - smiles: "CCCCCC=CCC=CC=CC=CC(C(CCCC(=O)O)O)O"
      - annotation: !!omap
          - chebi: "CHEBI:53026"
          - metanetx.chemical: "MNXM31306"
          - sbo: "SBO:0000247"
      - deltaG: 247.78
    - !!omap
      - id: "s_0014"
      - name: "(5S,6S)-di-HETE"
      - compartment: "n"
      - formula: "C20H31O4"
      - charge: -1
      - smiles: "CCCCCC=CCC=CC=CC=CC(C(CCCC(=O)O)O)O"
      - annotation: !!omap
          - chebi: "CHEBI:53026"
          - metanetx.chemical: "MNXM31306"
          - sbo: "SBO:0000247"
      - deltaG: 247.78
    - !!omap
      - id: "s_0015"
      - name: "(N(omega)-L-arginino)succinic acid"
      - compartment: "c"
      - formula: "C10H17N4O6"
      - charge: -1
      - smiles: "C(CC(C(=O)O)N)CN=C(N)NC(CC(=O)O)C(=O)O"
      - annotation: !!omap
          - bigg.metabolite: "argsuc"
          - chebi: "CHEBI:57472"
          - kegg.compound: "C03406"
          - metanetx.chemical: "MNXM722743"
          - sbo: "SBO:0000247"
      - deltaG: -69.25
    - !!omap
      - id: "s_0016"
      - name: "(R)-2,3-dihydroxy-3-methylbutanoate"
      - compartment: "m"
      - formula: "C5H9O4"
      - charge: -1
      - smiles: "CC(C)(C(C(=O)[O-])O)O"
      - annotation: !!omap
          - bigg.metabolite: "23dhmb"
          - chebi: "CHEBI:49072"
          - kegg.compound: "C04272"
          - metanetx.chemical: "MNXM114097"
          - sbo: "SBO:0000247"
      - deltaG: -67.63
    - !!omap
      - id: "s_0017"
      - name: "(R)-4'-phosphopantothenic acid"
      - compartment: "c"
      - formula: "C9H15NO8P"
      - charge: -3
      - smiles: "CC(C)(COP(=O)(O)O)C(C(=O)NCCC(=O)O)O"
      - annotation: !!omap
          - bigg.metabolite: "4ppan"
          - chebi: "CHEBI:15905"
          - kegg.compound: "C03492"
          - metanetx.chemical: "MNXM415"
          - sbo: "SBO:0000247"
      - deltaG: -256.78
    - !!omap
      - id: "s_0018"
      - name: "(R)-5-diphosphomevalonic acid"
      - compartment: "c"
      - formula: "C6H10O10P2"
      - charge: -4
      - smiles: "CC(CCOP(=O)(O)OP(=O)(O)O)(CC(=O)O)O"
      - annotation: !!omap
          - bigg.metabolite: "5dpmev"
          - chebi: "CHEBI:57557"
          - kegg.compound: "C01143"
          - metanetx.chemical: "MNXM689"
          - sbo: "SBO:0000247"
      - deltaG: -473.82
    - !!omap
      - id: "s_0019"
      - name: "(R)-5-phosphomevalonic acid"
      - compartment: "c"
      - formula: "C6H10O7P"
      - charge: -3
      - smiles: "CC(CCOP(=O)(O)O)(CC(=O)O)O"
      - annotation: !!omap
          - bigg.metabolite: "5pmev"
          - chebi: "CHEBI:58146"
          - kegg.compound: "C01107"
          - metanetx.chemical: "MNXM567"
          - sbo: "SBO:0000247"
      - deltaG: -274.7
    - !!omap
      - id: "s_0020"
      - name: "(R)-acetoin"
      - compartment: "c"
      - formula: "C4H8O2"
      - charge: 0
      - smiles: "CC(C(=O)C)O"
      - annotation: !!omap
          - bigg.metabolite: "actn__R"
          - chebi: "CHEBI:15686"
          - kegg.compound: "C00810"
          - metanetx.chemical: "MNXM664"
          - sbo: "SBO:0000247"
      - deltaG: 5.77
    - !!omap
      - id: "s_0021"
      - name: "(R)-carnitine"
      - compartment: "c"
      - formula: "C7H15NO3"
      - charge: 0
      - smiles: "C[N+](C)(C)CC(CC(=O)[O-])O"
      - annotation: !!omap
          - bigg.metabolite: "crn"
          - chebi: "CHEBI:16347"
          - kegg.compound: "C00318"
          - metanetx.chemical: "MNXM173"
          - sbo: "SBO:0000247"
      - deltaG: 76.02
    - !!omap
      - id: "s_0022"
      - name: "(R)-carnitine"
      - compartment: "e"
      - formula: "C7H15NO3"
      - charge: 0
      - smiles: "C[N+](C)(C)CC(CC(=O)[O-])O"
      - annotation: !!omap
          - bigg.metabolite: "crn"
          - chebi: "CHEBI:16347"
          - kegg.compound: "C00318"
          - metanetx.chemical: "MNXM173"
          - sbo: "SBO:0000247"
      - deltaG: 76.02
    - !!omap
      - id: "s_0023"
      - name: "(R)-carnitine"
      - compartment: "m"
      - formula: "C7H15NO3"
      - charge: 0
      - smiles: "C[N+](C)(C)CC(CC(=O)[O-])O"
      - annotation: !!omap
          - bigg.metabolite: "crn"
          - chebi: "CHEBI:16347"
          - kegg.compound: "C00318"
          - metanetx.chemical: "MNXM173"
          - sbo: "SBO:0000247"
      - deltaG: 76.02
    - !!omap
      - id: "s_0024"
      - name: "(R)-carnitine"
      - compartment: "p"
      - formula: "C7H15NO3"
      - charge: 0
      - smiles: "C[N+](C)(C)CC(CC(=O)[O-])O"
      - annotation: !!omap
          - bigg.metabolite: "crn"
          - chebi: "CHEBI:16347"
          - kegg.compound: "C00318"
          - metanetx.chemical: "MNXM173"
          - sbo: "SBO:0000247"
      - deltaG: 76.02
    - !!omap
      - id: "s_0025"
      - name: "(R)-lactate"
      - compartment: "c"
      - formula: "C3H5O3"
      - charge: -1
      - smiles: "CC(C(=O)O)O"
      - annotation: !!omap
          - bigg.metabolite: "lac__D"
          - chebi: "CHEBI:16004"
          - kegg.compound: "C00256"
          - metanetx.chemical: "MNXM285"
          - sbo: "SBO:0000247"
      - deltaG: -74.68
    - !!omap
      - id: "s_0026"
      - name: "(R)-lactate"
      - compartment: "e"
      - formula: "C3H5O3"
      - charge: -1
      - smiles: "CC(C(=O)O)O"
      - annotation: !!omap
          - bigg.metabolite: "lac__D"
          - chebi: "CHEBI:16004"
          - kegg.compound: "C00256"
          - metanetx.chemical: "MNXM285"
          - sbo: "SBO:0000247"
      - deltaG: -74.68
    - !!omap
      - id: "s_0027"
      - name: "(R)-lactate"
      - compartment: "m"
      - formula: "C3H5O3"
      - charge: -1
      - smiles: "CC(C(=O)O)O"
      - annotation: !!omap
          - bigg.metabolite: "lac__D"
          - chebi: "CHEBI:16004"
          - kegg.compound: "C00256"
          - metanetx.chemical: "MNXM285"
          - sbo: "SBO:0000247"
      - deltaG: -74.68
    - !!omap
      - id: "s_0028"
      - name: "(R)-mevalonate"
      - compartment: "c"
      - formula: "C6H11O4"
      - charge: -1
      - smiles: "CC(CCO)(CC(=O)[O-])O"
      - annotation: !!omap
          - bigg.metabolite: "mev__R"
          - chebi: "CHEBI:36464"
          - kegg.compound: "C00418"
          - metanetx.chemical: "MNXM333"
          - sbo: "SBO:0000247"
      - deltaG: -50.86
    - !!omap
      - id: "s_0029"
      - name: "(R)-mevalonate"
      - compartment: "e"
      - formula: "C6H11O4"
      - charge: -1
      - smiles: "CC(CCO)(CC(=O)[O-])O"
      - annotation: !!omap
          - bigg.metabolite: "mev__R"
          - chebi: "CHEBI:36464"
          - kegg.compound: "C00418"
          - metanetx.chemical: "MNXM333"
          - sbo: "SBO:0000247"
      - deltaG: -50.86
    - !!omap
      - id: "s_0030"
      - name: "(R)-pantoate"
      - compartment: "c"
      - formula: "C6H11O4"
      - charge: -1
      - smiles: "CC(C)(CO)C(C(=O)[O-])O"
      - annotation: !!omap
          - bigg.metabolite: "pant__R"
          - chebi: "CHEBI:15980"
          - kegg.compound: "C00522"
          - metanetx.chemical: "MNXM593"
          - sbo: "SBO:0000247"
      - deltaG: -52.67
    - !!omap
      - id: "s_0031"
      - name: "(R)-pantothenate"
      - compartment: "c"
      - formula: "C9H16NO5"
      - charge: -1
      - smiles: "CC(C)(CO)C(C(=O)NCCC(=O)O)O"
      - annotation: !!omap
          - bigg.metabolite: "pnto__R"
          - chebi: "CHEBI:29032"
          - kegg.compound: "C00864"
          - metanetx.chemical: "MNXM364"
          - sbo: "SBO:0000247"
      - deltaG: -32.94
    - !!omap
      - id: "s_0032"
      - name: "(R)-pantothenate"
      - compartment: "e"
      - formula: "C9H16NO5"
      - charge: -1
      - smiles: "CC(C)(CO)C(C(=O)NCCC(=O)O)O"
      - annotation: !!omap
          - bigg.metabolite: "pnto__R"
          - chebi: "CHEBI:29032"
          - kegg.compound: "C00864"
          - metanetx.chemical: "MNXM364"
          - sbo: "SBO:0000247"
      - deltaG: -32.94
    - !!omap
      - id: "s_0033"
      - name: "(R)-S-lactoylglutathione"
      - compartment: "c"
      - formula: "C13H20N3O8S"
      - charge: -1
      - smiles: "CC(C(=O)SCC(C(=O)NCC(=O)O)NC(=O)CCC(C(=O)O)N)O"
      - annotation: !!omap
          - bigg.metabolite: "lgt__S"
          - chebi: "CHEBI:57474"
          - kegg.compound: "C03451"
          - metanetx.chemical: "MNXM1253"
          - sbo: "SBO:0000247"
      - deltaG: -102.35
    - !!omap
      - id: "s_0034"
      - name: "(R)-S-lactoylglutathione"
      - compartment: "m"
      - formula: "C13H20N3O8S"
      - charge: -1
      - smiles: "CC(C(=O)SCC(C(=O)NCC(=O)O)NC(=O)CCC(C(=O)O)N)O"
      - annotation: !!omap
          - bigg.metabolite: "lgt__S"
          - chebi: "CHEBI:57474"
          - kegg.compound: "C03451"
          - metanetx.chemical: "MNXM1253"
          - sbo: "SBO:0000247"
      - deltaG: -102.35
    - !!omap
      - id: "s_0035"
      - name: "(R,R)-2,3-butanediol"
      - compartment: "c"
      - formula: "C4H10O2"
      - charge: 0
      - smiles: "CC(C(C)O)O"
      - annotation: !!omap
          - bigg.metabolite: "btd_RR"
          - chebi: "CHEBI:16982"
          - kegg.compound: "C03044"
          - metanetx.chemical: "MNXM2757"
          - sbo: "SBO:0000247"
      - deltaG: 17.23
    - !!omap
      - id: "s_0036"
      - name: "(R,R)-2,3-butanediol"
      - compartment: "e"
      - formula: "C4H10O2"
      - charge: 0
      - smiles: "CC(C(C)O)O"
      - annotation: !!omap
          - bigg.metabolite: "btd_RR"
          - chebi: "CHEBI:16982"
          - kegg.compound: "C03044"
          - metanetx.chemical: "MNXM2757"
          - sbo: "SBO:0000247"
      - deltaG: 17.23
    - !!omap
      - id: "s_0037"
      - name: "(S)-2,3-epoxysqualene"
      - compartment: "c"
      - formula: "C30H50O"
      - charge: 0
      - smiles: "CC(=CCCC(=CCCC(=CCCC=C(C)CCC=C(C)CCC1C(O1)(C)C)C)C)C"
      - annotation: !!omap
          - bigg.metabolite: "Ssq23epx"
          - chebi: "CHEBI:15441"
          - kegg.compound: "C01054"
          - metanetx.chemical: "MNXM130"
          - sbo: "SBO:0000247"
      - deltaG: 606.15
    - !!omap
      - id: "s_0038"
      - name: "(S)-2,3-epoxysqualene"
      - compartment: "er"
      - formula: "C30H50O"
      - charge: 0
      - smiles: "CC(=CCCC(=CCCC(=CCCC=C(C)CCC=C(C)CCC1C(O1)(C)C)C)C)C"
      - annotation: !!omap
          - bigg.metabolite: "Ssq23epx"
          - chebi: "CHEBI:15441"
          - kegg.compound: "C01054"
          - metanetx.chemical: "MNXM130"
          - sbo: "SBO:0000247"
      - deltaG: 606.15
    - !!omap
      - id: "s_0039"
      - name: "(S)-2-acetyl-2-hydroxybutanoate"
      - compartment: "m"
      - formula: "C6H9O4"
      - charge: -1
      - smiles: "CCC(C(=O)C)(C(=O)[O-])O"
      - annotation: !!omap
          - bigg.metabolite: "2ahbut"
          - chebi: "CHEBI:49256"
          - kegg.compound: "C06006"
          - metanetx.chemical: "MNXM114220"
          - sbo: "SBO:0000247"
      - deltaG: -57.99
    - !!omap
      - id: "s_0042"
      - name: "(R)-3-hydroxydecanoyl-CoA"
      - compartment: "p"
      - formula: "C31H50N7O18P3S"
      - charge: -4
      - smiles: "CCCCCCCC(CC(=O)SCCNC(=O)CCNC(=O)C(C(C)(C)COP(=O)(O)OP(=O)(O)OCC1C(C(C(O1)N2C=NC3=C(N=CN=C32)N)O)OP(=O)(O)O)O)O"
      - annotation: !!omap
          - bigg.metabolite: "3hdcoa"
          - chebi: "CHEBI:28325"
          - kegg.compound: "C05264"
          - metanetx.chemical: "MNXM674"
          - sbo: "SBO:0000247"
      - deltaG: -262.65
    - !!omap
      - id: "s_0045"
      - name: "(S)-3-hydroxyhexacosanoyl-CoA"
      - compartment: "p"
      - formula: "C47H82N7O18P3S"
      - charge: -4
      - smiles: "CCCCCCCCCCCCCCCCCCCCCCCC(CC(=O)SCCNC(=O)CCNC(=O)C(C(C)(C)COP(=O)(O)OP(=O)(O)OCC1C(C(C(O1)N2C=NC3=C(N=CN=C32)N)O)OP(=O)(O)O)O)O"
      - annotation: !!omap
          - chebi: "CHEBI:52976"
          - metanetx.chemical: "MNXM31741"
          - sbo: "SBO:0000247"
      - deltaG: 121.01
    - !!omap
      - id: "s_0048"
      - name: "(R)-3-hydroxylauroyl-CoA"
      - compartment: "p"
      - formula: "C33H54N7O18P3S"
      - charge: -4
      - smiles: "CCCCCCCCCC(CC(=O)SCCNC(=O)CCNC(=O)C(C(C)(C)COP(=O)(O)OP(=O)(O)OCC1C(C(C(O1)N2C=NC3=C(N=CN=C32)N)O)OP(=O)(O)O)O)O"
      - annotation: !!omap
          - bigg.metabolite: "3hddcoa"
          - chebi: "CHEBI:27668"
          - kegg.compound: "C05262"
          - metanetx.chemical: "MNXM733"
          - sbo: "SBO:0000247"
      - deltaG: -214.69
    - !!omap
      - id: "s_0051"
      - name: "(S)-3-hydroxypalmitoyl-CoA"
      - compartment: "p"
      - formula: "C37H62N7O18P3S"
      - charge: -4
      - smiles: "CCCCCCCCCCCCCC(CC(=O)SCCNC(=O)CCNC(=O)C(C(C)(C)COP(=O)(O)OP(=O)(O)OCC1C(C(C(O1)N2C=NC3=C(N=CN=C32)N)O)OP(=O)(O)O)O)O"
      - annotation: !!omap
          - bigg.metabolite: "3hhdcoa"
          - chebi: "CHEBI:27402"
          - kegg.compound: "C05258"
          - metanetx.chemical: "MNXM825"
          - sbo: "SBO:0000247"
      - deltaG: -118.77
    - !!omap
      - id: "s_0054"
      - name: "(S)-3-hydroxytetradecanoyl-CoA"
      - compartment: "p"
      - formula: "C35H58N7O18P3S"
      - charge: -4
      - smiles: "CCCCCCCCCCCC(CC(=O)SCCNC(=O)CCNC(=O)C(C(C)(C)COP(=O)(O)OP(=O)(O)OCC1C(C(C(O1)N2C=NC3=C(N=CN=C32)N)O)OP(=O)(O)O)O)O"
      - annotation: !!omap
          - bigg.metabolite: "3htdcoa"
          - chebi: "CHEBI:27466"
          - kegg.compound: "C05260"
          - metanetx.chemical: "MNXM767"
          - sbo: "SBO:0000247"
      - deltaG: -166.73
    - !!omap
      - id: "s_0056"
      - name: "(S)-3-methyl-2-oxopentanoate"
      - compartment: "c"
      - formula: "C6H9O3"
      - charge: -1
      - smiles: "CCC(C)C(=O)C(=O)O"
      - annotation: !!omap
          - bigg.metabolite: "3mop"
          - chebi: "CHEBI:35146"
          - kegg.compound: "C00671"
          - metanetx.chemical: "MNXM439"
          - sbo: "SBO:0000247"
      - deltaG: -22.2
    - !!omap
      - id: "s_0058"
      - name: "(S)-3-methyl-2-oxopentanoate"
      - compartment: "e"
      - formula: "C6H9O3"
      - charge: -1
      - smiles: "CCC(C)C(=O)C(=O)O"
      - annotation: !!omap
          - bigg.metabolite: "3mop"
          - chebi: "CHEBI:35146"
          - kegg.compound: "C00671"
          - metanetx.chemical: "MNXM439"
          - sbo: "SBO:0000247"
      - deltaG: -22.2
    - !!omap
      - id: "s_0060"
      - name: "(S)-3-methyl-2-oxopentanoate"
      - compartment: "m"
      - formula: "C6H9O3"
      - charge: -1
      - smiles: "CCC(C)C(=O)C(=O)O"
      - annotation: !!omap
          - bigg.metabolite: "3mop"
          - chebi: "CHEBI:35146"
          - kegg.compound: "C00671"
          - metanetx.chemical: "MNXM439"
          - sbo: "SBO:0000247"
      - deltaG: -22.2
    - !!omap
      - id: "s_0061"
      - name: "(S)-dihydroorotate"
      - compartment: "c"
      - formula: "C5H5N2O4"
      - charge: -1
      - smiles: "C1C(NC(=O)NC1=O)C(=O)O"
      - annotation: !!omap
          - bigg.metabolite: "dhor__S"
          - chebi: "CHEBI:30864"
          - kegg.compound: "C00337"
          - metanetx.chemical: "MNXM252"
          - sbo: "SBO:0000247"
      - deltaG: -103.03
    - !!omap
      - id: "s_0062"
      - name: "(S)-lactaldehyde"
      - compartment: "c"
      - formula: "C3H6O2"
      - charge: 0
      - smiles: "CC(C=O)O"
      - annotation: !!omap
          - bigg.metabolite: "lald__L"
          - chebi: "CHEBI:18041"
          - kegg.compound: "C00424"
          - metanetx.chemical: "MNXM2387"
          - sbo: "SBO:0000247"
      - deltaG: -12.05
    - !!omap
      - id: "s_0063"
      - name: "(S)-lactate"
      - compartment: "c"
      - formula: "C3H5O3"
      - charge: -1
      - smiles: "CC(C(=O)O)O"
      - annotation: !!omap
          - bigg.metabolite: "lac__L"
          - chebi: "CHEBI:16651"
          - kegg.compound: "C00186"
          - metanetx.chemical: "MNXM179"
          - sbo: "SBO:0000247"
      - deltaG: -74.68
    - !!omap
      - id: "s_0064"
      - name: "(S)-lactate"
      - compartment: "e"
      - formula: "C3H5O3"
      - charge: -1
      - smiles: "CC(C(=O)O)O"
      - annotation: !!omap
          - bigg.metabolite: "lac__L"
          - chebi: "CHEBI:16651"
          - kegg.compound: "C00186"
          - metanetx.chemical: "MNXM179"
          - sbo: "SBO:0000247"
      - deltaG: -74.68
    - !!omap
      - id: "s_0065"
      - name: "(S)-lactate"
      - compartment: "m"
      - formula: "C3H5O3"
      - charge: -1
      - smiles: "CC(C(=O)O)O"
      - annotation: !!omap
          - bigg.metabolite: "lac__L"
          - chebi: "CHEBI:16651"
          - kegg.compound: "C00186"
          - metanetx.chemical: "MNXM179"
          - sbo: "SBO:0000247"
      - deltaG: -74.68
    - !!omap
      - id: "s_0066"
      - name: "(S)-malate"
      - compartment: "c"
      - formula: "C4H4O5"
      - charge: -2
      - smiles: "C(C(C(=O)O)O)C(=O)O"
      - annotation: !!omap
          - bigg.metabolite: "mal__L"
          - chebi: "CHEBI:15589"
          - kegg.compound: "C00149"
          - metanetx.chemical: "MNXM98"
          - sbo: "SBO:0000247"
      - deltaG: -162.83
    - !!omap
      - id: "s_0067"
      - name: "(S)-malate"
      - compartment: "e"
      - formula: "C4H4O5"
      - charge: -2
      - smiles: "C(C(C(=O)O)O)C(=O)O"
      - annotation: !!omap
          - bigg.metabolite: "mal__L"
          - chebi: "CHEBI:15589"
          - kegg.compound: "C00149"
          - metanetx.chemical: "MNXM98"
          - sbo: "SBO:0000247"
      - deltaG: -162.83
    - !!omap
      - id: "s_0068"
      - name: "(S)-malate"
      - compartment: "m"
      - formula: "C4H4O5"
      - charge: -2
      - smiles: "C(C(C(=O)O)O)C(=O)O"
      - annotation: !!omap
          - bigg.metabolite: "mal__L"
          - chebi: "CHEBI:15589"
          - kegg.compound: "C00149"
          - metanetx.chemical: "MNXM98"
          - sbo: "SBO:0000247"
      - deltaG: -162.83
    - !!omap
      - id: "s_0069"
      - name: "(S)-malate"
      - compartment: "p"
      - formula: "C4H4O5"
      - charge: -2
      - smiles: "C(C(C(=O)O)O)C(=O)O"
      - annotation: !!omap
          - bigg.metabolite: "mal__L"
          - chebi: "CHEBI:15589"
          - kegg.compound: "C00149"
          - metanetx.chemical: "MNXM98"
          - sbo: "SBO:0000247"
      - deltaG: -162.83
    - !!omap
      - id: "s_0075"
      - name: "1,3-bisphospho-D-glycerate"
      - compartment: "c"
      - formula: "C3H4O10P2"
      - charge: -4
      - smiles: "C(C(C(=O)OP(=O)(O)O)O)OP(=O)(O)O"
      - annotation: !!omap
          - bigg.metabolite: "13dpg"
          - chebi: "CHEBI:57604"
          - kegg.compound: "C00236"
          - metanetx.chemical: "MNXM261"
          - sbo: "SBO:0000247"
      - deltaG: -550.97
    - !!omap
      - id: "s_0076"
      - name: "1-(2-carboxyphenylamino)-1-deoxy-D-ribulose 5-phosphate"
      - compartment: "c"
      - formula: "C12H13NO9P"
      - charge: -3
      - smiles: "C1=CC=C(C(=C1)C(=O)O)NCC(=O)C(C(COP(=O)(O)O)O)O"
      - annotation: !!omap
          - bigg.metabolite: "2cpr5p"
          - chebi: "CHEBI:58613"
          - kegg.compound: "C01302"
          - metanetx.chemical: "MNXM1455"
          - sbo: "SBO:0000247"
      - deltaG: -267.72
    - !!omap
      - id: "s_0077"
      - name: "1-(5-phospho-D-ribosyl)-5-[(5-phospho-D-ribosylamino)methylideneamino]imidazole-4-carboxamide"
      - compartment: "c"
      - formula: "C15H21N5O15P2"
      - charge: -4
      - annotation: !!omap
          - bigg.metabolite: "prfp"
          - chebi: "CHEBI:58435"
          - kegg.compound: "C04896"
          - metanetx.chemical: "MNXM1397"
          - sbo: "SBO:0000247"
      - deltaG: -456.95
    - !!omap
      - id: "s_0078"
      - name: "1-(5-phosphoribosyl)-5'-AMP"
      - compartment: "c"
      - formula: "C15H19N5O14P2"
      - charge: -4
      - annotation: !!omap
          - bigg.metabolite: "prbamp"
          - chebi: "CHEBI:59457"
          - kegg.compound: "C02741"
          - metanetx.chemical: "MNXM1515"
          - sbo: "SBO:0000247"
      - deltaG: -421.02
    - !!omap
      - id: "s_0079"
      - name: "1-(sn-glycero-3-phospho)-1D-myo-inositol"
      - compartment: "c"
      - formula: "C9H18O11P"
      - charge: -1
      - smiles: "C(C(COP(=O)(O)OC1C(C(C(C(C1O)O)O)O)O)O)O"
      - annotation: !!omap
          - bigg.metabolite: "g3pi"
          - chebi: "CHEBI:58444"
          - kegg.compound: "C01225"
          - metanetx.chemical: "MNXM1517"
          - sbo: "SBO:0000247"
      - deltaG: -312.07
    - !!omap
      - id: "s_0080"
      - name: "1-(sn-glycero-3-phospho)-1D-myo-inositol"
      - compartment: "e"
      - formula: "C9H18O11P"
      - charge: -1
      - smiles: "C(C(COP(=O)(O)OC1C(C(C(C(C1O)O)O)O)O)O)O"
      - annotation: !!omap
          - bigg.metabolite: "g3pi"
          - chebi: "CHEBI:58444"
          - kegg.compound: "C01225"
          - metanetx.chemical: "MNXM1517"
          - sbo: "SBO:0000247"
      - deltaG: -312.07
    - !!omap
      - id: "s_0084"
      - name: "1-acylglycerophosphocholine"
      - compartment: "e"
      - formula: "C9H20NO7PR"
      - charge: 1
      - annotation: !!omap
          - bigg.metabolite: "pchol_cho"
          - chebi: "CHEBI:11230"
          - kegg.compound: "C04230"
          - metanetx.chemical: "MNXM96952"
          - sbo: "SBO:0000247"
      - deltaG: 10000000
    - !!omap
      - id: "s_0086"
      - name: "1-C-(indol-3-yl)glycerol 3-phosphate"
      - compartment: "c"
      - formula: "C11H12NO6P"
      - charge: -2
      - smiles: "C1=CC=C2C(=C1)C(=CN2)C(C(COP(=O)(O)O)O)O"
      - annotation: !!omap
          - bigg.metabolite: "3ig3p"
          - chebi: "CHEBI:60820"
          - kegg.compound: "C03506"
          - metanetx.chemical: "MNXM163633"
          - sbo: "SBO:0000247"
      - deltaG: -141.84
    - !!omap
      - id: "s_0087"
      - name: "1-methylnicotinamide"
      - compartment: "c"
      - formula: "C7H9N2O"
      - charge: 1
      - smiles: "C[N+]1=CC=CC(=C1)C(=O)N"
      - annotation: !!omap
          - bigg.metabolite: "1mncam"
          - chebi: "CHEBI:16797"
          - kegg.compound: "C02918"
          - metanetx.chemical: "MNXM2172"
          - sbo: "SBO:0000247"
      - deltaG: 98.67
    - !!omap
      - id: "s_0089"
      - name: "1-phosphatidyl-1D-myo-inositol backbone"
      - compartment: "c"
      - formula: "C9H15O9P"
      - charge: -1
      - annotation: !!omap
          - bigg.metabolite: "pail_cho"
          - chebi: "CHEBI:57880"
          - kegg.compound: "C01194"
          - metanetx.chemical: "MNXM62"
          - sbo: "SBO:0000649"
      - deltaG: 10000000
    - !!omap
      - id: "s_0116"
      - name: "1-pyrroline-3-hydroxy-5-carboxylic acid"
      - compartment: "c"
      - formula: "C5H6NO3"
      - charge: -1
      - smiles: "C1C(C=NC1C(=O)O)O"
      - annotation: !!omap
          - bigg.metabolite: "1p3h5c"
          - chebi: "CHEBI:58509"
          - kegg.compound: "C04281"
          - metanetx.chemical: "MNXM114091"
          - sbo: "SBO:0000247"
      - deltaG: -40.88
    - !!omap
      - id: "s_0117"
      - name: "1-pyrroline-3-hydroxy-5-carboxylic acid"
      - compartment: "m"
      - formula: "C5H6NO3"
      - charge: -1
      - smiles: "C1C(C=NC1C(=O)O)O"
      - annotation: !!omap
          - bigg.metabolite: "1p3h5c"
          - chebi: "CHEBI:58509"
          - kegg.compound: "C04281"
          - metanetx.chemical: "MNXM114091"
          - sbo: "SBO:0000247"
      - deltaG: -40.88
    - !!omap
      - id: "s_0118"
      - name: "1-pyrroline-5-carboxylate"
      - compartment: "c"
      - formula: "C5H6NO2"
      - charge: -1
      - smiles: "C1CC(N=C1)C(=O)O"
      - annotation: !!omap
          - bigg.metabolite: "1pyr5c"
          - chebi: "CHEBI:15893"
          - kegg.compound: "C04322"
          - metanetx.chemical: "MNXM1617"
          - sbo: "SBO:0000247"
      - deltaG: -1.04
    - !!omap
      - id: "s_0119"
      - name: "1-pyrroline-5-carboxylate"
      - compartment: "m"
      - formula: "C5H6NO2"
      - charge: -1
      - smiles: "C1CC(N=C1)C(=O)O"
      - annotation: !!omap
          - bigg.metabolite: "1pyr5c"
          - chebi: "CHEBI:15893"
          - kegg.compound: "C04322"
          - metanetx.chemical: "MNXM1617"
          - sbo: "SBO:0000247"
      - deltaG: -1.04
    - !!omap
      - id: "s_0120"
      - name: "10-formyl-THF"
      - compartment: "c"
      - formula: "C20H21N7O7"
      - charge: -2
      - smiles: "C1C(NC2=C(N1)N=C(NC2=O)N)CN(C=O)C3=CC=C(C=C3)C(=O)NC(CCC(=O)O)C(=O)O"
      - annotation: !!omap
          - bigg.metabolite: "10fthf"
          - chebi: "CHEBI:57454"
          - kegg.compound: "C00234"
          - metanetx.chemical: "MNXM237"
          - sbo: "SBO:0000247"
      - deltaG: 73.23
    - !!omap
      - id: "s_0121"
      - name: "10-formyl-THF"
      - compartment: "m"
      - formula: "C20H21N7O7"
      - charge: -2
      - smiles: "C1C(NC2=C(N1)N=C(NC2=O)N)CN(C=O)C3=CC=C(C=C3)C(=O)NC(CCC(=O)O)C(=O)O"
      - annotation: !!omap
          - bigg.metabolite: "10fthf"
          - chebi: "CHEBI:57454"
          - kegg.compound: "C00234"
          - metanetx.chemical: "MNXM237"
          - sbo: "SBO:0000247"
      - deltaG: 73.23
    - !!omap
      - id: "s_0122"
      - name: "14-demethyllanosterol"
      - compartment: "c"
      - formula: "C29H48O"
      - charge: 0
      - smiles: "CC(CCC=C(C)C)C1CCC2C1(CCC3=C2CCC4C3(CCC(C4(C)C)O)C)C"
      - annotation: !!omap
          - bigg.metabolite: "44mzym"
          - chebi: "CHEBI:18364"
          - kegg.compound: "C05108"
          - metanetx.chemical: "MNXM913"
          - sbo: "SBO:0000247"
      - deltaG: 510.25
    - !!omap
      - id: "s_0123"
      - name: "1D-myo-inositol 1,3,4,5-tetrakisphosphate"
      - compartment: "n"
      - formula: "C6H8O18P4"
      - charge: -8
      - smiles: "C1(C(C(C(C(C1OP(=O)(O)O)OP(=O)(O)O)OP(=O)(O)O)O)OP(=O)(O)O)O"
      - annotation: !!omap
          - bigg.metabolite: "mi1345p"
          - chebi: "CHEBI:57895"
          - kegg.compound: "C01272"
          - metanetx.chemical: "MNXM624"
          - sbo: "SBO:0000247"
      - deltaG: -965
    - !!omap
      - id: "s_0124"
      - name: "1D-myo-inositol 1,4,5,6-tetrakisphosphate"
      - compartment: "n"
      - formula: "C6H8O18P4"
      - charge: -8
      - smiles: "C1(C(C(C(C(C1OP(=O)(O)O)OP(=O)(O)O)OP(=O)(O)O)OP(=O)(O)O)O)O"
      - annotation: !!omap
          - bigg.metabolite: "mi3456p"
          - chebi: "CHEBI:57627"
          - kegg.compound: "C11555"
          - metanetx.chemical: "MNXM1620"
          - sbo: "SBO:0000247"
      - deltaG: -965
    - !!omap
      - id: "s_0125"
      - name: "1D-myo-inositol 1,4,5-trisphosphate"
      - compartment: "n"
      - formula: "C6H9O15P3"
      - charge: -6
      - smiles: "C1(C(C(C(C(C1OP(=O)(O)O)O)OP(=O)(O)O)OP(=O)(O)O)O)O"
      - annotation: !!omap
          - bigg.metabolite: "mi145p"
          - chebi: "CHEBI:203600"
          - kegg.compound: "C01245"
          - metanetx.chemical: "MNXM200"
          - sbo: "SBO:0000247"
      - deltaG: -762.1
    - !!omap
      - id: "s_0126"
      - name: "1D-myo-inositol 1-phosphate"
      - compartment: "c"
      - formula: "C6H11O9P"
      - charge: -2
      - smiles: "C1(C(C(C(C(C1O)O)OP(=O)(O)O)O)O)O"
      - annotation: !!omap
          - bigg.metabolite: "mi1p__D"
          - chebi: "CHEBI:58433"
          - kegg.compound: "C01177"
          - metanetx.chemical: "MNXM646"
          - sbo: "SBO:0000247"
      - deltaG: -326.15
    - !!omap
      - id: "s_0127"
      - name: "1D-myo-inositol 1-phosphate"
      - compartment: "er"
      - formula: "C6H11O9P"
      - charge: -2
      - smiles: "C1(C(C(C(C(C1O)O)OP(=O)(O)O)O)O)O"
      - annotation: !!omap
          - bigg.metabolite: "mi1p__D"
          - chebi: "CHEBI:58433"
          - kegg.compound: "C01177"
          - metanetx.chemical: "MNXM646"
          - sbo: "SBO:0000247"
      - deltaG: -326.15
    - !!omap
      - id: "s_0129"
      - name: "1D-myo-inositol 3-phosphate"
      - compartment: "m"
      - formula: "C6H11O9P"
      - charge: -2
      - smiles: "C1(C(C(C(C(C1O)O)OP(=O)(O)O)O)O)O"
      - annotation: !!omap
          - bigg.metabolite: "mi3p__D"
          - chebi: "CHEBI:58401"
          - kegg.compound: "C04006"
          - metanetx.chemical: "MNXM540"
          - sbo: "SBO:0000247"
      - deltaG: -318.64
    - !!omap
      - id: "s_0131"
      - name: "2',3'-cyclic AMP"
      - compartment: "c"
      - formula: "C10H11N5O6P"
      - charge: -1
      - smiles: "C1=NC(=C2C(=N1)N(C=N2)C3C4C(C(O3)CO)OP(=O)(O4)O)N"
      - annotation: !!omap
          - bigg.metabolite: "23camp"
          - chebi: "CHEBI:60879"
          - kegg.compound: "C02353"
          - metanetx.chemical: "MNXM2598"
          - sbo: "SBO:0000247"
      - deltaG: -95.39
    - !!omap
      - id: "s_0132"
      - name: "2'-deoxyadenosine"
      - compartment: "c"
      - formula: "C10H13N5O3"
      - charge: 0
      - smiles: "C1C(C(OC1N2C=NC3=C(N=CN=C32)N)CO)O"
      - annotation: !!omap
          - bigg.metabolite: "dad_2"
          - chebi: "CHEBI:17256"
          - kegg.compound: "C00559"
          - metanetx.chemical: "MNXM625"
          - sbo: "SBO:0000247"
      - deltaG: 111.81
    - !!omap
      - id: "s_0133"
      - name: "2'-deoxyadenosine"
      - compartment: "e"
      - formula: "C10H13N5O3"
      - charge: 0
      - smiles: "C1C(C(OC1N2C=NC3=C(N=CN=C32)N)CO)O"
      - annotation: !!omap
          - bigg.metabolite: "dad_2"
          - chebi: "CHEBI:17256"
          - kegg.compound: "C00559"
          - metanetx.chemical: "MNXM625"
          - sbo: "SBO:0000247"
      - deltaG: 111.81
    - !!omap
      - id: "s_0134"
      - name: "2'-deoxyguanosine"
      - compartment: "c"
      - formula: "C10H13N5O4"
      - charge: 0
      - smiles: "C1C(C(OC1N2C=NC3=C2N=C(NC3=O)N)CO)O"
      - annotation: !!omap
          - bigg.metabolite: "dgsn"
          - chebi: "CHEBI:17172"
          - kegg.compound: "C00330"
          - metanetx.chemical: "MNXM647"
          - sbo: "SBO:0000247"
      - deltaG: 70.62
    - !!omap
      - id: "s_0135"
      - name: "2'-deoxyguanosine"
      - compartment: "e"
      - formula: "C10H13N5O4"
      - charge: 0
      - smiles: "C1C(C(OC1N2C=NC3=C2N=C(NC3=O)N)CO)O"
      - annotation: !!omap
          - bigg.metabolite: "dgsn"
          - chebi: "CHEBI:17172"
          - kegg.compound: "C00330"
          - metanetx.chemical: "MNXM647"
          - sbo: "SBO:0000247"
      - deltaG: 70.62
    - !!omap
      - id: "s_0136"
      - name: "2'-deoxyinosine"
      - compartment: "c"
      - formula: "C10H12N4O4"
      - charge: 0
      - smiles: "C1C(C(OC1N2C=NC3=C2N=CNC3=O)CO)O"
      - annotation: !!omap
          - bigg.metabolite: "din"
          - chebi: "CHEBI:28997"
          - kegg.compound: "C05512"
          - metanetx.chemical: "MNXM935"
          - sbo: "SBO:0000247"
      - deltaG: 33.37
    - !!omap
      - id: "s_0137"
      - name: "2'-deoxyinosine"
      - compartment: "e"
      - formula: "C10H12N4O4"
      - charge: 0
      - smiles: "C1C(C(OC1N2C=NC3=C2N=CNC3=O)CO)O"
      - annotation: !!omap
          - bigg.metabolite: "din"
          - chebi: "CHEBI:28997"
          - kegg.compound: "C05512"
          - metanetx.chemical: "MNXM935"
          - sbo: "SBO:0000247"
      - deltaG: 33.37
    - !!omap
      - id: "s_0138"
      - name: "2'-deoxyuridine"
      - compartment: "c"
      - formula: "C9H12N2O5"
      - charge: 0
      - smiles: "C1C(C(OC1N2C=CC(=O)NC2=O)CO)O"
      - annotation: !!omap
          - bigg.metabolite: "duri"
          - chebi: "CHEBI:16450"
          - kegg.compound: "C00526"
          - metanetx.chemical: "MNXM492"
          - sbo: "SBO:0000247"
      - deltaG: -28.59
    - !!omap
      - id: "s_0139"
      - name: "2'-deoxyuridine"
      - compartment: "e"
      - formula: "C9H12N2O5"
      - charge: 0
      - smiles: "C1C(C(OC1N2C=CC(=O)NC2=O)CO)O"
      - annotation: !!omap
          - bigg.metabolite: "duri"
          - chebi: "CHEBI:16450"
          - kegg.compound: "C00526"
          - metanetx.chemical: "MNXM492"
          - sbo: "SBO:0000247"
      - deltaG: -28.59
    - !!omap
      - id: "s_0140"
      - name: "2,3-bisphospho-D-glyceric acid"
      - compartment: "c"
      - formula: "C3H3O10P2"
      - charge: -5
      - smiles: "C(C(C(=O)O)OP(=O)(O)O)OP(=O)(O)O"
      - annotation: !!omap
          - bigg.metabolite: "23dpg"
          - chebi: "CHEBI:58248"
          - kegg.compound: "C01159"
          - metanetx.chemical: "MNXM892"
          - sbo: "SBO:0000247"
      - deltaG: -560.03
    - !!omap
      - id: "s_0141"
      - name: "2,5-diamino-4-hydroxy-6-(5-phosphoribosylamino)pyrimidine"
      - compartment: "c"
      - formula: "C9H14N5O8P"
      - charge: -2
      - smiles: "C(C1C(C(C(O1)NC2=C(C(=O)NC(=N2)N)N)O)O)OP(=O)(O)O"
      - annotation: !!omap
          - bigg.metabolite: "25drapp"
          - chebi: "CHEBI:58614"
          - kegg.compound: "C01304"
          - metanetx.chemical: "MNXM648"
          - sbo: "SBO:0000247"
      - deltaG: -190.84
    - !!omap
      - id: "s_0142"
      - name: "2,5-diamino-6-(5-phosphono)ribitylamino-4(3H)-pyrimidinone"
      - compartment: "c"
      - formula: "C9H16N5O8P"
      - charge: -2
      - smiles: "C(C(C(C(COP(=O)(O)O)O)O)O)NC1=C(C(=O)NC(=N1)N)N"
      - annotation: !!omap
          - bigg.metabolite: "25dthpp"
          - chebi: "CHEBI:52957"
          - kegg.compound: "C18910"
          - metanetx.chemical: "MNXM1099"
          - sbo: "SBO:0000247"
      - deltaG: -191.36
    - !!omap
      - id: "s_0143"
      - name: "2-(3-amino-3-carboxypropyl)-L-histidine"
      - compartment: "c"
      - formula: "C10H16N4O4"
      - charge: 0
      - smiles: "C1=C(NC(=N1)CCC(C(=O)O)N)CC(C(=O)O)N"
      - annotation: !!omap
          - bigg.metabolite: "caphis"
          - chebi: "CHEBI:17144"
          - kegg.compound: "C04441"
          - metanetx.chemical: "MNXM3804"
          - sbo: "SBO:0000247"
      - deltaG: 10000000
    - !!omap
      - id: "s_0144"
      - name: "2-[3-carboxy-3-(methylammonio)propyl]-L-histidine"
      - compartment: "c"
      - formula: "C11H18N4O4"
      - charge: 0
      - smiles: "C[NH2+]C(CCC1=NC=C(N1)CC(C(=O)O)N)C(=O)O"
      - annotation: !!omap
          - bigg.metabolite: "cmaphis"
          - chebi: "CHEBI:57784"
          - kegg.compound: "C04692"
          - metanetx.chemical: "MNXM3805"
          - sbo: "SBO:0000247"
      - deltaG: 10000000
    - !!omap
      - id: "s_0145"
      - name: "2-acetamido-5-oxopentanoate"
      - compartment: "m"
      - formula: "C7H10NO4"
      - charge: -1
      - smiles: "CC(=O)NC(CCC=O)C(=O)O"
      - annotation: !!omap
          - bigg.metabolite: "acg5sa"
          - chebi: "CHEBI:29123"
          - kegg.compound: "C01250"
          - metanetx.chemical: "MNXM1062"
          - sbo: "SBO:0000247"
      - deltaG: -37.04
    - !!omap
      - id: "s_0146"
      - name: "2-acetyllactic acid"
      - compartment: "m"
      - formula: "C5H7O4"
      - charge: -1
      - smiles: "CC(=O)C(C)(C(=O)O)O"
      - annotation: !!omap
          - chebi: "CHEBI:57774"
          - kegg.compound: "C00900"
          - metanetx.chemical: "MNXM426"
          - sbo: "SBO:0000247"
      - deltaG: -80.44
    - !!omap
      - id: "s_0147"
      - name: "2-amino-3-carboxymuconate-6-semialdehyde"
      - compartment: "c"
      - formula: "C7H6NO5"
      - charge: -1
      - smiles: "C(=CC(=C(C(=O)O)N)C(=O)O)C=O"
      - annotation: !!omap
          - bigg.metabolite: "cmusa"
          - chebi: "CHEBI:29044"
          - kegg.compound: "C04409"
          - metanetx.chemical: "MNXM1372"
          - sbo: "SBO:0000247"
      - deltaG: -100.45
    - !!omap
      - id: "s_0148"
      - name: "2-amino-6-(hydroxymethyl)-7,8-dihydropteridin-4-ol"
      - compartment: "m"
      - formula: "C7H9N5O2"
      - charge: 0
      - smiles: "C1C(=NC2=C(N1)N=C(NC2=O)N)CO"
      - annotation: !!omap
          - bigg.metabolite: "6hmhpt"
          - chebi: "CHEBI:17083"
          - kegg.compound: "C01300"
          - metanetx.chemical: "MNXM937"
          - sbo: "SBO:0000247"
      - deltaG: 102.85
    - !!omap
      - id: "s_0149"
      - name: "2-dehydropantoate"
      - compartment: "c"
      - formula: "C6H9O4"
      - charge: -1
      - smiles: "CC(C)(CO)C(=O)C(=O)O"
      - annotation: !!omap
          - bigg.metabolite: "2dhp"
          - chebi: "CHEBI:11561"
          - kegg.compound: "C00966"
          - metanetx.chemical: "MNXM959"
          - sbo: "SBO:0000247"
      - deltaG: -61.66
    - !!omap
      - id: "s_0150"
      - name: "2-dehydropantoate"
      - compartment: "m"
      - formula: "C6H9O4"
      - charge: -1
      - smiles: "CC(C)(CO)C(=O)C(=O)O"
      - annotation: !!omap
          - bigg.metabolite: "2dhp"
          - chebi: "CHEBI:11561"
          - kegg.compound: "C00966"
          - metanetx.chemical: "MNXM959"
          - sbo: "SBO:0000247"
      - deltaG: -61.66
    - !!omap
      - id: "s_0152"
      - name: "2-deoxy-D-ribose"
      - compartment: "c"
      - formula: "C5H10O4"
      - charge: 0
      - smiles: "C(C=O)C(C(CO)O)O"
      - annotation: !!omap
          - bigg.metabolite: "drib"
          - chebi: "CHEBI:28816"
          - kegg.compound: "C01801"
          - metanetx.chemical: "MNXM2474"
          - sbo: "SBO:0000247"
      - deltaG: -44.41
    - !!omap
      - id: "s_0153"
      - name: "2-deoxy-D-ribose 5-phosphate"
      - compartment: "c"
      - formula: "C5H9O7P"
      - charge: -2
      - annotation: !!omap
          - bigg.metabolite: "2dr5p"
          - chebi: "CHEBI:57651"
          - kegg.compound: "C00673"
          - metanetx.chemical: "MNXM2179"
          - sbo: "SBO:0000247"
      - deltaG: -267.45
    - !!omap
      - id: "s_0154"
      - name: "2-hexaprenyl-5-hydroxy-6-methoxy-3-methyl-1,4-benzoquinone"
      - compartment: "m"
      - formula: "C38H56O4"
      - charge: 0
      - smiles: "CC1=C(C(=O)C(=C(C1=O)O)OC)CC=C(C)CCC=C(C)CCC=C(C)CCC=C(C)CCC=C(C)CCC=C(C)C"
      - annotation: !!omap
          - bigg.metabolite: "2hpmhmbq"
          - chebi: "CHEBI:28753"
          - kegg.compound: "C05805"
          - metanetx.chemical: "MNXM5466"
          - sbo: "SBO:0000247"
      - deltaG: 653.19
    - !!omap
      - id: "s_0155"
      - name: "2-hexaprenyl-6-methoxy-1,4-benzoquinone"
      - compartment: "m"
      - formula: "C37H54O3"
      - charge: 0
      - smiles: "CC(=CCCC(=CCCC(=CCCC(=CCCC(=CCCC(=CCC1=CC(=O)C=C(C1=O)OC)C)C)C)C)C)C"
      - annotation: !!omap
          - bigg.metabolite: "2hp6mbq"
          - chebi: "CHEBI:27752"
          - kegg.compound: "C05803"
          - metanetx.chemical: "MNXM6707"
          - sbo: "SBO:0000247"
      - deltaG: 672.94
    - !!omap
      - id: "s_0156"
      - name: "2-hexaprenyl-6-methoxy-3-methyl-1,4-benzoquinone"
      - compartment: "m"
      - formula: "C38H56O3"
      - charge: 0
      - smiles: "CC1=C(C(=O)C(=CC1=O)OC)CC=C(C)CCC=C(C)CCC=C(C)CCC=C(C)CCC=C(C)CCC=C(C)C"
      - annotation: !!omap
          - bigg.metabolite: "2hpmmbq"
          - chebi: "CHEBI:28711"
          - kegg.compound: "C05804"
          - metanetx.chemical: "MNXM6067"
          - sbo: "SBO:0000247"
      - deltaG: 693.37
    - !!omap
      - id: "s_0157"
      - name: "2-hexaprenyl-6-methoxyphenol"
      - compartment: "m"
      - formula: "C37H56O2"
      - charge: 0
      - smiles: "CC(=CCCC(=CCCC(=CCCC(=CCCC(=CCCC(=CCC1=C(C(=CC=C1)OC)O)C)C)C)C)C)C"
      - annotation: !!omap
          - bigg.metabolite: "2hp6mp"
          - chebi: "CHEBI:1109"
          - kegg.compound: "C05802"
          - metanetx.chemical: "MNXM4291"
          - sbo: "SBO:0000247"
      - deltaG: 714
    - !!omap
      - id: "s_0158"
      - name: "2-hydroxy-3-oxobutyl phosphate"
      - compartment: "c"
      - formula: "C4H7O6P"
      - charge: -2
      - smiles: "CC(=O)C(COP(=O)(O)O)O"
      - annotation: !!omap
          - bigg.metabolite: "db4p"
          - chebi: "CHEBI:50606"
          - kegg.compound: "C15556"
          - metanetx.chemical: "MNXM2323"
          - sbo: "SBO:0000247"
      - deltaG: -253.51
    - !!omap
      - id: "s_0161"
      - name: "2-hydroxyhexadecanal"
      - compartment: "er"
      - formula: "C16H32O2"
      - charge: 0
      - smiles: "CCCCCCCCCCCCCCC(C=O)O"
      - annotation: !!omap
          - bigg.metabolite: "2hhxdal"
          - chebi: "CHEBI:50626"
          - metanetx.chemical: "MNXM35167"
          - sbo: "SBO:0000247"
      - deltaG: 264.7
    - !!omap
      - id: "s_0162"
      - name: "2-isopropylmalate"
      - compartment: "c"
      - formula: "C7H10O5"
      - charge: -2
      - smiles: "CC(C)C(CC(=O)O)(C(=O)O)O"
      - annotation: !!omap
          - bigg.metabolite: "3c3hmp"
          - chebi: "CHEBI:28107"
          - kegg.compound: "C02504"
          - metanetx.chemical: "MNXM164143"
          - sbo: "SBO:0000247"
      - deltaG: -104.59
    - !!omap
      - id: "s_0163"
      - name: "2-isopropylmalate"
      - compartment: "e"
      - formula: "C7H10O5"
      - charge: -2
      - smiles: "CC(C)C(CC(=O)O)(C(=O)O)O"
      - annotation: !!omap
          - bigg.metabolite: "3c3hmp"
          - chebi: "CHEBI:28107"
          - kegg.compound: "C02504"
          - metanetx.chemical: "MNXM164143"
          - sbo: "SBO:0000247"
      - deltaG: -104.59
    - !!omap
      - id: "s_0164"
      - name: "2-isopropylmalate"
      - compartment: "m"
      - formula: "C7H10O5"
      - charge: -2
      - smiles: "CC(C)C(CC(=O)O)(C(=O)O)O"
      - annotation: !!omap
          - bigg.metabolite: "3c3hmp"
          - chebi: "CHEBI:28107"
          - kegg.compound: "C02504"
          - metanetx.chemical: "MNXM164143"
          - sbo: "SBO:0000247"
      - deltaG: -104.59
    - !!omap
      - id: "s_0165"
      - name: "2-isopropylmaleic acid"
      - compartment: "c"
      - formula: "C7H8O4"
      - charge: -2
      - smiles: "CC(C)C(=CC(=O)O)C(=O)O"
      - annotation: !!omap
          - bigg.metabolite: "2ippm"
          - chebi: "CHEBI:58085"
          - kegg.compound: "C02631"
          - metanetx.chemical: "MNXM1706"
          - sbo: "SBO:0000247"
      - deltaG: -65.9
    - !!omap
      - id: "s_0166"
      - name: "2-methylbutanal"
      - compartment: "c"
      - formula: "C5H10O"
      - charge: 0
      - smiles: "CCC(C)C=O"
      - annotation: !!omap
          - bigg.metabolite: "2mbald"
          - chebi: "CHEBI:16182"
          - kegg.compound: "C02223"
          - metanetx.chemical: "MNXM2604"
          - sbo: "SBO:0000247"
      - deltaG: 66.37
    - !!omap
      - id: "s_0167"
      - name: "2-methylbutanal"
      - compartment: "e"
      - formula: "C5H10O"
      - charge: 0
      - smiles: "CCC(C)C=O"
      - annotation: !!omap
          - bigg.metabolite: "2mbald"
          - chebi: "CHEBI:16182"
          - kegg.compound: "C02223"
          - metanetx.chemical: "MNXM2604"
          - sbo: "SBO:0000247"
      - deltaG: 66.37
    - !!omap
      - id: "s_0168"
      - name: "2-methylbutanal"
      - compartment: "m"
      - formula: "C5H10O"
      - charge: 0
      - smiles: "CCC(C)C=O"
      - annotation: !!omap
          - bigg.metabolite: "2mbald"
          - chebi: "CHEBI:16182"
          - kegg.compound: "C02223"
          - metanetx.chemical: "MNXM2604"
          - sbo: "SBO:0000247"
      - deltaG: 66.37
    - !!omap
      - id: "s_0169"
      - name: "2-methylbutanol"
      - compartment: "c"
      - formula: "C5H12O"
      - charge: 0
      - smiles: "CCC(C)CO"
      - annotation: !!omap
          - bigg.metabolite: "2mbtoh"
          - chebi: "CHEBI:48945"
          - metanetx.chemical: "MNXM5476"
          - sbo: "SBO:0000247"
      - deltaG: 76.36
    - !!omap
      - id: "s_0170"
      - name: "2-methylbutanol"
      - compartment: "e"
      - formula: "C5H12O"
      - charge: 0
      - smiles: "CCC(C)CO"
      - annotation: !!omap
          - bigg.metabolite: "2mbtoh"
          - chebi: "CHEBI:48945"
          - metanetx.chemical: "MNXM5476"
          - sbo: "SBO:0000247"
      - deltaG: 76.36
    - !!omap
      - id: "s_0171"
      - name: "2-methylbutanol"
      - compartment: "m"
      - formula: "C5H12O"
      - charge: 0
      - smiles: "CCC(C)CO"
      - annotation: !!omap
          - bigg.metabolite: "2mbtoh"
          - chebi: "CHEBI:48945"
          - metanetx.chemical: "MNXM5476"
          - sbo: "SBO:0000247"
      - deltaG: 76.36
    - !!omap
      - id: "s_0172"
      - name: "2-methylbutyl acetate"
      - compartment: "c"
      - formula: "C7H14O2"
      - charge: 0
      - smiles: "CCC(C)COC(=O)C"
      - annotation: !!omap
          - bigg.metabolite: "2mbac"
          - chebi: "CHEBI:50585"
          - metanetx.chemical: "MNXM35277"
          - sbo: "SBO:0000247"
      - deltaG: 58.39
    - !!omap
      - id: "s_0173"
      - name: "2-methylbutyl acetate"
      - compartment: "e"
      - formula: "C7H14O2"
      - charge: 0
      - smiles: "CCC(C)COC(=O)C"
      - annotation: !!omap
          - bigg.metabolite: "2mbac"
          - chebi: "CHEBI:50585"
          - metanetx.chemical: "MNXM35277"
          - sbo: "SBO:0000247"
      - deltaG: 58.39
    - !!omap
      - id: "s_0174"
      - name: "2-methylcitrate"
      - compartment: "m"
      - formula: "C7H7O7"
      - charge: -3
      - smiles: "CC(C(=O)O)C(CC(=O)O)(C(=O)O)O"
      - annotation: !!omap
          - bigg.metabolite: "2mcit"
          - chebi: "CHEBI:15598"
          - kegg.compound: "C02225"
          - metanetx.chemical: "MNXM90279"
          - sbo: "SBO:0000247"
      - deltaG: -209.44
    - !!omap
      - id: "s_0176"
      - name: "2-oxoadipic acid"
      - compartment: "c"
      - formula: "C6H6O5"
      - charge: -2
      - smiles: "C(CC(=O)C(=O)O)CC(=O)O"
      - annotation: !!omap
          - bigg.metabolite: "2oxoadp"
          - chebi: "CHEBI:57499"
          - kegg.compound: "C00322"
          - metanetx.chemical: "MNXM263"
          - sbo: "SBO:0000247"
      - deltaG: -129.63
    - !!omap
      - id: "s_0177"
      - name: "2-oxoadipic acid"
      - compartment: "m"
      - formula: "C6H6O5"
      - charge: -2
      - smiles: "C(CC(=O)C(=O)O)CC(=O)O"
      - annotation: !!omap
          - bigg.metabolite: "2oxoadp"
          - chebi: "CHEBI:57499"
          - kegg.compound: "C00322"
          - metanetx.chemical: "MNXM263"
          - sbo: "SBO:0000247"
      - deltaG: -129.63
    - !!omap
      - id: "s_0178"
      - name: "2-oxobutanoate"
      - compartment: "c"
      - formula: "C4H5O3"
      - charge: -1
      - smiles: "CCC(=O)C(=O)[O-]"
      - annotation: !!omap
          - bigg.metabolite: "2obut"
          - chebi: "CHEBI:16763"
          - kegg.compound: "C00109"
          - metanetx.chemical: "MNXM159"
          - sbo: "SBO:0000247"
      - deltaG: -62.58
    - !!omap
      - id: "s_0179"
      - name: "2-oxobutanoate"
      - compartment: "m"
      - formula: "C4H5O3"
      - charge: -1
      - smiles: "CCC(=O)C(=O)[O-]"
      - annotation: !!omap
          - bigg.metabolite: "2obut"
          - chebi: "CHEBI:16763"
          - kegg.compound: "C00109"
          - metanetx.chemical: "MNXM159"
          - sbo: "SBO:0000247"
      - deltaG: -62.58
    - !!omap
      - id: "s_0180"
      - name: "2-oxoglutarate"
      - compartment: "c"
      - formula: "C5H4O5"
      - charge: -2
      - smiles: "C(CC(=O)O)C(=O)C(=O)O"
      - annotation: !!omap
          - bigg.metabolite: "akg"
          - chebi: "CHEBI:16810"
          - kegg.compound: "C00026"
          - metanetx.chemical: "MNXM20"
          - sbo: "SBO:0000247"
      - deltaG: -150.73
    - !!omap
      - id: "s_0181"
      - name: "2-oxoglutarate"
      - compartment: "e"
      - formula: "C5H4O5"
      - charge: -2
      - smiles: "C(CC(=O)O)C(=O)C(=O)O"
      - annotation: !!omap
          - bigg.metabolite: "akg"
          - chebi: "CHEBI:16810"
          - kegg.compound: "C00026"
          - metanetx.chemical: "MNXM20"
          - sbo: "SBO:0000247"
      - deltaG: -150.73
    - !!omap
      - id: "s_0182"
      - name: "2-oxoglutarate"
      - compartment: "m"
      - formula: "C5H4O5"
      - charge: -2
      - smiles: "C(CC(=O)O)C(=O)C(=O)O"
      - annotation: !!omap
          - bigg.metabolite: "akg"
          - chebi: "CHEBI:16810"
          - kegg.compound: "C00026"
          - metanetx.chemical: "MNXM20"
          - sbo: "SBO:0000247"
      - deltaG: -150.73
    - !!omap
      - id: "s_0183"
      - name: "2-oxoglutarate"
      - compartment: "n"
      - formula: "C5H4O5"
      - charge: -2
      - smiles: "C(CC(=O)O)C(=O)C(=O)O"
      - annotation: !!omap
          - bigg.metabolite: "akg"
          - chebi: "CHEBI:16810"
          - kegg.compound: "C00026"
          - metanetx.chemical: "MNXM20"
          - sbo: "SBO:0000247"
      - deltaG: -150.73
    - !!omap
      - id: "s_0184"
      - name: "2-oxoglutarate"
      - compartment: "p"
      - formula: "C5H4O5"
      - charge: -2
      - smiles: "C(CC(=O)O)C(=O)C(=O)O"
      - annotation: !!omap
          - bigg.metabolite: "akg"
          - chebi: "CHEBI:16810"
          - kegg.compound: "C00026"
          - metanetx.chemical: "MNXM20"
          - sbo: "SBO:0000247"
      - deltaG: -150.73
    - !!omap
      - id: "s_0185"
      - name: "2-phenylethanol"
      - compartment: "c"
      - formula: "C8H10O"
      - charge: 0
      - smiles: "C1=CC=C(C=C1)CCO"
      - annotation: !!omap
          - bigg.metabolite: "2phetoh"
          - chebi: "CHEBI:49000"
          - kegg.compound: "C05853"
          - metanetx.chemical: "MNXM2476"
          - sbo: "SBO:0000247"
      - deltaG: 90.71
    - !!omap
      - id: "s_0186"
      - name: "2-phenylethanol"
      - compartment: "e"
      - formula: "C8H10O"
      - charge: 0
      - smiles: "C1=CC=C(C=C1)CCO"
      - annotation: !!omap
          - bigg.metabolite: "2phetoh"
          - chebi: "CHEBI:49000"
          - kegg.compound: "C05853"
          - metanetx.chemical: "MNXM2476"
          - sbo: "SBO:0000247"
      - deltaG: 90.71
    - !!omap
      - id: "s_0187"
      - name: "2-phenylethanol"
      - compartment: "m"
      - formula: "C8H10O"
      - charge: 0
      - smiles: "C1=CC=C(C=C1)CCO"
      - annotation: !!omap
          - bigg.metabolite: "2phetoh"
          - chebi: "CHEBI:49000"
          - kegg.compound: "C05853"
          - metanetx.chemical: "MNXM2476"
          - sbo: "SBO:0000247"
      - deltaG: 90.71
    - !!omap
      - id: "s_0188"
      - name: "2-phospho-D-glyceric acid"
      - compartment: "c"
      - formula: "C3H4O7P"
      - charge: -3
      - smiles: "C(C(C(=O)O)OP(=O)(O)O)O"
      - annotation: !!omap
          - bigg.metabolite: "2pg"
          - chebi: "CHEBI:58289"
          - kegg.compound: "C00631"
          - metanetx.chemical: "MNXM275"
          - sbo: "SBO:0000247"
      - deltaG: -334.7
    - !!omap
      - id: "s_0189"
      - name: "geranylgeranyl diphosphate"
      - compartment: "c"
      - formula: "C20H33O7P2"
      - charge: -3
      - smiles: "CC(=CCCC(=CCCC(=CCCC(=CCOP(=O)(O)OP(=O)(O)O)C)C)C)C"
      - annotation: !!omap
          - bigg.metabolite: "ggdp"
          - chebi: "CHEBI:58756"
          - kegg.compound: "C00353"
          - metanetx.chemical: "MNXM139"
          - sbo: "SBO:0000247"
      - deltaG: -25.41
    - !!omap
      - id: "s_0190"
      - name: "farnesyl diphosphate"
      - compartment: "c"
      - formula: "C15H25O7P2"
      - charge: -3
      - smiles: "CC(=CCCC(=CCCC(=CCOP(=O)(O)OP(=O)(O)O)C)C)C"
      - annotation: !!omap
          - bigg.metabolite: "frdp"
          - chebi: "CHEBI:175763"
          - kegg.compound: "C00448"
          - metanetx.chemical: "MNXM34"
          - sbo: "SBO:0000247"
      - deltaG: -131.39
    - !!omap
      - id: "s_0191"
      - name: "farnesyl diphosphate"
      - compartment: "m"
      - formula: "C15H25O7P2"
      - charge: -3
      - smiles: "CC(=CCCC(=CCCC(=CCOP(=O)(O)OP(=O)(O)O)C)C)C"
      - annotation: !!omap
          - bigg.metabolite: "frdp"
          - chebi: "CHEBI:175763"
          - kegg.compound: "C00448"
          - metanetx.chemical: "MNXM34"
          - sbo: "SBO:0000247"
      - deltaG: -131.39
    - !!omap
      - id: "s_0192"
      - name: "3',5'-cyclic AMP"
      - compartment: "c"
      - formula: "C10H11N5O6P"
      - charge: -1
      - smiles: "C1C2C(C(C(O2)N3C=NC4=C(N=CN=C43)N)O)OP(=O)(O1)O"
      - annotation: !!omap
          - bigg.metabolite: "camp"
          - chebi: "CHEBI:58165"
          - kegg.compound: "C00575"
          - metanetx.chemical: "MNXM243"
          - sbo: "SBO:0000247"
      - deltaG: -93.83
    - !!omap
      - id: "s_0193"
      - name: "3',5'-cyclic CMP"
      - compartment: "c"
      - formula: "C9H11N3O7P"
      - charge: -1
      - smiles: "C1C2C(C(C(O2)N3C=CC(=NC3=O)N)O)OP(=O)(O1)O"
      - annotation: !!omap
          - bigg.metabolite: "35ccmp"
          - chebi: "CHEBI:58003"
          - kegg.compound: "C00941"
          - metanetx.chemical: "MNXM3162"
          - sbo: "SBO:0000247"
      - deltaG: -172.49
    - !!omap
      - id: "s_0194"
      - name: "3',5'-cyclic dAMP"
      - compartment: "c"
      - formula: "C10H11N5O5P"
      - charge: -1
      - smiles: "C1C2C(COP(=O)(O2)O)OC1N3C=NC4=C(N=CN=C43)N"
      - annotation: !!omap
          - bigg.metabolite: "35cdamp"
          - chebi: "CHEBI:28074"
          - kegg.compound: "C00968"
          - metanetx.chemical: "MNXM9931"
          - sbo: "SBO:0000247"
      - deltaG: -53.99
    - !!omap
      - id: "s_0195"
      - name: "3',5'-cyclic GMP"
      - compartment: "c"
      - formula: "C10H11N5O7P"
      - charge: -1
      - smiles: "C1C2C(C(C(O2)N3C=NC4=C3N=C(NC4=O)N)O)OP(=O)(O1)O"
      - annotation: !!omap
          - bigg.metabolite: "35cgmp"
          - chebi: "CHEBI:57746"
          - kegg.compound: "C00942"
          - metanetx.chemical: "MNXM665"
          - sbo: "SBO:0000247"
      - deltaG: -135.41
    - !!omap
      - id: "s_0196"
      - name: "3',5'-cyclic IMP"
      - compartment: "c"
      - formula: "C10H10N4O7P"
      - charge: -1
      - smiles: "C1C2C(C(C(O2)N3C=NC4=C3N=CNC4=O)O)OP(=O)(O1)O"
      - annotation: !!omap
          - bigg.metabolite: "35cimp"
          - chebi: "CHEBI:27541"
          - kegg.compound: "C00943"
          - metanetx.chemical: "MNXM9932"
          - sbo: "SBO:0000247"
      - deltaG: -170.13
    - !!omap
      - id: "s_0197"
      - name: "3'-dephospho-CoA"
      - compartment: "c"
      - formula: "C21H33N7O13P2S"
      - charge: -2
      - smiles: "CC(C)(COP(=O)(O)OP(=O)(O)OCC1C(C(C(O1)N2C=NC3=C(N=CN=C32)N)O)O)C(C(=O)NCCC(=O)NCCS)O"
      - annotation: !!omap
          - bigg.metabolite: "dpcoa"
          - chebi: "CHEBI:57328"
          - kegg.compound: "C00882"
          - metanetx.chemical: "MNXM481"
          - sbo: "SBO:0000247"
      - deltaG: -227.43
    - !!omap
      - id: "s_0200"
      - name: "3'-dephospho-CoA"
      - compartment: "m"
      - formula: "C21H33N7O13P2S"
      - charge: -2
      - smiles: "CC(C)(COP(=O)(O)OP(=O)(O)OCC1C(C(C(O1)N2C=NC3=C(N=CN=C32)N)O)O)C(C(=O)NCCC(=O)NCCS)O"
      - annotation: !!omap
          - bigg.metabolite: "dpcoa"
          - chebi: "CHEBI:57328"
          - kegg.compound: "C00882"
          - metanetx.chemical: "MNXM481"
          - sbo: "SBO:0000247"
      - deltaG: -227.43
    - !!omap
      - id: "s_0201"
      - name: "3'-phospho-5'-adenylyl sulfate"
      - compartment: "c"
      - formula: "C10H11N5O13P2S"
      - charge: -4
      - smiles: "C1=NC(=C2C(=N1)N(C=N2)C3C(C(C(O3)COP(=O)(O)OS(=O)(=O)O)OP(=O)(O)O)O)N"
      - annotation: !!omap
          - bigg.metabolite: "paps"
          - chebi: "CHEBI:58339"
          - kegg.compound: "C00053"
          - metanetx.chemical: "MNXM49"
          - sbo: "SBO:0000247"
      - deltaG: -487.05
    - !!omap
      - id: "s_0204"
      - name: "3-(4-hydroxyphenyl)pyruvate"
      - compartment: "c"
      - formula: "C9H7O4"
      - charge: -1
      - smiles: "C1=CC(=CC=C1CC(=O)C(=O)[O-])O"
      - annotation: !!omap
          - bigg.metabolite: "34hpp"
          - chebi: "CHEBI:36242"
          - kegg.compound: "C01179"
          - metanetx.chemical: "MNXM153"
          - sbo: "SBO:0000247"
      - deltaG: -47.34
    - !!omap
      - id: "s_0206"
      - name: "3-(4-hydroxyphenyl)pyruvate"
      - compartment: "p"
      - formula: "C9H7O4"
      - charge: -1
      - smiles: "C1=CC(=CC=C1CC(=O)C(=O)[O-])O"
      - annotation: !!omap
          - bigg.metabolite: "34hpp"
          - chebi: "CHEBI:36242"
          - kegg.compound: "C01179"
          - metanetx.chemical: "MNXM153"
          - sbo: "SBO:0000247"
      - deltaG: -47.34
    - !!omap
      - id: "s_0207"
      - name: "3-(imidazol-4-yl)-2-oxopropyl dihydrogen phosphate"
      - compartment: "c"
      - formula: "C6H7N2O5P"
      - charge: -2
      - smiles: "C1=C(NC=N1)CC(=O)COP(=O)(O)O"
      - annotation: !!omap
          - bigg.metabolite: "imacp"
          - chebi: "CHEBI:57766"
          - kegg.compound: "C01267"
          - metanetx.chemical: "MNXM1456"
          - sbo: "SBO:0000247"
      - deltaG: -174.77
    - !!omap
      - id: "s_0208"
      - name: "3-aminopropanal"
      - compartment: "c"
      - formula: "C3H8NO"
      - charge: 1
      - smiles: "C(CN)C=O"
      - annotation: !!omap
          - bigg.metabolite: "aproa"
          - chebi: "CHEBI:58374"
          - kegg.compound: "C02229"
          - metanetx.chemical: "MNXM493"
          - sbo: "SBO:0000247"
      - deltaG: 44.29
    - !!omap
      - id: "s_0209"
      - name: "3-dehydro-4-methylzymosterol"
      - compartment: "c"
      - formula: "C28H44O"
      - charge: 0
      - smiles: "CC1C2CCC3=C(C2(CCC1=O)C)CCC4(C3CCC4C(C)CCC=C(C)C)C"
      - annotation: !!omap
          - chebi: "CHEBI:50593"
          - kegg.compound: "C15816"
          - metanetx.chemical: "MNXM36392"
          - sbo: "SBO:0000247"
      - deltaG: 479.18
    - !!omap
      - id: "s_0210"
      - name: "3-dehydroquinate"
      - compartment: "c"
      - formula: "C7H9O6"
      - charge: -1
      - smiles: "C1C(C(C(=O)CC1(C(=O)[O-])O)O)O"
      - annotation: !!omap
          - bigg.metabolite: "3dhq"
          - chebi: "CHEBI:32364"
          - kegg.compound: "C00944"
          - metanetx.chemical: "MNXM478"
          - sbo: "SBO:0000247"
      - deltaG: -127.05
    - !!omap
      - id: "s_0211"
      - name: "3-dehydroshikimate"
      - compartment: "c"
      - formula: "C7H7O5"
      - charge: -1
      - smiles: "C1C(C(C(=O)C=C1C(=O)O)O)O"
      - annotation: !!omap
          - bigg.metabolite: "3dhsk"
          - chebi: "CHEBI:16630"
          - kegg.compound: "C02637"
          - metanetx.chemical: "MNXM611"
          - sbo: "SBO:0000247"
      - deltaG: -98.32
    - !!omap
      - id: "s_0212"
      - name: "3-hexaprenyl-4,5-dihydroxybenzoic acid"
      - compartment: "c"
      - formula: "C37H53O4"
      - charge: -1
      - smiles: "CC(=CCCC(=CCCC(=CCCC(=CCCC(=CCCC(=CCC1=C(C(=CC(=C1)C(=O)O)O)O)C)C)C)C)C)C"
      - annotation: !!omap
          - bigg.metabolite: "3dh5hpb"
          - chebi: "CHEBI:58373"
          - kegg.compound: "C05200"
          - metanetx.chemical: "MNXM2783"
          - sbo: "SBO:0000247"
      - deltaG: 549.14
    - !!omap
      - id: "s_0213"
      - name: "3-hexaprenyl-4,5-dihydroxybenzoic acid"
      - compartment: "m"
      - formula: "C37H53O4"
      - charge: -1
      - smiles: "CC(=CCCC(=CCCC(=CCCC(=CCCC(=CCCC(=CCC1=C(C(=CC(=C1)C(=O)O)O)O)C)C)C)C)C)C"
      - annotation: !!omap
          - bigg.metabolite: "3dh5hpb"
          - chebi: "CHEBI:58373"
          - kegg.compound: "C05200"
          - metanetx.chemical: "MNXM2783"
          - sbo: "SBO:0000247"
      - deltaG: 549.14
    - !!omap
      - id: "s_0214"
      - name: "3-hexaprenyl-4-hydroxy-5-methoxybenzoic acid"
      - compartment: "m"
      - formula: "C38H55O4"
      - charge: -1
      - smiles: "CC(=CCCC(=CCCC(=CCCC(=CCCC(=CCCC(=CCC1=C(C(=CC(=C1)C(=O)O)OC)O)C)C)C)C)C)C"
      - annotation: !!omap
          - bigg.metabolite: "3hph5mb"
          - chebi: "CHEBI:57916"
          - kegg.compound: "C05313"
          - metanetx.chemical: "MNXM3166"
          - sbo: "SBO:0000247"
      - deltaG: 620.76
    - !!omap
      - id: "s_0215"
      - name: "3-hexaprenyl-4-hydroxybenzoic acid"
      - compartment: "c"
      - formula: "C37H53O3"
      - charge: -1
      - smiles: "CC(=CCCC(=CCCC(=CCCC(=CCCC(=CCCC(=CCC1=C(C=CC(=C1)C(=O)O)O)C)C)C)C)C)C"
      - annotation: !!omap
          - bigg.metabolite: "3ophb_5"
          - chebi: "CHEBI:31116"
          - kegg.compound: "C13425"
          - metanetx.chemical: "MNXM3167"
          - sbo: "SBO:0000247"
      - deltaG: 588.62
    - !!omap
      - id: "s_0216"
      - name: "3-hexaprenyl-4-hydroxybenzoic acid"
      - compartment: "m"
      - formula: "C37H53O3"
      - charge: -1
      - smiles: "CC(=CCCC(=CCCC(=CCCC(=CCCC(=CCCC(=CCC1=C(C=CC(=C1)C(=O)O)O)C)C)C)C)C)C"
      - annotation: !!omap
          - bigg.metabolite: "3ophb_5"
          - chebi: "CHEBI:31116"
          - kegg.compound: "C13425"
          - metanetx.chemical: "MNXM3167"
          - sbo: "SBO:0000247"
      - deltaG: 588.62
    - !!omap
      - id: "s_0217"
      - name: "3-hydroxy-2-isopropyl-4-methoxy-4-oxobutanoate"
      - compartment: "c"
      - formula: "C8H13O5"
      - charge: -1
      - smiles: "CC(C)C(C(C(=O)OC)O)C(=O)[O-]"
      - annotation: !!omap
          - chebi: "CHEBI:52960"
          - metanetx.chemical: "MNXM36486"
          - sbo: "SBO:0000247"
      - deltaG: -69.22
    - !!omap
      - id: "s_0218"
      - name: "3-hydroxy-3-methylglutaryl-CoA"
      - compartment: "c"
      - formula: "C27H39N7O20P3S"
      - charge: -5
      - smiles: "CC(C)(COP(=O)(O)OP(=O)(O)OCC1C(C(C(O1)N2C=NC3=C(N=CN=C32)N)O)OP(=O)(O)O)C(C(=O)NCCC(=O)NCCSC(=O)CC(C)(CC(=O)O)O)O"
      - annotation: !!omap
          - bigg.metabolite: "hmgcoa"
          - chebi: "CHEBI:11814"
          - kegg.compound: "C00356"
          - metanetx.chemical: "MNXM197"
          - sbo: "SBO:0000247"
      - deltaG: -535.86
    - !!omap
      - id: "s_0221"
      - name: "3-hydroxy-3-methylglutaryl-CoA"
      - compartment: "m"
      - formula: "C27H39N7O20P3S"
      - charge: -5
      - smiles: "CC(C)(COP(=O)(O)OP(=O)(O)OCC1C(C(C(O1)N2C=NC3=C(N=CN=C32)N)O)OP(=O)(O)O)C(C(=O)NCCC(=O)NCCSC(=O)CC(C)(CC(=O)O)O)O"
      - annotation: !!omap
          - bigg.metabolite: "hmgcoa"
          - chebi: "CHEBI:11814"
          - kegg.compound: "C00356"
          - metanetx.chemical: "MNXM197"
          - sbo: "SBO:0000247"
      - deltaG: -535.86
    - !!omap
      - id: "s_0222"
      - name: "3-hydroxy-L-kynurenine"
      - compartment: "c"
      - formula: "C10H12N2O4"
      - charge: 0
      - smiles: "C1=CC(=C(C(=C1)O)N)C(=O)CC(C(=O)O)N"
      - annotation: !!omap
          - bigg.metabolite: "hLkynr"
          - chebi: "CHEBI:17380"
          - kegg.compound: "C03227"
          - metanetx.chemical: "MNXM576"
          - sbo: "SBO:0000247"
      - deltaG: 0.42
    - !!omap
      - id: "s_0224"
      - name: "3-hydroxyanthranilate"
      - compartment: "c"
      - formula: "C7H6NO3"
      - charge: -1
      - smiles: "C1=CC(=C(C(=C1)O)N)C(=O)O"
      - annotation: !!omap
          - bigg.metabolite: "3hanthrn"
          - chebi: "CHEBI:36559"
          - kegg.compound: "C00632"
          - metanetx.chemical: "MNXM359"
          - sbo: "SBO:0000247"
      - deltaG: -30.24
    - !!omap
      - id: "s_0229"
      - name: "3-hydroxyoctadecanoyl-CoA"
      - compartment: "p"
      - formula: "C39H66N7O18P3S"
      - charge: -4
      - smiles: "CCCCCCCCCCCCCCCC(CC(=O)SCCNC(=O)CCNC(=O)C(C(C)(C)COP(=O)(O)OP(=O)(O)OCC1C(C(C(O1)N2C=NC3=C(N=CN=C32)N)O)OP(=O)(O)O)O)O"
      - annotation: !!omap
          - bigg.metabolite: "CE2248"
          - chebi: "CHEBI:50583"
          - kegg.compound: "C16217"
          - metanetx.chemical: "MNXM1309"
          - sbo: "SBO:0000247"
      - deltaG: -70.82
    - !!omap
      - id: "s_0231"
      - name: "3-ketosphinganine"
      - compartment: "er"
      - formula: "C18H38NO2"
      - charge: 1
      - smiles: "CCCCCCCCCCCCCCCC(=O)C(CO)N"
      - annotation: !!omap
          - bigg.metabolite: "3dsphgn"
          - chebi: "CHEBI:58299"
          - kegg.compound: "C02934"
          - metanetx.chemical: "MNXM559"
          - sbo: "SBO:0000247"
      - deltaG: 322.59
    - !!omap
      - id: "s_0232"
      - name: "3-methyl-2-oxobutanoate"
      - compartment: "c"
      - formula: "C5H7O3"
      - charge: -1
      - smiles: "CC(C)C(=O)C(=O)O"
      - annotation: !!omap
          - bigg.metabolite: "3mob"
          - chebi: "CHEBI:11851"
          - kegg.compound: "C00141"
          - metanetx.chemical: "MNXM238"
          - sbo: "SBO:0000247"
      - deltaG: -43.3
    - !!omap
      - id: "s_0233"
      - name: "3-methyl-2-oxobutanoate"
      - compartment: "m"
      - formula: "C5H7O3"
      - charge: -1
      - smiles: "CC(C)C(=O)C(=O)O"
      - annotation: !!omap
          - bigg.metabolite: "3mob"
          - chebi: "CHEBI:11851"
          - kegg.compound: "C00141"
          - metanetx.chemical: "MNXM238"
          - sbo: "SBO:0000247"
      - deltaG: -43.3
    - !!omap
      - id: "s_0234"
      - name: "3-methylbutanal"
      - compartment: "c"
      - formula: "C5H10O"
      - charge: 0
      - smiles: "CC(C)CC=O"
      - annotation: !!omap
          - bigg.metabolite: "3mbald"
          - chebi: "CHEBI:16638"
          - kegg.compound: "C07329"
          - metanetx.chemical: "MNXM5280"
          - sbo: "SBO:0000247"
      - deltaG: 66.37
    - !!omap
      - id: "s_0235"
      - name: "3-methylbutanal"
      - compartment: "e"
      - formula: "C5H10O"
      - charge: 0
      - smiles: "CC(C)CC=O"
      - annotation: !!omap
          - bigg.metabolite: "3mbald"
          - chebi: "CHEBI:16638"
          - kegg.compound: "C07329"
          - metanetx.chemical: "MNXM5280"
          - sbo: "SBO:0000247"
      - deltaG: 66.37
    - !!omap
      - id: "s_0236"
      - name: "3-methylbutanal"
      - compartment: "m"
      - formula: "C5H10O"
      - charge: 0
      - smiles: "CC(C)CC=O"
      - annotation: !!omap
          - bigg.metabolite: "3mbald"
          - chebi: "CHEBI:16638"
          - kegg.compound: "C07329"
          - metanetx.chemical: "MNXM5280"
          - sbo: "SBO:0000247"
      - deltaG: 66.37
    - !!omap
      - id: "s_0239"
      - name: "3-oxodecanoyl-CoA"
      - compartment: "p"
      - formula: "C31H48N7O18P3S"
      - charge: -4
      - smiles: "CCCCCCCC(=O)CC(=O)SCCNC(=O)CCNC(=O)C(C(C)(C)COP(=O)(O)OP(=O)(O)OCC1C(C(C(O1)N2C=NC3=C(N=CN=C32)N)O)OP(=O)(O)O)O"
      - annotation: !!omap
          - bigg.metabolite: "3odcoa"
          - chebi: "CHEBI:28528"
          - kegg.compound: "C05265"
          - metanetx.chemical: "MNXM677"
          - sbo: "SBO:0000247"
      - deltaG: -276.99
    - !!omap
      - id: "s_0243"
      - name: "3-oxohexacosanoyl-CoA"
      - compartment: "p"
      - formula: "C47H80N7O18P3S"
      - charge: -4
      - smiles: "CCCCCCCCCCCCCCCCCCCCCCCC(=O)CC(=O)SCCNC(=O)CCNC(=O)C(C(C)(C)COP(=O)(O)OP(=O)(O)OCC1C(C(C(O1)N2C=NC3=C(N=CN=C32)N)O)OP(=O)(O)O)O"
      - annotation: !!omap
          - bigg.metabolite: "3ohxccoa"
          - chebi: "CHEBI:52977"
          - metanetx.chemical: "MNXM36758"
          - sbo: "SBO:0000247"
      - deltaG: 106.67
    - !!omap
      - id: "s_0247"
      - name: "3-oxolauroyl-CoA"
      - compartment: "p"
      - formula: "C33H52N7O18P3S"
      - charge: -4
      - smiles: "CCCCCCCCCC(=O)CC(=O)SCCNC(=O)CCNC(=O)C(C(C)(C)COP(=O)(O)OP(=O)(O)OCC1C(C(C(O1)N2C=NC3=C(N=CN=C32)N)O)OP(=O)(O)O)O"
      - annotation: !!omap
          - bigg.metabolite: "3oddcoa"
          - chebi: "CHEBI:27868"
          - kegg.compound: "C05263"
          - metanetx.chemical: "MNXM705"
          - sbo: "SBO:0000247"
      - deltaG: -229.03
    - !!omap
      - id: "s_0250"
      - name: "3-oxooctadecanoyl-CoA"
      - compartment: "p"
      - formula: "C39H64N7O18P3S"
      - charge: -4
      - smiles: "CCCCCCCCCCCCCCCC(=O)CC(=O)SCCNC(=O)CCNC(=O)C(C(C)(C)COP(=O)(O)OP(=O)(O)OCC1C(C(C(O1)N2C=NC3=C(N=CN=C32)N)O)OP(=O)(O)O)O"
      - annotation: !!omap
          - bigg.metabolite: "3ohodcoa"
          - chebi: "CHEBI:50571"
          - kegg.compound: "C16216"
          - metanetx.chemical: "MNXM513"
          - sbo: "SBO:0000247"
      - deltaG: -85.16
    - !!omap
      - id: "s_0253"
      - name: "3-oxopalmitoyl-CoA"
      - compartment: "p"
      - formula: "C37H60N7O18P3S"
      - charge: -4
      - smiles: "CCCCCCCCCCCCCC(=O)CC(=O)SCCNC(=O)CCNC(=O)C(C(C)(C)COP(=O)(O)OP(=O)(O)OCC1C(C(C(O1)N2C=NC3=C(N=CN=C32)N)O)OP(=O)(O)O)O"
      - annotation: !!omap
          - bigg.metabolite: "3ohdcoa"
          - chebi: "CHEBI:57349"
          - kegg.compound: "C05259"
          - metanetx.chemical: "MNXM738"
          - sbo: "SBO:0000247"
      - deltaG: -133.11
    - !!omap
      - id: "s_0257"
      - name: "3-oxotetradecanoyl-CoA"
      - compartment: "p"
      - formula: "C35H56N7O18P3S"
      - charge: -4
      - smiles: "CCCCCCCCCCCC(=O)CC(=O)SCCNC(=O)CCNC(=O)C(C(C)(C)COP(=O)(O)OP(=O)(O)OCC1C(C(C(O1)N2C=NC3=C(N=CN=C32)N)O)OP(=O)(O)O)O"
      - annotation: !!omap
          - bigg.metabolite: "3otdcoa"
          - chebi: "CHEBI:28726"
          - kegg.compound: "C05261"
          - metanetx.chemical: "MNXM707"
          - sbo: "SBO:0000247"
      - deltaG: -181.07
    - !!omap
      - id: "s_0258"
      - name: "3-phospho-hydroxypyruvate"
      - compartment: "c"
      - formula: "C3H2O7P"
      - charge: -3
      - smiles: "C(C(=O)C(=O)[O-])OP(=O)([O-])[O-]"
      - annotation: !!omap
          - bigg.metabolite: "3php"
          - chebi: "CHEBI:18110"
          - kegg.compound: "C03232"
          - metanetx.chemical: "MNXM541"
          - sbo: "SBO:0000247"
      - deltaG: -343.7
    - !!omap
      - id: "s_0259"
      - name: "3-phospho-serine"
      - compartment: "c"
      - formula: "C3H6NO6P"
      - charge: -2
      - smiles: "C(C(C(=O)O)N)OP(=O)(O)O"
      - annotation: !!omap
          - bigg.metabolite: "pser__L"
          - chebi: "CHEBI:57524"
          - kegg.compound: "C01005"
          - metanetx.chemical: "MNXM379"
          - sbo: "SBO:0000247"
      - deltaG: -281.93
    - !!omap
      - id: "s_0260"
      - name: "3-phosphonato-D-glycerate(3-)"
      - compartment: "c"
      - formula: "C3H4O7P"
      - charge: -3
      - smiles: "C(C(C(=O)[O-])O)OP(=O)([O-])[O-]"
      - annotation: !!omap
          - bigg.metabolite: "3pg"
          - chebi: "CHEBI:58272"
          - kegg.compound: "C00197"
          - metanetx.chemical: "MNXM126"
          - sbo: "SBO:0000247"
      - deltaG: -334.72
    - !!omap
      - id: "s_0261"
      - name: "3-phosphoshikimic acid"
      - compartment: "c"
      - formula: "C7H8O8P"
      - charge: -3
      - smiles: "C1C(C(C(C=C1C(=O)O)OP(=O)(O)O)O)O"
      - annotation: !!omap
          - bigg.metabolite: "skm5p"
          - chebi: "CHEBI:145989"
          - kegg.compound: "C03175"
          - metanetx.chemical: "MNXM1265"
          - sbo: "SBO:0000247"
      - deltaG: -307.66
    - !!omap
      - id: "s_0262"
      - name: "4,4-dimethyl-5alpha-cholesta-8,14,24-trien-3beta-ol"
      - compartment: "c"
      - formula: "C29H46O"
      - charge: 0
      - smiles: "CC(CCC=C(C)C)C1CC=C2C1(CCC3=C2CCC4C3(CCC(C4(C)C)O)C)C"
      - annotation: !!omap
          - bigg.metabolite: "44mctr"
          - chebi: "CHEBI:17813"
          - kegg.compound: "C11455"
          - metanetx.chemical: "MNXM1036"
          - sbo: "SBO:0000247"
      - deltaG: 505.34
    - !!omap
      - id: "s_0263"
      - name: "4,5-bis(diphospho)-1D-myo-inositol tetrakisphosphate"
      - compartment: "c"
      - formula: "C6H7O30P8"
      - charge: -13
      - smiles: "C1(C(C(C(C(C1OP(=O)(O)O)OP(=O)(O)OP(=O)(O)O)OP(=O)(O)OP(=O)(O)O)OP(=O)(O)O)OP(=O)(O)O)OP(=O)(O)O"
      - annotation: !!omap
          - chebi: "CHEBI:53066"
          - metanetx.chemical: "MNXM484827"
          - sbo: "SBO:0000247"
      - deltaG: -1769.75
    - !!omap
      - id: "s_0264"
      - name: "4-(phosphonooxy)-L-threonine"
      - compartment: "c"
      - formula: "C4H8NO7P"
      - charge: -2
      - smiles: "C(C(C(C(=O)O)N)O)OP(=O)(O)O"
      - annotation: !!omap
          - bigg.metabolite: "phthr"
          - chebi: "CHEBI:58452"
          - kegg.compound: "C06055"
          - metanetx.chemical: "MNXM759"
          - sbo: "SBO:0000247"
      - deltaG: -297.62
    - !!omap
      - id: "s_0267"
      - name: "4-amino-2-methyl-5-diphosphomethylpyrimidine"
      - compartment: "c"
      - formula: "C6H8N3O7P2"
      - charge: -3
      - smiles: "CC1=NC=C(C(=N1)N)COP(=O)(O)OP(=O)(O)O"
      - annotation: !!omap
          - bigg.metabolite: "2mahmp"
          - chebi: "CHEBI:57841"
          - kegg.compound: "C04752"
          - metanetx.chemical: "MNXM1135"
          - sbo: "SBO:0000247"
      - deltaG: -325.57
    - !!omap
      - id: "s_0268"
      - name: "4-amino-2-methyl-5-phosphomethylpyrimidine"
      - compartment: "c"
      - formula: "C6H8N3O4P"
      - charge: -2
      - smiles: "CC1=NC=C(C(=N1)N)COP(=O)(O)O"
      - annotation: !!omap
          - bigg.metabolite: "4ampm"
          - chebi: "CHEBI:58354"
          - kegg.compound: "C04556"
          - metanetx.chemical: "MNXM790"
          - sbo: "SBO:0000247"
      - deltaG: -119.06
    - !!omap
      - id: "s_0269"
      - name: "4-amino-4-deoxychorismate"
      - compartment: "c"
      - formula: "C10H10NO5"
      - charge: -1
      - smiles: "C=C(C(=O)[O-])OC1C=C(C=CC1[NH3+])C(=O)[O-]"
      - annotation: !!omap
          - bigg.metabolite: "4adcho"
          - chebi: "CHEBI:35181"
          - kegg.compound: "C11355"
          - metanetx.chemical: "MNXM1458"
          - sbo: "SBO:0000247"
      - deltaG: -42.22
    - !!omap
      - id: "s_0270"
      - name: "4-amino-5-hydroxymethyl-2-methylpyrimidine"
      - compartment: "c"
      - formula: "C6H9N3O"
      - charge: 0
      - smiles: "CC1=NC=C(C(=N1)N)CO"
      - annotation: !!omap
          - bigg.metabolite: "4ahmmp"
          - chebi: "CHEBI:16892"
          - kegg.compound: "C01279"
          - metanetx.chemical: "MNXM874"
          - sbo: "SBO:0000247"
      - deltaG: 105.84
    - !!omap
      - id: "s_0271"
      - name: "4-aminobenzoate"
      - compartment: "c"
      - formula: "C7H6NO2"
      - charge: -1
      - smiles: "C1=CC(=CC=C1C(=O)[O-])N"
      - annotation: !!omap
          - bigg.metabolite: "4abz"
          - chebi: "CHEBI:17836"
          - kegg.compound: "C00568"
          - metanetx.chemical: "MNXM421"
          - sbo: "SBO:0000247"
      - deltaG: 9.24
    - !!omap
      - id: "s_0272"
      - name: "4-aminobenzoate"
      - compartment: "e"
      - formula: "C7H6NO2"
      - charge: -1
      - smiles: "C1=CC(=CC=C1C(=O)[O-])N"
      - annotation: !!omap
          - bigg.metabolite: "4abz"
          - chebi: "CHEBI:17836"
          - kegg.compound: "C00568"
          - metanetx.chemical: "MNXM421"
          - sbo: "SBO:0000247"
      - deltaG: 9.24
    - !!omap
      - id: "s_0273"
      - name: "4-aminobenzoate"
      - compartment: "m"
      - formula: "C7H6NO2"
      - charge: -1
      - smiles: "C1=CC(=CC=C1C(=O)[O-])N"
      - annotation: !!omap
          - bigg.metabolite: "4abz"
          - chebi: "CHEBI:17836"
          - kegg.compound: "C00568"
          - metanetx.chemical: "MNXM421"
          - sbo: "SBO:0000247"
      - deltaG: 9.24
    - !!omap
      - id: "s_0274"
      - name: "4-aminobutanal"
      - compartment: "c"
      - formula: "C4H10NO"
      - charge: 1
      - smiles: "C(CC=O)CN"
      - annotation: !!omap
          - bigg.metabolite: "4abutn"
          - chebi: "CHEBI:58264"
          - kegg.compound: "C00555"
          - metanetx.chemical: "MNXM422"
          - sbo: "SBO:0000247"
      - deltaG: 65.39
    - !!omap
      - id: "s_0275"
      - name: "4-aminobutanal"
      - compartment: "m"
      - formula: "C4H10NO"
      - charge: 1
      - smiles: "C(CC=O)CN"
      - annotation: !!omap
          - bigg.metabolite: "4abutn"
          - chebi: "CHEBI:58264"
          - kegg.compound: "C00555"
          - metanetx.chemical: "MNXM422"
          - sbo: "SBO:0000247"
      - deltaG: 65.39
    - !!omap
      - id: "s_0279"
      - name: "4-diphospho-1D-myo-inositol pentakisphosphate"
      - compartment: "c"
      - formula: "C6H6O27P7"
      - charge: -13
      - smiles: "C1(C(C(C(C(C1OP(=O)(O)O)OP(=O)(O)O)OP(=O)(O)OP(=O)(O)O)OP(=O)(O)O)OP(=O)(O)O)OP(=O)(O)O"
      - annotation: !!omap
          - chebi: "CHEBI:53064"
          - kegg.compound: "C11526"
          - metanetx.chemical: "MNXM1715"
          - sbo: "SBO:0000247"
      - deltaG: -1571.48
    - !!omap
      - id: "s_0280"
      - name: "4-guanidinobutanamide"
      - compartment: "c"
      - formula: "C5H13N4O"
      - charge: 1
      - smiles: "C(CC(=O)N)CN=C(N)N"
      - annotation: !!omap
          - bigg.metabolite: "4gudbd"
          - chebi: "CHEBI:58365"
          - kegg.compound: "C03078"
          - metanetx.chemical: "MNXM2617"
          - sbo: "SBO:0000247"
      - deltaG: 96.23
    - !!omap
      - id: "s_0281"
      - name: "4-guanidinobutanoic acid"
      - compartment: "c"
      - formula: "C5H11N3O2"
      - charge: 0
      - smiles: "C(CC(=O)O)CN=C(N)N"
      - annotation: !!omap
          - bigg.metabolite: "4gudbutn"
          - chebi: "CHEBI:15728"
          - kegg.compound: "C01035"
          - metanetx.chemical: "MNXM1312"
          - sbo: "SBO:0000247"
      - deltaG: 30.23
    - !!omap
      - id: "s_0282"
      - name: "4-hydroxy-2-oxoglutarate"
      - compartment: "c"
      - formula: "C5H4O6"
      - charge: -2
      - smiles: "C(C(C(=O)O)O)C(=O)C(=O)O"
      - annotation: !!omap
          - bigg.metabolite: "4h2oglt"
          - chebi: "CHEBI:17742"
          - kegg.compound: "C01127"
          - metanetx.chemical: "MNXM894"
          - sbo: "SBO:0000247"
      - deltaG: -188.77
    - !!omap
      - id: "s_0283"
      - name: "4-hydroxy-2-oxoglutarate"
      - compartment: "m"
      - formula: "C5H4O6"
      - charge: -2
      - smiles: "C(C(C(=O)O)O)C(=O)C(=O)O"
      - annotation: !!omap
          - bigg.metabolite: "4h2oglt"
          - chebi: "CHEBI:17742"
          - kegg.compound: "C01127"
          - metanetx.chemical: "MNXM894"
          - sbo: "SBO:0000247"
      - deltaG: -188.77
    - !!omap
      - id: "s_0284"
      - name: "4-hydroxy-2-oxoglutarate"
      - compartment: "p"
      - formula: "C5H4O6"
      - charge: -2
      - smiles: "C(C(C(=O)O)O)C(=O)C(=O)O"
      - annotation: !!omap
          - bigg.metabolite: "4h2oglt"
          - chebi: "CHEBI:17742"
          - kegg.compound: "C01127"
          - metanetx.chemical: "MNXM894"
          - sbo: "SBO:0000247"
      - deltaG: -188.77
    - !!omap
      - id: "s_0285"
      - name: "4-hydroxy-L-threonine"
      - compartment: "c"
      - formula: "C4H9NO4"
      - charge: 0
      - smiles: "C(C(C(C(=O)O)N)O)O"
      - annotation: !!omap
          - bigg.metabolite: "4hthr"
          - chebi: "CHEBI:28330"
          - kegg.compound: "C06056"
          - metanetx.chemical: "MNXM1802"
          - sbo: "SBO:0000247"
      - deltaG: -72.94
    - !!omap
      - id: "s_0286"
      - name: "4-hydroxybenzoate"
      - compartment: "c"
      - formula: "C7H5O3"
      - charge: -1
      - smiles: "C1=CC(=CC=C1C(=O)O)[O-]"
      - annotation: !!omap
          - bigg.metabolite: "4hbz"
          - chebi: "CHEBI:17879"
          - kegg.compound: "C00156"
          - metanetx.chemical: "MNXM164"
          - sbo: "SBO:0000247"
      - deltaG: -44.04
    - !!omap
      - id: "s_0287"
      - name: "4-hydroxybenzoate"
      - compartment: "m"
      - formula: "C7H5O3"
      - charge: -1
      - smiles: "C1=CC(=CC=C1C(=O)O)[O-]"
      - annotation: !!omap
          - bigg.metabolite: "4hbz"
          - chebi: "CHEBI:17879"
          - kegg.compound: "C00156"
          - metanetx.chemical: "MNXM164"
          - sbo: "SBO:0000247"
      - deltaG: -44.04
    - !!omap
      - id: "s_0290"
      - name: "4-hydroxybenzoyl-CoA"
      - compartment: "m"
      - formula: "C28H36N7O18P3S"
      - charge: -4
      - smiles: "CC(C)(COP(=O)(O)OP(=O)(O)OCC1C(C(C(O1)N2C=NC3=C(N=CN=C32)N)O)OP(=O)(O)O)C(C(=O)NCCC(=O)NCCSC(=O)C4=CC=C(C=C4)O)O"
      - annotation: !!omap
          - bigg.metabolite: "4hbzcoa"
          - chebi: "CHEBI:57356"
          - kegg.compound: "C02949"
          - metanetx.chemical: "MNXM739"
          - sbo: "SBO:0000247"
      - deltaG: 10000000
    - !!omap
      - id: "s_0291"
      - name: "4-methyl-2-oxopentanoate"
      - compartment: "c"
      - formula: "C6H9O3"
      - charge: -1
      - smiles: "CC(C)CC(=O)C(=O)O"
      - annotation: !!omap
          - bigg.metabolite: "4mop"
          - chebi: "CHEBI:17865"
          - kegg.compound: "C00233"
          - metanetx.chemical: "MNXM404"
          - sbo: "SBO:0000247"
      - deltaG: -22.2
    - !!omap
      - id: "s_0292"
      - name: "4-methyl-2-oxopentanoate"
      - compartment: "m"
      - formula: "C6H9O3"
      - charge: -1
      - smiles: "CC(C)CC(=O)C(=O)O"
      - annotation: !!omap
          - bigg.metabolite: "4mop"
          - chebi: "CHEBI:17865"
          - kegg.compound: "C00233"
          - metanetx.chemical: "MNXM404"
          - sbo: "SBO:0000247"
      - deltaG: -22.2
    - !!omap
      - id: "s_0293"
      - name: "4-methyl-5-(2-phosphonooxyethyl)thiazole"
      - compartment: "c"
      - formula: "C6H8NO4PS"
      - charge: -2
      - smiles: "CC1=C(SC=N1)CCOP(=O)(O)O"
      - annotation: !!omap
          - bigg.metabolite: "4mpetz"
          - chebi: "CHEBI:58296"
          - kegg.compound: "C04327"
          - metanetx.chemical: "MNXM960"
          - sbo: "SBO:0000247"
      - deltaG: 10000000
    - !!omap
      - id: "s_0294"
      - name: "4-methylthio-2-oxobutanoate"
      - compartment: "c"
      - formula: "C5H7O3S"
      - charge: -1
      - smiles: "CSCCC(=O)C(=O)O"
      - annotation: !!omap
          - bigg.metabolite: "4met2obut"
          - chebi: "CHEBI:16723"
          - kegg.compound: "C01180"
          - metanetx.chemical: "MNXM276"
          - sbo: "SBO:0000247"
      - deltaG: -32.72
    - !!omap
      - id: "s_0295"
      - name: "4-phospho-L-aspartate"
      - compartment: "c"
      - formula: "C4H6NO7P"
      - charge: -2
      - smiles: "C(C(C(=O)O)N)C(=O)OP(=O)(O)O"
      - annotation: !!omap
          - bigg.metabolite: "4pasp"
          - chebi: "CHEBI:30407"
          - kegg.compound: "C03082"
          - metanetx.chemical: "MNXM1177"
          - sbo: "SBO:0000247"
      - deltaG: -325.03
    - !!omap
      - id: "s_0296"
      - name: "4alpha-methylzymosterol"
      - compartment: "c"
      - formula: "C28H46O"
      - charge: 0
      - smiles: "CC1C2CCC3=C(C2(CCC1O)C)CCC4(C3CCC4C(C)CCC=C(C)C)C"
      - annotation: !!omap
          - bigg.metabolite: "4mzym"
          - chebi: "CHEBI:1949"
          - kegg.compound: "C05103"
          - metanetx.chemical: "MNXM1804"
          - sbo: "SBO:0000247"
      - deltaG: 492.1
    - !!omap
      - id: "s_0297"
      - name: "4beta-methylzymosterol-4alpha-carboxylic acid"
      - compartment: "c"
      - formula: "C29H45O3"
      - charge: -1
      - smiles: "CC(CCC=C(C)C)C1CCC2C1(CCC3=C2CCC4C3(CCC(C4(C)C(=O)O)O)C)C"
      - annotation: !!omap
          - chebi: "CHEBI:50591"
          - kegg.compound: "C15808"
          - metanetx.chemical: "MNXM37762"
          - sbo: "SBO:0000247"
      - deltaG: 401.39
    - !!omap
      - id: "s_0298"
      - name: "5'-adenylyl sulfate"
      - compartment: "c"
      - formula: "C10H12N5O10PS"
      - charge: -2
      - smiles: "C1=NC(=C2C(=N1)N(C=N2)C3C(C(C(O3)COP(=O)(O)OS(=O)(=O)O)O)O)N"
      - annotation: !!omap
          - bigg.metabolite: "aps"
          - chebi: "CHEBI:58243"
          - kegg.compound: "C00224"
          - metanetx.chemical: "MNXM287"
          - sbo: "SBO:0000247"
      - deltaG: -260.26
    - !!omap
      - id: "s_0299"
      - name: "5'-phosphoribosyl-4-(N-succinocarboxamide)-5-aminoimidazole"
      - compartment: "c"
      - formula: "C13H15N4O12P"
      - charge: -4
      - smiles: "C1=NC(=C(N1C2C(C(C(O2)COP(=O)(O)O)O)O)N)C(=O)NC(CC(=O)O)C(=O)O"
      - annotation: !!omap
          - bigg.metabolite: "25aics"
          - chebi: "CHEBI:58443"
          - kegg.compound: "C04823"
          - metanetx.chemical: "MNXM607"
          - sbo: "SBO:0000247"
      - deltaG: -344.7
    - !!omap
      - id: "s_0300"
      - name: "5'-phosphoribosyl-5-aminoimidazole"
      - compartment: "c"
      - formula: "C8H13N3O7P"
      - charge: -1
      - smiles: "C1=C(N(C=N1)C2C(C(C(O2)COP(=O)(O)O)O)O)N"
      - annotation: !!omap
          - bigg.metabolite: "air"
          - chebi: "CHEBI:58592"
          - kegg.compound: "C03373"
          - metanetx.chemical: "MNXM388"
          - sbo: "SBO:0000247"
      - deltaG: -172.89
    - !!omap
      - id: "s_0301"
      - name: "5'-phosphoribosyl-N-formylglycineamide"
      - compartment: "c"
      - formula: "C8H13N2O9P"
      - charge: -2
      - smiles: "C(C1C(C(C(O1)NC(=O)CNC=O)O)O)OP(=O)(O)O"
      - annotation: !!omap
          - bigg.metabolite: "fgam"
          - chebi: "CHEBI:58426"
          - kegg.compound: "C04376"
          - metanetx.chemical: "MNXM453"
          - sbo: "SBO:0000247"
      - deltaG: -291.62
    - !!omap
      - id: "s_0302"
      - name: "5'-phosphoribosyl-N-formylglycineamidine"
      - compartment: "c"
      - formula: "C8H15N3O8P"
      - charge: -1
      - annotation: !!omap
          - bigg.metabolite: "fpram"
          - chebi: "CHEBI:58478"
          - kegg.compound: "C04640"
          - metanetx.chemical: "MNXM568"
          - sbo: "SBO:0000247"
      - deltaG: -237.6
    - !!omap
      - id: "s_0303"
      - name: "5'-S-methyl-5'-thioadenosine"
      - compartment: "c"
      - formula: "C11H15N5O3S"
      - charge: 0
      - smiles: "CSCC1C(C(C(O1)N2C=NC3=C(N=CN=C32)N)O)O"
      - annotation: !!omap
          - bigg.metabolite: "5mta"
          - chebi: "CHEBI:17509"
          - kegg.compound: "C00170"
          - metanetx.chemical: "MNXM150"
          - sbo: "SBO:0000247"
      - deltaG: 138.07
    - !!omap
      - id: "s_0304"
      - name: "5,10-methenyl-THF"
      - compartment: "c"
      - formula: "C20H20N7O6"
      - charge: -1
      - smiles: "C1C2CN(C=[N+]2C3=C(N1)N=C(NC3=O)N)C4=CC=C(C=C4)C(=O)NC(CCC(=O)O)C(=O)[O-]"
      - annotation: !!omap
          - bigg.metabolite: "methf"
          - chebi: "CHEBI:57455"
          - kegg.compound: "C00445"
          - metanetx.chemical: "MNXM511"
          - sbo: "SBO:0000247"
      - deltaG: 91.76
    - !!omap
      - id: "s_0305"
      - name: "5,10-methenyl-THF"
      - compartment: "m"
      - formula: "C20H20N7O6"
      - charge: -1
      - smiles: "C1C2CN(C=[N+]2C3=C(N1)N=C(NC3=O)N)C4=CC=C(C=C4)C(=O)NC(CCC(=O)O)C(=O)[O-]"
      - annotation: !!omap
          - bigg.metabolite: "methf"
          - chebi: "CHEBI:57455"
          - kegg.compound: "C00445"
          - metanetx.chemical: "MNXM511"
          - sbo: "SBO:0000247"
      - deltaG: 91.76
    - !!omap
      - id: "s_0306"
      - name: "5,10-methylenetetrahydrofolate"
      - compartment: "c"
      - formula: "C20H21N7O6"
      - charge: -2
      - smiles: "C1C2CN(CN2C3=C(N1)N=C(NC3=O)N)C4=CC=C(C=C4)C(=O)NC(CCC(=O)O)C(=O)O"
      - annotation: !!omap
          - bigg.metabolite: "mlthf"
          - chebi: "CHEBI:12071"
          - kegg.compound: "C00143"
          - metanetx.chemical: "MNXM183"
          - sbo: "SBO:0000247"
      - deltaG: 109.61
    - !!omap
      - id: "s_0307"
      - name: "5,10-methylenetetrahydrofolate"
      - compartment: "m"
      - formula: "C20H21N7O6"
      - charge: -2
      - smiles: "C1C2CN(CN2C3=C(N1)N=C(NC3=O)N)C4=CC=C(C=C4)C(=O)NC(CCC(=O)O)C(=O)O"
      - annotation: !!omap
          - bigg.metabolite: "mlthf"
          - chebi: "CHEBI:12071"
          - kegg.compound: "C00143"
          - metanetx.chemical: "MNXM183"
          - sbo: "SBO:0000247"
      - deltaG: 109.61
    - !!omap
      - id: "s_0308"
      - name: "5,6,7,8-tetrahydrofolyl-L-glutamic acid"
      - compartment: "c"
      - formula: "C24H27N8O9"
      - charge: -3
      - smiles: "C1C(NC2=C(N1)N=C(NC2=O)N)CNC3=CC=C(C=C3)C(=O)NC(CCC(=O)NC(CCC(=O)O)C(=O)O)C(=O)O"
      - annotation: !!omap
          - bigg.metabolite: "thfglu"
          - chebi: "CHEBI:27650"
          - kegg.compound: "C09332"
          - metanetx.chemical: "MNXM723480"
          - sbo: "SBO:0000247"
      - deltaG: 39.54
    - !!omap
      - id: "s_0309"
      - name: "5,6-bis(diphospho)-1D-myo-inositol tetrakisphosphate"
      - compartment: "c"
      - formula: "C6H7O30P8"
      - charge: -13
      - smiles: "C1(C(C(C(C(C1OP(=O)(O)O)OP(=O)(O)OP(=O)(O)O)OP(=O)(O)OP(=O)(O)O)OP(=O)(O)O)OP(=O)(O)O)OP(=O)(O)O"
      - annotation: !!omap
          - chebi: "CHEBI:52965"
          - metanetx.chemical: "MNXM487477"
          - sbo: "SBO:0000247"
      - deltaG: -1769.75
    - !!omap
      - id: "s_0310"
      - name: "5-(2-hydroxyethyl)-4-methylthiazole"
      - compartment: "c"
      - formula: "C6H9NOS"
      - charge: 0
      - smiles: "CC1=C(SC=N1)CCO"
      - annotation: !!omap
          - bigg.metabolite: "4mhetz"
          - chebi: "CHEBI:17957"
          - kegg.compound: "C04294"
          - metanetx.chemical: "MNXM962"
          - sbo: "SBO:0000247"
      - deltaG: 10000000
    - !!omap
      - id: "s_0311"
      - name: "5-(methylsulfanyl)-2,3-dioxopentyl phosphate"
      - compartment: "c"
      - formula: "C6H9O6PS"
      - charge: -2
      - smiles: "CSCCC(=O)C(=O)COP(=O)(O)O"
      - annotation: !!omap
          - bigg.metabolite: "dkmpp"
          - chebi: "CHEBI:58828"
          - kegg.compound: "C15650"
          - metanetx.chemical: "MNXM162358"
          - sbo: "SBO:0000247"
      - deltaG: -211.52
    - !!omap
      - id: "s_0312"
      - name: "5-[(5-phospho-1-deoxy-D-ribulos-1-ylamino)methylideneamino]-1-(5-phospho-D-ribosyl)imidazole-4-carboxamide"
      - compartment: "c"
      - formula: "C15H21N5O15P2"
      - charge: -4
      - annotation: !!omap
          - bigg.metabolite: "prlp"
          - chebi: "CHEBI:58525"
          - kegg.compound: "C04916"
          - metanetx.chemical: "MNXM1408"
          - sbo: "SBO:0000247"
      - deltaG: -455.74
    - !!omap
      - id: "s_0313"
      - name: "5-amino-6-(5-phosphoribitylamino)uracil"
      - compartment: "c"
      - formula: "C9H15N4O9P"
      - charge: -2
      - smiles: "C(C(C(C(COP(=O)(O)O)O)O)O)NC1=C(C(=O)NC(=O)N1)N"
      - annotation: !!omap
          - bigg.metabolite: "5aprbu"
          - chebi: "CHEBI:58421"
          - kegg.compound: "C04454"
          - metanetx.chemical: "MNXM1178"
          - sbo: "SBO:0000247"
      - deltaG: -256.84
    - !!omap
      - id: "s_0314"
      - name: "5-amino-6-(D-ribitylamino)uracil"
      - compartment: "c"
      - formula: "C9H16N4O6"
      - charge: 0
      - smiles: "C(C(C(C(CO)O)O)O)NC1=C(C(=O)NC(=O)N1)N"
      - annotation: !!omap
          - bigg.metabolite: "4r5au"
          - chebi: "CHEBI:15934"
          - kegg.compound: "C04732"
          - metanetx.chemical: "MNXM791"
          - sbo: "SBO:0000247"
      - deltaG: -33.78
    - !!omap
      - id: "s_0315"
      - name: "5-aminolevulinate"
      - compartment: "c"
      - formula: "C5H9NO3"
      - charge: 0
      - smiles: "C(CC(=O)O)C(=O)CN"
      - annotation: !!omap
          - bigg.metabolite: "5aop"
          - chebi: "CHEBI:17549"
          - kegg.compound: "C00430"
          - metanetx.chemical: "MNXM405"
          - sbo: "SBO:0000247"
      - deltaG: -29.67
    - !!omap
      - id: "s_0316"
      - name: "5-aminolevulinate"
      - compartment: "e"
      - formula: "C5H9NO3"
      - charge: 0
      - smiles: "C(CC(=O)O)C(=O)CN"
      - annotation: !!omap
          - bigg.metabolite: "5aop"
          - chebi: "CHEBI:17549"
          - kegg.compound: "C00430"
          - metanetx.chemical: "MNXM405"
          - sbo: "SBO:0000247"
      - deltaG: -29.67
    - !!omap
      - id: "s_0317"
      - name: "5-aminolevulinate"
      - compartment: "m"
      - formula: "C5H9NO3"
      - charge: 0
      - smiles: "C(CC(=O)O)C(=O)CN"
      - annotation: !!omap
          - bigg.metabolite: "5aop"
          - chebi: "CHEBI:17549"
          - kegg.compound: "C00430"
          - metanetx.chemical: "MNXM405"
          - sbo: "SBO:0000247"
      - deltaG: -29.67
    - !!omap
      - id: "s_0318"
      - name: "5-diphospho-1D-myo-inositol pentakisphosphate"
      - compartment: "c"
      - formula: "C6H6O27P7"
      - charge: -13
      - smiles: "C1(C(C(C(C(C1OP(=O)(O)O)OP(=O)(O)O)OP(=O)(O)OP(=O)(O)O)OP(=O)(O)O)OP(=O)(O)O)OP(=O)(O)O"
      - annotation: !!omap
          - bigg.metabolite: "ppmi12346p"
          - chebi: "CHEBI:58628"
          - kegg.compound: "C11526"
          - metanetx.chemical: "MNXM1715"
          - sbo: "SBO:0000247"
      - deltaG: -1571.48
    - !!omap
      - id: "s_0319"
      - name: "5-formyltetrahydrofolic acid"
      - compartment: "c"
      - formula: "C20H21N7O7"
      - charge: -2
      - smiles: "C1C(N(C2=C(N1)N=C(NC2=O)N)C=O)CNC3=CC=C(C=C3)C(=O)NC(CCC(=O)O)C(=O)O"
      - annotation: !!omap
          - bigg.metabolite: "5fthf"
          - chebi: "CHEBI:57457"
          - kegg.compound: "C03479"
          - metanetx.chemical: "MNXM1392"
          - sbo: "SBO:0000247"
      - deltaG: 75.25
    - !!omap
      - id: "s_0320"
      - name: "5-formyltetrahydrofolic acid"
      - compartment: "e"
      - formula: "C20H21N7O7"
      - charge: -2
      - smiles: "C1C(N(C2=C(N1)N=C(NC2=O)N)C=O)CNC3=CC=C(C=C3)C(=O)NC(CCC(=O)O)C(=O)O"
      - annotation: !!omap
          - bigg.metabolite: "5fthf"
          - chebi: "CHEBI:57457"
          - kegg.compound: "C03479"
          - metanetx.chemical: "MNXM1392"
          - sbo: "SBO:0000247"
      - deltaG: 75.25
    - !!omap
      - id: "s_0321"
      - name: "5-formyltetrahydrofolic acid"
      - compartment: "m"
      - formula: "C20H21N7O7"
      - charge: -2
      - smiles: "C1C(N(C2=C(N1)N=C(NC2=O)N)C=O)CNC3=CC=C(C=C3)C(=O)NC(CCC(=O)O)C(=O)O"
      - annotation: !!omap
          - bigg.metabolite: "5fthf"
          - chebi: "CHEBI:57457"
          - kegg.compound: "C03479"
          - metanetx.chemical: "MNXM1392"
          - sbo: "SBO:0000247"
      - deltaG: 75.25
    - !!omap
      - id: "s_0322"
      - name: "5-methyltetrahydrofolate"
      - compartment: "c"
      - formula: "C20H23N7O6"
      - charge: -2
      - smiles: "CN1C(CNC2=C1C(=O)NC(=N2)N)CNC3=CC=C(C=C3)C(=O)NC(CCC(=O)O)C(=O)O"
      - annotation: !!omap
          - bigg.metabolite: "5mthf"
          - chebi: "CHEBI:18608"
          - kegg.compound: "C00440"
          - metanetx.chemical: "MNXM318"
          - sbo: "SBO:0000247"
      - deltaG: 121.03
    - !!omap
      - id: "s_0323"
      - name: "5-methyltetrahydropteroyltri-L-glutamic acid"
      - compartment: "c"
      - formula: "C30H35N9O12"
      - charge: -4
      - smiles: "CN1C(CNC2=C1C(=O)NC(=N2)N)CNC3=CC=C(C=C3)C(=O)NC(CCC(=O)NC(CCC(=O)NC(CCC(=O)O)C(=O)O)C(=O)O)C(=O)O"
      - annotation: !!omap
          - bigg.metabolite: "mhpglu"
          - chebi: "CHEBI:58207"
          - kegg.compound: "C04489"
          - metanetx.chemical: "MNXM2623"
          - sbo: "SBO:0000247"
      - deltaG: 21.07
    - !!omap
      - id: "s_0324"
      - name: "5-O-(1-carboxyvinyl)-3-phosphoshikimic acid"
      - compartment: "c"
      - formula: "C10H9O10P"
      - charge: -4
      - smiles: "C=C(C(=O)O)OC1CC(=CC(C1O)OP(=O)(O)O)C(=O)O"
      - annotation: !!omap
          - bigg.metabolite: "3psme"
          - chebi: "CHEBI:57701"
          - kegg.compound: "C01269"
          - metanetx.chemical: "MNXM1365"
          - sbo: "SBO:0000247"
      - deltaG: -343.05
    - !!omap
      - id: "s_0325"
      - name: "5-phospho-ribosyl-glycineamide"
      - compartment: "c"
      - formula: "C7H14N2O8P"
      - charge: -1
      - annotation: !!omap
          - bigg.metabolite: "gar"
          - chebi: "CHEBI:58457"
          - kegg.compound: "C03838"
          - metanetx.chemical: "MNXM463"
          - sbo: "SBO:0000247"
      - deltaG: -255.15
    - !!omap
      - id: "s_0326"
      - name: "5-phosphoribosyl-ATP"
      - compartment: "c"
      - formula: "C15H19N5O20P4"
      - charge: -6
      - smiles: "C1=NC2=C(N1C3C(C(C(O3)COP(=O)([O-])OP(=O)([O-])OP(=O)([O-])[O-])O)O)N=CN(C2=N)C4C(C(C(O4)COP(=O)([O-])[O-])O)O"
      - annotation: !!omap
          - bigg.metabolite: "prbatp"
          - chebi: "CHEBI:58424"
          - kegg.compound: "C02739"
          - metanetx.chemical: "MNXM1351"
          - sbo: "SBO:0000247"
      - deltaG: -825.32
    - !!omap
      - id: "s_0327"
      - name: "5-phosphoribosylamine"
      - compartment: "c"
      - formula: "C5H11NO7P"
      - charge: -1
      - smiles: "C(C1C(C(C(O1)N)O)O)OP(=O)(O)O"
      - annotation: !!omap
          - bigg.metabolite: "pram"
          - chebi: "CHEBI:58681"
          - kegg.compound: "C03090"
          - metanetx.chemical: "MNXM90003"
          - sbo: "SBO:0000247"
      - deltaG: -253.8
    - !!omap
      - id: "s_0328"
      - name: "6,7-dimethyl-8-(1-D-ribityl)lumazine"
      - compartment: "c"
      - formula: "C13H17N4O6"
      - charge: -1
      - smiles: "CC1=C(N(C2=NC(=O)NC(=O)C2=N1)CC(C(C(CO)O)O)O)C"
      - annotation: !!omap
          - bigg.metabolite: "dmlz"
          - chebi: "CHEBI:58201"
          - kegg.compound: "C04332"
          - metanetx.chemical: "MNXM1313"
          - sbo: "SBO:0000247"
      - deltaG: 10000000
    - !!omap
      - id: "s_0329"
      - name: "6-(alpha-D-glucosaminyl)-1-phosphatidyl-1D-myo-inositol"
      - compartment: "er"
      - formula: "C17H28NO17PR2"
      - charge: 0
      - smiles: "C(C1C(C(C(C(O1)OC2C(C(C(C(C2OP(=O)(O)OCC(COC=O)OC=O)O)O)O)O)N)O)O)O"
      - annotation: !!omap
          - bigg.metabolite: "gpail_hs"
          - chebi: "CHEBI:17049"
          - kegg.compound: "C04248"
          - metanetx.chemical: "MNXM1411"
          - sbo: "SBO:0000247"
      - deltaG: 10000000
    - !!omap
      - id: "s_0330"
      - name: "6-(alpha-D-glucosaminyl)-O-acyl-1-phosphatidyl-1D-myo-inositol"
      - compartment: "er"
      - charge: 0
      - annotation: !!omap
          - chebi: "CHEBI:53055"
          - metanetx.chemical: "MNXM38512"
          - sbo: "SBO:0000247"
      - deltaG: 10000000
    - !!omap
      - id: "s_0331"
      - name: "6-(N-acetyl-alpha-D-glucosaminyl)-1-phosphatidyl-1D-myo-inositol"
      - compartment: "er"
      - formula: "C19H30NO18PR2"
      - charge: 0
      - smiles: "CC(=O)NC1C(C(C(OC1OC2C(C(C(C(C2OP(=O)(O)OCC(COC(=O)C)OC(=O)C)O)O)O)O)CO)O)O"
      - annotation: !!omap
          - bigg.metabolite: "acgpail_hs"
          - chebi: "CHEBI:57265"
          - kegg.compound: "C01288"
          - metanetx.chemical: "MNXM999"
          - sbo: "SBO:0000247"
      - deltaG: 10000000
    - !!omap
      - id: "s_0332"
      - name: "6-[6-(2-aminoethylphosphoryl)-alpha-mannosyl-(1->6)-2-(2-aminoethylphosphoryl)-alpha-mannosyl-(1->6)-alpha-2-(2-aminoethylphosphoryl)mannosyl-(1->4)-alpha-glucosaminyl]-O-acyl-1-phosphatidyl-1D-myo-inositol"
      - compartment: "er"
      - formula: "C43H44N9O"
      - charge: 0
      - annotation: !!omap
          - chebi: "CHEBI:53060"
          - metanetx.chemical: "MNXM38542"
          - sbo: "SBO:0000247"
      - deltaG: 10000000
    - !!omap
      - id: "s_0333"
      - name: "6-diphospho-1D-myo-inositol pentakisphosphate"
      - compartment: "c"
      - formula: "C6H6O27P7"
      - charge: -13
      - smiles: "C1(C(C(C(C(C1OP(=O)(O)O)OP(=O)(O)O)OP(=O)(O)OP(=O)(O)O)OP(=O)(O)O)OP(=O)(O)O)OP(=O)(O)O"
      - annotation: !!omap
          - chebi: "CHEBI:53065"
          - kegg.compound: "C11526"
          - metanetx.chemical: "MNXM1715"
          - sbo: "SBO:0000247"
      - deltaG: -1571.48
    - !!omap
      - id: "s_0334"
      - name: "6-O-[alpha-D-mannosyl-(1->4)-alpha-D-glucosaminyl]-O-acyl-1-phosphatidyl-1D-myo-inositol"
      - compartment: "er"
      - charge: 0
      - annotation: !!omap
          - chebi: "CHEBI:53056"
          - metanetx.chemical: "MNXM1644"
          - sbo: "SBO:0000247"
      - deltaG: 10000000
    - !!omap
      - id: "s_0335"
      - name: "6-O-phosphono-D-glucono-1,5-lactone"
      - compartment: "c"
      - formula: "C6H9O9P"
      - charge: -2
      - smiles: "C(C1C(C(C(C(=O)O1)O)O)O)OP(=O)(O)O"
      - annotation: !!omap
          - bigg.metabolite: "6pgl"
          - chebi: "CHEBI:57955"
          - kegg.compound: "C01236"
          - metanetx.chemical: "MNXM429"
          - sbo: "SBO:0000247"
      - deltaG: -341.7
    - !!omap
      - id: "s_0337"
      - name: "6-O-{2-O-[(2-aminoethyl)phosphoryl]-alpha-D-mannosyl-(1->4)-alpha-D-glucosaminyl}-O-acyl-1-phosphatidyl-1D-myo-inositol"
      - compartment: "er"
      - charge: 0
      - annotation: !!omap
          - chebi: "CHEBI:53057"
          - metanetx.chemical: "MNXM38904"
          - sbo: "SBO:0000247"
      - deltaG: 10000000
    - !!omap
      - id: "s_0338"
      - name: "6-O-{alpha-D-mannosyl-(1->2)-alpha-D-mannosyl-(1->6)-2-O-[(2-aminoethyl)phosphoryl]-alpha-D-mannosyl-(1->4)-alpha-D-glucosaminyl}-O-acyl-1-phosphatidyl-1D-myo-inositol"
      - compartment: "er"
      - charge: 0
      - annotation: !!omap
          - chebi: "CHEBI:53059"
          - metanetx.chemical: "MNXM1179"
          - sbo: "SBO:0000247"
      - deltaG: 10000000
    - !!omap
      - id: "s_0339"
      - name: "6-O-{alpha-D-mannosyl-(1->6)-2-O-[(2-aminoethyl)phosphoryl]-alpha-D-mannosyl-(1->4)-alpha-D-glucosaminyl}-O-acyl-1-phosphatidyl-1D-myo-inositol"
      - compartment: "er"
      - charge: 0
      - annotation: !!omap
          - chebi: "CHEBI:53058"
          - metanetx.chemical: "MNXM1139"
          - sbo: "SBO:0000247"
      - deltaG: 10000000
    - !!omap
      - id: "s_0340"
      - name: "6-phospho-D-gluconate"
      - compartment: "c"
      - formula: "C6H10O10P"
      - charge: -3
      - smiles: "C(C(C(C(C(C(=O)O)O)O)O)O)OP(=O)(O)O"
      - annotation: !!omap
          - bigg.metabolite: "6pgc"
          - chebi: "CHEBI:58759"
          - kegg.compound: "C00345"
          - metanetx.chemical: "MNXM325"
          - sbo: "SBO:0000247"
      - deltaG: -385.57
    - !!omap
      - id: "s_0341"
      - name: "7,8-diaminononanoate"
      - compartment: "c"
      - formula: "C9H21N2O2"
      - charge: 1
      - smiles: "CC(C(CCCCCC(=O)O)N)N"
      - annotation: !!omap
          - bigg.metabolite: "dann"
          - chebi: "CHEBI:17830"
          - kegg.compound: "C01037"
          - metanetx.chemical: "MNXM1140"
          - sbo: "SBO:0000247"
      - deltaG: 110.26
    - !!omap
      - id: "s_0342"
      - name: "7,8-diaminononanoate"
      - compartment: "e"
      - formula: "C9H21N2O2"
      - charge: 1
      - smiles: "CC(C(CCCCCC(=O)O)N)N"
      - annotation: !!omap
          - bigg.metabolite: "dann"
          - chebi: "CHEBI:17830"
          - kegg.compound: "C01037"
          - metanetx.chemical: "MNXM1140"
          - sbo: "SBO:0000247"
      - deltaG: 110.26
    - !!omap
      - id: "s_0343"
      - name: "7,8-dihydroneopterin"
      - compartment: "c"
      - formula: "C9H13N5O4"
      - charge: 0
      - smiles: "C1C(=NC2=C(N1)N=C(NC2=O)N)C(C(CO)O)O"
      - annotation: !!omap
          - bigg.metabolite: "dhnpt"
          - chebi: "CHEBI:17001"
          - kegg.compound: "C04874"
          - metanetx.chemical: "MNXM162359"
          - sbo: "SBO:0000247"
      - deltaG: 50.4
    - !!omap
      - id: "s_0344"
      - name: "7,8-dihydroneopterin"
      - compartment: "m"
      - formula: "C9H13N5O4"
      - charge: 0
      - smiles: "C1C(=NC2=C(N1)N=C(NC2=O)N)C(C(CO)O)O"
      - annotation: !!omap
          - bigg.metabolite: "dhnpt"
          - chebi: "CHEBI:17001"
          - kegg.compound: "C04874"
          - metanetx.chemical: "MNXM162359"
          - sbo: "SBO:0000247"
      - deltaG: 50.4
    - !!omap
      - id: "s_0345"
      - name: "7,8-dihydroneopterin 3'-phosphate"
      - compartment: "c"
      - formula: "C9H12N5O7P"
      - charge: -2
      - smiles: "C1C(=NC2=C(N1)N=C(NC2=O)N)C(C(COP(=O)(O)O)O)O"
      - annotation: !!omap
          - bigg.metabolite: "dhpmp"
          - chebi: "CHEBI:58762"
          - kegg.compound: "C05925"
          - metanetx.chemical: "MNXM1645"
          - sbo: "SBO:0000247"
      - deltaG: -172.96
    - !!omap
      - id: "s_0346"
      - name: "7,8-dihydroneopterin 3'-triphosphate"
      - compartment: "c"
      - formula: "C9H12N5O13P3"
      - charge: -4
      - smiles: "C1C(=NC2=C(N1)N=C(NC2=O)N)C(C(COP(=O)(O)OP(=O)(O)OP(=O)(O)O)O)O"
      - annotation: !!omap
          - bigg.metabolite: "ahdt"
          - chebi: "CHEBI:58462"
          - kegg.compound: "C04895"
          - metanetx.chemical: "MNXM397"
          - sbo: "SBO:0000247"
      - deltaG: -591.65
    - !!omap
      - id: "s_0347"
      - name: "7,8-dihydropteroate"
      - compartment: "c"
      - formula: "C14H13N6O3"
      - charge: -1
      - smiles: "C1C(=NC2=C(N1)N=C(NC2=O)N)CNC3=CC=C(C=C3)C(=O)O"
      - annotation: !!omap
          - bigg.metabolite: "dhpt"
          - chebi: "CHEBI:17839"
          - kegg.compound: "C00921"
          - metanetx.chemical: "MNXM722728"
          - sbo: "SBO:0000247"
      - deltaG: 123.88
    - !!omap
      - id: "s_0348"
      - name: "7,8-dihydropteroate"
      - compartment: "m"
      - formula: "C14H13N6O3"
      - charge: -1
      - smiles: "C1C(=NC2=C(N1)N=C(NC2=O)N)CNC3=CC=C(C=C3)C(=O)O"
      - annotation: !!omap
          - bigg.metabolite: "dhpt"
          - chebi: "CHEBI:17839"
          - kegg.compound: "C00921"
          - metanetx.chemical: "MNXM722728"
          - sbo: "SBO:0000247"
      - deltaG: 123.88
    - !!omap
      - id: "s_0349"
      - name: "7-phospho-2-dehydro-3-deoxy-D-arabino-heptonic acid"
      - compartment: "c"
      - formula: "C7H10O10P"
      - charge: -3
      - smiles: "C(C(C(C(COP(=O)(O)O)O)O)O)C(=O)C(=O)O"
      - annotation: !!omap
          - bigg.metabolite: "2dda7p"
          - chebi: "CHEBI:58394"
          - kegg.compound: "C04691"
          - metanetx.chemical: "MNXM1219"
          - sbo: "SBO:0000247"
      - deltaG: -373.47
    - !!omap
      - id: "s_0350"
      - name: "7-phospho-2-dehydro-3-deoxy-D-arabino-heptonic acid"
      - compartment: "m"
      - formula: "C7H10O10P"
      - charge: -3
      - smiles: "C(C(C(C(COP(=O)(O)O)O)O)O)C(=O)C(=O)O"
      - annotation: !!omap
          - bigg.metabolite: "2dda7p"
          - chebi: "CHEBI:58394"
          - kegg.compound: "C04691"
          - metanetx.chemical: "MNXM1219"
          - sbo: "SBO:0000247"
      - deltaG: -373.47
    - !!omap
      - id: "s_0352"
      - name: "8-amino-7-oxononanoate"
      - compartment: "c"
      - formula: "C9H17NO3"
      - charge: 0
      - smiles: "CC(C(=O)CCCCCC(=O)[O-])N"
      - annotation: !!omap
          - bigg.metabolite: "8aonn"
          - chebi: "CHEBI:15830"
          - kegg.compound: "C01092"
          - metanetx.chemical: "MNXM1141"
          - sbo: "SBO:0000247"
      - deltaG: 53.44
    - !!omap
      - id: "s_0354"
      - name: "8-amino-7-oxononanoate"
      - compartment: "e"
      - formula: "C9H17NO3"
      - charge: 0
      - smiles: "CC(C(=O)CCCCCC(=O)[O-])N"
      - annotation: !!omap
          - bigg.metabolite: "8aonn"
          - chebi: "CHEBI:15830"
          - kegg.compound: "C01092"
          - metanetx.chemical: "MNXM1141"
          - sbo: "SBO:0000247"
      - deltaG: 53.44
    - !!omap
      - id: "s_0357"
      - name: "9H-xanthine"
      - compartment: "c"
      - formula: "C5H4N4O2"
      - charge: 0
      - smiles: "C1=NC2=C(N1)C(=O)NC(=O)N2"
      - annotation: !!omap
          - bigg.metabolite: "xan"
          - chebi: "CHEBI:17712"
          - kegg.compound: "C00385"
          - metanetx.chemical: "MNXM174"
          - sbo: "SBO:0000247"
      - deltaG: -1.51
    - !!omap
      - id: "s_0358"
      - name: "9H-xanthine"
      - compartment: "e"
      - formula: "C5H4N4O2"
      - charge: 0
      - smiles: "C1=NC2=C(N1)C(=O)NC(=O)N2"
      - annotation: !!omap
          - bigg.metabolite: "xan"
          - chebi: "CHEBI:17712"
          - kegg.compound: "C00385"
          - metanetx.chemical: "MNXM174"
          - sbo: "SBO:0000247"
      - deltaG: -1.51
    - !!omap
      - id: "s_0359"
      - name: "acetaldehyde"
      - compartment: "c"
      - formula: "C2H4O"
      - charge: 0
      - smiles: "CC=O"
      - annotation: !!omap
          - bigg.metabolite: "acald"
          - chebi: "CHEBI:15343"
          - kegg.compound: "C00084"
          - metanetx.chemical: "MNXM75"
          - sbo: "SBO:0000247"
      - deltaG: 5.55
    - !!omap
      - id: "s_0360"
      - name: "acetaldehyde"
      - compartment: "e"
      - formula: "C2H4O"
      - charge: 0
      - smiles: "CC=O"
      - annotation: !!omap
          - bigg.metabolite: "acald"
          - chebi: "CHEBI:15343"
          - kegg.compound: "C00084"
          - metanetx.chemical: "MNXM75"
          - sbo: "SBO:0000247"
      - deltaG: 5.55
    - !!omap
      - id: "s_0361"
      - name: "acetaldehyde"
      - compartment: "m"
      - formula: "C2H4O"
      - charge: 0
      - smiles: "CC=O"
      - annotation: !!omap
          - bigg.metabolite: "acald"
          - chebi: "CHEBI:15343"
          - kegg.compound: "C00084"
          - metanetx.chemical: "MNXM75"
          - sbo: "SBO:0000247"
      - deltaG: 5.55
    - !!omap
      - id: "s_0362"
      - name: "acetate"
      - compartment: "c"
      - formula: "C2H3O2"
      - charge: -1
      - smiles: "CC(=O)[O-]"
      - annotation: !!omap
          - bigg.metabolite: "ac"
          - chebi: "CHEBI:30089"
          - kegg.compound: "C00033"
          - metanetx.chemical: "MNXM26"
          - sbo: "SBO:0000247"
      - deltaG: -59.27
    - !!omap
      - id: "s_0363"
      - name: "acetate"
      - compartment: "er"
      - formula: "C2H3O2"
      - charge: -1
      - smiles: "CC(=O)[O-]"
      - annotation: !!omap
          - bigg.metabolite: "ac"
          - chebi: "CHEBI:30089"
          - kegg.compound: "C00033"
          - metanetx.chemical: "MNXM26"
          - sbo: "SBO:0000247"
      - deltaG: -59.27
    - !!omap
      - id: "s_0364"
      - name: "acetate"
      - compartment: "e"
      - formula: "C2H3O2"
      - charge: -1
      - smiles: "CC(=O)[O-]"
      - annotation: !!omap
          - bigg.metabolite: "ac"
          - chebi: "CHEBI:30089"
          - kegg.compound: "C00033"
          - metanetx.chemical: "MNXM26"
          - sbo: "SBO:0000247"
      - deltaG: -59.27
    - !!omap
      - id: "s_0365"
      - name: "acetate"
      - compartment: "m"
      - formula: "C2H3O2"
      - charge: -1
      - smiles: "CC(=O)[O-]"
      - annotation: !!omap
          - bigg.metabolite: "ac"
          - chebi: "CHEBI:30089"
          - kegg.compound: "C00033"
          - metanetx.chemical: "MNXM26"
          - sbo: "SBO:0000247"
      - deltaG: -59.27
    - !!omap
      - id: "s_0366"
      - name: "acetate"
      - compartment: "n"
      - formula: "C2H3O2"
      - charge: -1
      - smiles: "CC(=O)[O-]"
      - annotation: !!omap
          - bigg.metabolite: "ac"
          - chebi: "CHEBI:30089"
          - kegg.compound: "C00033"
          - metanetx.chemical: "MNXM26"
          - sbo: "SBO:0000247"
      - deltaG: -59.27
    - !!omap
      - id: "s_0367"
      - name: "acetoacetyl-CoA"
      - compartment: "c"
      - formula: "C25H36N7O18P3S"
      - charge: -4
      - smiles: "CC(=O)CC(=O)SCCNC(=O)CCNC(=O)C(C(C)(C)COP(=O)(O)OP(=O)(O)OCC1C(C(C(O1)N2C=NC3=C(N=CN=C32)N)O)OP(=O)(O)O)O"
      - annotation: !!omap
          - bigg.metabolite: "aacoa"
          - chebi: "CHEBI:57286"
          - kegg.compound: "C00332"
          - metanetx.chemical: "MNXM133"
          - sbo: "SBO:0000247"
      - deltaG: -475.88
    - !!omap
      - id: "s_0370"
      - name: "acetoacetyl-CoA"
      - compartment: "m"
      - formula: "C25H36N7O18P3S"
      - charge: -4
      - smiles: "CC(=O)CC(=O)SCCNC(=O)CCNC(=O)C(C(C)(C)COP(=O)(O)OP(=O)(O)OCC1C(C(C(O1)N2C=NC3=C(N=CN=C32)N)O)OP(=O)(O)O)O"
      - annotation: !!omap
          - bigg.metabolite: "aacoa"
          - chebi: "CHEBI:57286"
          - kegg.compound: "C00332"
          - metanetx.chemical: "MNXM133"
          - sbo: "SBO:0000247"
      - deltaG: -475.88
    - !!omap
      - id: "s_0372"
      - name: "acetyl-ACP"
      - compartment: "m"
      - formula: "C2H3OSR"
      - charge: 0
      - annotation: !!omap
          - bigg.metabolite: "acACP"
          - chebi: "CHEBI:17093"
          - kegg.compound: "C03939"
          - metanetx.chemical: "MNXM1269"
          - sbo: "SBO:0000247"
      - deltaG: 10000000
    - !!omap
      - id: "s_0373"
      - name: "acetyl-CoA"
      - compartment: "c"
      - formula: "C23H34N7O17P3S"
      - charge: -4
      - smiles: "CC(=O)SCCNC(=O)CCNC(=O)C(C(C)(C)COP(=O)(O)OP(=O)(O)OCC1C(C(C(O1)N2C=NC3=C(N=CN=C32)N)O)OP(=O)(O)O)O"
      - annotation: !!omap
          - bigg.metabolite: "accoa"
          - chebi: "CHEBI:57288"
          - kegg.compound: "C00024"
          - metanetx.chemical: "MNXM21"
          - sbo: "SBO:0000247"
      - deltaG: -468.58
    - !!omap
      - id: "s_0376"
      - name: "acetyl-CoA"
      - compartment: "m"
      - formula: "C23H34N7O17P3S"
      - charge: -4
      - smiles: "CC(=O)SCCNC(=O)CCNC(=O)C(C(C)(C)COP(=O)(O)OP(=O)(O)OCC1C(C(C(O1)N2C=NC3=C(N=CN=C32)N)O)OP(=O)(O)O)O"
      - annotation: !!omap
          - bigg.metabolite: "accoa"
          - chebi: "CHEBI:57288"
          - kegg.compound: "C00024"
          - metanetx.chemical: "MNXM21"
          - sbo: "SBO:0000247"
      - deltaG: -468.58
    - !!omap
      - id: "s_0377"
      - name: "acetyl-CoA"
      - compartment: "n"
      - formula: "C23H34N7O17P3S"
      - charge: -4
      - smiles: "CC(=O)SCCNC(=O)CCNC(=O)C(C(C)(C)COP(=O)(O)OP(=O)(O)OCC1C(C(C(O1)N2C=NC3=C(N=CN=C32)N)O)OP(=O)(O)O)O"
      - annotation: !!omap
          - bigg.metabolite: "accoa"
          - chebi: "CHEBI:57288"
          - kegg.compound: "C00024"
          - metanetx.chemical: "MNXM21"
          - sbo: "SBO:0000247"
      - deltaG: -468.58
    - !!omap
      - id: "s_0378"
      - name: "acetyl-CoA"
      - compartment: "p"
      - formula: "C23H34N7O17P3S"
      - charge: -4
      - smiles: "CC(=O)SCCNC(=O)CCNC(=O)C(C(C)(C)COP(=O)(O)OP(=O)(O)OCC1C(C(C(O1)N2C=NC3=C(N=CN=C32)N)O)OP(=O)(O)O)O"
      - annotation: !!omap
          - bigg.metabolite: "accoa"
          - chebi: "CHEBI:57288"
          - kegg.compound: "C00024"
          - metanetx.chemical: "MNXM21"
          - sbo: "SBO:0000247"
      - deltaG: -468.58
    - !!omap
      - id: "s_0383"
      - name: "adenine"
      - compartment: "c"
      - formula: "C5H5N5"
      - charge: 0
      - smiles: "C1=NC2=NC=NC(=C2N1)N"
      - annotation: !!omap
          - bigg.metabolite: "ade"
          - chebi: "CHEBI:16708"
          - kegg.compound: "C00147"
          - metanetx.chemical: "MNXM168"
          - sbo: "SBO:0000247"
      - deltaG: 115.32
    - !!omap
      - id: "s_0384"
      - name: "adenine"
      - compartment: "e"
      - formula: "C5H5N5"
      - charge: 0
      - smiles: "C1=NC2=NC=NC(=C2N1)N"
      - annotation: !!omap
          - bigg.metabolite: "ade"
          - chebi: "CHEBI:16708"
          - kegg.compound: "C00147"
          - metanetx.chemical: "MNXM168"
          - sbo: "SBO:0000247"
      - deltaG: 115.32
    - !!omap
      - id: "s_0385"
      - name: "adenine"
      - compartment: "m"
      - formula: "C5H5N5"
      - charge: 0
      - smiles: "C1=NC2=NC=NC(=C2N1)N"
      - annotation: !!omap
          - bigg.metabolite: "ade"
          - chebi: "CHEBI:16708"
          - kegg.compound: "C00147"
          - metanetx.chemical: "MNXM168"
          - sbo: "SBO:0000247"
      - deltaG: 115.32
    - !!omap
      - id: "s_0386"
      - name: "adenosine"
      - compartment: "c"
      - formula: "C10H13N5O4"
      - charge: 0
      - smiles: "C1=NC(=C2C(=N1)N(C=N2)C3C(C(C(O3)CO)O)O)N"
      - annotation: !!omap
          - bigg.metabolite: "adn"
          - chebi: "CHEBI:16335"
          - kegg.compound: "C00212"
          - metanetx.chemical: "MNXM212"
          - sbo: "SBO:0000247"
      - deltaG: 71.97
    - !!omap
      - id: "s_0387"
      - name: "adenosine"
      - compartment: "e"
      - formula: "C10H13N5O4"
      - charge: 0
      - smiles: "C1=NC(=C2C(=N1)N(C=N2)C3C(C(C(O3)CO)O)O)N"
      - annotation: !!omap
          - bigg.metabolite: "adn"
          - chebi: "CHEBI:16335"
          - kegg.compound: "C00212"
          - metanetx.chemical: "MNXM212"
          - sbo: "SBO:0000247"
      - deltaG: 71.97
    - !!omap
      - id: "s_0389"
      - name: "adenosine 2'-phosphate"
      - compartment: "c"
      - formula: "C10H12N5O7P"
      - charge: -2
      - smiles: "C1=NC(=C2C(=N1)N(C=N2)C3C(C(C(O3)CO)O)OP(=O)(O)O)N"
      - annotation: !!omap
          - bigg.metabolite: "amp2p"
          - chebi: "CHEBI:28223"
          - kegg.compound: "C00946"
          - metanetx.chemical: "MNXM7028"
          - sbo: "SBO:0000247"
      - deltaG: -152.71
    - !!omap
      - id: "s_0390"
      - name: "adenosine 3',5'-bismonophosphate"
      - compartment: "c"
      - formula: "C10H11N5O10P2"
      - charge: -4
      - smiles: "C1=NC(=C2C(=N1)N(C=N2)C3C(C(C(O3)COP(=O)(O)O)OP(=O)(O)O)O)N"
      - annotation: !!omap
          - bigg.metabolite: "pap"
          - chebi: "CHEBI:58343"
          - kegg.compound: "C00054"
          - metanetx.chemical: "MNXM45"
          - sbo: "SBO:0000247"
      - deltaG: -376.4
    - !!omap
      - id: "s_0391"
      - name: "adenosine 3',5'-bismonophosphate"
      - compartment: "e"
      - formula: "C10H11N5O10P2"
      - charge: -4
      - smiles: "C1=NC(=C2C(=N1)N(C=N2)C3C(C(C(O3)COP(=O)(O)O)OP(=O)(O)O)O)N"
      - annotation: !!omap
          - bigg.metabolite: "pap"
          - chebi: "CHEBI:58343"
          - kegg.compound: "C00054"
          - metanetx.chemical: "MNXM45"
          - sbo: "SBO:0000247"
      - deltaG: -376.4
    - !!omap
      - id: "s_0392"
      - name: "adenosine 3',5'-bismonophosphate"
      - compartment: "m"
      - formula: "C10H11N5O10P2"
      - charge: -4
      - smiles: "C1=NC(=C2C(=N1)N(C=N2)C3C(C(C(O3)COP(=O)(O)O)OP(=O)(O)O)O)N"
      - annotation: !!omap
          - bigg.metabolite: "pap"
          - chebi: "CHEBI:58343"
          - kegg.compound: "C00054"
          - metanetx.chemical: "MNXM45"
          - sbo: "SBO:0000247"
      - deltaG: -376.4
    - !!omap
      - id: "s_0393"
      - name: "adenylo-succinate"
      - compartment: "c"
      - formula: "C14H14N5O11P"
      - charge: -4
      - smiles: "C1=NC(=C2C(=N1)N(C=N2)C3C(C(C(O3)COP(=O)([O-])[O-])O)O)NC(CC(=O)[O-])C(=O)[O-]"
      - annotation: !!omap
          - bigg.metabolite: "dcamp"
          - chebi: "CHEBI:57567"
          - kegg.compound: "C03794"
          - metanetx.chemical: "MNXM565"
          - sbo: "SBO:0000247"
      - deltaG: -285.92
    - !!omap
      - id: "s_0394"
      - name: "ADP"
      - compartment: "c"
      - formula: "C10H12N5O10P2"
      - charge: -3
      - smiles: "C1=NC(=C2C(=N1)N(C=N2)C3C(C(C(O3)COP(=O)(O)OP(=O)(O)O)O)O)N"
      - annotation: !!omap
          - bigg.metabolite: "adp"
          - chebi: "CHEBI:456216"
          - kegg.compound: "C00008"
          - metanetx.chemical: "MNXM7"
          - sbo: "SBO:0000247"
      - deltaG: -356.59
    - !!omap
      - id: "s_0395"
      - name: "ADP"
      - compartment: "er"
      - formula: "C10H12N5O10P2"
      - charge: -3
      - smiles: "C1=NC(=C2C(=N1)N(C=N2)C3C(C(C(O3)COP(=O)(O)OP(=O)(O)O)O)O)N"
      - annotation: !!omap
          - bigg.metabolite: "adp"
          - chebi: "CHEBI:456216"
          - kegg.compound: "C00008"
          - metanetx.chemical: "MNXM7"
          - sbo: "SBO:0000247"
      - deltaG: -356.59
    - !!omap
      - id: "s_0397"
      - name: "ADP"
      - compartment: "m"
      - formula: "C10H12N5O10P2"
      - charge: -3
      - smiles: "C1=NC(=C2C(=N1)N(C=N2)C3C(C(C(O3)COP(=O)(O)OP(=O)(O)O)O)O)N"
      - annotation: !!omap
          - bigg.metabolite: "adp"
          - chebi: "CHEBI:456216"
          - kegg.compound: "C00008"
          - metanetx.chemical: "MNXM7"
          - sbo: "SBO:0000247"
      - deltaG: -356.59
    - !!omap
      - id: "s_0398"
      - name: "ADP"
      - compartment: "n"
      - formula: "C10H12N5O10P2"
      - charge: -3
      - smiles: "C1=NC(=C2C(=N1)N(C=N2)C3C(C(C(O3)COP(=O)(O)OP(=O)(O)O)O)O)N"
      - annotation: !!omap
          - bigg.metabolite: "adp"
          - chebi: "CHEBI:456216"
          - kegg.compound: "C00008"
          - metanetx.chemical: "MNXM7"
          - sbo: "SBO:0000247"
      - deltaG: -356.59
    - !!omap
      - id: "s_0399"
      - name: "ADP"
      - compartment: "p"
      - formula: "C10H12N5O10P2"
      - charge: -3
      - smiles: "C1=NC(=C2C(=N1)N(C=N2)C3C(C(C(O3)COP(=O)(O)OP(=O)(O)O)O)O)N"
      - annotation: !!omap
          - bigg.metabolite: "adp"
          - chebi: "CHEBI:456216"
          - kegg.compound: "C00008"
          - metanetx.chemical: "MNXM7"
          - sbo: "SBO:0000247"
      - deltaG: -356.59
    - !!omap
      - id: "s_0402"
      - name: "ADP-ribose"
      - compartment: "n"
      - formula: "C15H21N5O14P2"
      - charge: -2
      - smiles: "C1=NC(=C2C(=N1)N(C=N2)C3C(C(C(O3)COP(=O)(O)OP(=O)(O)OCC4C(C(C(O4)O)O)O)O)O)N"
      - annotation: !!omap
          - bigg.metabolite: "adprib"
          - chebi: "CHEBI:57967"
          - kegg.compound: "C00301"
          - metanetx.chemical: "MNXM48596"
          - sbo: "SBO:0000247"
      - deltaG: -394.82
    - !!omap
      - id: "s_0403"
      - name: "AICAR"
      - compartment: "c"
      - formula: "C9H13N4O8P"
      - charge: -2
      - smiles: "C1=NC(=C(N1C2C(C(C(O2)CO)O)O)N)C(=O)N"
      - annotation: !!omap
          - bigg.metabolite: "aicar"
          - chebi: "CHEBI:58475"
          - kegg.compound: "C04677"
          - metanetx.chemical: "MNXM365"
          - sbo: "SBO:0000247"
      - deltaG: -217.27
    - !!omap
      - id: "s_0404"
      - name: "Ala-tRNA(Ala)"
      - compartment: "c"
      - formula: "C3H7NOR"
      - charge: 1
      - annotation: !!omap
          - bigg.metabolite: "alatrna"
          - chebi: "CHEBI:17732"
          - kegg.compound: "C00886"
          - metanetx.chemical: "MNXM89815"
          - sbo: "SBO:0000247"
      - deltaG: 10000000
    - !!omap
      - id: "s_0405"
      - name: "allantoate"
      - compartment: "c"
      - formula: "C4H7N4O4"
      - charge: -1
      - smiles: "C(C(=O)[O-])(NC(=O)N)NC(=O)N"
      - annotation: !!omap
          - bigg.metabolite: "alltt"
          - chebi: "CHEBI:17536"
          - kegg.compound: "C00499"
          - metanetx.chemical: "MNXM584"
          - sbo: "SBO:0000247"
      - deltaG: -98.98
    - !!omap
      - id: "s_0406"
      - name: "allantoate"
      - compartment: "e"
      - formula: "C4H7N4O4"
      - charge: -1
      - smiles: "C(C(=O)[O-])(NC(=O)N)NC(=O)N"
      - annotation: !!omap
          - bigg.metabolite: "alltt"
          - chebi: "CHEBI:17536"
          - kegg.compound: "C00499"
          - metanetx.chemical: "MNXM584"
          - sbo: "SBO:0000247"
      - deltaG: -98.98
    - !!omap
      - id: "s_0407"
      - name: "allantoin"
      - compartment: "c"
      - formula: "C4H6N4O3"
      - charge: 0
      - smiles: "C1(C(=O)NC(=O)N1)NC(=O)N"
      - annotation: !!omap
          - bigg.metabolite: "alltn"
          - chebi: "CHEBI:15676"
          - kegg.compound: "C01551"
          - metanetx.chemical: "MNXM612"
          - sbo: "SBO:0000247"
      - deltaG: -57.75
    - !!omap
      - id: "s_0408"
      - name: "allantoin"
      - compartment: "e"
      - formula: "C4H6N4O3"
      - charge: 0
      - smiles: "C1(C(=O)NC(=O)N1)NC(=O)N"
      - annotation: !!omap
          - bigg.metabolite: "alltn"
          - chebi: "CHEBI:15676"
          - kegg.compound: "C01551"
          - metanetx.chemical: "MNXM612"
          - sbo: "SBO:0000247"
      - deltaG: -57.75
    - !!omap
      - id: "s_0409"
      - name: "alpha,alpha-trehalose 6-phosphate"
      - compartment: "c"
      - formula: "C12H21O14P"
      - charge: -2
      - smiles: "C(C1C(C(C(C(O1)OC2C(C(C(C(O2)COP(=O)(O)O)O)O)O)O)O)O)O"
      - annotation: !!omap
          - bigg.metabolite: "tre6p"
          - chebi: "CHEBI:58429"
          - kegg.compound: "C00689"
          - metanetx.chemical: "MNXM448"
          - sbo: "SBO:0000247"
      - deltaG: -385.59
    - !!omap
      - id: "s_0410"
      - name: "alpha-D-galactose 1-phosphate"
      - compartment: "c"
      - formula: "C6H11O9P"
      - charge: -2
      - smiles: "C(C1C(C(C(C(O1)OP(=O)(O)O)O)O)O)O"
      - annotation: !!omap
          - bigg.metabolite: "gal1p"
          - chebi: "CHEBI:58336"
          - kegg.compound: "C00446"
          - metanetx.chemical: "MNXM336"
          - sbo: "SBO:0000247"
      - deltaG: -324.47
    - !!omap
      - id: "s_0411"
      - name: "alpha-D-glucosamine 1-phosphate"
      - compartment: "c"
      - formula: "C6H13NO8P"
      - charge: -1
      - smiles: "C(C1C(C(C(C(O1)OP(=O)(O)O)N)O)O)O"
      - annotation: !!omap
          - bigg.metabolite: "gam1p"
          - chebi: "CHEBI:58516"
          - kegg.compound: "C06156"
          - metanetx.chemical: "MNXM969"
          - sbo: "SBO:0000247"
      - deltaG: -272.01
    - !!omap
      - id: "s_0412"
      - name: "alpha-D-glucosamine 6-phosphate"
      - compartment: "c"
      - formula: "C6H13NO8P"
      - charge: -1
      - smiles: "C(C1C(C(C(C(O1)O)N)O)O)OP(=O)(O)O"
      - annotation: !!omap
          - bigg.metabolite: "gam6p"
          - chebi: "CHEBI:15873"
          - kegg.compound: "C00352"
          - metanetx.chemical: "MNXM370"
          - sbo: "SBO:0000247"
      - deltaG: -270.97
    - !!omap
      - id: "s_0413"
      - name: "alpha-D-glucosamine 6-phosphate"
      - compartment: "e"
      - formula: "C6H13NO8P"
      - charge: -1
      - smiles: "C(C1C(C(C(C(O1)O)N)O)O)OP(=O)(O)O"
      - annotation: !!omap
          - bigg.metabolite: "gam6p"
          - chebi: "CHEBI:15873"
          - kegg.compound: "C00352"
          - metanetx.chemical: "MNXM370"
          - sbo: "SBO:0000247"
      - deltaG: -270.97
    - !!omap
      - id: "s_0414"
      - name: "alpha-D-mannosyl-beta-D-mannosyldiacetylchitobiosyldiphosphodolichol"
      - compartment: "g"
      - formula: "C48H84N2O27P2"
      - charge: 0
      - smiles: "CC(CCC=C(C)CCC=C(C)CCC=C(C)CCC=C(C)C)CCOP(=O)(O)OP(=O)(O)OC1C(C(C(C(O1)CO)OC2C(C(C(C(O2)CO)OC3C(C(C(C(O3)CO)O)OC4C(C(C(C(O4)CO)O)O)O)O)O)NC(=O)C)O)NC(=O)C"
      - annotation: !!omap
          - chebi: "CHEBI:28067"
          - kegg.compound: "C05861"
          - metanetx.chemical: "MNXM41231"
          - sbo: "SBO:0000247"
      - deltaG: -190.39
    - !!omap
      - id: "s_0415"
      - name: "alpha-D-ribose 1-phosphate(2-)"
      - compartment: "c"
      - formula: "C5H9O8P"
      - charge: -2
      - smiles: "C(C1C(C(C(O1)OP(=O)([O-])[O-])O)O)O"
      - annotation: !!omap
          - bigg.metabolite: "r1p"
          - chebi: "CHEBI:57720"
          - kegg.compound: "C00620"
          - metanetx.chemical: "MNXM295"
          - sbo: "SBO:0000247"
      - deltaG: -307.29
    - !!omap
      - id: "s_0416"
      - name: "alpha-D-ribose 1-phosphate(2-)"
      - compartment: "m"
      - formula: "C5H9O8P"
      - charge: -2
      - smiles: "C(C1C(C(C(O1)OP(=O)([O-])[O-])O)O)O"
      - annotation: !!omap
          - bigg.metabolite: "r1p"
          - chebi: "CHEBI:57720"
          - kegg.compound: "C00620"
          - metanetx.chemical: "MNXM295"
          - sbo: "SBO:0000247"
      - deltaG: -307.29
    - !!omap
      - id: "s_0417"
      - name: "aminoacetaldehyde"
      - compartment: "c"
      - formula: "C2H6NO"
      - charge: 1
      - smiles: "C(C=O)N"
      - annotation: !!omap
          - bigg.metabolite: "aacald"
          - chebi: "CHEBI:58213"
          - kegg.compound: "C06735"
          - metanetx.chemical: "MNXM2212"
          - sbo: "SBO:0000247"
      - deltaG: 13.62
    - !!omap
      - id: "s_0418"
      - name: "aminoacetone"
      - compartment: "c"
      - formula: "C3H8NO"
      - charge: 1
      - smiles: "CC(=O)CN"
      - annotation: !!omap
          - bigg.metabolite: "aact"
          - chebi: "CHEBI:58320"
          - kegg.compound: "C01888"
          - metanetx.chemical: "MNXM1106"
          - sbo: "SBO:0000247"
      - deltaG: 41.02
    - !!omap
      - id: "s_0419"
      - name: "ammonium"
      - compartment: "c"
      - formula: "H4N"
      - charge: 1
      - smiles: "[NH4+]"
      - annotation: !!omap
          - bigg.metabolite: "nh4"
          - chebi: "CHEBI:28938"
          - kegg.compound: "C01342"
          - metanetx.chemical: "MNXM15"
          - sbo: "SBO:0000247"
      - deltaG: 19.78
    - !!omap
      - id: "s_0420"
      - name: "ammonium"
      - compartment: "e"
      - formula: "H4N"
      - charge: 1
      - smiles: "[NH4+]"
      - annotation: !!omap
          - bigg.metabolite: "nh4"
          - chebi: "CHEBI:28938"
          - kegg.compound: "C01342"
          - metanetx.chemical: "MNXM15"
          - sbo: "SBO:0000247"
      - deltaG: 19.78
    - !!omap
      - id: "s_0421"
      - name: "ammonium"
      - compartment: "m"
      - formula: "H4N"
      - charge: 1
      - smiles: "[NH4+]"
      - annotation: !!omap
          - bigg.metabolite: "nh4"
          - chebi: "CHEBI:28938"
          - kegg.compound: "C01342"
          - metanetx.chemical: "MNXM15"
          - sbo: "SBO:0000247"
      - deltaG: 19.78
    - !!omap
      - id: "s_0423"
      - name: "AMP"
      - compartment: "c"
      - formula: "C10H12N5O7P"
      - charge: -2
      - smiles: "C1=NC(=C2C(=N1)N(C=N2)C3C(C(C(O3)COP(=O)(O)O)O)O)N"
      - annotation: !!omap
          - bigg.metabolite: "amp"
          - chebi: "CHEBI:456215"
          - kegg.compound: "C00020"
          - metanetx.chemical: "MNXM14"
          - sbo: "SBO:0000247"
      - deltaG: -152.06
    - !!omap
      - id: "s_0424"
      - name: "AMP"
      - compartment: "m"
      - formula: "C10H12N5O7P"
      - charge: -2
      - smiles: "C1=NC(=C2C(=N1)N(C=N2)C3C(C(C(O3)COP(=O)(O)O)O)O)N"
      - annotation: !!omap
          - bigg.metabolite: "amp"
          - chebi: "CHEBI:456215"
          - kegg.compound: "C00020"
          - metanetx.chemical: "MNXM14"
          - sbo: "SBO:0000247"
      - deltaG: -152.06
    - !!omap
      - id: "s_0425"
      - name: "AMP"
      - compartment: "n"
      - formula: "C10H12N5O7P"
      - charge: -2
      - smiles: "C1=NC(=C2C(=N1)N(C=N2)C3C(C(C(O3)COP(=O)(O)O)O)O)N"
      - annotation: !!omap
          - bigg.metabolite: "amp"
          - chebi: "CHEBI:456215"
          - kegg.compound: "C00020"
          - metanetx.chemical: "MNXM14"
          - sbo: "SBO:0000247"
      - deltaG: -152.06
    - !!omap
      - id: "s_0426"
      - name: "AMP"
      - compartment: "p"
      - formula: "C10H12N5O7P"
      - charge: -2
      - smiles: "C1=NC(=C2C(=N1)N(C=N2)C3C(C(C(O3)COP(=O)(O)O)O)O)N"
      - annotation: !!omap
          - bigg.metabolite: "amp"
          - chebi: "CHEBI:456215"
          - kegg.compound: "C00020"
          - metanetx.chemical: "MNXM14"
          - sbo: "SBO:0000247"
      - deltaG: -152.06
    - !!omap
      - id: "s_0427"
      - name: "anthranilate"
      - compartment: "c"
      - formula: "C7H6NO2"
      - charge: -1
      - smiles: "C1=CC=C(C(=C1)C(=O)O)N"
      - annotation: !!omap
          - bigg.metabolite: "anth"
          - chebi: "CHEBI:16567"
          - kegg.compound: "C00108"
          - metanetx.chemical: "MNXM188"
          - sbo: "SBO:0000247"
      - deltaG: 9.24
    - !!omap
      - id: "s_0428"
      - name: "Arg-tRNA(Arg)"
      - compartment: "c"
      - formula: "C6H15N4OR"
      - charge: 2
      - annotation: !!omap
          - bigg.metabolite: "argtrna"
          - chebi: "CHEBI:18366"
          - kegg.compound: "C02163"
          - metanetx.chemical: "MNXM89870"
          - sbo: "SBO:0000247"
      - deltaG: 10000000
    - !!omap
      - id: "s_0429"
      - name: "Arg-tRNA(Arg)"
      - compartment: "m"
      - formula: "C6H15N4OR"
      - charge: 2
      - annotation: !!omap
          - bigg.metabolite: "argtrna"
          - chebi: "CHEBI:18366"
          - kegg.compound: "C02163"
          - metanetx.chemical: "MNXM89870"
          - sbo: "SBO:0000247"
      - deltaG: 10000000
    - !!omap
      - id: "s_0430"
      - name: "Asn-tRNA(Asn)"
      - compartment: "c"
      - formula: "C4H8N2O2R"
      - charge: 1
      - annotation: !!omap
          - bigg.metabolite: "asntrna"
          - chebi: "CHEBI:29265"
          - kegg.compound: "C03402"
          - metanetx.chemical: "MNXM89761"
          - sbo: "SBO:0000247"
      - deltaG: 10000000
    - !!omap
      - id: "s_0431"
      - name: "Asn-tRNA(Asn)"
      - compartment: "m"
      - formula: "C4H8N2O2R"
      - charge: 1
      - annotation: !!omap
          - bigg.metabolite: "asntrna"
          - chebi: "CHEBI:29265"
          - kegg.compound: "C03402"
          - metanetx.chemical: "MNXM89761"
          - sbo: "SBO:0000247"
      - deltaG: 10000000
    - !!omap
      - id: "s_0432"
      - name: "Asp-tRNA(Asp)"
      - compartment: "c"
      - formula: "C4H6NO3R"
      - charge: 0
      - annotation: !!omap
          - bigg.metabolite: "asptrna"
          - chebi: "CHEBI:29158"
          - kegg.compound: "C02984"
          - metanetx.chemical: "MNXM164570"
          - sbo: "SBO:0000247"
      - deltaG: 10000000
    - !!omap
      - id: "s_0433"
      - name: "Asp-tRNA(Asp)"
      - compartment: "m"
      - formula: "C4H6NO3R"
      - charge: 0
      - annotation: !!omap
          - bigg.metabolite: "asptrna"
          - chebi: "CHEBI:29158"
          - kegg.compound: "C02984"
          - metanetx.chemical: "MNXM164570"
          - sbo: "SBO:0000247"
      - deltaG: 10000000
    - !!omap
      - id: "s_0434"
      - name: "ATP"
      - compartment: "c"
      - formula: "C10H12N5O13P3"
      - charge: -4
      - smiles: "C1=NC(=C2C(=N1)N(C=N2)C3C(C(C(O3)COP(=O)(O)OP(=O)(O)OP(=O)(O)O)O)O)N"
      - annotation: !!omap
          - bigg.metabolite: "atp"
          - chebi: "CHEBI:30616"
          - kegg.compound: "C00002"
          - metanetx.chemical: "MNXM3"
          - sbo: "SBO:0000247"
      - deltaG: -564.41
    - !!omap
      - id: "s_0435"
      - name: "ATP"
      - compartment: "er"
      - formula: "C10H12N5O13P3"
      - charge: -4
      - smiles: "C1=NC(=C2C(=N1)N(C=N2)C3C(C(C(O3)COP(=O)(O)OP(=O)(O)OP(=O)(O)O)O)O)N"
      - annotation: !!omap
          - bigg.metabolite: "atp"
          - chebi: "CHEBI:30616"
          - kegg.compound: "C00002"
          - metanetx.chemical: "MNXM3"
          - sbo: "SBO:0000247"
      - deltaG: -564.41
    - !!omap
      - id: "s_0437"
      - name: "ATP"
      - compartment: "m"
      - formula: "C10H12N5O13P3"
      - charge: -4
      - smiles: "C1=NC(=C2C(=N1)N(C=N2)C3C(C(C(O3)COP(=O)(O)OP(=O)(O)OP(=O)(O)O)O)O)N"
      - annotation: !!omap
          - bigg.metabolite: "atp"
          - chebi: "CHEBI:30616"
          - kegg.compound: "C00002"
          - metanetx.chemical: "MNXM3"
          - sbo: "SBO:0000247"
      - deltaG: -564.41
    - !!omap
      - id: "s_0438"
      - name: "ATP"
      - compartment: "n"
      - formula: "C10H12N5O13P3"
      - charge: -4
      - smiles: "C1=NC(=C2C(=N1)N(C=N2)C3C(C(C(O3)COP(=O)(O)OP(=O)(O)OP(=O)(O)O)O)O)N"
      - annotation: !!omap
          - bigg.metabolite: "atp"
          - chebi: "CHEBI:30616"
          - kegg.compound: "C00002"
          - metanetx.chemical: "MNXM3"
          - sbo: "SBO:0000247"
      - deltaG: -564.41
    - !!omap
      - id: "s_0439"
      - name: "ATP"
      - compartment: "p"
      - formula: "C10H12N5O13P3"
      - charge: -4
      - smiles: "C1=NC(=C2C(=N1)N(C=N2)C3C(C(C(O3)COP(=O)(O)OP(=O)(O)OP(=O)(O)O)O)O)N"
      - annotation: !!omap
          - bigg.metabolite: "atp"
          - chebi: "CHEBI:30616"
          - kegg.compound: "C00002"
          - metanetx.chemical: "MNXM3"
          - sbo: "SBO:0000247"
      - deltaG: -564.41
    - !!omap
      - id: "s_0441"
      - name: "beta-alanine"
      - compartment: "c"
      - formula: "C3H7NO2"
      - charge: 0
      - smiles: "C(CN)C(=O)O"
      - annotation: !!omap
          - bigg.metabolite: "ala_B"
          - chebi: "CHEBI:16958"
          - kegg.compound: "C00099"
          - metanetx.chemical: "MNXM144"
          - sbo: "SBO:0000247"
      - deltaG: -22.32
    - !!omap
      - id: "s_0442"
      - name: "beta-D-fructose 2,6-bisphosphate"
      - compartment: "c"
      - formula: "C6H10O12P2"
      - charge: -4
      - smiles: "C(C1C(C(C(O1)(CO)OP(=O)(O)O)O)O)OP(=O)(O)O"
      - annotation: !!omap
          - bigg.metabolite: "f26bp"
          - chebi: "CHEBI:58579"
          - kegg.compound: "C00665"
          - metanetx.chemical: "MNXM651"
          - sbo: "SBO:0000247"
      - deltaG: -549.91
    - !!omap
      - id: "s_0443"
      - name: "beta-D-mannosyldiacetylchitobiosyldiphosphodolichol"
      - compartment: "c"
      - formula: "C42H72N2O22P2"
      - charge: -2
      - smiles: "CC(CCC=C(C)CCC=C(C)CCC=C(C)CCC=C(C)C)CCOP(=O)(O)OP(=O)(O)OC1C(C(C(C(O1)CO)OC2C(C(C(C(O2)CO)OC3C(C(C(C(O3)CO)O)O)O)O)NC(=O)C)O)NC(=O)C"
      - annotation: !!omap
          - bigg.metabolite: "mpdol"
          - chebi: "CHEBI:18396"
          - kegg.compound: "C05860"
          - metanetx.chemical: "MNXM1078"
          - sbo: "SBO:0000247"
      - deltaG: -32.88
    - !!omap
      - id: "s_0444"
      - name: "beta-D-mannosyldiacetylchitobiosyldiphosphodolichol"
      - compartment: "g"
      - formula: "C42H72N2O22P2"
      - charge: -2
      - smiles: "CC(CCC=C(C)CCC=C(C)CCC=C(C)CCC=C(C)C)CCOP(=O)(O)OP(=O)(O)OC1C(C(C(C(O1)CO)OC2C(C(C(C(O2)CO)OC3C(C(C(C(O3)CO)O)O)O)O)NC(=O)C)O)NC(=O)C"
      - annotation: !!omap
          - bigg.metabolite: "mpdol"
          - chebi: "CHEBI:18396"
          - kegg.compound: "C05860"
          - metanetx.chemical: "MNXM1078"
          - sbo: "SBO:0000247"
      - deltaG: -32.88
    - !!omap
      - id: "s_0445"
      - name: "bicarbonate"
      - compartment: "c"
      - formula: "CHO3"
      - charge: -1
      - smiles: "C(=O)(O)[O-]"
      - annotation: !!omap
          - bigg.metabolite: "hco3"
          - chebi: "CHEBI:17544"
          - kegg.compound: "C00288"
          - metanetx.chemical: "MNXM60"
          - sbo: "SBO:0000247"
      - deltaG: -130.72
    - !!omap
      - id: "s_0446"
      - name: "bicarbonate"
      - compartment: "e"
      - formula: "CHO3"
      - charge: -1
      - smiles: "C(=O)(O)[O-]"
      - annotation: !!omap
          - bigg.metabolite: "hco3"
          - chebi: "CHEBI:17544"
          - kegg.compound: "C00288"
          - metanetx.chemical: "MNXM60"
          - sbo: "SBO:0000247"
      - deltaG: -130.72
    - !!omap
      - id: "s_0447"
      - name: "bicarbonate"
      - compartment: "m"
      - formula: "CHO3"
      - charge: -1
      - smiles: "C(=O)(O)[O-]"
      - annotation: !!omap
          - bigg.metabolite: "hco3"
          - chebi: "CHEBI:17544"
          - kegg.compound: "C00288"
          - metanetx.chemical: "MNXM60"
          - sbo: "SBO:0000247"
      - deltaG: -130.72
    - !!omap
      - id: "s_0448"
      - name: "bicarbonate"
      - compartment: "n"
      - formula: "CHO3"
      - charge: -1
      - smiles: "C(=O)(O)[O-]"
      - annotation: !!omap
          - bigg.metabolite: "hco3"
          - chebi: "CHEBI:17544"
          - kegg.compound: "C00288"
          - metanetx.chemical: "MNXM60"
          - sbo: "SBO:0000247"
      - deltaG: -130.72
    - !!omap
      - id: "s_0450"
      - name: "biomass"
      - compartment: "c"
      - charge: 0
      - annotation: !!omap
          - bigg.metabolite: "biomass"
          - sbo: "SBO:0000649"
      - deltaG: 10000000
    - !!omap
      - id: "s_0451"
      - name: "biotin"
      - compartment: "c"
      - formula: "C10H15N2O3S"
      - charge: -1
      - smiles: "C1C2C(C(S1)CCCCC(=O)O)NC(=O)N2"
      - annotation: !!omap
          - bigg.metabolite: "btn"
          - chebi: "CHEBI:57586"
          - kegg.compound: "C00120"
          - metanetx.chemical: "MNXM304"
          - sbo: "SBO:0000247"
      - deltaG: 36.84
    - !!omap
      - id: "s_0452"
      - name: "biotin"
      - compartment: "e"
      - formula: "C10H15N2O3S"
      - charge: -1
      - smiles: "C1C2C(C(S1)CCCCC(=O)O)NC(=O)N2"
      - annotation: !!omap
          - bigg.metabolite: "btn"
          - chebi: "CHEBI:57586"
          - kegg.compound: "C00120"
          - metanetx.chemical: "MNXM304"
          - sbo: "SBO:0000247"
      - deltaG: 36.84
    - !!omap
      - id: "s_0453"
      - name: "biotinyl-5'-AMP"
      - compartment: "c"
      - formula: "C20H27N7O9PS"
      - charge: -1
      - smiles: "C1C2C(C(S1)CCCCC(=O)OP(=O)(O)OCC3C(C(C(O3)N4C=NC5=C(N=CN=C54)N)O)O)NC(=O)N2"
      - annotation: !!omap
          - bigg.metabolite: "btamp"
          - chebi: "CHEBI:62414"
          - kegg.compound: "C05921"
          - metanetx.chemical: "MNXM2351"
          - sbo: "SBO:0000247"
      - deltaG: -59.24
    - !!omap
      - id: "s_0454"
      - name: "but-1-ene-1,2,4-tricarboxylic acid"
      - compartment: "m"
      - formula: "C7H5O6"
      - charge: -3
      - smiles: "C(CC(=O)O)C(=CC(=O)O)C(=O)O"
      - annotation: !!omap
          - bigg.metabolite: "b124tc"
          - chebi: "CHEBI:58174"
          - kegg.compound: "C04002"
          - metanetx.chemical: "MNXM920"
          - sbo: "SBO:0000247"
      - deltaG: -170.31
    - !!omap
      - id: "s_0455"
      - name: "carbamoyl phosphate"
      - compartment: "c"
      - formula: "CH2NO5P"
      - charge: -2
      - smiles: "C(=O)(N)OP(=O)(O)O"
      - annotation: !!omap
          - bigg.metabolite: "cbp"
          - chebi: "CHEBI:58228"
          - kegg.compound: "C00169"
          - metanetx.chemical: "MNXM138"
          - sbo: "SBO:0000247"
      - deltaG: -291.58
    - !!omap
      - id: "s_0456"
      - name: "carbon dioxide"
      - compartment: "c"
      - formula: "CO2"
      - charge: 0
      - smiles: "C(=O)=O"
      - annotation: !!omap
          - bigg.metabolite: "co2"
          - chebi: "CHEBI:16526"
          - kegg.compound: "C00011"
          - metanetx.chemical: "MNXM13"
          - sbo: "SBO:0000247"
      - deltaG: -92.26
    - !!omap
      - id: "s_0457"
      - name: "carbon dioxide"
      - compartment: "er"
      - formula: "CO2"
      - charge: 0
      - smiles: "C(=O)=O"
      - annotation: !!omap
          - bigg.metabolite: "co2"
          - chebi: "CHEBI:16526"
          - kegg.compound: "C00011"
          - metanetx.chemical: "MNXM13"
          - sbo: "SBO:0000247"
      - deltaG: -92.26
    - !!omap
      - id: "s_0458"
      - name: "carbon dioxide"
      - compartment: "e"
      - formula: "CO2"
      - charge: 0
      - smiles: "C(=O)=O"
      - annotation: !!omap
          - bigg.metabolite: "co2"
          - chebi: "CHEBI:16526"
          - kegg.compound: "C00011"
          - metanetx.chemical: "MNXM13"
          - sbo: "SBO:0000247"
      - deltaG: -92.26
    - !!omap
      - id: "s_0460"
      - name: "carbon dioxide"
      - compartment: "m"
      - formula: "CO2"
      - charge: 0
      - smiles: "C(=O)=O"
      - annotation: !!omap
          - bigg.metabolite: "co2"
          - chebi: "CHEBI:16526"
          - kegg.compound: "C00011"
          - metanetx.chemical: "MNXM13"
          - sbo: "SBO:0000247"
      - deltaG: -92.26
    - !!omap
      - id: "s_0461"
      - name: "carbon dioxide"
      - compartment: "n"
      - formula: "CO2"
      - charge: 0
      - smiles: "C(=O)=O"
      - annotation: !!omap
          - bigg.metabolite: "co2"
          - chebi: "CHEBI:16526"
          - kegg.compound: "C00011"
          - metanetx.chemical: "MNXM13"
          - sbo: "SBO:0000247"
      - deltaG: -92.26
    - !!omap
      - id: "s_0462"
      - name: "carbon dioxide"
      - compartment: "p"
      - formula: "CO2"
      - charge: 0
      - smiles: "C(=O)=O"
      - annotation: !!omap
          - bigg.metabolite: "co2"
          - chebi: "CHEBI:16526"
          - kegg.compound: "C00011"
          - metanetx.chemical: "MNXM13"
          - sbo: "SBO:0000247"
      - deltaG: -92.26
    - !!omap
      - id: "s_0465"
      - name: "carboxyacetyl-ACP"
      - compartment: "m"
      - formula: "C3H3O3SR"
      - charge: 0
      - annotation: !!omap
          - bigg.metabolite: "malACP"
          - chebi: "CHEBI:17330"
          - kegg.compound: "C01209"
          - metanetx.chemical: "MNXM184"
          - sbo: "SBO:0000247"
      - deltaG: 10000000
    - !!omap
      - id: "s_0467"
      - name: "CDP"
      - compartment: "c"
      - formula: "C9H12N3O11P2"
      - charge: -3
      - smiles: "C1=CN(C(=O)N=C1N)C2C(C(C(O2)COP(=O)(O)OP(=O)(O)O)O)O"
      - annotation: !!omap
          - bigg.metabolite: "cdp"
          - chebi: "CHEBI:58069"
          - kegg.compound: "C00112"
          - metanetx.chemical: "MNXM220"
          - sbo: "SBO:0000247"
      - deltaG: -428.35
    - !!omap
      - id: "s_0468"
      - name: "CDP"
      - compartment: "n"
      - formula: "C9H12N3O11P2"
      - charge: -3
      - smiles: "C1=CN(C(=O)N=C1N)C2C(C(C(O2)COP(=O)(O)OP(=O)(O)O)O)O"
      - annotation: !!omap
          - bigg.metabolite: "cdp"
          - chebi: "CHEBI:58069"
          - kegg.compound: "C00112"
          - metanetx.chemical: "MNXM220"
          - sbo: "SBO:0000247"
      - deltaG: -428.35
    - !!omap
      - id: "s_0469"
      - name: "CDP-choline"
      - compartment: "c"
      - formula: "C14H25N4O11P2"
      - charge: -1
      - smiles: "C[N+](C)(C)CCOP(=O)([O-])OP(=O)(O)OCC1C(C(C(O1)N2C=CC(=NC2=O)N)O)O"
      - annotation: !!omap
          - bigg.metabolite: "cdpchol"
          - chebi: "CHEBI:16436"
          - kegg.compound: "C00307"
          - metanetx.chemical: "MNXM283"
          - sbo: "SBO:0000247"
      - deltaG: -242.43
    - !!omap
      - id: "s_0474"
      - name: "CDP-ethanolamine"
      - compartment: "c"
      - formula: "C11H19N4O11P2"
      - charge: -1
      - smiles: "C1=CN(C(=O)N=C1N)C2C(C(C(O2)COP(=O)(O)OP(=O)(O)OCCN)O)O"
      - annotation: !!omap
          - bigg.metabolite: "cdpea"
          - chebi: "CHEBI:57876"
          - kegg.compound: "C00570"
          - metanetx.chemical: "MNXM449"
          - sbo: "SBO:0000247"
      - deltaG: -357.55
    - !!omap
      - id: "s_0475"
      - name: "ceramide-1 (C24)"
      - compartment: "er"
      - formula: "C42H85NO3"
      - charge: 0
      - annotation: !!omap
          - bigg.metabolite: "cer1_24"
          - chebi: "CHEBI:52961"
          - metanetx.chemical: "MNXM64020"
          - sbo: "SBO:0000247"
      - deltaG: 781.96
    - !!omap
      - id: "s_0476"
      - name: "ceramide-1 (C24)"
      - compartment: "g"
      - formula: "C42H85NO3"
      - charge: 0
      - annotation: !!omap
          - bigg.metabolite: "cer1_24"
          - chebi: "CHEBI:52961"
          - metanetx.chemical: "MNXM64020"
          - sbo: "SBO:0000247"
      - deltaG: 781.96
    - !!omap
      - id: "s_0477"
      - name: "ceramide-1 (C24)"
      - compartment: "m"
      - formula: "C42H85NO3"
      - charge: 0
      - annotation: !!omap
          - bigg.metabolite: "cer1_24"
          - chebi: "CHEBI:52961"
          - metanetx.chemical: "MNXM64020"
          - sbo: "SBO:0000247"
      - deltaG: 781.96
    - !!omap
      - id: "s_0478"
      - name: "ceramide-1 (C26)"
      - compartment: "er"
      - formula: "C44H89NO3"
      - charge: 0
      - smiles: "CCCCCCCCCCCCCCCCCCCCCCCCCC(=O)NC(CO)C(CCCCCCCCCCCCCCC)O"
      - annotation: !!omap
          - bigg.metabolite: "cer1_26"
          - chebi: "CHEBI:52962"
          - metanetx.chemical: "MNXM63798"
          - sbo: "SBO:0000247"
      - deltaG: 824.47
    - !!omap
      - id: "s_0479"
      - name: "ceramide-1 (C26)"
      - compartment: "g"
      - formula: "C44H89NO3"
      - charge: 0
      - smiles: "CCCCCCCCCCCCCCCCCCCCCCCCCC(=O)NC(CO)C(CCCCCCCCCCCCCCC)O"
      - annotation: !!omap
          - bigg.metabolite: "cer1_26"
          - chebi: "CHEBI:52962"
          - metanetx.chemical: "MNXM63798"
          - sbo: "SBO:0000247"
      - deltaG: 824.47
    - !!omap
      - id: "s_0480"
      - name: "ceramide-1 (C26)"
      - compartment: "m"
      - formula: "C44H89NO3"
      - charge: 0
      - smiles: "CCCCCCCCCCCCCCCCCCCCCCCCCC(=O)NC(CO)C(CCCCCCCCCCCCCCC)O"
      - annotation: !!omap
          - bigg.metabolite: "cer1_26"
          - chebi: "CHEBI:52962"
          - metanetx.chemical: "MNXM63798"
          - sbo: "SBO:0000247"
      - deltaG: 824.47
    - !!omap
      - id: "s_0481"
      - name: "ceramide-2 (C24)"
      - compartment: "er"
      - formula: "C42H85NO4"
      - charge: 0
      - smiles: "CCCCCCCCCCCCCCCCCCCCCCCC(=O)NC(CO)C(C(CCCCCCCCCCCCCC)O)O"
      - annotation: !!omap
          - bigg.metabolite: "cer2_24"
          - chebi: "CHEBI:52979"
          - metanetx.chemical: "MNXM64019"
          - sbo: "SBO:0000247"
      - deltaG: 743.92
    - !!omap
      - id: "s_0482"
      - name: "ceramide-2 (C24)"
      - compartment: "g"
      - formula: "C42H85NO4"
      - charge: 0
      - smiles: "CCCCCCCCCCCCCCCCCCCCCCCC(=O)NC(CO)C(C(CCCCCCCCCCCCCC)O)O"
      - annotation: !!omap
          - bigg.metabolite: "cer2_24"
          - chebi: "CHEBI:52979"
          - metanetx.chemical: "MNXM64019"
          - sbo: "SBO:0000247"
      - deltaG: 743.92
    - !!omap
      - id: "s_0483"
      - name: "ceramide-2 (C24)"
      - compartment: "m"
      - formula: "C42H85NO4"
      - charge: 0
      - smiles: "CCCCCCCCCCCCCCCCCCCCCCCC(=O)NC(CO)C(C(CCCCCCCCCCCCCC)O)O"
      - annotation: !!omap
          - bigg.metabolite: "cer2_24"
          - chebi: "CHEBI:52979"
          - metanetx.chemical: "MNXM64019"
          - sbo: "SBO:0000247"
      - deltaG: 743.92
    - !!omap
      - id: "s_0484"
      - name: "ceramide-2 (C26)"
      - compartment: "er"
      - formula: "C44H89NO4"
      - charge: 0
      - smiles: "CCCCCCCCCCCCCCCCCCCCCCCCCC(=O)NC(CO)C(C(CCCCCCCCCCCCCC)O)O"
      - annotation: !!omap
          - bigg.metabolite: "cer2_26"
          - chebi: "CHEBI:52980"
          - metanetx.chemical: "MNXM63797"
          - sbo: "SBO:0000247"
      - deltaG: 786.43
    - !!omap
      - id: "s_0485"
      - name: "ceramide-2 (C26)"
      - compartment: "g"
      - formula: "C44H89NO4"
      - charge: 0
      - smiles: "CCCCCCCCCCCCCCCCCCCCCCCCCC(=O)NC(CO)C(C(CCCCCCCCCCCCCC)O)O"
      - annotation: !!omap
          - bigg.metabolite: "cer2_26"
          - chebi: "CHEBI:52980"
          - metanetx.chemical: "MNXM63797"
          - sbo: "SBO:0000247"
      - deltaG: 786.43
    - !!omap
      - id: "s_0486"
      - name: "ceramide-2 (C26)"
      - compartment: "m"
      - formula: "C44H89NO4"
      - charge: 0
      - smiles: "CCCCCCCCCCCCCCCCCCCCCCCCCC(=O)NC(CO)C(C(CCCCCCCCCCCCCC)O)O"
      - annotation: !!omap
          - bigg.metabolite: "cer2_26"
          - chebi: "CHEBI:52980"
          - metanetx.chemical: "MNXM63797"
          - sbo: "SBO:0000247"
      - deltaG: 786.43
    - !!omap
      - id: "s_0487"
      - name: "ceramide-2' (C24)"
      - compartment: "er"
      - formula: "C42H85NO4"
      - charge: 0
      - smiles: "CCCCCCCCCCCCCCCCCCCCCCC(C(=O)NC(CO)C(CCCCCCCCCCCCCCC)O)O"
      - annotation: !!omap
          - chebi: "CHEBI:52371"
          - metanetx.chemical: "MNXM63175"
          - sbo: "SBO:0000247"
      - deltaG: 743.92
    - !!omap
      - id: "s_0488"
      - name: "ceramide-2' (C24)"
      - compartment: "g"
      - formula: "C42H85NO4"
      - charge: 0
      - smiles: "CCCCCCCCCCCCCCCCCCCCCCC(C(=O)NC(CO)C(CCCCCCCCCCCCCCC)O)O"
      - annotation: !!omap
          - chebi: "CHEBI:52371"
          - metanetx.chemical: "MNXM63175"
          - sbo: "SBO:0000247"
      - deltaG: 743.92
    - !!omap
      - id: "s_0489"
      - name: "ceramide-2' (C24)"
      - compartment: "m"
      - formula: "C42H85NO4"
      - charge: 0
      - smiles: "CCCCCCCCCCCCCCCCCCCCCCC(C(=O)NC(CO)C(CCCCCCCCCCCCCCC)O)O"
      - annotation: !!omap
          - chebi: "CHEBI:52371"
          - metanetx.chemical: "MNXM63175"
          - sbo: "SBO:0000247"
      - deltaG: 743.92
    - !!omap
      - id: "s_0490"
      - name: "ceramide-2' (C26)"
      - compartment: "er"
      - formula: "C44H89NO4"
      - charge: 0
      - annotation: !!omap
          - chebi: "CHEBI:52372"
          - metanetx.chemical: "MNXM63156"
          - sbo: "SBO:0000247"
      - deltaG: 786.43
    - !!omap
      - id: "s_0491"
      - name: "ceramide-2' (C26)"
      - compartment: "g"
      - formula: "C44H89NO4"
      - charge: 0
      - annotation: !!omap
          - chebi: "CHEBI:52372"
          - metanetx.chemical: "MNXM63156"
          - sbo: "SBO:0000247"
      - deltaG: 786.43
    - !!omap
      - id: "s_0492"
      - name: "ceramide-2' (C26)"
      - compartment: "m"
      - formula: "C44H89NO4"
      - charge: 0
      - annotation: !!omap
          - chebi: "CHEBI:52372"
          - metanetx.chemical: "MNXM63156"
          - sbo: "SBO:0000247"
      - deltaG: 786.43
    - !!omap
      - id: "s_0493"
      - name: "ceramide-3 (C24)"
      - compartment: "er"
      - formula: "C42H85NO5"
      - charge: 0
      - smiles: "CCCCCCCCCCCCCCCCCCCCCCC(C(=O)NC(CO)C(C(CCCCCCCCCCCCCC)O)O)O"
      - annotation: !!omap
          - bigg.metabolite: "cer3_24"
          - chebi: "CHEBI:52373"
          - metanetx.chemical: "MNXM63174"
          - sbo: "SBO:0000247"
      - deltaG: 705.88
    - !!omap
      - id: "s_0494"
      - name: "ceramide-3 (C24)"
      - compartment: "g"
      - formula: "C42H85NO5"
      - charge: 0
      - smiles: "CCCCCCCCCCCCCCCCCCCCCCC(C(=O)NC(CO)C(C(CCCCCCCCCCCCCC)O)O)O"
      - annotation: !!omap
          - bigg.metabolite: "cer3_24"
          - chebi: "CHEBI:52373"
          - metanetx.chemical: "MNXM63174"
          - sbo: "SBO:0000247"
      - deltaG: 705.88
    - !!omap
      - id: "s_0495"
      - name: "ceramide-3 (C24)"
      - compartment: "m"
      - formula: "C42H85NO5"
      - charge: 0
      - smiles: "CCCCCCCCCCCCCCCCCCCCCCC(C(=O)NC(CO)C(C(CCCCCCCCCCCCCC)O)O)O"
      - annotation: !!omap
          - bigg.metabolite: "cer3_24"
          - chebi: "CHEBI:52373"
          - metanetx.chemical: "MNXM63174"
          - sbo: "SBO:0000247"
      - deltaG: 705.88
    - !!omap
      - id: "s_0496"
      - name: "ceramide-3 (C26)"
      - compartment: "er"
      - formula: "C44H89NO5"
      - charge: 0
      - smiles: "CCCCCCCCCCCCCCCCCCCCCCCCC(C(=O)NC(CO)C(C(CCCCCCCCCCCCCC)O)O)O"
      - annotation: !!omap
          - bigg.metabolite: "cer3_26"
          - chebi: "CHEBI:52374"
          - metanetx.chemical: "MNXM63157"
          - sbo: "SBO:0000247"
      - deltaG: 748.39
    - !!omap
      - id: "s_0497"
      - name: "ceramide-3 (C26)"
      - compartment: "g"
      - formula: "C44H89NO5"
      - charge: 0
      - smiles: "CCCCCCCCCCCCCCCCCCCCCCCCC(C(=O)NC(CO)C(C(CCCCCCCCCCCCCC)O)O)O"
      - annotation: !!omap
          - bigg.metabolite: "cer3_26"
          - chebi: "CHEBI:52374"
          - metanetx.chemical: "MNXM63157"
          - sbo: "SBO:0000247"
      - deltaG: 748.39
    - !!omap
      - id: "s_0498"
      - name: "ceramide-3 (C26)"
      - compartment: "m"
      - formula: "C44H89NO5"
      - charge: 0
      - smiles: "CCCCCCCCCCCCCCCCCCCCCCCCC(C(=O)NC(CO)C(C(CCCCCCCCCCCCCC)O)O)O"
      - annotation: !!omap
          - bigg.metabolite: "cer3_26"
          - chebi: "CHEBI:52374"
          - metanetx.chemical: "MNXM63157"
          - sbo: "SBO:0000247"
      - deltaG: 748.39
    - !!omap
      - id: "s_0499"
      - name: "ceramide-4 (C24)"
      - compartment: "er"
      - formula: "C42H85NO6"
      - charge: 0
      - smiles: "CCCCCCCCCCCCCCCCCCCCCC(C(C(=O)NC(CO)C(C(CCCCCCCCCCCCCC)O)O)O)O"
      - annotation: !!omap
          - chebi: "CHEBI:60256"
          - metanetx.chemical: "MNXM31230"
          - sbo: "SBO:0000247"
      - deltaG: 667.84
    - !!omap
      - id: "s_0500"
      - name: "ceramide-4 (C24)"
      - compartment: "g"
      - formula: "C42H85NO6"
      - charge: 0
      - smiles: "CCCCCCCCCCCCCCCCCCCCCC(C(C(=O)NC(CO)C(C(CCCCCCCCCCCCCC)O)O)O)O"
      - annotation: !!omap
          - chebi: "CHEBI:60256"
          - metanetx.chemical: "MNXM31230"
          - sbo: "SBO:0000247"
      - deltaG: 667.84
    - !!omap
      - id: "s_0501"
      - name: "ceramide-4 (C24)"
      - compartment: "m"
      - formula: "C42H85NO6"
      - charge: 0
      - smiles: "CCCCCCCCCCCCCCCCCCCCCC(C(C(=O)NC(CO)C(C(CCCCCCCCCCCCCC)O)O)O)O"
      - annotation: !!omap
          - chebi: "CHEBI:60256"
          - metanetx.chemical: "MNXM31230"
          - sbo: "SBO:0000247"
      - deltaG: 667.84
    - !!omap
      - id: "s_0502"
      - name: "ceramide-4 (C26)"
      - compartment: "er"
      - formula: "C44H89NO6"
      - charge: 0
      - smiles: "CCCCCCCCCCCCCCCCCCCCCCCC(C(C(=O)NC(CO)C(C(CCCCCCCCCCCCCC)O)O)O)O"
      - annotation: !!omap
          - chebi: "CHEBI:60384"
          - metanetx.chemical: "MNXM63127"
          - sbo: "SBO:0000247"
      - deltaG: 710.35
    - !!omap
      - id: "s_0503"
      - name: "ceramide-4 (C26)"
      - compartment: "g"
      - formula: "C44H89NO6"
      - charge: 0
      - smiles: "CCCCCCCCCCCCCCCCCCCCCCCC(C(C(=O)NC(CO)C(C(CCCCCCCCCCCCCC)O)O)O)O"
      - annotation: !!omap
          - chebi: "CHEBI:60384"
          - metanetx.chemical: "MNXM63127"
          - sbo: "SBO:0000247"
      - deltaG: 710.35
    - !!omap
      - id: "s_0504"
      - name: "ceramide-4 (C26)"
      - compartment: "m"
      - formula: "C44H89NO6"
      - charge: 0
      - smiles: "CCCCCCCCCCCCCCCCCCCCCCCC(C(C(=O)NC(CO)C(C(CCCCCCCCCCCCCC)O)O)O)O"
      - annotation: !!omap
          - chebi: "CHEBI:60384"
          - metanetx.chemical: "MNXM63127"
          - sbo: "SBO:0000247"
      - deltaG: 710.35
    - !!omap
      - id: "s_0507"
      - name: "cerotic acid"
      - compartment: "er"
      - formula: "C26H51O2"
      - charge: -1
      - smiles: "CCCCCCCCCCCCCCCCCCCCCCCCCC(=O)O"
      - annotation: !!omap
          - bigg.metabolite: "hexc"
          - chebi: "CHEBI:31013"
          - metanetx.chemical: "MNXM46158"
          - sbo: "SBO:0000247"
      - deltaG: 452.74
    - !!omap
      - id: "s_0509"
      - name: "chitin"
      - compartment: "c"
      - formula: "C8H13NO5"
      - charge: 0
      - annotation: !!omap
          - bigg.metabolite: "chtn"
          - chebi: "CHEBI:17029"
          - kegg.compound: "C00461"
          - metanetx.chemical: "MNXM1271"
          - sbo: "SBO:0000247"
      - deltaG: 10000000
    - !!omap
      - id: "s_0510"
      - name: "chitosan"
      - compartment: "ce"
      - formula: "C6H11NO4"
      - charge: 0
      - smiles: "COC(=O)NC1C(C(C(OC1OC2C(OC(C(C2O)N)OC3C(OC(C(C3O)N)O)CO)CO)CO)OC4C(C(C(C(O4)CO)OC5C(C(C(C(O5)CO)OC6C(C(C(C(O6)CO)OC7C(C(C(C(O7)CO)OC8C(C(C(C(O8)CO)OC9C(C(C(C(O9)CO)O)O)N)O)N)O)N)O)N)O)N)O)N)O"
      - annotation: !!omap
          - bigg.metabolite: "chitos"
          - chebi: "CHEBI:16261"
          - kegg.compound: "C00734"
          - metanetx.chemical: "MNXM2106"
          - sbo: "SBO:0000247"
      - deltaG: -66.84
    - !!omap
      - id: "s_0511"
      - name: "choline"
      - compartment: "ce"
      - formula: "C5H14NO"
      - charge: 1
      - smiles: "C[N+](C)(C)CCO"
      - annotation: !!omap
          - bigg.metabolite: "chol"
          - chebi: "CHEBI:15354"
          - kegg.compound: "C00114"
          - metanetx.chemical: "MNXM90"
          - sbo: "SBO:0000247"
      - deltaG: 145.8
    - !!omap
      - id: "s_0512"
      - name: "choline"
      - compartment: "c"
      - formula: "C5H14NO"
      - charge: 1
      - smiles: "C[N+](C)(C)CCO"
      - annotation: !!omap
          - bigg.metabolite: "chol"
          - chebi: "CHEBI:15354"
          - kegg.compound: "C00114"
          - metanetx.chemical: "MNXM90"
          - sbo: "SBO:0000247"
      - deltaG: 145.8
    - !!omap
      - id: "s_0513"
      - name: "choline"
      - compartment: "e"
      - formula: "C5H14NO"
      - charge: 1
      - smiles: "C[N+](C)(C)CCO"
      - annotation: !!omap
          - bigg.metabolite: "chol"
          - chebi: "CHEBI:15354"
          - kegg.compound: "C00114"
          - metanetx.chemical: "MNXM90"
          - sbo: "SBO:0000247"
      - deltaG: 145.8
    - !!omap
      - id: "s_0514"
      - name: "choline phosphate"
      - compartment: "c"
      - formula: "C5H13NO4P"
      - charge: -1
      - smiles: "C[N+](C)(C)CCOP(=O)(O)O.[Cl-]"
      - annotation: !!omap
          - bigg.metabolite: "cholp"
          - chebi: "CHEBI:295975"
          - kegg.compound: "C00588"
          - metanetx.chemical: "MNXM229"
          - sbo: "SBO:0000247"
      - deltaG: -75.84
    - !!omap
      - id: "s_0515"
      - name: "chorismate"
      - compartment: "c"
      - formula: "C10H8O6"
      - charge: -2
      - smiles: "C=C(C(=O)O)OC1C=C(C=CC1O)C(=O)O"
      - annotation: !!omap
          - bigg.metabolite: "chor"
          - chebi: "CHEBI:29748"
          - kegg.compound: "C00251"
          - metanetx.chemical: "MNXM337"
          - sbo: "SBO:0000247"
      - deltaG: -92.37
    - !!omap
      - id: "s_0516"
      - name: "cis-aconitate"
      - compartment: "c"
      - formula: "C6H3O6"
      - charge: -3
      - smiles: "C(C(=CC(=O)O)C(=O)O)C(=O)O"
      - annotation: !!omap
          - bigg.metabolite: "acon_C"
          - chebi: "CHEBI:16383"
          - kegg.compound: "C00417"
          - metanetx.chemical: "MNXM813"
          - sbo: "SBO:0000247"
      - deltaG: -194.81
    - !!omap
      - id: "s_0517"
      - name: "cis-aconitate"
      - compartment: "m"
      - formula: "C6H3O6"
      - charge: -3
      - smiles: "C(C(=CC(=O)O)C(=O)O)C(=O)O"
      - annotation: !!omap
          - bigg.metabolite: "acon_C"
          - chebi: "CHEBI:16383"
          - kegg.compound: "C00417"
          - metanetx.chemical: "MNXM813"
          - sbo: "SBO:0000247"
      - deltaG: -194.81
    - !!omap
      - id: "s_0522"
      - name: "citrate"
      - compartment: "c"
      - formula: "C6H5O7"
      - charge: -3
      - smiles: "C(C(=O)[O-])C(CC(=O)[O-])(C(=O)[O-])O"
      - annotation: !!omap
          - bigg.metabolite: "cit"
          - chebi: "CHEBI:16947"
          - kegg.compound: "C00158"
          - metanetx.chemical: "MNXM131"
          - sbo: "SBO:0000247"
      - deltaG: -233.5
    - !!omap
      - id: "s_0523"
      - name: "citrate"
      - compartment: "e"
      - formula: "C6H5O7"
      - charge: -3
      - smiles: "C(C(=O)[O-])C(CC(=O)[O-])(C(=O)[O-])O"
      - annotation: !!omap
          - bigg.metabolite: "cit"
          - chebi: "CHEBI:16947"
          - kegg.compound: "C00158"
          - metanetx.chemical: "MNXM131"
          - sbo: "SBO:0000247"
      - deltaG: -233.5
    - !!omap
      - id: "s_0524"
      - name: "citrate"
      - compartment: "m"
      - formula: "C6H5O7"
      - charge: -3
      - smiles: "C(C(=O)[O-])C(CC(=O)[O-])(C(=O)[O-])O"
      - annotation: !!omap
          - bigg.metabolite: "cit"
          - chebi: "CHEBI:16947"
          - kegg.compound: "C00158"
          - metanetx.chemical: "MNXM131"
          - sbo: "SBO:0000247"
      - deltaG: -233.5
    - !!omap
      - id: "s_0525"
      - name: "citrate"
      - compartment: "p"
      - formula: "C6H5O7"
      - charge: -3
      - smiles: "C(C(=O)[O-])C(CC(=O)[O-])(C(=O)[O-])O"
      - annotation: !!omap
          - bigg.metabolite: "cit"
          - chebi: "CHEBI:16947"
          - kegg.compound: "C00158"
          - metanetx.chemical: "MNXM131"
          - sbo: "SBO:0000247"
      - deltaG: -233.5
    - !!omap
      - id: "s_0526"
      - name: "CMP"
      - compartment: "c"
      - formula: "C9H12N3O8P"
      - charge: -2
      - smiles: "C1=CN(C(=O)N=C1N)C2C(C(C(O2)COP(=O)(O)O)O)O"
      - annotation: !!omap
          - bigg.metabolite: "cmp"
          - chebi: "CHEBI:60377"
          - kegg.compound: "C00055"
          - metanetx.chemical: "MNXM31"
          - sbo: "SBO:0000247"
      - deltaG: -229.34
    - !!omap
      - id: "s_0528"
      - name: "CMP"
      - compartment: "m"
      - formula: "C9H12N3O8P"
      - charge: -2
      - smiles: "C1=CN(C(=O)N=C1N)C2C(C(C(O2)COP(=O)(O)O)O)O"
      - annotation: !!omap
          - bigg.metabolite: "cmp"
          - chebi: "CHEBI:60377"
          - kegg.compound: "C00055"
          - metanetx.chemical: "MNXM31"
          - sbo: "SBO:0000247"
      - deltaG: -229.34
    - !!omap
      - id: "s_0529"
      - name: "coenzyme A"
      - compartment: "c"
      - formula: "C21H32N7O16P3S"
      - charge: -4
      - smiles: "CC(C)(COP(=O)(O)OP(=O)(O)OCC1C(C(C(O1)N2C=NC3=C(N=CN=C32)N)O)OP(=O)(O)O)C(C(=O)NCCC(=O)NCCS)O"
      - annotation: !!omap
          - bigg.metabolite: "coa"
          - chebi: "CHEBI:57287"
          - kegg.compound: "C00010"
          - metanetx.chemical: "MNXM12"
          - sbo: "SBO:0000247"
      - deltaG: -454.22
    - !!omap
      - id: "s_0530"
      - name: "coenzyme A"
      - compartment: "er"
      - formula: "C21H32N7O16P3S"
      - charge: -4
      - smiles: "CC(C)(COP(=O)(O)OP(=O)(O)OCC1C(C(C(O1)N2C=NC3=C(N=CN=C32)N)O)OP(=O)(O)O)C(C(=O)NCCC(=O)NCCS)O"
      - annotation: !!omap
          - bigg.metabolite: "coa"
          - chebi: "CHEBI:57287"
          - kegg.compound: "C00010"
          - metanetx.chemical: "MNXM12"
          - sbo: "SBO:0000247"
      - deltaG: -454.22
    - !!omap
      - id: "s_0531"
      - name: "coenzyme A"
      - compartment: "lp"
      - formula: "C21H32N7O16P3S"
      - charge: -4
      - smiles: "CC(C)(COP(=O)(O)OP(=O)(O)OCC1C(C(C(O1)N2C=NC3=C(N=CN=C32)N)O)OP(=O)(O)O)C(C(=O)NCCC(=O)NCCS)O"
      - annotation: !!omap
          - bigg.metabolite: "coa"
          - chebi: "CHEBI:57287"
          - kegg.compound: "C00010"
          - metanetx.chemical: "MNXM12"
          - sbo: "SBO:0000247"
      - deltaG: -454.22
    - !!omap
      - id: "s_0532"
      - name: "coenzyme A"
      - compartment: "m"
      - formula: "C21H32N7O16P3S"
      - charge: -4
      - smiles: "CC(C)(COP(=O)(O)OP(=O)(O)OCC1C(C(C(O1)N2C=NC3=C(N=CN=C32)N)O)OP(=O)(O)O)C(C(=O)NCCC(=O)NCCS)O"
      - annotation: !!omap
          - bigg.metabolite: "coa"
          - chebi: "CHEBI:57287"
          - kegg.compound: "C00010"
          - metanetx.chemical: "MNXM12"
          - sbo: "SBO:0000247"
      - deltaG: -454.22
    - !!omap
      - id: "s_0533"
      - name: "coenzyme A"
      - compartment: "n"
      - formula: "C21H32N7O16P3S"
      - charge: -4
      - smiles: "CC(C)(COP(=O)(O)OP(=O)(O)OCC1C(C(C(O1)N2C=NC3=C(N=CN=C32)N)O)OP(=O)(O)O)C(C(=O)NCCC(=O)NCCS)O"
      - annotation: !!omap
          - bigg.metabolite: "coa"
          - chebi: "CHEBI:57287"
          - kegg.compound: "C00010"
          - metanetx.chemical: "MNXM12"
          - sbo: "SBO:0000247"
      - deltaG: -454.22
    - !!omap
      - id: "s_0534"
      - name: "coenzyme A"
      - compartment: "p"
      - formula: "C21H32N7O16P3S"
      - charge: -4
      - smiles: "CC(C)(COP(=O)(O)OP(=O)(O)OCC1C(C(C(O1)N2C=NC3=C(N=CN=C32)N)O)OP(=O)(O)O)C(C(=O)NCCC(=O)NCCS)O"
      - annotation: !!omap
          - bigg.metabolite: "coa"
          - chebi: "CHEBI:57287"
          - kegg.compound: "C00010"
          - metanetx.chemical: "MNXM12"
          - sbo: "SBO:0000247"
      - deltaG: -454.22
    - !!omap
      - id: "s_0538"
      - name: "coproporphyrinogen III"
      - compartment: "c"
      - formula: "C36H40N4O8"
      - charge: -4
      - smiles: "CC1=C2CC3=C(C(=C(N3)CC4=C(C(=C(N4)CC5=C(C(=C(N5)CC(=C1CCC(=O)O)N2)C)CCC(=O)O)C)CCC(=O)O)CCC(=O)O)C"
      - annotation: !!omap
          - bigg.metabolite: "cpppg3"
          - chebi: "CHEBI:57309"
          - kegg.compound: "C03263"
          - metanetx.chemical: "MNXM410"
          - sbo: "SBO:0000247"
      - deltaG: 10000000
    - !!omap
      - id: "s_0539"
      - name: "CTP"
      - compartment: "c"
      - formula: "C9H12N3O14P3"
      - charge: -4
      - smiles: "C1=CN(C(=O)N=C1N)C2C(C(C(O2)COP(=O)(O)OP(=O)(O)OP(=O)(O)O)O)O"
      - annotation: !!omap
          - bigg.metabolite: "ctp"
          - chebi: "CHEBI:37563"
          - kegg.compound: "C00063"
          - metanetx.chemical: "MNXM63"
          - sbo: "SBO:0000247"
      - deltaG: -637.62
    - !!omap
      - id: "s_0541"
      - name: "CTP"
      - compartment: "m"
      - formula: "C9H12N3O14P3"
      - charge: -4
      - smiles: "C1=CN(C(=O)N=C1N)C2C(C(C(O2)COP(=O)(O)OP(=O)(O)OP(=O)(O)O)O)O"
      - annotation: !!omap
          - bigg.metabolite: "ctp"
          - chebi: "CHEBI:37563"
          - kegg.compound: "C00063"
          - metanetx.chemical: "MNXM63"
          - sbo: "SBO:0000247"
      - deltaG: -637.62
    - !!omap
      - id: "s_0542"
      - name: "Cys-tRNA(Cys)"
      - compartment: "c"
      - formula: "C3H7NOSR"
      - charge: 1
      - annotation: !!omap
          - bigg.metabolite: "cystrna"
          - chebi: "CHEBI:29152"
          - kegg.compound: "C03125"
          - metanetx.chemical: "MNXM155005"
          - sbo: "SBO:0000247"
      - deltaG: 10000000
    - !!omap
      - id: "s_0543"
      - name: "cytidine"
      - compartment: "c"
      - formula: "C9H13N3O5"
      - charge: 0
      - smiles: "C1=CN(C(=O)N=C1N)C2C(C(C(O2)CO)O)O"
      - annotation: !!omap
          - bigg.metabolite: "cytd"
          - chebi: "CHEBI:17562"
          - kegg.compound: "C00475"
          - metanetx.chemical: "MNXM338"
          - sbo: "SBO:0000247"
      - deltaG: -6.3
    - !!omap
      - id: "s_0544"
      - name: "cytidine"
      - compartment: "e"
      - formula: "C9H13N3O5"
      - charge: 0
      - smiles: "C1=CN(C(=O)N=C1N)C2C(C(C(O2)CO)O)O"
      - annotation: !!omap
          - bigg.metabolite: "cytd"
          - chebi: "CHEBI:17562"
          - kegg.compound: "C00475"
          - metanetx.chemical: "MNXM338"
          - sbo: "SBO:0000247"
      - deltaG: -6.3
    - !!omap
      - id: "s_0545"
      - name: "cytosine"
      - compartment: "c"
      - formula: "C4H5N3O"
      - charge: 0
      - smiles: "C1=C(NC(=O)N=C1)N"
      - annotation: !!omap
          - bigg.metabolite: "csn"
          - chebi: "CHEBI:16040"
          - kegg.compound: "C00380"
          - metanetx.chemical: "MNXM761"
          - sbo: "SBO:0000247"
      - deltaG: 40
    - !!omap
      - id: "s_0546"
      - name: "cytosine"
      - compartment: "e"
      - formula: "C4H5N3O"
      - charge: 0
      - smiles: "C1=C(NC(=O)N=C1)N"
      - annotation: !!omap
          - bigg.metabolite: "csn"
          - chebi: "CHEBI:16040"
          - kegg.compound: "C00380"
          - metanetx.chemical: "MNXM761"
          - sbo: "SBO:0000247"
      - deltaG: 40
    - !!omap
      - id: "s_0547"
      - name: "D-arabinono-1,4-lactone"
      - compartment: "c"
      - formula: "C5H8O5"
      - charge: 0
      - smiles: "C(C1C(C(C(=O)O1)O)O)O"
      - annotation: !!omap
          - bigg.metabolite: "Dara14lac"
          - chebi: "CHEBI:16292"
          - kegg.compound: "C00652"
          - metanetx.chemical: "MNXM778"
          - sbo: "SBO:0000247"
      - deltaG: -101.47
    - !!omap
      - id: "s_0548"
      - name: "D-arabinose"
      - compartment: "c"
      - formula: "C5H10O5"
      - charge: 0
      - annotation: !!omap
          - bigg.metabolite: "arab__D"
          - chebi: "CHEBI:17108"
          - kegg.compound: "C00216"
          - metanetx.chemical: "MNXM40571"
          - sbo: "SBO:0000247"
      - deltaG: -82.69
    - !!omap
      - id: "s_0549"
      - name: "D-arabinose"
      - compartment: "e"
      - formula: "C5H10O5"
      - charge: 0
      - annotation: !!omap
          - bigg.metabolite: "arab__D"
          - chebi: "CHEBI:17108"
          - kegg.compound: "C00216"
          - metanetx.chemical: "MNXM40571"
          - sbo: "SBO:0000247"
      - deltaG: -82.69
    - !!omap
      - id: "s_0550"
      - name: "D-erythro-1-(imidazol-4-yl)glycerol 3-phosphate"
      - compartment: "c"
      - formula: "C6H9N2O6P"
      - charge: -2
      - smiles: "C1=C(NC=N1)C(C(COP(=O)(O)O)O)O"
      - annotation: !!omap
          - bigg.metabolite: "eig3p"
          - chebi: "CHEBI:58278"
          - kegg.compound: "C04666"
          - metanetx.chemical: "MNXM1422"
          - sbo: "SBO:0000247"
      - deltaG: -201.4
    - !!omap
      - id: "s_0551"
      - name: "D-erythrose 4-phosphate"
      - compartment: "c"
      - formula: "C4H7O7P"
      - charge: -2
      - smiles: "C(C(C(C=O)O)O)OP(=O)(O)O"
      - annotation: !!omap
          - bigg.metabolite: "e4p"
          - chebi: "CHEBI:16897"
          - kegg.compound: "C00279"
          - metanetx.chemical: "MNXM258"
          - sbo: "SBO:0000247"
      - deltaG: -288.29
    - !!omap
      - id: "s_0552"
      - name: "D-erythrose 4-phosphate"
      - compartment: "m"
      - formula: "C4H7O7P"
      - charge: -2
      - smiles: "C(C(C(C=O)O)O)OP(=O)(O)O"
      - annotation: !!omap
          - bigg.metabolite: "e4p"
          - chebi: "CHEBI:16897"
          - kegg.compound: "C00279"
          - metanetx.chemical: "MNXM258"
          - sbo: "SBO:0000247"
      - deltaG: -288.29
    - !!omap
      - id: "s_0553"
      - name: "D-fructose"
      - compartment: "c"
      - formula: "C6H12O6"
      - charge: 0
      - smiles: "C1C(C(C(C(O1)(CO)O)O)O)O"
      - annotation: !!omap
          - bigg.metabolite: "fru"
          - chebi: "CHEBI:15824"
          - kegg.compound: "C00095"
          - metanetx.chemical: "MNXM175"
          - sbo: "SBO:0000247"
      - deltaG: -102.33
    - !!omap
      - id: "s_0554"
      - name: "D-fructose"
      - compartment: "e"
      - formula: "C6H12O6"
      - charge: 0
      - smiles: "C1C(C(C(C(O1)(CO)O)O)O)O"
      - annotation: !!omap
          - bigg.metabolite: "fru"
          - chebi: "CHEBI:15824"
          - kegg.compound: "C00095"
          - metanetx.chemical: "MNXM175"
          - sbo: "SBO:0000247"
      - deltaG: -102.33
    - !!omap
      - id: "s_0555"
      - name: "D-fructose 1,6-bisphosphate"
      - compartment: "c"
      - formula: "C6H10O12P2"
      - charge: -4
      - annotation: !!omap
          - bigg.metabolite: "fdp"
          - chebi: "CHEBI:16905"
          - kegg.compound: "C00354"
          - metanetx.chemical: "MNXM417"
          - sbo: "SBO:0000247"
      - deltaG: -549.91
    - !!omap
      - id: "s_0556"
      - name: "D-fructose 1-phosphate"
      - compartment: "c"
      - formula: "C6H11O9P"
      - charge: -2
      - smiles: "C(C(C(C(C(=O)COP(=O)(O)O)O)O)O)O"
      - annotation: !!omap
          - bigg.metabolite: "f1p"
          - chebi: "CHEBI:18105"
          - kegg.compound: "C01094"
          - metanetx.chemical: "MNXM145568"
          - sbo: "SBO:0000247"
      - deltaG: -325.36
    - !!omap
      - id: "s_0557"
      - name: "D-fructose 6-phosphate"
      - compartment: "c"
      - formula: "C6H11O9P"
      - charge: -2
      - smiles: "C(C(C(C(C(=O)CO)O)O)O)OP(=O)(O)O"
      - annotation: !!omap
          - bigg.metabolite: "f6p"
          - chebi: "CHEBI:57579"
          - kegg.compound: "C00085"
          - metanetx.chemical: "MNXM162235"
          - sbo: "SBO:0000247"
      - deltaG: -325.37
    - !!omap
      - id: "s_0558"
      - name: "D-galactose"
      - compartment: "c"
      - formula: "C6H12O6"
      - charge: 0
      - smiles: "C(C1C(C(C(C(O1)O)O)O)O)O"
      - annotation: !!omap
          - bigg.metabolite: "gal"
          - chebi: "CHEBI:12936"
          - kegg.compound: "C00124"
          - metanetx.chemical: "MNXM390"
          - sbo: "SBO:0000247"
      - deltaG: -101.44
    - !!omap
      - id: "s_0559"
      - name: "D-galactose"
      - compartment: "e"
      - formula: "C6H12O6"
      - charge: 0
      - smiles: "C(C1C(C(C(C(O1)O)O)O)O)O"
      - annotation: !!omap
          - bigg.metabolite: "gal"
          - chebi: "CHEBI:12936"
          - kegg.compound: "C00124"
          - metanetx.chemical: "MNXM390"
          - sbo: "SBO:0000247"
      - deltaG: -101.44
    - !!omap
      - id: "s_0560"
      - name: "D-galacturonate"
      - compartment: "e"
      - formula: "C6H9O7"
      - charge: -1
      - smiles: "C1(C(C(OC(C1O)O)C(=O)O)O)O"
      - annotation: !!omap
          - bigg.metabolite: "galur"
          - chebi: "CHEBI:12952"
          - kegg.compound: "C00333"
          - metanetx.chemical: "MNXM693"
          - sbo: "SBO:0000247"
      - deltaG: -200.16
    - !!omap
      - id: "s_0561"
      - name: "D-glucitol"
      - compartment: "c"
      - formula: "C6H14O6"
      - charge: 0
      - smiles: "C(C(C(C(C(CO)O)O)O)O)O"
      - annotation: !!omap
          - bigg.metabolite: "sbt__D"
          - chebi: "CHEBI:17924"
          - kegg.compound: "C00794"
          - metanetx.chemical: "MNXM469"
          - sbo: "SBO:0000247"
      - deltaG: -89.12
    - !!omap
      - id: "s_0562"
      - name: "D-glucitol"
      - compartment: "e"
      - formula: "C6H14O6"
      - charge: 0
      - smiles: "C(C(C(C(C(CO)O)O)O)O)O"
      - annotation: !!omap
          - bigg.metabolite: "sbt__D"
          - chebi: "CHEBI:17924"
          - kegg.compound: "C00794"
          - metanetx.chemical: "MNXM469"
          - sbo: "SBO:0000247"
      - deltaG: -89.12
    - !!omap
      - id: "s_0563"
      - name: "D-glucose"
      - compartment: "c"
      - formula: "C6H12O6"
      - charge: 0
      - smiles: "C(C1C(C(C(C(O1)O)O)O)O)O"
      - annotation: !!omap
          - bigg.metabolite: "glc__D"
          - chebi: "CHEBI:4167"
          - kegg.compound: "C00031"
          - metanetx.chemical: "MNXM41"
          - sbo: "SBO:0000247"
      - deltaG: -101.44
    - !!omap
      - id: "s_0565"
      - name: "D-glucose"
      - compartment: "e"
      - formula: "C6H12O6"
      - charge: 0
      - smiles: "C(C1C(C(C(C(O1)O)O)O)O)O"
      - annotation: !!omap
          - bigg.metabolite: "glc__D"
          - chebi: "CHEBI:4167"
          - kegg.compound: "C00031"
          - metanetx.chemical: "MNXM41"
          - sbo: "SBO:0000247"
      - deltaG: -101.44
    - !!omap
      - id: "s_0566"
      - name: "D-glucose"
      - compartment: "v"
      - formula: "C6H12O6"
      - charge: 0
      - smiles: "C(C1C(C(C(C(O1)O)O)O)O)O"
      - annotation: !!omap
          - bigg.metabolite: "glc__D"
          - chebi: "CHEBI:4167"
          - kegg.compound: "C00031"
          - metanetx.chemical: "MNXM41"
          - sbo: "SBO:0000247"
      - deltaG: -101.44
    - !!omap
      - id: "s_0567"
      - name: "D-glucose 1-phosphate"
      - compartment: "c"
      - formula: "C6H11O9P"
      - charge: -2
      - smiles: "C(C1C(C(C(C(O1)OP(=O)(O)O)O)O)O)O"
      - annotation: !!omap
          - bigg.metabolite: "g1p"
          - chebi: "CHEBI:57629"
          - kegg.compound: "C00103"
          - metanetx.chemical: "MNXM89588"
          - sbo: "SBO:0000247"
      - deltaG: -324.47
    - !!omap
      - id: "s_0568"
      - name: "D-glucose 6-phosphate"
      - compartment: "c"
      - formula: "C6H11O9P"
      - charge: -2
      - smiles: "C(C(C(C(C(C=O)O)O)O)O)OP(=O)(O)O"
      - annotation: !!omap
          - bigg.metabolite: "g6p"
          - chebi: "CHEBI:14314"
          - kegg.compound: "C00092"
          - metanetx.chemical: "MNXM160"
          - sbo: "SBO:0000247"
      - deltaG: -324.48
    - !!omap
      - id: "s_0570"
      - name: "D-glyceraldehyde"
      - compartment: "c"
      - formula: "C3H6O3"
      - charge: 0
      - smiles: "C(C(C=O)O)O"
      - annotation: !!omap
          - bigg.metabolite: "glyald"
          - chebi: "CHEBI:17378"
          - kegg.compound: "C00577"
          - metanetx.chemical: "MNXM435"
          - sbo: "SBO:0000247"
      - deltaG: -48.28
    - !!omap
      - id: "s_0571"
      - name: "D-mannose"
      - compartment: "c"
      - formula: "C6H12O6"
      - charge: 0
      - smiles: "C(C1C(C(C(C(O1)O)O)O)O)O"
      - annotation: !!omap
          - bigg.metabolite: "man"
          - chebi: "CHEBI:16024"
          - kegg.compound: "C00159"
          - metanetx.chemical: "MNXM182"
          - sbo: "SBO:0000247"
      - deltaG: -101.44
    - !!omap
      - id: "s_0572"
      - name: "D-mannose"
      - compartment: "e"
      - formula: "C6H12O6"
      - charge: 0
      - smiles: "C(C1C(C(C(C(O1)O)O)O)O)O"
      - annotation: !!omap
          - bigg.metabolite: "man"
          - chebi: "CHEBI:16024"
          - kegg.compound: "C00159"
          - metanetx.chemical: "MNXM182"
          - sbo: "SBO:0000247"
      - deltaG: -101.44
    - !!omap
      - id: "s_0573"
      - name: "D-mannose 1-phosphate"
      - compartment: "c"
      - formula: "C6H11O9P"
      - charge: -2
      - smiles: "C(C1C(C(C(C(O1)OP(=O)(O)O)O)O)O)O"
      - annotation: !!omap
          - bigg.metabolite: "man1p"
          - chebi: "CHEBI:35374"
          - kegg.compound: "C00636"
          - metanetx.chemical: "MNXM721"
          - sbo: "SBO:0000247"
      - deltaG: -324.47
    - !!omap
      - id: "s_0574"
      - name: "D-mannose 6-phosphate"
      - compartment: "c"
      - formula: "C6H11O9P"
      - charge: -2
      - smiles: "C(C1C(C(C(C(O1)O)O)O)O)OP(=O)(O)O"
      - annotation: !!omap
          - bigg.metabolite: "man6p"
          - chebi: "CHEBI:17369"
          - kegg.compound: "C00275"
          - metanetx.chemical: "MNXM427"
          - sbo: "SBO:0000247"
      - deltaG: -324.48
    - !!omap
      - id: "s_0575"
      - name: "D-ribose"
      - compartment: "c"
      - formula: "C5H10O5"
      - charge: 0
      - smiles: "C1C(C(C(C(O1)O)O)O)O"
      - annotation: !!omap
          - bigg.metabolite: "rib__D"
          - chebi: "CHEBI:16988"
          - kegg.compound: "C00121"
          - metanetx.chemical: "MNXM242"
          - sbo: "SBO:0000247"
      - deltaG: -84.25
    - !!omap
      - id: "s_0576"
      - name: "D-ribose"
      - compartment: "e"
      - formula: "C5H10O5"
      - charge: 0
      - smiles: "C1C(C(C(C(O1)O)O)O)O"
      - annotation: !!omap
          - bigg.metabolite: "rib__D"
          - chebi: "CHEBI:16988"
          - kegg.compound: "C00121"
          - metanetx.chemical: "MNXM242"
          - sbo: "SBO:0000247"
      - deltaG: -84.25
    - !!omap
      - id: "s_0577"
      - name: "D-ribulose 5-phosphate"
      - compartment: "c"
      - formula: "C5H9O8P"
      - charge: -2
      - smiles: "C(C(C(C(=O)CO)O)O)OP(=O)(O)O"
      - annotation: !!omap
          - bigg.metabolite: "ru5p__D"
          - chebi: "CHEBI:58121"
          - kegg.compound: "C00199"
          - metanetx.chemical: "MNXM145"
          - sbo: "SBO:0000247"
      - deltaG: -306.69
    - !!omap
      - id: "s_0578"
      - name: "D-xylose"
      - compartment: "c"
      - formula: "C5H10O5"
      - charge: 0
      - smiles: "C1C(C(C(C(O1)O)O)O)O"
      - annotation: !!omap
          - bigg.metabolite: "xyl__D"
          - chebi: "CHEBI:15936"
          - kegg.compound: "C00181"
          - metanetx.chemical: "MNXM90941"
          - sbo: "SBO:0000247"
      - deltaG: -82.69
    - !!omap
      - id: "s_0579"
      - name: "D-xylose"
      - compartment: "e"
      - formula: "C5H10O5"
      - charge: 0
      - smiles: "C1C(C(C(C(O1)O)O)O)O"
      - annotation: !!omap
          - bigg.metabolite: "xyl__D"
          - chebi: "CHEBI:15936"
          - kegg.compound: "C00181"
          - metanetx.chemical: "MNXM90941"
          - sbo: "SBO:0000247"
      - deltaG: -82.69
    - !!omap
      - id: "s_0580"
      - name: "D-xylulose"
      - compartment: "c"
      - formula: "C5H10O5"
      - charge: 0
      - smiles: "C(C(C(C(=O)CO)O)O)O"
      - annotation: !!omap
          - bigg.metabolite: "xylu__D"
          - chebi: "CHEBI:17140"
          - kegg.compound: "C00310"
          - metanetx.chemical: "MNXM597"
          - sbo: "SBO:0000247"
      - deltaG: -83.58
    - !!omap
      - id: "s_0581"
      - name: "D-xylulose 5-phosphate"
      - compartment: "c"
      - formula: "C5H9O8P"
      - charge: -2
      - smiles: "C(C(C(C(=O)CO)O)O)OP(=O)(O)O"
      - annotation: !!omap
          - bigg.metabolite: "xu5p__D"
          - chebi: "CHEBI:57737"
          - kegg.compound: "C00231"
          - metanetx.chemical: "MNXM186"
          - sbo: "SBO:0000247"
      - deltaG: -306.69
    - !!omap
      - id: "s_0582"
      - name: "dADP"
      - compartment: "c"
      - formula: "C10H12N5O9P2"
      - charge: -3
      - smiles: "C1C(C(OC1N2C=NC3=C(N=CN=C32)N)COP(=O)(O)OP(=O)(O)O)O"
      - annotation: !!omap
          - bigg.metabolite: "dadp"
          - chebi: "CHEBI:57667"
          - kegg.compound: "C00206"
          - metanetx.chemical: "MNXM374"
          - sbo: "SBO:0000247"
      - deltaG: -316.75
    - !!omap
      - id: "s_0583"
      - name: "dADP"
      - compartment: "n"
      - formula: "C10H12N5O9P2"
      - charge: -3
      - smiles: "C1C(C(OC1N2C=NC3=C(N=CN=C32)N)COP(=O)(O)OP(=O)(O)O)O"
      - annotation: !!omap
          - bigg.metabolite: "dadp"
          - chebi: "CHEBI:57667"
          - kegg.compound: "C00206"
          - metanetx.chemical: "MNXM374"
          - sbo: "SBO:0000247"
      - deltaG: -316.75
    - !!omap
      - id: "s_0584"
      - name: "dAMP"
      - compartment: "c"
      - formula: "C10H12N5O6P"
      - charge: -2
      - smiles: "C1C(C(OC1N2C=NC3=C(N=CN=C32)N)COP(=O)(O)O)O"
      - annotation: !!omap
          - bigg.metabolite: "damp"
          - chebi: "CHEBI:58245"
          - kegg.compound: "C00360"
          - metanetx.chemical: "MNXM432"
          - sbo: "SBO:0000247"
      - deltaG: -112.21
    - !!omap
      - id: "s_0585"
      - name: "dAMP"
      - compartment: "m"
      - formula: "C10H12N5O6P"
      - charge: -2
      - smiles: "C1C(C(OC1N2C=NC3=C(N=CN=C32)N)COP(=O)(O)O)O"
      - annotation: !!omap
          - bigg.metabolite: "damp"
          - chebi: "CHEBI:58245"
          - kegg.compound: "C00360"
          - metanetx.chemical: "MNXM432"
          - sbo: "SBO:0000247"
      - deltaG: -112.21
    - !!omap
      - id: "s_0586"
      - name: "dATP"
      - compartment: "c"
      - formula: "C10H12N5O12P3"
      - charge: -4
      - smiles: "C1C(C(OC1N2C=NC3=C(N=CN=C32)N)COP(=O)(O)OP(=O)(O)OP(=O)(O)O)O"
      - annotation: !!omap
          - bigg.metabolite: "datp"
          - chebi: "CHEBI:61404"
          - kegg.compound: "C00131"
          - metanetx.chemical: "MNXM286"
          - sbo: "SBO:0000247"
      - deltaG: -524.57
    - !!omap
      - id: "s_0587"
      - name: "dCDP"
      - compartment: "c"
      - formula: "C9H12N3O10P2"
      - charge: -3
      - smiles: "C1C(C(OC1N2C=CC(=NC2=O)N)COP(=O)(O)OP(=O)(O)O)O"
      - annotation: !!omap
          - bigg.metabolite: "dcdp"
          - chebi: "CHEBI:58593"
          - kegg.compound: "C00705"
          - metanetx.chemical: "MNXM411"
          - sbo: "SBO:0000247"
      - deltaG: -388.51
    - !!omap
      - id: "s_0588"
      - name: "dCDP"
      - compartment: "n"
      - formula: "C9H12N3O10P2"
      - charge: -3
      - smiles: "C1C(C(OC1N2C=CC(=NC2=O)N)COP(=O)(O)OP(=O)(O)O)O"
      - annotation: !!omap
          - bigg.metabolite: "dcdp"
          - chebi: "CHEBI:58593"
          - kegg.compound: "C00705"
          - metanetx.chemical: "MNXM411"
          - sbo: "SBO:0000247"
      - deltaG: -388.51
    - !!omap
      - id: "s_0589"
      - name: "dCMP"
      - compartment: "c"
      - formula: "C9H12N3O7P"
      - charge: -2
      - smiles: "C1C(C(OC1N2C=CC(=NC2=O)N)COP(=O)(O)O)O"
      - annotation: !!omap
          - bigg.metabolite: "dcmp"
          - chebi: "CHEBI:57566"
          - kegg.compound: "C00239"
          - metanetx.chemical: "MNXM266"
          - sbo: "SBO:0000247"
      - deltaG: -189.5
    - !!omap
      - id: "s_0590"
      - name: "dCTP"
      - compartment: "c"
      - formula: "C9H12N3O13P3"
      - charge: -4
      - smiles: "C1C(C(OC1N2C=CC(=NC2=O)N)COP(=O)(O)OP(=O)(O)OP(=O)(O)O)O"
      - annotation: !!omap
          - bigg.metabolite: "dctp"
          - chebi: "CHEBI:57724"
          - kegg.compound: "C00458"
          - metanetx.chemical: "MNXM360"
          - sbo: "SBO:0000247"
      - deltaG: -597.78
    - !!omap
      - id: "s_0591"
      - name: "deamido-NAD(+)"
      - compartment: "c"
      - formula: "C21H24N6O15P2"
      - charge: -2
      - smiles: "C1=CC(=C[N+](=C1)C2C(C(C(O2)COP(=O)(O)OP(=O)(O)OCC3C(C(C(O3)N4C=NC5=C(N=CN=C54)N)O)O)O)O)C(=O)O"
      - annotation: !!omap
          - bigg.metabolite: "dnad"
          - chebi: "CHEBI:18304"
          - kegg.compound: "C00857"
          - metanetx.chemical: "MNXM309"
          - sbo: "SBO:0000247"
      - deltaG: -361.56
    - !!omap
      - id: "s_0593"
      - name: "deamido-NAD(+)"
      - compartment: "n"
      - formula: "C21H24N6O15P2"
      - charge: -2
      - smiles: "C1=CC(=C[N+](=C1)C2C(C(C(O2)COP(=O)(O)OP(=O)(O)OCC3C(C(C(O3)N4C=NC5=C(N=CN=C54)N)O)O)O)O)C(=O)O"
      - annotation: !!omap
          - bigg.metabolite: "dnad"
          - chebi: "CHEBI:18304"
          - kegg.compound: "C00857"
          - metanetx.chemical: "MNXM309"
          - sbo: "SBO:0000247"
      - deltaG: -361.56
    - !!omap
      - id: "s_0595"
      - name: "decanoate"
      - compartment: "c"
      - formula: "C10H19O2"
      - charge: -1
      - smiles: "CCCCCCCCCC(=O)[O-]"
      - annotation: !!omap
          - bigg.metabolite: "dca"
          - chebi: "CHEBI:27689"
          - kegg.compound: "C01571"
          - metanetx.chemical: "MNXM1043"
          - sbo: "SBO:0000247"
      - deltaG: 111.01
    - !!omap
      - id: "s_0597"
      - name: "decanoate"
      - compartment: "e"
      - formula: "C10H19O2"
      - charge: -1
      - smiles: "CCCCCCCCCC(=O)[O-]"
      - annotation: !!omap
          - bigg.metabolite: "dca"
          - chebi: "CHEBI:27689"
          - kegg.compound: "C01571"
          - metanetx.chemical: "MNXM1043"
          - sbo: "SBO:0000247"
      - deltaG: 111.01
    - !!omap
      - id: "s_0600"
      - name: "decanoate"
      - compartment: "p"
      - formula: "C10H19O2"
      - charge: -1
      - smiles: "CCCCCCCCCC(=O)[O-]"
      - annotation: !!omap
          - bigg.metabolite: "dca"
          - chebi: "CHEBI:27689"
          - kegg.compound: "C01571"
          - metanetx.chemical: "MNXM1043"
          - sbo: "SBO:0000247"
      - deltaG: 111.01
    - !!omap
      - id: "s_0605"
      - name: "decanoyl-CoA"
      - compartment: "p"
      - formula: "C31H50N7O17P3S"
      - charge: -4
      - smiles: "CCCCCCCCCC(=O)SCCNC(=O)CCNC(=O)C(C(C)(C)COP(=O)(O)OP(=O)(O)OCC1C(C(C(O1)N2C=NC3=C(N=CN=C32)N)O)OP(=O)(O)O)O"
      - annotation: !!omap
          - bigg.metabolite: "dccoa"
          - chebi: "CHEBI:61430"
          - kegg.compound: "C05274"
          - metanetx.chemical: "MNXM486"
          - sbo: "SBO:0000247"
      - deltaG: -224.61
    - !!omap
      - id: "s_0606"
      - name: "decaprenyl diphosphate"
      - compartment: "lp"
      - formula: "C50H81O7P2"
      - charge: -3
      - smiles: "CC(=CCCC(=CCCC(=CCCC(=CCCC(=CCCC(=CCCC(=CCCC(=CCCC(=CCCC(=CCOP(=O)(O)OP(=O)(O)O)C)C)C)C)C)C)C)C)C)C"
      - annotation: !!omap
          - bigg.metabolite: "decdp"
          - chebi: "CHEBI:53043"
          - kegg.compound: "C17432"
          - metanetx.chemical: "MNXM1721"
          - sbo: "SBO:0000247"
      - deltaG: 610.5
    - !!omap
      - id: "s_0607"
      - name: "dehydro-D-arabinono-1,4-lactone"
      - compartment: "c"
      - formula: "C5H5O5"
      - charge: -1
      - smiles: "C(C1C(=C(C(=O)O1)O)O)O"
      - annotation: !!omap
          - chebi: "CHEBI:58277"
          - kegg.compound: "C06316"
          - metanetx.chemical: "MNXM3254"
          - sbo: "SBO:0000247"
      - deltaG: -110
    - !!omap
      - id: "s_0608"
      - name: "Delta(6)-trans,Delta(8)-cis-leukotriene B4"
      - compartment: "c"
      - formula: "C20H31O4"
      - charge: -1
      - smiles: "CCCCCC=CCC(C=CC=CC=CC(CCCC(=O)O)O)O"
      - annotation: !!omap
          - chebi: "CHEBI:53027"
          - metanetx.chemical: "MNXM507621"
          - sbo: "SBO:0000247"
      - deltaG: 247.78
    - !!omap
      - id: "s_0609"
      - name: "Delta(6)-trans,Delta(8)-cis-leukotriene B4"
      - compartment: "n"
      - formula: "C20H31O4"
      - charge: -1
      - smiles: "CCCCCC=CCC(C=CC=CC=CC(CCCC(=O)O)O)O"
      - annotation: !!omap
          - chebi: "CHEBI:53027"
          - metanetx.chemical: "MNXM507621"
          - sbo: "SBO:0000247"
      - deltaG: 247.78
    - !!omap
      - id: "s_0610"
      - name: "deoxycytidine"
      - compartment: "c"
      - formula: "C9H13N3O4"
      - charge: 0
      - smiles: "C1C(C(OC1N2C=CC(=NC2=O)N)CO)O"
      - annotation: !!omap
          - bigg.metabolite: "dcyt"
          - chebi: "CHEBI:15698"
          - kegg.compound: "C00881"
          - metanetx.chemical: "MNXM704"
          - sbo: "SBO:0000247"
      - deltaG: 33.54
    - !!omap
      - id: "s_0611"
      - name: "deoxycytidine"
      - compartment: "e"
      - formula: "C9H13N3O4"
      - charge: 0
      - smiles: "C1C(C(OC1N2C=CC(=NC2=O)N)CO)O"
      - annotation: !!omap
          - bigg.metabolite: "dcyt"
          - chebi: "CHEBI:15698"
          - kegg.compound: "C00881"
          - metanetx.chemical: "MNXM704"
          - sbo: "SBO:0000247"
      - deltaG: 33.54
    - !!omap
      - id: "s_0612"
      - name: "dethiobiotin"
      - compartment: "c"
      - formula: "C10H17N2O3"
      - charge: -1
      - smiles: "CC1C(NC(=O)N1)CCCCCC(=O)O"
      - annotation: !!omap
          - bigg.metabolite: "dtbt"
          - chebi: "CHEBI:57861"
          - kegg.compound: "C01909"
          - metanetx.chemical: "MNXM1020"
          - sbo: "SBO:0000247"
      - deltaG: 50.02
    - !!omap
      - id: "s_0613"
      - name: "dGDP"
      - compartment: "c"
      - formula: "C10H12N5O10P2"
      - charge: -3
      - smiles: "C1C(C(OC1N2C=NC3=C2N=C(NC3=O)N)COP(=O)(O)OP(=O)(O)O)O"
      - annotation: !!omap
          - bigg.metabolite: "dgdp"
          - chebi: "CHEBI:58595"
          - kegg.compound: "C00361"
          - metanetx.chemical: "MNXM436"
          - sbo: "SBO:0000247"
      - deltaG: -351.43
    - !!omap
      - id: "s_0614"
      - name: "dGDP"
      - compartment: "n"
      - formula: "C10H12N5O10P2"
      - charge: -3
      - smiles: "C1C(C(OC1N2C=NC3=C2N=C(NC3=O)N)COP(=O)(O)OP(=O)(O)O)O"
      - annotation: !!omap
          - bigg.metabolite: "dgdp"
          - chebi: "CHEBI:58595"
          - kegg.compound: "C00361"
          - metanetx.chemical: "MNXM436"
          - sbo: "SBO:0000247"
      - deltaG: -351.43
    - !!omap
      - id: "s_0615"
      - name: "dGMP"
      - compartment: "c"
      - formula: "C10H12N5O7P"
      - charge: -2
      - smiles: "C1C(C(OC1N2C=NC3=C2N=C(NC3=O)N)COP(=O)(O)O)O"
      - annotation: !!omap
          - bigg.metabolite: "dgmp"
          - chebi: "CHEBI:57673"
          - kegg.compound: "C00362"
          - metanetx.chemical: "MNXM546"
          - sbo: "SBO:0000247"
      - deltaG: -152.42
    - !!omap
      - id: "s_0616"
      - name: "dGMP"
      - compartment: "m"
      - formula: "C10H12N5O7P"
      - charge: -2
      - smiles: "C1C(C(OC1N2C=NC3=C2N=C(NC3=O)N)COP(=O)(O)O)O"
      - annotation: !!omap
          - bigg.metabolite: "dgmp"
          - chebi: "CHEBI:57673"
          - kegg.compound: "C00362"
          - metanetx.chemical: "MNXM546"
          - sbo: "SBO:0000247"
      - deltaG: -152.42
    - !!omap
      - id: "s_0617"
      - name: "dGTP"
      - compartment: "c"
      - formula: "C10H12N5O13P3"
      - charge: -4
      - smiles: "C1C(C(OC1N2C=NC3=C2N=C(NC3=O)N)COP(=O)(O)OP(=O)(O)OP(=O)(O)O)O"
      - annotation: !!omap
          - bigg.metabolite: "dgtp"
          - chebi: "CHEBI:57794"
          - kegg.compound: "C00286"
          - metanetx.chemical: "MNXM344"
          - sbo: "SBO:0000247"
      - deltaG: -560.7
    - !!omap
      - id: "s_0618"
      - name: "dIDP"
      - compartment: "c"
      - formula: "C10H11N4O10P2"
      - charge: -3
      - annotation: !!omap
          - bigg.metabolite: "didp"
          - chebi: "CHEBI:28823"
          - kegg.compound: "C01344"
          - metanetx.chemical: "MNXM2174"
          - sbo: "SBO:0000247"
      - deltaG: -369.81
    - !!omap
      - id: "s_0620"
      - name: "diglyceride backbone"
      - compartment: "er"
      - formula: "C3H4O"
      - annotation: !!omap
          - bigg.metabolite: "dag_hs"
          - chebi: "CHEBI:18035"
          - kegg.compound: "C00165"
          - metanetx.chemical: "MNXM59"
          - sbo: "SBO:0000649"
      - deltaG: 10000000
    - !!omap
      - id: "s_0625"
      - name: "dihydrofolic acid"
      - compartment: "c"
      - formula: "C19H19N7O6"
      - charge: -2
      - smiles: "C1C(=NC2=C(N1)N=C(NC2=O)N)CNC3=CC=C(C=C3)C(=O)NC(CCC(=O)O)C(=O)O"
      - annotation: !!omap
          - bigg.metabolite: "dhf"
          - chebi: "CHEBI:57451"
          - kegg.compound: "C00415"
          - metanetx.chemical: "MNXM281"
          - sbo: "SBO:0000247"
      - deltaG: 74.53
    - !!omap
      - id: "s_0626"
      - name: "dihydrofolic acid"
      - compartment: "m"
      - formula: "C19H19N7O6"
      - charge: -2
      - smiles: "C1C(=NC2=C(N1)N=C(NC2=O)N)CNC3=CC=C(C=C3)C(=O)NC(CCC(=O)O)C(=O)O"
      - annotation: !!omap
          - bigg.metabolite: "dhf"
          - chebi: "CHEBI:57451"
          - kegg.compound: "C00415"
          - metanetx.chemical: "MNXM281"
          - sbo: "SBO:0000247"
      - deltaG: 74.53
    - !!omap
      - id: "s_0627"
      - name: "dihydrolipoamide"
      - compartment: "m"
      - formula: "C8H17NOS2"
      - charge: 0
      - smiles: "C(CCC(=O)N)CC(CCS)S"
      - annotation: !!omap
          - bigg.metabolite: "dhlam"
          - chebi: "CHEBI:17694"
          - kegg.compound: "C00579"
          - metanetx.chemical: "MNXM1277"
          - sbo: "SBO:0000247"
      - deltaG: 149.77
    - !!omap
      - id: "s_0628"
      - name: "dihydrolipoylprotein"
      - compartment: "m"
      - formula: "C8H16NOS2R"
      - charge: 0
      - annotation: !!omap
          - bigg.metabolite: "dhlpro"
          - chebi: "CHEBI:16194"
          - kegg.compound: "C02972"
          - metanetx.chemical: "MNXM1663"
          - sbo: "SBO:0000247"
      - deltaG: 10000000
    - !!omap
      - id: "s_0629"
      - name: "dihydroxyacetone phosphate"
      - compartment: "c"
      - formula: "C3H5O6P"
      - charge: -2
      - smiles: "C(C(=O)COP(=O)(O)O)O"
      - annotation: !!omap
          - bigg.metabolite: "dhap"
          - chebi: "CHEBI:57642"
          - kegg.compound: "C00111"
          - metanetx.chemical: "MNXM77"
          - sbo: "SBO:0000247"
      - deltaG: -272.78
    - !!omap
      - id: "s_0631"
      - name: "dihydroxyacetone phosphate"
      - compartment: "lp"
      - formula: "C3H5O6P"
      - charge: -2
      - smiles: "C(C(=O)COP(=O)(O)O)O"
      - annotation: !!omap
          - bigg.metabolite: "dhap"
          - chebi: "CHEBI:57642"
          - kegg.compound: "C00111"
          - metanetx.chemical: "MNXM77"
          - sbo: "SBO:0000247"
      - deltaG: -272.78
    - !!omap
      - id: "s_0632"
      - name: "dihydroxyacetone phosphate"
      - compartment: "m"
      - formula: "C3H5O6P"
      - charge: -2
      - smiles: "C(C(=O)COP(=O)(O)O)O"
      - annotation: !!omap
          - bigg.metabolite: "dhap"
          - chebi: "CHEBI:57642"
          - kegg.compound: "C00111"
          - metanetx.chemical: "MNXM77"
          - sbo: "SBO:0000247"
      - deltaG: -272.78
    - !!omap
      - id: "s_0633"
      - name: "diphosphate"
      - compartment: "c"
      - formula: "HO7P2"
      - charge: -3
      - smiles: "[O-]P(=O)([O-])OP(=O)([O-])[O-]"
      - annotation: !!omap
          - bigg.metabolite: "ppi"
          - chebi: "CHEBI:33019"
          - kegg.compound: "C00013"
          - metanetx.chemical: "MNXM11"
          - sbo: "SBO:0000247"
      - deltaG: -473.36
    - !!omap
      - id: "s_0635"
      - name: "diphosphate"
      - compartment: "lp"
      - formula: "HO7P2"
      - charge: -3
      - smiles: "[O-]P(=O)([O-])OP(=O)([O-])[O-]"
      - annotation: !!omap
          - bigg.metabolite: "ppi"
          - chebi: "CHEBI:33019"
          - kegg.compound: "C00013"
          - metanetx.chemical: "MNXM11"
          - sbo: "SBO:0000247"
      - deltaG: -473.36
    - !!omap
      - id: "s_0636"
      - name: "diphosphate"
      - compartment: "m"
      - formula: "HO7P2"
      - charge: -3
      - smiles: "[O-]P(=O)([O-])OP(=O)([O-])[O-]"
      - annotation: !!omap
          - bigg.metabolite: "ppi"
          - chebi: "CHEBI:33019"
          - kegg.compound: "C00013"
          - metanetx.chemical: "MNXM11"
          - sbo: "SBO:0000247"
      - deltaG: -473.36
    - !!omap
      - id: "s_0637"
      - name: "diphosphate"
      - compartment: "n"
      - formula: "HO7P2"
      - charge: -3
      - smiles: "[O-]P(=O)([O-])OP(=O)([O-])[O-]"
      - annotation: !!omap
          - bigg.metabolite: "ppi"
          - chebi: "CHEBI:33019"
          - kegg.compound: "C00013"
          - metanetx.chemical: "MNXM11"
          - sbo: "SBO:0000247"
      - deltaG: -473.36
    - !!omap
      - id: "s_0638"
      - name: "diphosphate"
      - compartment: "p"
      - formula: "HO7P2"
      - charge: -3
      - smiles: "[O-]P(=O)([O-])OP(=O)([O-])[O-]"
      - annotation: !!omap
          - bigg.metabolite: "ppi"
          - chebi: "CHEBI:33019"
          - kegg.compound: "C00013"
          - metanetx.chemical: "MNXM11"
          - sbo: "SBO:0000247"
      - deltaG: -473.36
    - !!omap
      - id: "s_0639"
      - name: "dITP"
      - compartment: "c"
      - formula: "C10H11N4O13P3"
      - charge: -4
      - smiles: "C1C(C(OC1N2C=NC3=C2N=CNC3=O)COP(=O)(O)OP(=O)(O)OP(=O)(O)O)O"
      - annotation: !!omap
          - bigg.metabolite: "ditp"
          - chebi: "CHEBI:61382"
          - kegg.compound: "C01345"
          - metanetx.chemical: "MNXM1325"
          - sbo: "SBO:0000247"
      - deltaG: -593.58
    - !!omap
      - id: "s_0640"
      - name: "docosaprenyl diphosphate"
      - compartment: "lp"
      - formula: "C110H177O7P2"
      - charge: -3
      - smiles: "CC(=CCCC(=CCCC(=CCCC(=CCCC(=CCCC(=CCCC(=CCCC(=CCCC(=CCCC(=CCCC(=CCCC(=CCCC(=CCCC(=CCCC(=CCCC(=CCCC(=CCCC(=CCCC(=CCCC(=CCCC(=CCCC(=CCOP(=O)(O)OP(=O)(O)O)C)C)C)C)C)C)C)C)C)C)C)C)C)C)C)C)C)C)C)C)C)C"
      - annotation: !!omap
          - chebi: "CHEBI:53031"
          - metanetx.chemical: "MNXM51449"
          - sbo: "SBO:0000247"
      - deltaG: 1882.32
    - !!omap
      - id: "s_0641"
      - name: "dodecaprenyl diphosphate"
      - compartment: "lp"
      - formula: "C60H97O7P2"
      - charge: -3
      - smiles: "CC(=CCCC(=CCCC(=CCCC(=CCCC(=CCCC(=CCCC(=CCCC(=CCCC(=CCCC(=CCCC(=CCCC(=CCOP(=O)(O)OP(=O)(O)O)C)C)C)C)C)C)C)C)C)C)C)C"
      - annotation: !!omap
          - chebi: "CHEBI:53041"
          - kegg.compound: "C05847"
          - metanetx.chemical: "MNXM11433"
          - sbo: "SBO:0000247"
      - deltaG: 822.47
    - !!omap
      - id: "s_0642"
      - name: "dolichol"
      - compartment: "c"
      - formula: "C20H36O"
      - charge: 0
      - annotation: !!omap
          - bigg.metabolite: "dolichol"
          - chebi: "CHEBI:16091"
          - kegg.compound: "C00381"
          - metanetx.chemical: "MNXM797"
          - sbo: "SBO:0000247"
      - deltaG: 500.3
    - !!omap
      - id: "s_0644"
      - name: "dolichyl D-mannosyl phosphate"
      - compartment: "er"
      - formula: "C26H47O9P"
      - charge: 0
      - smiles: "CC(CCC=C(C)CCC=C(C)CCC=C(C)CCC=C(C)C)CCOP(=O)(O)OC1C(C(C(C(O1)CO)O)O)O"
      - annotation: !!omap
          - bigg.metabolite: "dolmanp"
          - chebi: "CHEBI:15809"
          - kegg.compound: "C03862"
          - metanetx.chemical: "MNXM296"
          - sbo: "SBO:0000247"
      - deltaG: 232.67
    - !!omap
      - id: "s_0645"
      - name: "dolichyl phosphate"
      - compartment: "c"
      - formula: "C20H37O4P"
      - charge: 0
      - smiles: "CC(CCC=C(C)CCC=C(C)CCC=C(C)CCC=C(C)C)CCOP(=O)(O)O"
      - annotation: !!omap
          - bigg.metabolite: "dolp"
          - chebi: "CHEBI:16214"
          - kegg.compound: "C00110"
          - metanetx.chemical: "MNXM278"
          - sbo: "SBO:0000247"
      - deltaG: 277.19
    - !!omap
      - id: "s_0646"
      - name: "dolichyl phosphate"
      - compartment: "er"
      - formula: "C20H37O4P"
      - charge: 0
      - smiles: "CC(CCC=C(C)CCC=C(C)CCC=C(C)CCC=C(C)C)CCOP(=O)(O)O"
      - annotation: !!omap
          - bigg.metabolite: "dolp"
          - chebi: "CHEBI:16214"
          - kegg.compound: "C00110"
          - metanetx.chemical: "MNXM278"
          - sbo: "SBO:0000247"
      - deltaG: 277.19
    - !!omap
      - id: "s_0647"
      - name: "dTDP"
      - compartment: "c"
      - formula: "C10H13N2O11P2"
      - charge: -3
      - smiles: "CC1=CN(C(=O)NC1=O)C2CC(C(O2)COP(=O)(O)OP(=O)(O)O)O"
      - annotation: !!omap
          - bigg.metabolite: "dtdp"
          - chebi: "CHEBI:58369"
          - kegg.compound: "C00363"
          - metanetx.chemical: "MNXM152"
          - sbo: "SBO:0000247"
      - deltaG: -431.57
    - !!omap
      - id: "s_0649"
      - name: "dTMP"
      - compartment: "c"
      - formula: "C10H13N2O8P"
      - charge: -2
      - smiles: "CC1=CN(C(=O)NC1=O)C2CC(C(O2)COP(=O)(O)O)O"
      - annotation: !!omap
          - bigg.metabolite: "dtmp"
          - chebi: "CHEBI:17013"
          - kegg.compound: "C00364"
          - metanetx.chemical: "MNXM257"
          - sbo: "SBO:0000247"
      - deltaG: -232.56
    - !!omap
      - id: "s_0650"
      - name: "dTTP"
      - compartment: "c"
      - formula: "C10H13N2O14P3"
      - charge: -4
      - smiles: "CC1=CN(C(=O)NC1=O)C2CC(C(O2)COP(=O)(O)OP(=O)(O)OP(=O)(O)O)O"
      - annotation: !!omap
          - bigg.metabolite: "dttp"
          - chebi: "CHEBI:58370"
          - kegg.compound: "C00459"
          - metanetx.chemical: "MNXM394"
          - sbo: "SBO:0000247"
      - deltaG: -640.85
    - !!omap
      - id: "s_0651"
      - name: "dTTP"
      - compartment: "e"
      - formula: "C10H13N2O14P3"
      - charge: -4
      - smiles: "CC1=CN(C(=O)NC1=O)C2CC(C(O2)COP(=O)(O)OP(=O)(O)OP(=O)(O)O)O"
      - annotation: !!omap
          - bigg.metabolite: "dttp"
          - chebi: "CHEBI:58370"
          - kegg.compound: "C00459"
          - metanetx.chemical: "MNXM394"
          - sbo: "SBO:0000247"
      - deltaG: -640.85
    - !!omap
      - id: "s_0652"
      - name: "dUDP"
      - compartment: "c"
      - formula: "C9H11N2O11P2"
      - charge: -3
      - smiles: "C1C(C(OC1N2C=CC(=O)NC2=O)COP(=O)(O)OP(=O)(O)O)O"
      - annotation: !!omap
          - bigg.metabolite: "dudp"
          - chebi: "CHEBI:60471"
          - kegg.compound: "C01346"
          - metanetx.chemical: "MNXM572"
          - sbo: "SBO:0000247"
      - deltaG: -450.64
    - !!omap
      - id: "s_0653"
      - name: "dUDP"
      - compartment: "n"
      - formula: "C9H11N2O11P2"
      - charge: -3
      - smiles: "C1C(C(OC1N2C=CC(=O)NC2=O)COP(=O)(O)OP(=O)(O)O)O"
      - annotation: !!omap
          - bigg.metabolite: "dudp"
          - chebi: "CHEBI:60471"
          - kegg.compound: "C01346"
          - metanetx.chemical: "MNXM572"
          - sbo: "SBO:0000247"
      - deltaG: -450.64
    - !!omap
      - id: "s_0654"
      - name: "dUMP"
      - compartment: "c"
      - formula: "C9H11N2O8P"
      - charge: -2
      - smiles: "C1C(C(OC1N2C=CC(=O)NC2=O)COP(=O)(O)O)O"
      - annotation: !!omap
          - bigg.metabolite: "dump"
          - chebi: "CHEBI:246422"
          - kegg.compound: "C00365"
          - metanetx.chemical: "MNXM234"
          - sbo: "SBO:0000247"
      - deltaG: -251.63
    - !!omap
      - id: "s_0655"
      - name: "dUMP"
      - compartment: "n"
      - formula: "C9H11N2O8P"
      - charge: -2
      - smiles: "C1C(C(OC1N2C=CC(=O)NC2=O)COP(=O)(O)O)O"
      - annotation: !!omap
          - bigg.metabolite: "dump"
          - chebi: "CHEBI:246422"
          - kegg.compound: "C00365"
          - metanetx.chemical: "MNXM234"
          - sbo: "SBO:0000247"
      - deltaG: -251.63
    - !!omap
      - id: "s_0656"
      - name: "dUTP"
      - compartment: "c"
      - formula: "C9H11N2O14P3"
      - charge: -4
      - smiles: "C1C(C(OC1N2C=CC(=O)NC2=O)COP(=O)(O)OP(=O)(O)OP(=O)(O)O)O"
      - annotation: !!omap
          - bigg.metabolite: "dutp"
          - chebi: "CHEBI:58212"
          - kegg.compound: "C00460"
          - metanetx.chemical: "MNXM452"
          - sbo: "SBO:0000247"
      - deltaG: -659.91
    - !!omap
      - id: "s_0657"
      - name: "episterol"
      - compartment: "c"
      - formula: "C28H46O"
      - charge: 0
      - smiles: "CC(C)C(=C)CCC(C)C1CCC2C1(CCC3C2=CCC4C3(CCC(C4)O)C)C"
      - annotation: !!omap
          - bigg.metabolite: "epist"
          - chebi: "CHEBI:23929"
          - kegg.compound: "C15777"
          - metanetx.chemical: "MNXM52365"
          - sbo: "SBO:0000247"
      - deltaG: 484.42
    - !!omap
      - id: "s_0659"
      - name: "episterol"
      - compartment: "e"
      - formula: "C28H46O"
      - charge: 0
      - smiles: "CC(C)C(=C)CCC(C)C1CCC2C1(CCC3C2=CCC4C3(CCC(C4)O)C)C"
      - annotation: !!omap
          - bigg.metabolite: "epist"
          - chebi: "CHEBI:23929"
          - kegg.compound: "C15777"
          - metanetx.chemical: "MNXM52365"
          - sbo: "SBO:0000247"
      - deltaG: 484.42
    - !!omap
      - id: "s_0662"
      - name: "ergosta-5,7,22,24(28)-tetraen-3beta-ol"
      - compartment: "c"
      - formula: "C28H42O"
      - charge: 0
      - smiles: "CC(C)C(=C)C=CC(C)C1CCC2C1(CCC3C2=CC=C4C3(CCC(C4)O)C)C"
      - annotation: !!omap
          - bigg.metabolite: "ergtetrol"
          - chebi: "CHEBI:18249"
          - kegg.compound: "C05440"
          - metanetx.chemical: "MNXM1109"
          - sbo: "SBO:0000247"
      - deltaG: 477.57
    - !!omap
      - id: "s_0663"
      - name: "ergosta-5,7,22,24(28)-tetraen-3beta-ol"
      - compartment: "er"
      - formula: "C28H42O"
      - charge: 0
      - smiles: "CC(C)C(=C)C=CC(C)C1CCC2C1(CCC3C2=CC=C4C3(CCC(C4)O)C)C"
      - annotation: !!omap
          - bigg.metabolite: "ergtetrol"
          - chebi: "CHEBI:18249"
          - kegg.compound: "C05440"
          - metanetx.chemical: "MNXM1109"
          - sbo: "SBO:0000247"
      - deltaG: 477.57
    - !!omap
      - id: "s_0664"
      - name: "ergosta-5,7,24(28)-trien-3beta-ol"
      - compartment: "c"
      - formula: "C28H44O"
      - charge: 0
      - smiles: "CC(C)C(=C)CCC(C)C1CCC2C1(CCC3C2=CC=C4C3(CCC(C4)O)C)C"
      - annotation: !!omap
          - bigg.metabolite: "ergtrol"
          - chebi: "CHEBI:52972"
          - kegg.compound: "C15780"
          - metanetx.chemical: "MNXM5738"
          - sbo: "SBO:0000247"
      - deltaG: 479.5
    - !!omap
      - id: "s_0665"
      - name: "ergosterol"
      - compartment: "ce"
      - formula: "C28H44O"
      - charge: 0
      - smiles: "CC(C)C(C)C=CC(C)C1CCC2C1(CCC3C2=CC=C4C3(CCC(C4)O)C)C"
      - annotation: !!omap
          - bigg.metabolite: "ergst"
          - chebi: "CHEBI:16933"
          - kegg.compound: "C01694"
          - metanetx.chemical: "MNXM922"
          - sbo: "SBO:0000247"
      - deltaG: 472.21
    - !!omap
      - id: "s_0666"
      - name: "ergosterol"
      - compartment: "c"
      - formula: "C28H44O"
      - charge: 0
      - smiles: "CC(C)C(C)C=CC(C)C1CCC2C1(CCC3C2=CC=C4C3(CCC(C4)O)C)C"
      - annotation: !!omap
          - bigg.metabolite: "ergst"
          - chebi: "CHEBI:16933"
          - kegg.compound: "C01694"
          - metanetx.chemical: "MNXM922"
          - sbo: "SBO:0000247"
      - deltaG: 472.21
    - !!omap
      - id: "s_0667"
      - name: "ergosterol"
      - compartment: "er"
      - formula: "C28H44O"
      - charge: 0
      - smiles: "CC(C)C(C)C=CC(C)C1CCC2C1(CCC3C2=CC=C4C3(CCC(C4)O)C)C"
      - annotation: !!omap
          - bigg.metabolite: "ergst"
          - chebi: "CHEBI:16933"
          - kegg.compound: "C01694"
          - metanetx.chemical: "MNXM922"
          - sbo: "SBO:0000247"
      - deltaG: 472.21
    - !!omap
      - id: "s_0668"
      - name: "ergosterol"
      - compartment: "e"
      - formula: "C28H44O"
      - charge: 0
      - smiles: "CC(C)C(C)C=CC(C)C1CCC2C1(CCC3C2=CC=C4C3(CCC(C4)O)C)C"
      - annotation: !!omap
          - bigg.metabolite: "ergst"
          - chebi: "CHEBI:16933"
          - kegg.compound: "C01694"
          - metanetx.chemical: "MNXM922"
          - sbo: "SBO:0000247"
      - deltaG: 472.21
    - !!omap
      - id: "s_0669"
      - name: "ergosterol"
      - compartment: "lp"
      - formula: "C28H44O"
      - charge: 0
      - smiles: "CC(C)C(C)C=CC(C)C1CCC2C1(CCC3C2=CC=C4C3(CCC(C4)O)C)C"
      - annotation: !!omap
          - bigg.metabolite: "ergst"
          - chebi: "CHEBI:16933"
          - kegg.compound: "C01694"
          - metanetx.chemical: "MNXM922"
          - sbo: "SBO:0000247"
      - deltaG: 472.21
    - !!omap
      - id: "s_0670"
      - name: "ergosterol 3-beta-D-glucoside"
      - compartment: "c"
      - formula: "C34H54O6"
      - charge: 0
      - annotation: !!omap
          - bigg.metabolite: "ergst3glc"
          - chebi: "CHEBI:52973"
          - metanetx.chemical: "MNXM52465"
          - sbo: "SBO:0000247"
      - deltaG: 419.61
    - !!omap
      - id: "s_0672"
      - name: "ergosterol ester backbone"
      - compartment: "c"
      - formula: "C28H42"
      - charge: 0
      - annotation: !!omap
          - chebi: "CHEBI:52320"
          - metanetx.chemical: "MNXM5739"
          - sbo: "SBO:0000649"
      - deltaG: 10000000
    - !!omap
      - id: "s_0677"
      - name: "erythro-4-hydroxy-L-glutamic acid"
      - compartment: "c"
      - formula: "C5H8NO5"
      - charge: -1
      - smiles: "C(C(C(=O)O)N)C(C(=O)O)O"
      - annotation: !!omap
          - bigg.metabolite: "e4hglu"
          - chebi: "CHEBI:21285"
          - kegg.compound: "C05947"
          - metanetx.chemical: "MNXM923"
          - sbo: "SBO:0000247"
      - deltaG: -129.86
    - !!omap
      - id: "s_0678"
      - name: "erythro-4-hydroxy-L-glutamic acid"
      - compartment: "m"
      - formula: "C5H8NO5"
      - charge: -1
      - smiles: "C(C(C(=O)O)N)C(C(=O)O)O"
      - annotation: !!omap
          - bigg.metabolite: "e4hglu"
          - chebi: "CHEBI:21285"
          - kegg.compound: "C05947"
          - metanetx.chemical: "MNXM923"
          - sbo: "SBO:0000247"
      - deltaG: -129.86
    - !!omap
      - id: "s_0679"
      - name: "erythro-4-hydroxy-L-glutamic acid"
      - compartment: "p"
      - formula: "C5H8NO5"
      - charge: -1
      - smiles: "C(C(C(=O)O)N)C(C(=O)O)O"
      - annotation: !!omap
          - bigg.metabolite: "e4hglu"
          - chebi: "CHEBI:21285"
          - kegg.compound: "C05947"
          - metanetx.chemical: "MNXM923"
          - sbo: "SBO:0000247"
      - deltaG: -129.86
    - !!omap
      - id: "s_0680"
      - name: "ethanol"
      - compartment: "c"
      - formula: "C2H6O"
      - charge: 0
      - smiles: "CCO"
      - annotation: !!omap
          - bigg.metabolite: "etoh"
          - chebi: "CHEBI:16236"
          - kegg.compound: "C00469"
          - metanetx.chemical: "MNXM303"
          - sbo: "SBO:0000247"
      - deltaG: 14.98
    - !!omap
      - id: "s_0681"
      - name: "ethanol"
      - compartment: "e"
      - formula: "C2H6O"
      - charge: 0
      - smiles: "CCO"
      - annotation: !!omap
          - bigg.metabolite: "etoh"
          - chebi: "CHEBI:16236"
          - kegg.compound: "C00469"
          - metanetx.chemical: "MNXM303"
          - sbo: "SBO:0000247"
      - deltaG: 14.98
    - !!omap
      - id: "s_0682"
      - name: "ethanol"
      - compartment: "m"
      - formula: "C2H6O"
      - charge: 0
      - smiles: "CCO"
      - annotation: !!omap
          - bigg.metabolite: "etoh"
          - chebi: "CHEBI:16236"
          - kegg.compound: "C00469"
          - metanetx.chemical: "MNXM303"
          - sbo: "SBO:0000247"
      - deltaG: 14.98
    - !!omap
      - id: "s_0683"
      - name: "ethanolamine"
      - compartment: "c"
      - formula: "C2H8NO"
      - charge: 1
      - smiles: "C(CO)N"
      - annotation: !!omap
          - bigg.metabolite: "etha"
          - chebi: "CHEBI:57603"
          - kegg.compound: "C00189"
          - metanetx.chemical: "MNXM218"
          - sbo: "SBO:0000247"
      - deltaG: 33.19
    - !!omap
      - id: "s_0684"
      - name: "ethanolamine"
      - compartment: "e"
      - formula: "C2H8NO"
      - charge: 1
      - smiles: "C(CO)N"
      - annotation: !!omap
          - bigg.metabolite: "etha"
          - chebi: "CHEBI:57603"
          - kegg.compound: "C00189"
          - metanetx.chemical: "MNXM218"
          - sbo: "SBO:0000247"
      - deltaG: 33.19
    - !!omap
      - id: "s_0685"
      - name: "ethyl acetate"
      - compartment: "c"
      - formula: "C4H8O2"
      - charge: 0
      - smiles: "CCOC(=O)C"
      - annotation: !!omap
          - chebi: "CHEBI:27750"
          - kegg.compound: "C00849"
          - metanetx.chemical: "MNXM8595"
          - sbo: "SBO:0000247"
      - deltaG: -3.09
    - !!omap
      - id: "s_0686"
      - name: "ethyl acetate"
      - compartment: "e"
      - formula: "C4H8O2"
      - charge: 0
      - smiles: "CCOC(=O)C"
      - annotation: !!omap
          - chebi: "CHEBI:27750"
          - kegg.compound: "C00849"
          - metanetx.chemical: "MNXM8595"
          - sbo: "SBO:0000247"
      - deltaG: -3.09
    - !!omap
      - id: "s_0687"
      - name: "FAD"
      - compartment: "c"
      - formula: "C27H30N9O15P2"
      - charge: -3
      - smiles: "CC1=CC2=C(C=C1C)N(C3=NC(=O)NC(=O)C3=N2)CC(C(C(COP(=O)(O)OP(=O)(O)OCC4C(C(C(O4)N5C=NC6=C(N=CN=C65)N)O)O)O)O)O"
      - annotation: !!omap
          - bigg.metabolite: "fad"
          - chebi: "CHEBI:57692"
          - kegg.compound: "C00016"
          - metanetx.chemical: "MNXM33"
          - sbo: "SBO:0000247"
      - deltaG: 10000000
    - !!omap
      - id: "s_0688"
      - name: "FAD"
      - compartment: "m"
      - formula: "C27H30N9O15P2"
      - charge: -3
      - smiles: "CC1=CC2=C(C=C1C)N(C3=NC(=O)NC(=O)C3=N2)CC(C(C(COP(=O)(O)OP(=O)(O)OCC4C(C(C(O4)N5C=NC6=C(N=CN=C65)N)O)O)O)O)O"
      - annotation: !!omap
          - bigg.metabolite: "fad"
          - chebi: "CHEBI:57692"
          - kegg.compound: "C00016"
          - metanetx.chemical: "MNXM33"
          - sbo: "SBO:0000247"
      - deltaG: 10000000
    - !!omap
      - id: "s_0689"
      - name: "FADH2"
      - compartment: "c"
      - formula: "C27H33N9O15P2"
      - charge: -2
      - smiles: "CC1=CC2=C(C=C1C)N(C3=C(N2)C(=O)NC(=O)N3)CC(C(C(COP(=O)(O)OP(=O)(O)OCC4C(C(C(O4)N5C=NC6=C(N=CN=C65)N)O)O)O)O)O"
      - annotation: !!omap
          - bigg.metabolite: "fadh2"
          - chebi: "CHEBI:58307"
          - kegg.compound: "C01352"
          - metanetx.chemical: "MNXM38"
          - sbo: "SBO:0000247"
      - deltaG: -233.44
    - !!omap
      - id: "s_0690"
      - name: "FADH2"
      - compartment: "m"
      - formula: "C27H33N9O15P2"
      - charge: -2
      - smiles: "CC1=CC2=C(C=C1C)N(C3=C(N2)C(=O)NC(=O)N3)CC(C(C(COP(=O)(O)OP(=O)(O)OCC4C(C(C(O4)N5C=NC6=C(N=CN=C65)N)O)O)O)O)O"
      - annotation: !!omap
          - bigg.metabolite: "fadh2"
          - chebi: "CHEBI:58307"
          - kegg.compound: "C01352"
          - metanetx.chemical: "MNXM38"
          - sbo: "SBO:0000247"
      - deltaG: -233.44
    - !!omap
      - id: "s_0692"
      - name: "farnesyl diphosphate"
      - compartment: "lp"
      - formula: "C15H25O7P2"
      - charge: -3
      - smiles: "CC(=CCCC(=CCCC(=CCOP(=O)(O)OP(=O)(O)O)C)C)C"
      - annotation: !!omap
          - bigg.metabolite: "frdp"
          - chebi: "CHEBI:175763"
          - kegg.compound: "C00448"
          - metanetx.chemical: "MNXM34"
          - sbo: "SBO:0000247"
      - deltaG: -131.39
    - !!omap
      - id: "s_0694"
      - name: "fatty acid backbone"
      - compartment: "c"
      - charge: 0
      - annotation: !!omap
          - chebi: "CHEBI:35366"
          - kegg.compound: "C00162"
          - metanetx.chemical: "MNXM72"
          - sbo: "SBO:0000649"
      - deltaG: 10000000
    - !!omap
      - id: "s_0699"
      - name: "fecosterol"
      - compartment: "ce"
      - formula: "C28H46O"
      - charge: 0
      - smiles: "CC(C)C(=C)CCC(C)C1CCC2C1(CCC3=C2CCC4C3(CCC(C4)O)C)C"
      - annotation: !!omap
          - bigg.metabolite: "fecost"
          - chebi: "CHEBI:17038"
          - kegg.compound: "C04525"
          - metanetx.chemical: "MNXM1741"
          - sbo: "SBO:0000247"
      - deltaG: 484.33
    - !!omap
      - id: "s_0700"
      - name: "fecosterol"
      - compartment: "c"
      - formula: "C28H46O"
      - charge: 0
      - smiles: "CC(C)C(=C)CCC(C)C1CCC2C1(CCC3=C2CCC4C3(CCC(C4)O)C)C"
      - annotation: !!omap
          - bigg.metabolite: "fecost"
          - chebi: "CHEBI:17038"
          - kegg.compound: "C04525"
          - metanetx.chemical: "MNXM1741"
          - sbo: "SBO:0000247"
      - deltaG: 484.33
    - !!omap
      - id: "s_0702"
      - name: "fecosterol"
      - compartment: "e"
      - formula: "C28H46O"
      - charge: 0
      - smiles: "CC(C)C(=C)CCC(C)C1CCC2C1(CCC3=C2CCC4C3(CCC(C4)O)C)C"
      - annotation: !!omap
          - bigg.metabolite: "fecost"
          - chebi: "CHEBI:17038"
          - kegg.compound: "C04525"
          - metanetx.chemical: "MNXM1741"
          - sbo: "SBO:0000247"
      - deltaG: 484.33
    - !!omap
      - id: "s_0703"
      - name: "fecosterol"
      - compartment: "lp"
      - formula: "C28H46O"
      - charge: 0
      - smiles: "CC(C)C(=C)CCC(C)C1CCC2C1(CCC3=C2CCC4C3(CCC(C4)O)C)C"
      - annotation: !!omap
          - bigg.metabolite: "fecost"
          - chebi: "CHEBI:17038"
          - kegg.compound: "C04525"
          - metanetx.chemical: "MNXM1741"
          - sbo: "SBO:0000247"
      - deltaG: 484.33
    - !!omap
      - id: "s_0709"
      - name: "ferricytochrome c"
      - compartment: "m"
      - formula: "C42H44FeN8O8S2R4"
      - charge: 0
      - smiles: "CC1=C(C2=CC3=NC(=CC4=C(C(=C([N-]4)C=C5C(=C(C(=N5)C=C1[N-]2)C(C)SCC(C(=O)NC)N)C)C(C)SCC(C(=O)NC)N)C)C(=C3CCC(=O)O)C)CCC(=O)O.[Fe+2]"
      - annotation: !!omap
          - bigg.metabolite: "ficytc"
          - chebi: "CHEBI:15991"
          - kegg.compound: "C00125"
          - metanetx.chemical: "MNXM5749"
          - sbo: "SBO:0000247"
      - deltaG: 10000000
    - !!omap
      - id: "s_0710"
      - name: "ferrocytochrome c"
      - compartment: "m"
      - formula: "C42H45FeN8O8S2R4"
      - charge: 0
      - smiles: "CC1=C(C2=CC3=NC(=CC4=C(C(=C([N-]4)C=C5C(=C(C(=N5)C=C1[N-]2)C(C)SCC(C(=O)NC)N)C)C(C)SCC(C(=O)NC)N)C)C(=C3CCC(=O)O)C)CCC(=O)O.[Fe+2]"
      - annotation: !!omap
          - bigg.metabolite: "focytc"
          - chebi: "CHEBI:16928"
          - kegg.compound: "C00126"
          - metanetx.chemical: "MNXM746"
          - sbo: "SBO:0000247"
      - deltaG: 10000000
    - !!omap
      - id: "s_0712"
      - name: "ferroheme b"
      - compartment: "m"
      - formula: "C34H30FeN4O4"
      - charge: -2
      - smiles: "CC1=C(C2=CC3=NC(=CC4=C(C(=C([N-]4)C=C5C(=C(C(=N5)C=C1[N-]2)C=C)C)C=C)C)C(=C3CCC(=O)O)C)CCC(=O)O.[Fe]"
      - annotation: !!omap
          - bigg.metabolite: "pheme"
          - chebi: "CHEBI:60344"
          - kegg.compound: "C00032"
          - metanetx.chemical: "MNXM249"
          - sbo: "SBO:0000247"
      - deltaG: 10000000
    - !!omap
      - id: "s_0713"
      - name: "fMet-tRNA(fMet)"
      - compartment: "m"
      - formula: "C6H10NO2SR"
      - charge: 0
      - annotation: !!omap
          - bigg.metabolite: "fmettrna"
          - chebi: "CHEBI:17119"
          - kegg.compound: "C03294"
          - metanetx.chemical: "MNXM95381"
          - sbo: "SBO:0000247"
      - deltaG: 10000000
    - !!omap
      - id: "s_0714"
      - name: "FMN"
      - compartment: "c"
      - formula: "C17H18N4O9P"
      - charge: -3
      - smiles: "CC1=CC2=C(C=C1C)N(C3=NC(=O)NC(=O)C3=N2)CC(C(C(COP(=O)(O)O)O)O)O"
      - annotation: !!omap
          - bigg.metabolite: "fmn"
          - chebi: "CHEBI:58210"
          - kegg.compound: "C00061"
          - metanetx.chemical: "MNXM119"
          - sbo: "SBO:0000247"
      - deltaG: 10000000
    - !!omap
      - id: "s_0715"
      - name: "FMN"
      - compartment: "e"
      - formula: "C17H18N4O9P"
      - charge: -3
      - smiles: "CC1=CC2=C(C=C1C)N(C3=NC(=O)NC(=O)C3=N2)CC(C(C(COP(=O)(O)O)O)O)O"
      - annotation: !!omap
          - bigg.metabolite: "fmn"
          - chebi: "CHEBI:58210"
          - kegg.compound: "C00061"
          - metanetx.chemical: "MNXM119"
          - sbo: "SBO:0000247"
      - deltaG: 10000000
    - !!omap
      - id: "s_0716"
      - name: "FMN"
      - compartment: "m"
      - formula: "C17H18N4O9P"
      - charge: -3
      - smiles: "CC1=CC2=C(C=C1C)N(C3=NC(=O)NC(=O)C3=N2)CC(C(C(COP(=O)(O)O)O)O)O"
      - annotation: !!omap
          - bigg.metabolite: "fmn"
          - chebi: "CHEBI:58210"
          - kegg.compound: "C00061"
          - metanetx.chemical: "MNXM119"
          - sbo: "SBO:0000247"
      - deltaG: 10000000
    - !!omap
      - id: "s_0717"
      - name: "FMNH2"
      - compartment: "c"
      - formula: "C17H21N4O9P"
      - charge: -2
      - smiles: "CC1=CC2=C(C=C1C)N(C3=C(N2)C(=O)NC(=O)N3)CC(C(C(COP(=O)(O)O)O)O)O"
      - annotation: !!omap
          - bigg.metabolite: "fmnh2"
          - chebi: "CHEBI:57618"
          - kegg.compound: "C01847"
          - metanetx.chemical: "MNXM208"
          - sbo: "SBO:0000247"
      - deltaG: -142.78
    - !!omap
      - id: "s_0719"
      - name: "folate"
      - compartment: "c"
      - formula: "C19H17N7O6"
      - charge: -2
      - smiles: "C1=CC(=CC=C1C(=O)NC(CCC(=O)O)C(=O)O)NCC2=CN=C3C(=N2)C(=O)NC(=N3)N"
      - annotation: !!omap
          - bigg.metabolite: "fol"
          - chebi: "CHEBI:62501"
          - kegg.compound: "C00504"
          - metanetx.chemical: "MNXM617"
          - sbo: "SBO:0000247"
      - deltaG: 69.78
    - !!omap
      - id: "s_0720"
      - name: "folate"
      - compartment: "e"
      - formula: "C19H17N7O6"
      - charge: -2
      - smiles: "C1=CC(=CC=C1C(=O)NC(CCC(=O)O)C(=O)O)NCC2=CN=C3C(=N2)C(=O)NC(=N3)N"
      - annotation: !!omap
          - bigg.metabolite: "fol"
          - chebi: "CHEBI:62501"
          - kegg.compound: "C00504"
          - metanetx.chemical: "MNXM617"
          - sbo: "SBO:0000247"
      - deltaG: 69.78
    - !!omap
      - id: "s_0721"
      - name: "formaldehyde"
      - compartment: "c"
      - formula: "CH2O"
      - charge: 0
      - smiles: "C=O"
      - annotation: !!omap
          - bigg.metabolite: "fald"
          - chebi: "CHEBI:16842"
          - kegg.compound: "C00067"
          - metanetx.chemical: "MNXM56"
          - sbo: "SBO:0000247"
      - deltaG: -11.73
    - !!omap
      - id: "s_0722"
      - name: "formate"
      - compartment: "c"
      - formula: "CHO2"
      - charge: -1
      - smiles: "C(=O)[O-]"
      - annotation: !!omap
          - bigg.metabolite: "for"
          - chebi: "CHEBI:15740"
          - kegg.compound: "C00058"
          - metanetx.chemical: "MNXM39"
          - sbo: "SBO:0000247"
      - deltaG: -74.36
    - !!omap
      - id: "s_0723"
      - name: "formate"
      - compartment: "e"
      - formula: "CHO2"
      - charge: -1
      - smiles: "C(=O)[O-]"
      - annotation: !!omap
          - bigg.metabolite: "for"
          - chebi: "CHEBI:15740"
          - kegg.compound: "C00058"
          - metanetx.chemical: "MNXM39"
          - sbo: "SBO:0000247"
      - deltaG: -74.36
    - !!omap
      - id: "s_0724"
      - name: "formate"
      - compartment: "m"
      - formula: "CHO2"
      - charge: -1
      - smiles: "C(=O)[O-]"
      - annotation: !!omap
          - bigg.metabolite: "for"
          - chebi: "CHEBI:15740"
          - kegg.compound: "C00058"
          - metanetx.chemical: "MNXM39"
          - sbo: "SBO:0000247"
      - deltaG: -74.36
    - !!omap
      - id: "s_0725"
      - name: "fumarate"
      - compartment: "c"
      - formula: "C4H2O4"
      - charge: -2
      - smiles: "C(=CC(=O)[O-])C(=O)[O-]"
      - annotation: !!omap
          - bigg.metabolite: "fum"
          - chebi: "CHEBI:29806"
          - kegg.compound: "C00122"
          - metanetx.chemical: "MNXM93"
          - sbo: "SBO:0000247"
      - deltaG: -125
    - !!omap
      - id: "s_0726"
      - name: "fumarate"
      - compartment: "e"
      - formula: "C4H2O4"
      - charge: -2
      - smiles: "C(=CC(=O)[O-])C(=O)[O-]"
      - annotation: !!omap
          - bigg.metabolite: "fum"
          - chebi: "CHEBI:29806"
          - kegg.compound: "C00122"
          - metanetx.chemical: "MNXM93"
          - sbo: "SBO:0000247"
      - deltaG: -125
    - !!omap
      - id: "s_0727"
      - name: "fumarate"
      - compartment: "m"
      - formula: "C4H2O4"
      - charge: -2
      - smiles: "C(=CC(=O)[O-])C(=O)[O-]"
      - annotation: !!omap
          - bigg.metabolite: "fum"
          - chebi: "CHEBI:29806"
          - kegg.compound: "C00122"
          - metanetx.chemical: "MNXM93"
          - sbo: "SBO:0000247"
      - deltaG: -125
    - !!omap
      - id: "s_0734"
      - name: "gamma-aminobutyrate"
      - compartment: "c"
      - formula: "C4H9NO2"
      - charge: 0
      - smiles: "C(CC(=O)O)CN"
      - annotation: !!omap
          - bigg.metabolite: "4abut"
          - chebi: "CHEBI:16865"
          - kegg.compound: "C00334"
          - metanetx.chemical: "MNXM192"
          - sbo: "SBO:0000247"
      - deltaG: -0.61
    - !!omap
      - id: "s_0736"
      - name: "gamma-aminobutyrate"
      - compartment: "e"
      - formula: "C4H9NO2"
      - charge: 0
      - smiles: "C(CC(=O)O)CN"
      - annotation: !!omap
          - bigg.metabolite: "4abut"
          - chebi: "CHEBI:16865"
          - kegg.compound: "C00334"
          - metanetx.chemical: "MNXM192"
          - sbo: "SBO:0000247"
      - deltaG: -0.61
    - !!omap
      - id: "s_0738"
      - name: "gamma-aminobutyrate"
      - compartment: "m"
      - formula: "C4H9NO2"
      - charge: 0
      - smiles: "C(CC(=O)O)CN"
      - annotation: !!omap
          - bigg.metabolite: "4abut"
          - chebi: "CHEBI:16865"
          - kegg.compound: "C00334"
          - metanetx.chemical: "MNXM192"
          - sbo: "SBO:0000247"
      - deltaG: -0.61
    - !!omap
      - id: "s_0739"
      - name: "GDP"
      - compartment: "c"
      - formula: "C10H12N5O11P2"
      - charge: -3
      - smiles: "C1=NC2=C(N1C3C(C(C(O3)COP(=O)(O)OP(=O)(O)O)O)O)N=C(NC2=O)N"
      - annotation: !!omap
          - bigg.metabolite: "gdp"
          - chebi: "CHEBI:58189"
          - kegg.compound: "C00035"
          - metanetx.chemical: "MNXM30"
          - sbo: "SBO:0000247"
      - deltaG: -391.27
    - !!omap
      - id: "s_0740"
      - name: "GDP"
      - compartment: "g"
      - formula: "C10H12N5O11P2"
      - charge: -3
      - smiles: "C1=NC2=C(N1C3C(C(C(O3)COP(=O)(O)OP(=O)(O)O)O)O)N=C(NC2=O)N"
      - annotation: !!omap
          - bigg.metabolite: "gdp"
          - chebi: "CHEBI:58189"
          - kegg.compound: "C00035"
          - metanetx.chemical: "MNXM30"
          - sbo: "SBO:0000247"
      - deltaG: -391.27
    - !!omap
      - id: "s_0741"
      - name: "GDP"
      - compartment: "m"
      - formula: "C10H12N5O11P2"
      - charge: -3
      - smiles: "C1=NC2=C(N1C3C(C(C(O3)COP(=O)(O)OP(=O)(O)O)O)O)N=C(NC2=O)N"
      - annotation: !!omap
          - bigg.metabolite: "gdp"
          - chebi: "CHEBI:58189"
          - kegg.compound: "C00035"
          - metanetx.chemical: "MNXM30"
          - sbo: "SBO:0000247"
      - deltaG: -391.27
    - !!omap
      - id: "s_0742"
      - name: "GDP"
      - compartment: "n"
      - formula: "C10H12N5O11P2"
      - charge: -3
      - smiles: "C1=NC2=C(N1C3C(C(C(O3)COP(=O)(O)OP(=O)(O)O)O)O)N=C(NC2=O)N"
      - annotation: !!omap
          - bigg.metabolite: "gdp"
          - chebi: "CHEBI:58189"
          - kegg.compound: "C00035"
          - metanetx.chemical: "MNXM30"
          - sbo: "SBO:0000247"
      - deltaG: -391.27
    - !!omap
      - id: "s_0743"
      - name: "GDP-alpha-D-mannose"
      - compartment: "c"
      - formula: "C16H23N5O16P2"
      - charge: -2
      - smiles: "C1=NC2=C(N1C3C(C(C(O3)COP(=O)(O)OP(=O)(O)OC4C(C(C(C(O4)CO)O)O)O)O)O)N=C(NC2=O)N"
      - annotation: !!omap
          - bigg.metabolite: "gdpmann"
          - chebi: "CHEBI:57527"
          - kegg.compound: "C00096"
          - metanetx.chemical: "MNXM82"
          - sbo: "SBO:0000247"
      - deltaG: -453.27
    - !!omap
      - id: "s_0744"
      - name: "GDP-alpha-D-mannose"
      - compartment: "g"
      - formula: "C16H23N5O16P2"
      - charge: -2
      - smiles: "C1=NC2=C(N1C3C(C(C(O3)COP(=O)(O)OP(=O)(O)OC4C(C(C(C(O4)CO)O)O)O)O)O)N=C(NC2=O)N"
      - annotation: !!omap
          - bigg.metabolite: "gdpmann"
          - chebi: "CHEBI:57527"
          - kegg.compound: "C00096"
          - metanetx.chemical: "MNXM82"
          - sbo: "SBO:0000247"
      - deltaG: -453.27
    - !!omap
      - id: "s_0745"
      - name: "geranyl diphosphate"
      - compartment: "c"
      - formula: "C10H17O7P2"
      - charge: -3
      - smiles: "CC(=CCCC(=CCOP(=O)(O)OP(=O)(O)O)C)C"
      - annotation: !!omap
          - bigg.metabolite: "grdp"
          - chebi: "CHEBI:58057"
          - kegg.compound: "C00341"
          - metanetx.chemical: "MNXM100"
          - sbo: "SBO:0000247"
      - deltaG: -237.38
    - !!omap
      - id: "s_0746"
      - name: "geranylgeranyl diphosphate"
      - compartment: "lp"
      - formula: "C20H33O7P2"
      - charge: -3
      - smiles: "CC(=CCCC(=CCCC(=CCCC(=CCOP(=O)(O)OP(=O)(O)O)C)C)C)C"
      - annotation: !!omap
          - bigg.metabolite: "ggdp"
          - chebi: "CHEBI:58756"
          - kegg.compound: "C00353"
          - metanetx.chemical: "MNXM139"
          - sbo: "SBO:0000247"
      - deltaG: -25.41
    - !!omap
      - id: "s_0747"
      - name: "Gln-tRNA(Gln)"
      - compartment: "c"
      - formula: "C5H10N2O2R"
      - charge: 1
      - annotation: !!omap
          - bigg.metabolite: "glntrna"
          - chebi: "CHEBI:29166"
          - kegg.compound: "C02282"
          - metanetx.chemical: "MNXM89810"
          - sbo: "SBO:0000247"
      - deltaG: 10000000
    - !!omap
      - id: "s_0748"
      - name: "Glu-tRNA(Glu)"
      - compartment: "c"
      - formula: "C5H8NO3R"
      - charge: 0
      - annotation: !!omap
          - bigg.metabolite: "glutrna"
          - chebi: "CHEBI:29157"
          - kegg.compound: "C02987"
          - metanetx.chemical: "MNXM89752"
          - sbo: "SBO:0000247"
      - deltaG: 10000000
    - !!omap
      - id: "s_0749"
      - name: "Glu-tRNA(Glu)"
      - compartment: "m"
      - formula: "C5H8NO3R"
      - charge: 0
      - annotation: !!omap
          - bigg.metabolite: "glutrna"
          - chebi: "CHEBI:29157"
          - kegg.compound: "C02987"
          - metanetx.chemical: "MNXM89752"
          - sbo: "SBO:0000247"
      - deltaG: 10000000
    - !!omap
      - id: "s_0750"
      - name: "glutathione"
      - compartment: "c"
      - formula: "C10H16N3O6S"
      - charge: -1
      - smiles: "C(CC(=O)NC(CS)C(=O)NCC(=O)O)C(C(=O)O)N"
      - annotation: !!omap
          - bigg.metabolite: "gthrd"
          - chebi: "CHEBI:57925"
          - kegg.compound: "C00051"
          - metanetx.chemical: "MNXM57"
          - sbo: "SBO:0000247"
      - deltaG: -70.58
    - !!omap
      - id: "s_0751"
      - name: "glutathione"
      - compartment: "e"
      - formula: "C10H16N3O6S"
      - charge: -1
      - smiles: "C(CC(=O)NC(CS)C(=O)NCC(=O)O)C(C(=O)O)N"
      - annotation: !!omap
          - bigg.metabolite: "gthrd"
          - chebi: "CHEBI:57925"
          - kegg.compound: "C00051"
          - metanetx.chemical: "MNXM57"
          - sbo: "SBO:0000247"
      - deltaG: -70.58
    - !!omap
      - id: "s_0752"
      - name: "glutathione"
      - compartment: "m"
      - formula: "C10H16N3O6S"
      - charge: -1
      - smiles: "C(CC(=O)NC(CS)C(=O)NCC(=O)O)C(C(=O)O)N"
      - annotation: !!omap
          - bigg.metabolite: "gthrd"
          - chebi: "CHEBI:57925"
          - kegg.compound: "C00051"
          - metanetx.chemical: "MNXM57"
          - sbo: "SBO:0000247"
      - deltaG: -70.58
    - !!omap
      - id: "s_0753"
      - name: "glutathione"
      - compartment: "v"
      - formula: "C10H16N3O6S"
      - charge: -1
      - smiles: "C(CC(=O)NC(CS)C(=O)NCC(=O)O)C(C(=O)O)N"
      - annotation: !!omap
          - bigg.metabolite: "gthrd"
          - chebi: "CHEBI:57925"
          - kegg.compound: "C00051"
          - metanetx.chemical: "MNXM57"
          - sbo: "SBO:0000247"
      - deltaG: -70.58
    - !!omap
      - id: "s_0754"
      - name: "glutathione disulfide"
      - compartment: "c"
      - formula: "C20H30N6O12S2"
      - charge: -2
      - smiles: "C(CC(=O)NC(CSSCC(C(=O)NCC(=O)O)NC(=O)CCC(C(=O)O)N)C(=O)NCC(=O)O)C(C(=O)O)N"
      - annotation: !!omap
          - bigg.metabolite: "gthox"
          - chebi: "CHEBI:58297"
          - kegg.compound: "C00127"
          - metanetx.chemical: "MNXM151"
          - sbo: "SBO:0000247"
      - deltaG: -155.26
    - !!omap
      - id: "s_0755"
      - name: "glutathione disulfide"
      - compartment: "e"
      - formula: "C20H30N6O12S2"
      - charge: -2
      - smiles: "C(CC(=O)NC(CSSCC(C(=O)NCC(=O)O)NC(=O)CCC(C(=O)O)N)C(=O)NCC(=O)O)C(C(=O)O)N"
      - annotation: !!omap
          - bigg.metabolite: "gthox"
          - chebi: "CHEBI:58297"
          - kegg.compound: "C00127"
          - metanetx.chemical: "MNXM151"
          - sbo: "SBO:0000247"
      - deltaG: -155.26
    - !!omap
      - id: "s_0756"
      - name: "glutathione disulfide"
      - compartment: "m"
      - formula: "C20H30N6O12S2"
      - charge: -2
      - smiles: "C(CC(=O)NC(CSSCC(C(=O)NCC(=O)O)NC(=O)CCC(C(=O)O)N)C(=O)NCC(=O)O)C(C(=O)O)N"
      - annotation: !!omap
          - bigg.metabolite: "gthox"
          - chebi: "CHEBI:58297"
          - kegg.compound: "C00127"
          - metanetx.chemical: "MNXM151"
          - sbo: "SBO:0000247"
      - deltaG: -155.26
    - !!omap
      - id: "s_0757"
      - name: "Gly-tRNA(Gly)"
      - compartment: "c"
      - formula: "C2H5NOR"
      - charge: 1
      - annotation: !!omap
          - bigg.metabolite: "glytrna"
          - chebi: "CHEBI:29156"
          - kegg.compound: "C02412"
          - metanetx.chemical: "MNXM89763"
          - sbo: "SBO:0000247"
      - deltaG: 10000000
    - !!omap
      - id: "s_0764"
      - name: "glyceraldehyde 3-phosphate"
      - compartment: "c"
      - formula: "C3H5O6P"
      - charge: -2
      - smiles: "C(C(C=O)O)OP(=O)(O)O"
      - annotation: !!omap
          - bigg.metabolite: "g3p"
          - chebi: "CHEBI:58027"
          - kegg.compound: "C00661"
          - metanetx.chemical: "MNXM74"
          - sbo: "SBO:0000247"
      - deltaG: -271.33
    - !!omap
      - id: "s_0765"
      - name: "glycerol"
      - compartment: "c"
      - formula: "C3H8O3"
      - charge: 0
      - smiles: "C(C(CO)O)O"
      - annotation: !!omap
          - bigg.metabolite: "glyc"
          - chebi: "CHEBI:17754"
          - kegg.compound: "C00116"
          - metanetx.chemical: "MNXM89612"
          - sbo: "SBO:0000247"
      - deltaG: -38.29
    - !!omap
      - id: "s_0766"
      - name: "glycerol"
      - compartment: "e"
      - formula: "C3H8O3"
      - charge: 0
      - smiles: "C(C(CO)O)O"
      - annotation: !!omap
          - bigg.metabolite: "glyc"
          - chebi: "CHEBI:17754"
          - kegg.compound: "C00116"
          - metanetx.chemical: "MNXM89612"
          - sbo: "SBO:0000247"
      - deltaG: -38.29
    - !!omap
      - id: "s_0767"
      - name: "glycerol 3-phosphate"
      - compartment: "c"
      - formula: "C3H7O6P"
      - charge: -2
      - smiles: "C(C(COP(=O)(O)O)O)O"
      - annotation: !!omap
          - bigg.metabolite: "glyc3p"
          - chebi: "CHEBI:57597"
          - kegg.compound: "C00093"
          - metanetx.chemical: "MNXM66"
          - sbo: "SBO:0000247"
      - deltaG: -261.35
    - !!omap
      - id: "s_0769"
      - name: "glycerol 3-phosphate"
      - compartment: "lp"
      - formula: "C3H7O6P"
      - charge: -2
      - smiles: "C(C(COP(=O)(O)O)O)O"
      - annotation: !!omap
          - bigg.metabolite: "glyc3p"
          - chebi: "CHEBI:57597"
          - kegg.compound: "C00093"
          - metanetx.chemical: "MNXM66"
          - sbo: "SBO:0000247"
      - deltaG: -261.35
    - !!omap
      - id: "s_0770"
      - name: "glycerol 3-phosphate"
      - compartment: "m"
      - formula: "C3H7O6P"
      - charge: -2
      - smiles: "C(C(COP(=O)(O)O)O)O"
      - annotation: !!omap
          - bigg.metabolite: "glyc3p"
          - chebi: "CHEBI:57597"
          - kegg.compound: "C00093"
          - metanetx.chemical: "MNXM66"
          - sbo: "SBO:0000247"
      - deltaG: -261.35
    - !!omap
      - id: "s_0771"
      - name: "glycerone"
      - compartment: "c"
      - formula: "C3H6O3"
      - charge: 0
      - smiles: "C(C(=O)CO)O"
      - annotation: !!omap
          - bigg.metabolite: "dha"
          - chebi: "CHEBI:16016"
          - kegg.compound: "C00184"
          - metanetx.chemical: "MNXM460"
          - sbo: "SBO:0000247"
      - deltaG: -49.74
    - !!omap
      - id: "s_0773"
      - name: "glycogen"
      - compartment: "c"
      - formula: "C6H10O5"
      - charge: 0
      - smiles: "C(C1C(C(C(C(O1)OCC2C(C(C(C(O2)OC3C(OC(C(C3O)O)O)CO)O)O)OC4C(C(C(C(O4)CO)O)O)O)O)O)O)O"
      - annotation: !!omap
          - bigg.metabolite: "glycogen"
          - chebi: "CHEBI:28087"
          - kegg.compound: "C00182"
          - metanetx.chemical: "MNXM55375"
          - sbo: "SBO:0000247"
      - deltaG: -284.77
    - !!omap
      - id: "s_0774"
      - name: "glycogen"
      - compartment: "v"
      - formula: "C6H10O5"
      - charge: 0
      - smiles: "C(C1C(C(C(C(O1)OCC2C(C(C(C(O2)OC3C(OC(C(C3O)O)O)CO)O)O)OC4C(C(C(C(O4)CO)O)O)O)O)O)O)O"
      - annotation: !!omap
          - bigg.metabolite: "glycogen"
          - chebi: "CHEBI:28087"
          - kegg.compound: "C00182"
          - metanetx.chemical: "MNXM55375"
          - sbo: "SBO:0000247"
      - deltaG: -284.77
    - !!omap
      - id: "s_0775"
      - name: "glycolaldehyde"
      - compartment: "c"
      - formula: "C2H4O2"
      - charge: 0
      - smiles: "C(C=O)O"
      - annotation: !!omap
          - bigg.metabolite: "gcald"
          - chebi: "CHEBI:17071"
          - kegg.compound: "C00266"
          - metanetx.chemical: "MNXM349"
          - sbo: "SBO:0000247"
      - deltaG: -31.33
    - !!omap
      - id: "s_0776"
      - name: "glycolaldehyde"
      - compartment: "e"
      - formula: "C2H4O2"
      - charge: 0
      - smiles: "C(C=O)O"
      - annotation: !!omap
          - bigg.metabolite: "gcald"
          - chebi: "CHEBI:17071"
          - kegg.compound: "C00266"
          - metanetx.chemical: "MNXM349"
          - sbo: "SBO:0000247"
      - deltaG: -31.33
    - !!omap
      - id: "s_0777"
      - name: "glycolaldehyde"
      - compartment: "m"
      - formula: "C2H4O2"
      - charge: 0
      - smiles: "C(C=O)O"
      - annotation: !!omap
          - bigg.metabolite: "gcald"
          - chebi: "CHEBI:17071"
          - kegg.compound: "C00266"
          - metanetx.chemical: "MNXM349"
          - sbo: "SBO:0000247"
      - deltaG: -31.33
    - !!omap
      - id: "s_0779"
      - name: "glyoxylate"
      - compartment: "c"
      - formula: "C2HO3"
      - charge: -1
      - smiles: "C(=O)C(=O)[O-]"
      - annotation: !!omap
          - bigg.metabolite: "glx"
          - chebi: "CHEBI:36655"
          - kegg.compound: "C00048"
          - metanetx.chemical: "MNXM69"
          - sbo: "SBO:0000247"
      - deltaG: -101.5
    - !!omap
      - id: "s_0780"
      - name: "glyoxylate"
      - compartment: "e"
      - formula: "C2HO3"
      - charge: -1
      - smiles: "C(=O)C(=O)[O-]"
      - annotation: !!omap
          - bigg.metabolite: "glx"
          - chebi: "CHEBI:36655"
          - kegg.compound: "C00048"
          - metanetx.chemical: "MNXM69"
          - sbo: "SBO:0000247"
      - deltaG: -101.5
    - !!omap
      - id: "s_0781"
      - name: "glyoxylate"
      - compartment: "p"
      - formula: "C2HO3"
      - charge: -1
      - smiles: "C(=O)C(=O)[O-]"
      - annotation: !!omap
          - bigg.metabolite: "glx"
          - chebi: "CHEBI:36655"
          - kegg.compound: "C00048"
          - metanetx.chemical: "MNXM69"
          - sbo: "SBO:0000247"
      - deltaG: -101.5
    - !!omap
      - id: "s_0782"
      - name: "GMP"
      - compartment: "c"
      - formula: "C10H12N5O8P"
      - charge: -2
      - smiles: "C1=NC2=C(N1C3C(C(C(O3)COP(=O)(O)O)O)O)N=C(NC2=O)N"
      - annotation: !!omap
          - bigg.metabolite: "gmp"
          - chebi: "CHEBI:58115"
          - kegg.compound: "C00144"
          - metanetx.chemical: "MNXM113"
          - sbo: "SBO:0000247"
      - deltaG: -192.26
    - !!omap
      - id: "s_0783"
      - name: "GMP"
      - compartment: "g"
      - formula: "C10H12N5O8P"
      - charge: -2
      - smiles: "C1=NC2=C(N1C3C(C(C(O3)COP(=O)(O)O)O)O)N=C(NC2=O)N"
      - annotation: !!omap
          - bigg.metabolite: "gmp"
          - chebi: "CHEBI:58115"
          - kegg.compound: "C00144"
          - metanetx.chemical: "MNXM113"
          - sbo: "SBO:0000247"
      - deltaG: -192.26
    - !!omap
      - id: "s_0785"
      - name: "GTP"
      - compartment: "c"
      - formula: "C10H12N5O14P3"
      - charge: -4
      - smiles: "C1=NC2=C(N1C3C(C(C(O3)COP(=O)(O)OP(=O)(O)OP(=O)(O)O)O)O)N=C(NC2=O)N"
      - annotation: !!omap
          - bigg.metabolite: "gtp"
          - chebi: "CHEBI:57600"
          - kegg.compound: "C00044"
          - metanetx.chemical: "MNXM51"
          - sbo: "SBO:0000247"
      - deltaG: -600.54
    - !!omap
      - id: "s_0786"
      - name: "GTP"
      - compartment: "m"
      - formula: "C10H12N5O14P3"
      - charge: -4
      - smiles: "C1=NC2=C(N1C3C(C(C(O3)COP(=O)(O)OP(=O)(O)OP(=O)(O)O)O)O)N=C(NC2=O)N"
      - annotation: !!omap
          - bigg.metabolite: "gtp"
          - chebi: "CHEBI:57600"
          - kegg.compound: "C00044"
          - metanetx.chemical: "MNXM51"
          - sbo: "SBO:0000247"
      - deltaG: -600.54
    - !!omap
      - id: "s_0787"
      - name: "guanine"
      - compartment: "c"
      - formula: "C5H5N5O"
      - charge: 0
      - smiles: "C1=NC2=C(N1)C(=O)NC(=N2)N"
      - annotation: !!omap
          - bigg.metabolite: "gua"
          - chebi: "CHEBI:16235"
          - kegg.compound: "C00242"
          - metanetx.chemical: "MNXM259"
          - sbo: "SBO:0000247"
      - deltaG: 64.47
    - !!omap
      - id: "s_0788"
      - name: "guanine"
      - compartment: "e"
      - formula: "C5H5N5O"
      - charge: 0
      - smiles: "C1=NC2=C(N1)C(=O)NC(=N2)N"
      - annotation: !!omap
          - bigg.metabolite: "gua"
          - chebi: "CHEBI:16235"
          - kegg.compound: "C00242"
          - metanetx.chemical: "MNXM259"
          - sbo: "SBO:0000247"
      - deltaG: 64.47
    - !!omap
      - id: "s_0789"
      - name: "guanine"
      - compartment: "m"
      - formula: "C5H5N5O"
      - charge: 0
      - smiles: "C1=NC2=C(N1)C(=O)NC(=N2)N"
      - annotation: !!omap
          - bigg.metabolite: "gua"
          - chebi: "CHEBI:16235"
          - kegg.compound: "C00242"
          - metanetx.chemical: "MNXM259"
          - sbo: "SBO:0000247"
      - deltaG: 64.47
    - !!omap
      - id: "s_0790"
      - name: "guanosine"
      - compartment: "c"
      - formula: "C10H13N5O5"
      - charge: 0
      - smiles: "C1=NC2=C(N1C3C(C(C(O3)CO)O)O)N=C(NC2=O)N"
      - annotation: !!omap
          - bigg.metabolite: "gsn"
          - chebi: "CHEBI:16750"
          - kegg.compound: "C00387"
          - metanetx.chemical: "MNXM401"
          - sbo: "SBO:0000247"
      - deltaG: 30.78
    - !!omap
      - id: "s_0791"
      - name: "guanosine"
      - compartment: "e"
      - formula: "C10H13N5O5"
      - charge: 0
      - smiles: "C1=NC2=C(N1C3C(C(C(O3)CO)O)O)N=C(NC2=O)N"
      - annotation: !!omap
          - bigg.metabolite: "gsn"
          - chebi: "CHEBI:16750"
          - kegg.compound: "C00387"
          - metanetx.chemical: "MNXM401"
          - sbo: "SBO:0000247"
      - deltaG: 30.78
    - !!omap
      - id: "s_0792"
      - name: "guanosine"
      - compartment: "m"
      - formula: "C10H13N5O5"
      - charge: 0
      - smiles: "C1=NC2=C(N1C3C(C(C(O3)CO)O)O)N=C(NC2=O)N"
      - annotation: !!omap
          - bigg.metabolite: "gsn"
          - chebi: "CHEBI:16750"
          - kegg.compound: "C00387"
          - metanetx.chemical: "MNXM401"
          - sbo: "SBO:0000247"
      - deltaG: 30.78
    - !!omap
      - id: "s_0793"
      - name: "H+"
      - compartment: "ce"
      - formula: "H"
      - charge: 1
      - smiles: "[H+]"
      - annotation: !!omap
          - bigg.metabolite: "h"
          - chebi: "CHEBI:24636"
          - kegg.compound: "C00080"
          - metanetx.chemical: "MNXM1"
          - sbo: "SBO:0000247"
      - deltaG: 9.54
    - !!omap
      - id: "s_0794"
      - name: "H+"
      - compartment: "c"
      - formula: "H"
      - charge: 1
      - smiles: "[H+]"
      - annotation: !!omap
          - bigg.metabolite: "h"
          - chebi: "CHEBI:24636"
          - kegg.compound: "C00080"
          - metanetx.chemical: "MNXM1"
          - sbo: "SBO:0000247"
      - deltaG: 9.54
    - !!omap
      - id: "s_0795"
      - name: "H+"
      - compartment: "er"
      - formula: "H"
      - charge: 1
      - smiles: "[H+]"
      - annotation: !!omap
          - bigg.metabolite: "h"
          - chebi: "CHEBI:24636"
          - kegg.compound: "C00080"
          - metanetx.chemical: "MNXM1"
          - sbo: "SBO:0000247"
      - deltaG: 9.54
    - !!omap
      - id: "s_0796"
      - name: "H+"
      - compartment: "e"
      - formula: "H"
      - charge: 1
      - smiles: "[H+]"
      - annotation: !!omap
          - bigg.metabolite: "h"
          - chebi: "CHEBI:24636"
          - kegg.compound: "C00080"
          - metanetx.chemical: "MNXM1"
          - sbo: "SBO:0000247"
      - deltaG: 9.54
    - !!omap
      - id: "s_0797"
      - name: "H+"
      - compartment: "g"
      - formula: "H"
      - charge: 1
      - smiles: "[H+]"
      - annotation: !!omap
          - bigg.metabolite: "h"
          - chebi: "CHEBI:24636"
          - kegg.compound: "C00080"
          - metanetx.chemical: "MNXM1"
          - sbo: "SBO:0000247"
      - deltaG: 9.54
    - !!omap
      - id: "s_0798"
      - name: "H+"
      - compartment: "lp"
      - formula: "H"
      - charge: 1
      - smiles: "[H+]"
      - annotation: !!omap
          - bigg.metabolite: "h"
          - chebi: "CHEBI:24636"
          - kegg.compound: "C00080"
          - metanetx.chemical: "MNXM1"
          - sbo: "SBO:0000247"
      - deltaG: 9.54
    - !!omap
      - id: "s_0799"
      - name: "H+"
      - compartment: "m"
      - formula: "H"
      - charge: 1
      - smiles: "[H+]"
      - annotation: !!omap
          - bigg.metabolite: "h"
          - chebi: "CHEBI:24636"
          - kegg.compound: "C00080"
          - metanetx.chemical: "MNXM1"
          - sbo: "SBO:0000247"
      - deltaG: 9.54
    - !!omap
      - id: "s_0800"
      - name: "H+"
      - compartment: "n"
      - formula: "H"
      - charge: 1
      - smiles: "[H+]"
      - annotation: !!omap
          - bigg.metabolite: "h"
          - chebi: "CHEBI:24636"
          - kegg.compound: "C00080"
          - metanetx.chemical: "MNXM1"
          - sbo: "SBO:0000247"
      - deltaG: 9.54
    - !!omap
      - id: "s_0801"
      - name: "H+"
      - compartment: "p"
      - formula: "H"
      - charge: 1
      - smiles: "[H+]"
      - annotation: !!omap
          - bigg.metabolite: "h"
          - chebi: "CHEBI:24636"
          - kegg.compound: "C00080"
          - metanetx.chemical: "MNXM1"
          - sbo: "SBO:0000247"
      - deltaG: 9.54
    - !!omap
      - id: "s_0802"
      - name: "H+"
      - compartment: "v"
      - formula: "H"
      - charge: 1
      - smiles: "[H+]"
      - annotation: !!omap
          - bigg.metabolite: "h"
          - chebi: "CHEBI:24636"
          - kegg.compound: "C00080"
          - metanetx.chemical: "MNXM1"
          - sbo: "SBO:0000247"
      - deltaG: 9.54
    - !!omap
      - id: "s_0803"
      - name: "H2O"
      - compartment: "c"
      - formula: "H2O"
      - charge: 0
      - smiles: "O"
      - annotation: !!omap
          - bigg.metabolite: "h2o"
          - chebi: "CHEBI:15377"
          - kegg.compound: "C00001"
          - metanetx.chemical: "MNXM2"
          - sbo: "SBO:0000247"
      - deltaG: -37.21
    - !!omap
      - id: "s_0804"
      - name: "H2O"
      - compartment: "er"
      - formula: "H2O"
      - charge: 0
      - smiles: "O"
      - annotation: !!omap
          - bigg.metabolite: "h2o"
          - chebi: "CHEBI:15377"
          - kegg.compound: "C00001"
          - metanetx.chemical: "MNXM2"
          - sbo: "SBO:0000247"
      - deltaG: -37.21
    - !!omap
      - id: "s_0805"
      - name: "H2O"
      - compartment: "e"
      - formula: "H2O"
      - charge: 0
      - smiles: "O"
      - annotation: !!omap
          - bigg.metabolite: "h2o"
          - chebi: "CHEBI:15377"
          - kegg.compound: "C00001"
          - metanetx.chemical: "MNXM2"
          - sbo: "SBO:0000247"
      - deltaG: -37.21
    - !!omap
      - id: "s_0806"
      - name: "H2O"
      - compartment: "g"
      - formula: "H2O"
      - charge: 0
      - smiles: "O"
      - annotation: !!omap
          - bigg.metabolite: "h2o"
          - chebi: "CHEBI:15377"
          - kegg.compound: "C00001"
          - metanetx.chemical: "MNXM2"
          - sbo: "SBO:0000247"
      - deltaG: -37.21
    - !!omap
      - id: "s_0807"
      - name: "H2O"
      - compartment: "m"
      - formula: "H2O"
      - charge: 0
      - smiles: "O"
      - annotation: !!omap
          - bigg.metabolite: "h2o"
          - chebi: "CHEBI:15377"
          - kegg.compound: "C00001"
          - metanetx.chemical: "MNXM2"
          - sbo: "SBO:0000247"
      - deltaG: -37.21
    - !!omap
      - id: "s_0808"
      - name: "H2O"
      - compartment: "n"
      - formula: "H2O"
      - charge: 0
      - smiles: "O"
      - annotation: !!omap
          - bigg.metabolite: "h2o"
          - chebi: "CHEBI:15377"
          - kegg.compound: "C00001"
          - metanetx.chemical: "MNXM2"
          - sbo: "SBO:0000247"
      - deltaG: -37.21
    - !!omap
      - id: "s_0809"
      - name: "H2O"
      - compartment: "p"
      - formula: "H2O"
      - charge: 0
      - smiles: "O"
      - annotation: !!omap
          - bigg.metabolite: "h2o"
          - chebi: "CHEBI:15377"
          - kegg.compound: "C00001"
          - metanetx.chemical: "MNXM2"
          - sbo: "SBO:0000247"
      - deltaG: -37.21
    - !!omap
      - id: "s_0810"
      - name: "H2O"
      - compartment: "v"
      - formula: "H2O"
      - charge: 0
      - smiles: "O"
      - annotation: !!omap
          - bigg.metabolite: "h2o"
          - chebi: "CHEBI:15377"
          - kegg.compound: "C00001"
          - metanetx.chemical: "MNXM2"
          - sbo: "SBO:0000247"
      - deltaG: -37.21
    - !!omap
      - id: "s_0811"
      - name: "heme a"
      - compartment: "m"
      - formula: "C49H55FeN4O6"
      - charge: -3
      - smiles: "CC1=C(C2=CC3=NC(=CC4=C(C(=C([N-]4)C=C5C(=C(C(=N5)C=C1[N-]2)C(CCC=C(C)CCC=C(C)CCC=C(C)C)O)C)C=O)CCC(=O)O)C(=C3C)CCC(=O)O)C=C.[Fe]"
      - annotation: !!omap
          - bigg.metabolite: "hemeA"
          - chebi: "CHEBI:24479"
          - kegg.compound: "C15670"
          - metanetx.chemical: "MNXM53309"
          - sbo: "SBO:0000247"
      - deltaG: 10000000
    - !!omap
      - id: "s_0812"
      - name: "heme o"
      - compartment: "m"
      - formula: "C49H56FeN4O5"
      - charge: -2
      - smiles: "CC1=C(C2=CC3=NC(=CC4=C(C(=C([N-]4)C=C5C(=C(C(=N5)C=C1[N-]2)C=C)C)C(CCC=C(C)CCC=C(C)CCC=C(C)C)O)C)C(=C3CCC(=O)O)C)CCC(=O)O.[Fe+2]"
      - annotation: !!omap
          - bigg.metabolite: "hemeO"
          - chebi: "CHEBI:24480"
          - kegg.compound: "C15672"
          - metanetx.chemical: "MNXM1278"
          - sbo: "SBO:0000247"
      - deltaG: 10000000
    - !!omap
      - id: "s_0813"
      - name: "henicosaprenyl diphosphate"
      - compartment: "lp"
      - formula: "C105H169O7P2"
      - charge: -3
      - smiles: "CC(=CCCC(=CCCC(=CCCC(=CCCC(=CCCC(=CCCC(=CCCC(=CCCC(=CCCC(=CCCC(=CCCC(=CCCC(=CCCC(=CCCC(=CCCC(=CCCC(=CCCC(=CCCC(=CCCC(=CCCC(=CCOP(=O)(O)OP(=O)(O)O)C)C)C)C)C)C)C)C)C)C)C)C)C)C)C)C)C)C)C)C)C"
      - annotation: !!omap
          - chebi: "CHEBI:53032"
          - metanetx.chemical: "MNXM56060"
          - sbo: "SBO:0000247"
      - deltaG: 1776.34
    - !!omap
      - id: "s_0814"
      - name: "heptadecaprenyl diphosphate"
      - compartment: "lp"
      - formula: "C85H137O7P2"
      - charge: -3
      - smiles: "CC(=CCCC(=CCCC(=CCCC(=CCCC(=CCCC(=CCCC(=CCCC(=CCCC(=CCCC(=CCCC(=CCCC(=CCCC(=CCCC(=CCCC(=CCCC(=CCCC(=CCOP(=O)(O)OP(=O)(O)O)C)C)C)C)C)C)C)C)C)C)C)C)C)C)C)C)C"
      - annotation: !!omap
          - chebi: "CHEBI:53036"
          - metanetx.chemical: "MNXM56121"
          - sbo: "SBO:0000247"
      - deltaG: 1352.4
    - !!omap
      - id: "s_0815"
      - name: "heptaprenyl diphosphate"
      - compartment: "lp"
      - formula: "C35H57O7P2"
      - charge: -3
      - smiles: "CC(=CCCC(=CCCC(=CCCC(=CCCC(=CCCC(=CCCC(=CCOP(=O)(O)OP(=O)(O)O)C)C)C)C)C)C)C"
      - annotation: !!omap
          - bigg.metabolite: "hepdp"
          - chebi: "CHEBI:53046"
          - kegg.compound: "C04216"
          - metanetx.chemical: "MNXM1722"
          - sbo: "SBO:0000247"
      - deltaG: 292.55
    - !!omap
      - id: "s_0816"
      - name: "hexacosanoyl-CoA"
      - compartment: "c"
      - formula: "C47H82N7O17P3S"
      - charge: -4
      - smiles: "CCCCCCCCCCCCCCCCCCCCCCCCCC(=O)SCCNC(=O)CCNC(=O)C(C(C)(C)COP(=O)(O)OP(=O)(O)OCC1C(C(C(O1)N2C=NC3=C(N=CN=C32)N)O)OP(=O)(O)O)O"
      - annotation: !!omap
          - bigg.metabolite: "hexccoa"
          - chebi: "CHEBI:52966"
          - metanetx.chemical: "MNXM1190"
          - sbo: "SBO:0000247"
      - deltaG: 37.67
    - !!omap
      - id: "s_0817"
      - name: "hexacosanoyl-CoA"
      - compartment: "er"
      - formula: "C47H82N7O17P3S"
      - charge: -4
      - smiles: "CCCCCCCCCCCCCCCCCCCCCCCCCC(=O)SCCNC(=O)CCNC(=O)C(C(C)(C)COP(=O)(O)OP(=O)(O)OCC1C(C(C(O1)N2C=NC3=C(N=CN=C32)N)O)OP(=O)(O)O)O"
      - annotation: !!omap
          - bigg.metabolite: "hexccoa"
          - chebi: "CHEBI:52966"
          - metanetx.chemical: "MNXM1190"
          - sbo: "SBO:0000247"
      - deltaG: 37.67
    - !!omap
      - id: "s_0819"
      - name: "hexacosanoyl-CoA"
      - compartment: "p"
      - formula: "C47H82N7O17P3S"
      - charge: -4
      - smiles: "CCCCCCCCCCCCCCCCCCCCCCCCCC(=O)SCCNC(=O)CCNC(=O)C(C(C)(C)COP(=O)(O)OP(=O)(O)OCC1C(C(C(O1)N2C=NC3=C(N=CN=C32)N)O)OP(=O)(O)O)O"
      - annotation: !!omap
          - bigg.metabolite: "hexccoa"
          - chebi: "CHEBI:52966"
          - metanetx.chemical: "MNXM1190"
          - sbo: "SBO:0000247"
      - deltaG: 37.67
    - !!omap
      - id: "s_0823"
      - name: "hexadec-2-enoyl-CoA"
      - compartment: "p"
      - formula: "C37H60N7O17P3S"
      - charge: -4
      - smiles: "CCCCCCCCCCCCCC=CC(=O)SCCNC(=O)CCNC(=O)C(C(C)(C)COP(=O)(O)OP(=O)(O)OCC1C(C(C(O1)N2C=NC3=C(N=CN=C32)N)O)OP(=O)(O)O)O"
      - annotation: !!omap
          - bigg.metabolite: "hdd2coa"
          - kegg.compound: "C05272"
          - metanetx.chemical: "MNXM581"
          - sbo: "SBO:0000247"
      - deltaG: -82.28
    - !!omap
      - id: "s_0825"
      - name: "hexadecanal"
      - compartment: "er"
      - formula: "C16H32O"
      - charge: 0
      - smiles: "CCCCCCCCCCCCCCCC=O"
      - annotation: !!omap
          - bigg.metabolite: "hxdcal"
          - chebi: "CHEBI:17600"
          - kegg.compound: "C00517"
          - metanetx.chemical: "MNXM528"
          - sbo: "SBO:0000247"
      - deltaG: 302.74
    - !!omap
      - id: "s_0826"
      - name: "hexadecanal"
      - compartment: "e"
      - formula: "C16H32O"
      - charge: 0
      - smiles: "CCCCCCCCCCCCCCCC=O"
      - annotation: !!omap
          - bigg.metabolite: "hxdcal"
          - chebi: "CHEBI:17600"
          - kegg.compound: "C00517"
          - metanetx.chemical: "MNXM528"
          - sbo: "SBO:0000247"
      - deltaG: 302.74
    - !!omap
      - id: "s_0829"
      - name: "hexadecaprenyl diphosphate"
      - compartment: "lp"
      - formula: "C80H129O7P2"
      - charge: -3
      - smiles: "CC(=CCCC(=CCCC(=CCCC(=CCCC(=CCCC(=CCCC(=CCCC(=CCCC(=CCCC(=CCCC(=CCCC(=CCCC(=CCCC(=CCCC(=CCCC(=CCOP(=O)(O)OP(=O)(O)O)C)C)C)C)C)C)C)C)C)C)C)C)C)C)C)C"
      - annotation: !!omap
          - chebi: "CHEBI:53037"
          - metanetx.chemical: "MNXM56347"
          - sbo: "SBO:0000247"
      - deltaG: 1246.41
    - !!omap
      - id: "s_0830"
      - name: "hexaprenyl diphosphate"
      - compartment: "lp"
      - formula: "C30H49O7P2"
      - charge: -3
      - smiles: "CC(=CCCC(=CCCC(=CCCC(=CCCC(=CCCC(=CCOP(=O)(O)OP(=O)(O)O)C)C)C)C)C)C"
      - annotation: !!omap
          - bigg.metabolite: "hexdp"
          - chebi: "CHEBI:53047"
          - kegg.compound: "C01230"
          - metanetx.chemical: "MNXM1067"
          - sbo: "SBO:0000247"
      - deltaG: 186.56
    - !!omap
      - id: "s_0831"
      - name: "hexaprenyl diphosphate"
      - compartment: "m"
      - formula: "C30H49O7P2"
      - charge: -3
      - smiles: "CC(=CCCC(=CCCC(=CCCC(=CCCC(=CCCC(=CCOP(=O)(O)OP(=O)(O)O)C)C)C)C)C)C"
      - annotation: !!omap
          - bigg.metabolite: "hexdp"
          - chebi: "CHEBI:53047"
          - kegg.compound: "C01230"
          - metanetx.chemical: "MNXM1067"
          - sbo: "SBO:0000247"
      - deltaG: 186.56
    - !!omap
      - id: "s_0832"
      - name: "His-tRNA(His)"
      - compartment: "c"
      - formula: "C6H9N3OR"
      - charge: 1
      - annotation: !!omap
          - bigg.metabolite: "histrna"
          - chebi: "CHEBI:29155"
          - kegg.compound: "C02988"
          - metanetx.chemical: "MNXM89831"
          - sbo: "SBO:0000247"
      - deltaG: 10000000
    - !!omap
      - id: "s_0833"
      - name: "His-tRNA(His)"
      - compartment: "m"
      - formula: "C6H9N3OR"
      - charge: 1
      - annotation: !!omap
          - bigg.metabolite: "histrna"
          - chebi: "CHEBI:29155"
          - kegg.compound: "C02988"
          - metanetx.chemical: "MNXM89831"
          - sbo: "SBO:0000247"
      - deltaG: 10000000
    - !!omap
      - id: "s_0834"
      - name: "homocitrate"
      - compartment: "m"
      - formula: "C7H7O7"
      - charge: -3
      - smiles: "C(CC(CC(=O)O)(C(=O)O)O)C(=O)O"
      - annotation: !!omap
          - chebi: "CHEBI:36457"
          - kegg.compound: "C01251"
          - metanetx.chemical: "MNXM722779"
          - sbo: "SBO:0000247"
      - deltaG: -207.63
    - !!omap
      - id: "s_0835"
      - name: "homocitrate"
      - compartment: "n"
      - formula: "C7H7O7"
      - charge: -3
      - smiles: "C(CC(CC(=O)O)(C(=O)O)O)C(=O)O"
      - annotation: !!omap
          - chebi: "CHEBI:36457"
          - kegg.compound: "C01251"
          - metanetx.chemical: "MNXM722779"
          - sbo: "SBO:0000247"
      - deltaG: -207.63
    - !!omap
      - id: "s_0836"
      - name: "homoisocitrate"
      - compartment: "m"
      - formula: "C7H7O7"
      - charge: -3
      - smiles: "C(CC(=O)O)C(C(C(=O)O)O)C(=O)O"
      - annotation: !!omap
          - bigg.metabolite: "hicit"
          - chebi: "CHEBI:30904"
          - kegg.compound: "C05662"
          - metanetx.chemical: "MNXM984"
          - sbo: "SBO:0000247"
      - deltaG: -206.21
    - !!omap
      - id: "s_0837"
      - name: "hydrogen peroxide"
      - compartment: "c"
      - formula: "H2O2"
      - charge: 0
      - smiles: "OO"
      - annotation: !!omap
          - bigg.metabolite: "h2o2"
          - chebi: "CHEBI:16240"
          - kegg.compound: "C00027"
          - metanetx.chemical: "MNXM22"
          - sbo: "SBO:0000247"
      - deltaG: -12.58
    - !!omap
      - id: "s_0838"
      - name: "hydrogen peroxide"
      - compartment: "m"
      - formula: "H2O2"
      - charge: 0
      - smiles: "OO"
      - annotation: !!omap
          - bigg.metabolite: "h2o2"
          - chebi: "CHEBI:16240"
          - kegg.compound: "C00027"
          - metanetx.chemical: "MNXM22"
          - sbo: "SBO:0000247"
      - deltaG: -12.58
    - !!omap
      - id: "s_0839"
      - name: "hydrogen peroxide"
      - compartment: "n"
      - formula: "H2O2"
      - charge: 0
      - smiles: "OO"
      - annotation: !!omap
          - bigg.metabolite: "h2o2"
          - chebi: "CHEBI:16240"
          - kegg.compound: "C00027"
          - metanetx.chemical: "MNXM22"
          - sbo: "SBO:0000247"
      - deltaG: -12.58
    - !!omap
      - id: "s_0840"
      - name: "hydrogen peroxide"
      - compartment: "p"
      - formula: "H2O2"
      - charge: 0
      - smiles: "OO"
      - annotation: !!omap
          - bigg.metabolite: "h2o2"
          - chebi: "CHEBI:16240"
          - kegg.compound: "C00027"
          - metanetx.chemical: "MNXM22"
          - sbo: "SBO:0000247"
      - deltaG: -12.58
    - !!omap
      - id: "s_0841"
      - name: "hydrogen sulfide"
      - compartment: "c"
      - formula: "HS"
      - charge: 0
      - smiles: "S"
      - annotation: !!omap
          - bigg.metabolite: "h2s"
          - biocyc: "HS"
          - chebi: "CHEBI:16136"
          - kegg.compound: "C00283"
          - metanetx.chemical: "MNXM124865"
          - sbo: "SBO:0000247"
      - deltaG: 10000000
    - !!omap
      - id: "s_0843"
      - name: "hypoxanthine"
      - compartment: "c"
      - formula: "C5H4N4O"
      - charge: 0
      - smiles: "C1=NC2=C(N1)C(=O)NC=N2"
      - annotation: !!omap
          - bigg.metabolite: "hxan"
          - chebi: "CHEBI:17368"
          - kegg.compound: "C00262"
          - metanetx.chemical: "MNXM213"
          - sbo: "SBO:0000247"
      - deltaG: 62.03
    - !!omap
      - id: "s_0844"
      - name: "hypoxanthine"
      - compartment: "e"
      - formula: "C5H4N4O"
      - charge: 0
      - smiles: "C1=NC2=C(N1)C(=O)NC=N2"
      - annotation: !!omap
          - bigg.metabolite: "hxan"
          - chebi: "CHEBI:17368"
          - kegg.compound: "C00262"
          - metanetx.chemical: "MNXM213"
          - sbo: "SBO:0000247"
      - deltaG: 62.03
    - !!omap
      - id: "s_0845"
      - name: "icosaprenyl diphosphate"
      - compartment: "lp"
      - formula: "C100H161O7P2"
      - charge: -3
      - smiles: "CC(=CCCC(=CCCC(=CCCC(=CCCC(=CCCC(=CCCC(=CCCC(=CCCC(=CCCC(=CCCC(=CCCC(=CCCC(=CCCC(=CCCC(=CCCC(=CCCC(=CCCC(=CCCC(=CCCC(=CCOP(=O)(O)OP(=O)(O)O)C)C)C)C)C)C)C)C)C)C)C)C)C)C)C)C)C)C)C)C"
      - annotation: !!omap
          - chebi: "CHEBI:53033"
          - metanetx.chemical: "MNXM57076"
          - sbo: "SBO:0000247"
      - deltaG: 1670.35
    - !!omap
      - id: "s_0846"
      - name: "IDP"
      - compartment: "c"
      - formula: "C10H11N4O11P2"
      - charge: -3
      - smiles: "C1=NC2=C(C(=O)N1)N=CN2C3C(C(C(O3)COP(=O)(O)OP(=O)(O)O)O)O"
      - annotation: !!omap
          - bigg.metabolite: "idp"
          - chebi: "CHEBI:58280"
          - kegg.compound: "C00104"
          - metanetx.chemical: "MNXM495"
          - sbo: "SBO:0000247"
      - deltaG: -424.2
    - !!omap
      - id: "s_0847"
      - name: "Ile-tRNA(Ile)"
      - compartment: "c"
      - formula: "C6H13NOR"
      - charge: 1
      - annotation: !!omap
          - bigg.metabolite: "iletrna"
          - chebi: "CHEBI:29160"
          - kegg.compound: "C03127"
          - metanetx.chemical: "MNXM89832"
          - sbo: "SBO:0000247"
      - deltaG: 10000000
    - !!omap
      - id: "s_0848"
      - name: "Ile-tRNA(Ile)"
      - compartment: "m"
      - formula: "C6H13NOR"
      - charge: 1
      - annotation: !!omap
          - bigg.metabolite: "iletrna"
          - chebi: "CHEBI:29160"
          - kegg.compound: "C03127"
          - metanetx.chemical: "MNXM89832"
          - sbo: "SBO:0000247"
      - deltaG: 10000000
    - !!omap
      - id: "s_0849"
      - name: "IMP"
      - compartment: "c"
      - formula: "C10H11N4O8P"
      - charge: -2
      - smiles: "C1=NC2=C(C(=O)N1)N=CN2C3C(C(C(O3)COP(=O)(O)O)O)O"
      - annotation: !!omap
          - bigg.metabolite: "imp"
          - chebi: "CHEBI:58053"
          - kegg.compound: "C00130"
          - metanetx.chemical: "MNXM125"
          - sbo: "SBO:0000247"
      - deltaG: -221.02
    - !!omap
      - id: "s_0850"
      - name: "indol-3-ylacetaldehyde"
      - compartment: "c"
      - formula: "C10H9NO"
      - charge: 0
      - smiles: "C1=CC=C2C(=C1)C(=CN2)CC=O"
      - annotation: !!omap
          - bigg.metabolite: "id3acald"
          - chebi: "CHEBI:18086"
          - kegg.compound: "C00637"
          - metanetx.chemical: "MNXM518"
          - sbo: "SBO:0000247"
      - deltaG: 126.21
    - !!omap
      - id: "s_0851"
      - name: "indol-3-ylacetaldehyde"
      - compartment: "e"
      - formula: "C10H9NO"
      - charge: 0
      - smiles: "C1=CC=C2C(=C1)C(=CN2)CC=O"
      - annotation: !!omap
          - bigg.metabolite: "id3acald"
          - chebi: "CHEBI:18086"
          - kegg.compound: "C00637"
          - metanetx.chemical: "MNXM518"
          - sbo: "SBO:0000247"
      - deltaG: 126.21
    - !!omap
      - id: "s_0852"
      - name: "indol-3-ylacetaldehyde"
      - compartment: "m"
      - formula: "C10H9NO"
      - charge: 0
      - smiles: "C1=CC=C2C(=C1)C(=CN2)CC=O"
      - annotation: !!omap
          - bigg.metabolite: "id3acald"
          - chebi: "CHEBI:18086"
          - kegg.compound: "C00637"
          - metanetx.chemical: "MNXM518"
          - sbo: "SBO:0000247"
      - deltaG: 126.21
    - !!omap
      - id: "s_0853"
      - name: "indole-3-acetate"
      - compartment: "c"
      - formula: "C10H8NO2"
      - charge: -1
      - smiles: "C1=CC=C2C(=C1)C(=CN2)CC(=O)[O-]"
      - annotation: !!omap
          - bigg.metabolite: "ind3ac"
          - chebi: "CHEBI:30854"
          - kegg.compound: "C00954"
          - metanetx.chemical: "MNXM383"
          - sbo: "SBO:0000247"
      - deltaG: 63.58
    - !!omap
      - id: "s_0854"
      - name: "indole-3-acetate"
      - compartment: "m"
      - formula: "C10H8NO2"
      - charge: -1
      - smiles: "C1=CC=C2C(=C1)C(=CN2)CC(=O)[O-]"
      - annotation: !!omap
          - bigg.metabolite: "ind3ac"
          - chebi: "CHEBI:30854"
          - kegg.compound: "C00954"
          - metanetx.chemical: "MNXM383"
          - sbo: "SBO:0000247"
      - deltaG: 63.58
    - !!omap
      - id: "s_0855"
      - name: "indole-3-pyruvate"
      - compartment: "c"
      - formula: "C11H8NO3"
      - charge: -1
      - smiles: "C1=CC=C2C(=C1)C(=CN2)CC(=O)C(=O)O"
      - annotation: !!omap
          - bigg.metabolite: "indpyr"
          - chebi: "CHEBI:17640"
          - kegg.compound: "C00331"
          - metanetx.chemical: "MNXM315"
          - sbo: "SBO:0000247"
      - deltaG: 37.64
    - !!omap
      - id: "s_0856"
      - name: "inosine"
      - compartment: "c"
      - formula: "C10H12N4O5"
      - charge: 0
      - smiles: "C1=NC2=C(C(=O)N1)N=CN2C3C(C(C(O3)CO)O)O"
      - annotation: !!omap
          - bigg.metabolite: "ins"
          - chebi: "CHEBI:17596"
          - kegg.compound: "C00294"
          - metanetx.chemical: "MNXM334"
          - sbo: "SBO:0000247"
      - deltaG: 17.39
    - !!omap
      - id: "s_0857"
      - name: "inosine"
      - compartment: "e"
      - formula: "C10H12N4O5"
      - charge: 0
      - smiles: "C1=NC2=C(C(=O)N1)N=CN2C3C(C(C(O3)CO)O)O"
      - annotation: !!omap
          - bigg.metabolite: "ins"
          - chebi: "CHEBI:17596"
          - kegg.compound: "C00294"
          - metanetx.chemical: "MNXM334"
          - sbo: "SBO:0000247"
      - deltaG: 17.39
    - !!omap
      - id: "s_0859"
      - name: "inositol phosphomannosylinositol phosphoceramide backbone"
      - compartment: "g"
      - formula: "C18H33NO23P2"
      - charge: 0
      - annotation: !!omap
          - chebi: "CHEBI:62682"
          - metanetx.chemical: "MNXM57447"
          - sbo: "SBO:0000649"
      - deltaG: 10000000
    - !!omap
      - id: "s_0861"
      - name: "inositol phosphomannosylinositol phosphoceramide A (C24)"
      - compartment: "er"
      - formula: "C60H117NO24P2"
      - charge: 0
      - smiles: "CCCCCCCCCCCCCCCCCCCCCCCC(=O)NC(COP(=O)(O)OC1C(C(C(C(C1OC2C(C(C(C(O2)COP(=O)(O)OC3C(C(C(C(C3O)O)O)O)O)O)O)O)O)O)O)O)C(CCCCCCCCCCCCCCC)O"
      - annotation: !!omap
          - chebi: "CHEBI:60439"
          - metanetx.chemical: "MNXM57464"
          - sbo: "SBO:0000247"
      - deltaG: 176.04
    - !!omap
      - id: "s_0862"
      - name: "inositol phosphomannosylinositol phosphoceramide A (C24)"
      - compartment: "g"
      - formula: "C60H117NO24P2"
      - charge: 0
      - smiles: "CCCCCCCCCCCCCCCCCCCCCCCC(=O)NC(COP(=O)(O)OC1C(C(C(C(C1OC2C(C(C(C(O2)COP(=O)(O)OC3C(C(C(C(C3O)O)O)O)O)O)O)O)O)O)O)O)C(CCCCCCCCCCCCCCC)O"
      - annotation: !!omap
          - chebi: "CHEBI:60439"
          - metanetx.chemical: "MNXM57464"
          - sbo: "SBO:0000247"
      - deltaG: 176.04
    - !!omap
      - id: "s_0863"
      - name: "inositol phosphomannosylinositol phosphoceramide A (C24)"
      - compartment: "m"
      - formula: "C60H117NO24P2"
      - charge: 0
      - smiles: "CCCCCCCCCCCCCCCCCCCCCCCC(=O)NC(COP(=O)(O)OC1C(C(C(C(C1OC2C(C(C(C(O2)COP(=O)(O)OC3C(C(C(C(C3O)O)O)O)O)O)O)O)O)O)O)O)C(CCCCCCCCCCCCCCC)O"
      - annotation: !!omap
          - chebi: "CHEBI:60439"
          - metanetx.chemical: "MNXM57464"
          - sbo: "SBO:0000247"
      - deltaG: 176.04
    - !!omap
      - id: "s_0864"
      - name: "inositol phosphomannosylinositol phosphoceramide A (C26)"
      - compartment: "er"
      - formula: "C62H121NO24P2"
      - charge: 0
      - smiles: "CCCCCCCCCCCCCCCCCCCCCCCCCC(=O)NC(COP(=O)(O)OC1C(C(C(C(C1OC2C(C(C(C(O2)COP(=O)(O)OC3C(C(C(C(C3O)O)O)O)O)O)O)O)O)O)O)O)C(CCCCCCCCCCCCCCCCC)O"
      - annotation: !!omap
          - chebi: "CHEBI:62715"
          - metanetx.chemical: "MNXM527419"
          - sbo: "SBO:0000247"
      - deltaG: 218.55
    - !!omap
      - id: "s_0865"
      - name: "inositol phosphomannosylinositol phosphoceramide A (C26)"
      - compartment: "g"
      - formula: "C62H121NO24P2"
      - charge: 0
      - smiles: "CCCCCCCCCCCCCCCCCCCCCCCCCC(=O)NC(COP(=O)(O)OC1C(C(C(C(C1OC2C(C(C(C(O2)COP(=O)(O)OC3C(C(C(C(C3O)O)O)O)O)O)O)O)O)O)O)O)C(CCCCCCCCCCCCCCCCC)O"
      - annotation: !!omap
          - chebi: "CHEBI:62715"
          - metanetx.chemical: "MNXM527419"
          - sbo: "SBO:0000247"
      - deltaG: 218.55
    - !!omap
      - id: "s_0866"
      - name: "inositol phosphomannosylinositol phosphoceramide A (C26)"
      - compartment: "m"
      - formula: "C62H121NO24P2"
      - charge: 0
      - smiles: "CCCCCCCCCCCCCCCCCCCCCCCCCC(=O)NC(COP(=O)(O)OC1C(C(C(C(C1OC2C(C(C(C(O2)COP(=O)(O)OC3C(C(C(C(C3O)O)O)O)O)O)O)O)O)O)O)O)C(CCCCCCCCCCCCCCCCC)O"
      - annotation: !!omap
          - chebi: "CHEBI:62715"
          - metanetx.chemical: "MNXM527419"
          - sbo: "SBO:0000247"
      - deltaG: 218.55
    - !!omap
      - id: "s_0867"
      - name: "inositol phosphomannosylinositol phosphoceramide B (C24)"
      - compartment: "er"
      - formula: "C60H117NO25P2"
      - charge: 0
      - smiles: "CCCCCCCCCCCCCCCCCCCCCCCC(=O)NC(COP(=O)(O)OC1C(C(C(C(C1OC2C(C(C(C(O2)COP(=O)(O)OC3C(C(C(C(C3O)O)O)O)O)O)O)O)O)O)O)O)C(C(CCCCCCCCCCCCCC)O)O"
      - annotation: !!omap
          - chebi: "CHEBI:60443"
          - metanetx.chemical: "MNXM527425"
          - sbo: "SBO:0000247"
      - deltaG: 138
    - !!omap
      - id: "s_0868"
      - name: "inositol phosphomannosylinositol phosphoceramide B (C24)"
      - compartment: "g"
      - formula: "C60H117NO25P2"
      - charge: 0
      - smiles: "CCCCCCCCCCCCCCCCCCCCCCCC(=O)NC(COP(=O)(O)OC1C(C(C(C(C1OC2C(C(C(C(O2)COP(=O)(O)OC3C(C(C(C(C3O)O)O)O)O)O)O)O)O)O)O)O)C(C(CCCCCCCCCCCCCC)O)O"
      - annotation: !!omap
          - chebi: "CHEBI:60443"
          - metanetx.chemical: "MNXM527425"
          - sbo: "SBO:0000247"
      - deltaG: 138
    - !!omap
      - id: "s_0869"
      - name: "inositol phosphomannosylinositol phosphoceramide B (C24)"
      - compartment: "m"
      - formula: "C60H117NO25P2"
      - charge: 0
      - smiles: "CCCCCCCCCCCCCCCCCCCCCCCC(=O)NC(COP(=O)(O)OC1C(C(C(C(C1OC2C(C(C(C(O2)COP(=O)(O)OC3C(C(C(C(C3O)O)O)O)O)O)O)O)O)O)O)O)C(C(CCCCCCCCCCCCCC)O)O"
      - annotation: !!omap
          - chebi: "CHEBI:60443"
          - metanetx.chemical: "MNXM527425"
          - sbo: "SBO:0000247"
      - deltaG: 138
    - !!omap
      - id: "s_0870"
      - name: "inositol phosphomannosylinositol phosphoceramide B (C26)"
      - compartment: "er"
      - formula: "C62H121NO25P2"
      - charge: 0
      - smiles: "CCCCCCCCCCCCCCCCCCCCCCCCCC(=O)NC(COP(=O)(O)OC1C(C(C(C(C1OC2C(C(C(C(O2)COP(=O)(O)OC3C(C(C(C(C3O)O)O)O)O)O)O)O)O)O)O)O)C(C(CCCCCCCCCCCCCCCC)O)O"
      - annotation: !!omap
          - chebi: "CHEBI:62719"
          - metanetx.chemical: "MNXM527421"
          - sbo: "SBO:0000247"
      - deltaG: 180.51
    - !!omap
      - id: "s_0871"
      - name: "inositol phosphomannosylinositol phosphoceramide B (C26)"
      - compartment: "g"
      - formula: "C62H121NO25P2"
      - charge: 0
      - smiles: "CCCCCCCCCCCCCCCCCCCCCCCCCC(=O)NC(COP(=O)(O)OC1C(C(C(C(C1OC2C(C(C(C(O2)COP(=O)(O)OC3C(C(C(C(C3O)O)O)O)O)O)O)O)O)O)O)O)C(C(CCCCCCCCCCCCCCCC)O)O"
      - annotation: !!omap
          - chebi: "CHEBI:62719"
          - metanetx.chemical: "MNXM527421"
          - sbo: "SBO:0000247"
      - deltaG: 180.51
    - !!omap
      - id: "s_0872"
      - name: "inositol phosphomannosylinositol phosphoceramide B (C26)"
      - compartment: "m"
      - formula: "C62H121NO25P2"
      - charge: 0
      - smiles: "CCCCCCCCCCCCCCCCCCCCCCCCCC(=O)NC(COP(=O)(O)OC1C(C(C(C(C1OC2C(C(C(C(O2)COP(=O)(O)OC3C(C(C(C(C3O)O)O)O)O)O)O)O)O)O)O)O)C(C(CCCCCCCCCCCCCCCC)O)O"
      - annotation: !!omap
          - chebi: "CHEBI:62719"
          - metanetx.chemical: "MNXM527421"
          - sbo: "SBO:0000247"
      - deltaG: 180.51
    - !!omap
      - id: "s_0873"
      - name: "inositol phosphomannosylinositol phosphoceramide B' (C24)"
      - compartment: "er"
      - formula: "C60H117NO25P2"
      - charge: 0
      - smiles: "CCCCCCCCCCCCCCCCCCCCCCC(C(=O)NC(COP(=O)(O)OC1C(C(C(C(C1OC2C(C(C(C(O2)COP(=O)(O)OC3C(C(C(C(C3O)O)O)O)O)O)O)O)O)O)O)O)C(CCCCCCCCCCCCCCC)O)O"
      - annotation: !!omap
          - chebi: "CHEBI:60441"
          - metanetx.chemical: "MNXM527422"
          - sbo: "SBO:0000247"
      - deltaG: 138
    - !!omap
      - id: "s_0874"
      - name: "inositol phosphomannosylinositol phosphoceramide B' (C24)"
      - compartment: "g"
      - formula: "C60H117NO25P2"
      - charge: 0
      - smiles: "CCCCCCCCCCCCCCCCCCCCCCC(C(=O)NC(COP(=O)(O)OC1C(C(C(C(C1OC2C(C(C(C(O2)COP(=O)(O)OC3C(C(C(C(C3O)O)O)O)O)O)O)O)O)O)O)O)C(CCCCCCCCCCCCCCC)O)O"
      - annotation: !!omap
          - chebi: "CHEBI:60441"
          - metanetx.chemical: "MNXM527422"
          - sbo: "SBO:0000247"
      - deltaG: 138
    - !!omap
      - id: "s_0875"
      - name: "inositol phosphomannosylinositol phosphoceramide B' (C24)"
      - compartment: "m"
      - formula: "C60H117NO25P2"
      - charge: 0
      - smiles: "CCCCCCCCCCCCCCCCCCCCCCC(C(=O)NC(COP(=O)(O)OC1C(C(C(C(C1OC2C(C(C(C(O2)COP(=O)(O)OC3C(C(C(C(C3O)O)O)O)O)O)O)O)O)O)O)O)C(CCCCCCCCCCCCCCC)O)O"
      - annotation: !!omap
          - chebi: "CHEBI:60441"
          - metanetx.chemical: "MNXM527422"
          - sbo: "SBO:0000247"
      - deltaG: 138
    - !!omap
      - id: "s_0876"
      - name: "inositol phosphomannosylinositol phosphoceramide B' (C26)"
      - compartment: "er"
      - formula: "C62H121NO25P2"
      - charge: 0
      - smiles: "CCCCCCCCCCCCCCCCCCCCCCCCC(C(=O)NC(COP(=O)(O)OC1C(C(C(C(C1OC2C(C(C(C(O2)COP(=O)(O)OC3C(C(C(C(C3O)O)O)O)O)O)O)O)O)O)O)O)C(CCCCCCCCCCCCCCCCC)O)O"
      - annotation: !!omap
          - chebi: "CHEBI:62717"
          - metanetx.chemical: "MNXM527418"
          - sbo: "SBO:0000247"
      - deltaG: 180.51
    - !!omap
      - id: "s_0877"
      - name: "inositol phosphomannosylinositol phosphoceramide B' (C26)"
      - compartment: "g"
      - formula: "C62H121NO25P2"
      - charge: 0
      - smiles: "CCCCCCCCCCCCCCCCCCCCCCCCC(C(=O)NC(COP(=O)(O)OC1C(C(C(C(C1OC2C(C(C(C(O2)COP(=O)(O)OC3C(C(C(C(C3O)O)O)O)O)O)O)O)O)O)O)O)C(CCCCCCCCCCCCCCCCC)O)O"
      - annotation: !!omap
          - chebi: "CHEBI:62717"
          - metanetx.chemical: "MNXM527418"
          - sbo: "SBO:0000247"
      - deltaG: 180.51
    - !!omap
      - id: "s_0878"
      - name: "inositol phosphomannosylinositol phosphoceramide B' (C26)"
      - compartment: "m"
      - formula: "C62H121NO25P2"
      - charge: 0
      - smiles: "CCCCCCCCCCCCCCCCCCCCCCCCC(C(=O)NC(COP(=O)(O)OC1C(C(C(C(C1OC2C(C(C(C(O2)COP(=O)(O)OC3C(C(C(C(C3O)O)O)O)O)O)O)O)O)O)O)O)C(CCCCCCCCCCCCCCCCC)O)O"
      - annotation: !!omap
          - chebi: "CHEBI:62717"
          - metanetx.chemical: "MNXM527418"
          - sbo: "SBO:0000247"
      - deltaG: 180.51
    - !!omap
      - id: "s_0879"
      - name: "inositol phosphomannosylinositol phosphoceramide C (C24)"
      - compartment: "er"
      - formula: "C60H117NO26P2"
      - charge: 0
      - smiles: "CCCCCCCCCCCCCCCCCCCCCCC(C(=O)NC(COP(=O)(O)OC1C(C(C(C(C1OC2C(C(C(C(O2)COP(=O)(O)OC3C(C(C(C(C3O)O)O)O)O)O)O)O)O)O)O)O)C(C(CCCCCCCCCCCCCC)O)O)O"
      - annotation: !!omap
          - chebi: "CHEBI:60445"
          - metanetx.chemical: "MNXM527424"
          - sbo: "SBO:0000247"
      - deltaG: 99.96
    - !!omap
      - id: "s_0880"
      - name: "inositol phosphomannosylinositol phosphoceramide C (C24)"
      - compartment: "g"
      - formula: "C60H117NO26P2"
      - charge: 0
      - smiles: "CCCCCCCCCCCCCCCCCCCCCCC(C(=O)NC(COP(=O)(O)OC1C(C(C(C(C1OC2C(C(C(C(O2)COP(=O)(O)OC3C(C(C(C(C3O)O)O)O)O)O)O)O)O)O)O)O)C(C(CCCCCCCCCCCCCC)O)O)O"
      - annotation: !!omap
          - chebi: "CHEBI:60445"
          - metanetx.chemical: "MNXM527424"
          - sbo: "SBO:0000247"
      - deltaG: 99.96
    - !!omap
      - id: "s_0881"
      - name: "inositol phosphomannosylinositol phosphoceramide C (C24)"
      - compartment: "m"
      - formula: "C60H117NO26P2"
      - charge: 0
      - smiles: "CCCCCCCCCCCCCCCCCCCCCCC(C(=O)NC(COP(=O)(O)OC1C(C(C(C(C1OC2C(C(C(C(O2)COP(=O)(O)OC3C(C(C(C(C3O)O)O)O)O)O)O)O)O)O)O)O)C(C(CCCCCCCCCCCCCC)O)O)O"
      - annotation: !!omap
          - chebi: "CHEBI:60445"
          - metanetx.chemical: "MNXM527424"
          - sbo: "SBO:0000247"
      - deltaG: 99.96
    - !!omap
      - id: "s_0882"
      - name: "inositol phosphomannosylinositol phosphoceramide C (C26)"
      - compartment: "er"
      - formula: "C62H121NO26P2"
      - charge: 0
      - smiles: "CCCCCCCCCCCCCCCCCCCCCCCCC(C(=O)NC(COP(=O)([O-])OC1C(C(C(C(C1OC2C(C(C(C(O2)COP(=O)([O-])OC3C(C(C(C(C3O)O)O)O)O)O)O)O)O)O)O)O)C(C(CCCCCCCCCCCCCC)O)O)O"
      - annotation: !!omap
          - chebi: "CHEBI:53005"
          - metanetx.chemical: "MNXM57469"
          - sbo: "SBO:0000247"
      - deltaG: 142.47
    - !!omap
      - id: "s_0883"
      - name: "inositol phosphomannosylinositol phosphoceramide C (C26)"
      - compartment: "g"
      - formula: "C62H121NO26P2"
      - charge: 0
      - smiles: "CCCCCCCCCCCCCCCCCCCCCCCCC(C(=O)NC(COP(=O)([O-])OC1C(C(C(C(C1OC2C(C(C(C(O2)COP(=O)([O-])OC3C(C(C(C(C3O)O)O)O)O)O)O)O)O)O)O)O)C(C(CCCCCCCCCCCCCC)O)O)O"
      - annotation: !!omap
          - chebi: "CHEBI:53005"
          - metanetx.chemical: "MNXM57469"
          - sbo: "SBO:0000247"
      - deltaG: 142.47
    - !!omap
      - id: "s_0884"
      - name: "inositol phosphomannosylinositol phosphoceramide C (C26)"
      - compartment: "m"
      - formula: "C62H121NO26P2"
      - charge: 0
      - smiles: "CCCCCCCCCCCCCCCCCCCCCCCCC(C(=O)NC(COP(=O)([O-])OC1C(C(C(C(C1OC2C(C(C(C(O2)COP(=O)([O-])OC3C(C(C(C(C3O)O)O)O)O)O)O)O)O)O)O)O)C(C(CCCCCCCCCCCCCC)O)O)O"
      - annotation: !!omap
          - chebi: "CHEBI:53005"
          - metanetx.chemical: "MNXM57469"
          - sbo: "SBO:0000247"
      - deltaG: 142.47
    - !!omap
      - id: "s_0885"
      - name: "inositol phosphomannosylinositol phosphoceramide D (C24)"
      - compartment: "er"
      - formula: "C60H117NO27P2"
      - charge: 0
      - smiles: "CCCCCCCCCCCCCCCCCCCCCC(C(C(=O)NC(COP(=O)(O)OC1C(C(C(C(C1OC2C(C(C(C(O2)COP(=O)(O)OC3C(C(C(C(C3O)O)O)O)O)O)O)O)O)O)O)O)C(C(CCCCCCCCCCCCCC)O)O)O)O"
      - annotation: !!omap
          - chebi: "CHEBI:60446"
          - metanetx.chemical: "MNXM527423"
          - sbo: "SBO:0000247"
      - deltaG: 61.92
    - !!omap
      - id: "s_0886"
      - name: "inositol phosphomannosylinositol phosphoceramide D (C24)"
      - compartment: "g"
      - formula: "C60H117NO27P2"
      - charge: 0
      - smiles: "CCCCCCCCCCCCCCCCCCCCCC(C(C(=O)NC(COP(=O)(O)OC1C(C(C(C(C1OC2C(C(C(C(O2)COP(=O)(O)OC3C(C(C(C(C3O)O)O)O)O)O)O)O)O)O)O)O)C(C(CCCCCCCCCCCCCC)O)O)O)O"
      - annotation: !!omap
          - chebi: "CHEBI:60446"
          - metanetx.chemical: "MNXM527423"
          - sbo: "SBO:0000247"
      - deltaG: 61.92
    - !!omap
      - id: "s_0887"
      - name: "inositol phosphomannosylinositol phosphoceramide D (C24)"
      - compartment: "m"
      - formula: "C60H117NO27P2"
      - charge: 0
      - smiles: "CCCCCCCCCCCCCCCCCCCCCC(C(C(=O)NC(COP(=O)(O)OC1C(C(C(C(C1OC2C(C(C(C(O2)COP(=O)(O)OC3C(C(C(C(C3O)O)O)O)O)O)O)O)O)O)O)O)C(C(CCCCCCCCCCCCCC)O)O)O)O"
      - annotation: !!omap
          - chebi: "CHEBI:60446"
          - metanetx.chemical: "MNXM527423"
          - sbo: "SBO:0000247"
      - deltaG: 61.92
    - !!omap
      - id: "s_0888"
      - name: "inositol phosphomannosylinositol phosphoceramide D (C26)"
      - compartment: "er"
      - formula: "C62H121NO27P2"
      - charge: 0
      - smiles: "CCCCCCCCCCCCCCCCCCCCCCCC(C(C(=O)NC(COP(=O)(O)OC1C(C(C(C(C1OC2C(C(C(C(O2)COP(=O)(O)OC3C(C(C(C(C3O)O)O)O)O)O)O)O)O)O)O)O)C(C(CCCCCCCCCCCCCCCC)O)O)O)O"
      - annotation: !!omap
          - chebi: "CHEBI:62684"
          - metanetx.chemical: "MNXM527420"
          - sbo: "SBO:0000247"
      - deltaG: 104.43
    - !!omap
      - id: "s_0889"
      - name: "inositol phosphomannosylinositol phosphoceramide D (C26)"
      - compartment: "g"
      - formula: "C62H121NO27P2"
      - charge: 0
      - smiles: "CCCCCCCCCCCCCCCCCCCCCCCC(C(C(=O)NC(COP(=O)(O)OC1C(C(C(C(C1OC2C(C(C(C(O2)COP(=O)(O)OC3C(C(C(C(C3O)O)O)O)O)O)O)O)O)O)O)O)C(C(CCCCCCCCCCCCCCCC)O)O)O)O"
      - annotation: !!omap
          - chebi: "CHEBI:62684"
          - metanetx.chemical: "MNXM527420"
          - sbo: "SBO:0000247"
      - deltaG: 104.43
    - !!omap
      - id: "s_0890"
      - name: "inositol phosphomannosylinositol phosphoceramide D (C26)"
      - compartment: "m"
      - formula: "C62H121NO27P2"
      - charge: 0
      - smiles: "CCCCCCCCCCCCCCCCCCCCCCCC(C(C(=O)NC(COP(=O)(O)OC1C(C(C(C(C1OC2C(C(C(C(O2)COP(=O)(O)OC3C(C(C(C(C3O)O)O)O)O)O)O)O)O)O)O)O)C(C(CCCCCCCCCCCCCCCC)O)O)O)O"
      - annotation: !!omap
          - chebi: "CHEBI:62684"
          - metanetx.chemical: "MNXM527420"
          - sbo: "SBO:0000247"
      - deltaG: 104.43
    - !!omap
      - id: "s_0892"
      - name: "inositol-P-ceramide backbone"
      - compartment: "g"
      - formula: "C6H12NO10P"
      - annotation: !!omap
          - chebi: "CHEBI:60245"
          - metanetx.chemical: "MNXM57446"
          - sbo: "SBO:0000649"
      - deltaG: 10000000
    - !!omap
      - id: "s_0894"
      - name: "inositol-P-ceramide A (C24)"
      - compartment: "er"
      - formula: "C48H96NO11P"
      - charge: 0
      - smiles: "CCCCCCCCCCCCCCCCCCCCCCCC(=O)NC(COP(=O)([O-])OC1C(C(C(C(C1O)O)O)O)O)C(CCCCCCCCCCCCCCC)O"
      - annotation: !!omap
          - chebi: "CHEBI:60402"
          - metanetx.chemical: "MNXM57476"
          - sbo: "SBO:0000247"
      - deltaG: 509.16
    - !!omap
      - id: "s_0895"
      - name: "inositol-P-ceramide A (C24)"
      - compartment: "g"
      - formula: "C48H96NO11P"
      - charge: 0
      - smiles: "CCCCCCCCCCCCCCCCCCCCCCCC(=O)NC(COP(=O)([O-])OC1C(C(C(C(C1O)O)O)O)O)C(CCCCCCCCCCCCCCC)O"
      - annotation: !!omap
          - chebi: "CHEBI:60402"
          - metanetx.chemical: "MNXM57476"
          - sbo: "SBO:0000247"
      - deltaG: 509.16
    - !!omap
      - id: "s_0896"
      - name: "inositol-P-ceramide A (C24)"
      - compartment: "m"
      - formula: "C48H96NO11P"
      - charge: 0
      - smiles: "CCCCCCCCCCCCCCCCCCCCCCCC(=O)NC(COP(=O)([O-])OC1C(C(C(C(C1O)O)O)O)O)C(CCCCCCCCCCCCCCC)O"
      - annotation: !!omap
          - chebi: "CHEBI:60402"
          - metanetx.chemical: "MNXM57476"
          - sbo: "SBO:0000247"
      - deltaG: 509.16
    - !!omap
      - id: "s_0897"
      - name: "inositol-P-ceramide A (C26)"
      - compartment: "er"
      - formula: "C50H100NO11P"
      - charge: 0
      - smiles: "CCCCCCCCCCCCCCCCCCCCCCCCCC(=O)NC(COP(=O)(O)OC1C(C(C(C(C1O)O)O)O)O)C(CCCCCCCCCCCCCCC)O"
      - annotation: !!omap
          - chebi: "CHEBI:60405"
          - metanetx.chemical: "MNXM57477"
          - sbo: "SBO:0000247"
      - deltaG: 551.67
    - !!omap
      - id: "s_0898"
      - name: "inositol-P-ceramide A (C26)"
      - compartment: "g"
      - formula: "C50H100NO11P"
      - charge: 0
      - smiles: "CCCCCCCCCCCCCCCCCCCCCCCCCC(=O)NC(COP(=O)(O)OC1C(C(C(C(C1O)O)O)O)O)C(CCCCCCCCCCCCCCC)O"
      - annotation: !!omap
          - chebi: "CHEBI:60405"
          - metanetx.chemical: "MNXM57477"
          - sbo: "SBO:0000247"
      - deltaG: 551.67
    - !!omap
      - id: "s_0899"
      - name: "inositol-P-ceramide A (C26)"
      - compartment: "m"
      - formula: "C50H100NO11P"
      - charge: 0
      - smiles: "CCCCCCCCCCCCCCCCCCCCCCCCCC(=O)NC(COP(=O)(O)OC1C(C(C(C(C1O)O)O)O)O)C(CCCCCCCCCCCCCCC)O"
      - annotation: !!omap
          - chebi: "CHEBI:60405"
          - metanetx.chemical: "MNXM57477"
          - sbo: "SBO:0000247"
      - deltaG: 551.67
    - !!omap
      - id: "s_0900"
      - name: "inositol-P-ceramide B (C24)"
      - compartment: "er"
      - formula: "C48H96NO12P"
      - charge: 0
      - smiles: "CCCCCCCCCCCCCCCCCCCCCCCC(=O)NC(COP(=O)(O)OC1C(C(C(C(C1O)O)O)O)O)C(C(CCCCCCCCCCCCCC)O)O"
      - annotation: !!omap
          - chebi: "CHEBI:60410"
          - metanetx.chemical: "MNXM527429"
          - sbo: "SBO:0000247"
      - deltaG: 471.12
    - !!omap
      - id: "s_0901"
      - name: "inositol-P-ceramide B (C24)"
      - compartment: "g"
      - formula: "C48H96NO12P"
      - charge: 0
      - smiles: "CCCCCCCCCCCCCCCCCCCCCCCC(=O)NC(COP(=O)(O)OC1C(C(C(C(C1O)O)O)O)O)C(C(CCCCCCCCCCCCCC)O)O"
      - annotation: !!omap
          - chebi: "CHEBI:60410"
          - metanetx.chemical: "MNXM527429"
          - sbo: "SBO:0000247"
      - deltaG: 471.12
    - !!omap
      - id: "s_0902"
      - name: "inositol-P-ceramide B (C24)"
      - compartment: "m"
      - formula: "C48H96NO12P"
      - charge: 0
      - smiles: "CCCCCCCCCCCCCCCCCCCCCCCC(=O)NC(COP(=O)(O)OC1C(C(C(C(C1O)O)O)O)O)C(C(CCCCCCCCCCCCCC)O)O"
      - annotation: !!omap
          - chebi: "CHEBI:60410"
          - metanetx.chemical: "MNXM527429"
          - sbo: "SBO:0000247"
      - deltaG: 471.12
    - !!omap
      - id: "s_0903"
      - name: "inositol-P-ceramide B (C26)"
      - compartment: "er"
      - formula: "C50H100NO12P"
      - charge: 0
      - smiles: "CCCCCCCCCCCCCCCCCCCCCCCCCC(=O)NC(COP(=O)(O)OC1C(C(C(C(C1O)O)O)O)O)C(C(CCCCCCCCCCCCCC)O)O"
      - annotation: !!omap
          - chebi: "CHEBI:60412"
          - metanetx.chemical: "MNXM527430"
          - sbo: "SBO:0000247"
      - deltaG: 513.63
    - !!omap
      - id: "s_0904"
      - name: "inositol-P-ceramide B (C26)"
      - compartment: "g"
      - formula: "C50H100NO12P"
      - charge: 0
      - smiles: "CCCCCCCCCCCCCCCCCCCCCCCCCC(=O)NC(COP(=O)(O)OC1C(C(C(C(C1O)O)O)O)O)C(C(CCCCCCCCCCCCCC)O)O"
      - annotation: !!omap
          - chebi: "CHEBI:60412"
          - metanetx.chemical: "MNXM527430"
          - sbo: "SBO:0000247"
      - deltaG: 513.63
    - !!omap
      - id: "s_0905"
      - name: "inositol-P-ceramide B (C26)"
      - compartment: "m"
      - formula: "C50H100NO12P"
      - charge: 0
      - smiles: "CCCCCCCCCCCCCCCCCCCCCCCCCC(=O)NC(COP(=O)(O)OC1C(C(C(C(C1O)O)O)O)O)C(C(CCCCCCCCCCCCCC)O)O"
      - annotation: !!omap
          - chebi: "CHEBI:60412"
          - metanetx.chemical: "MNXM527430"
          - sbo: "SBO:0000247"
      - deltaG: 513.63
    - !!omap
      - id: "s_0906"
      - name: "inositol-P-ceramide B' (C24)"
      - compartment: "er"
      - formula: "C48H96NO12P"
      - charge: 0
      - smiles: "CCCCCCCCCCCCCCCCCCCCCCC(C(=O)NC(COP(=O)(O)OC1C(C(C(C(C1O)O)O)O)O)C(CCCCCCCCCCCCCCC)O)O"
      - annotation: !!omap
          - chebi: "CHEBI:60406"
          - metanetx.chemical: "MNXM57474"
          - sbo: "SBO:0000247"
      - deltaG: 471.12
    - !!omap
      - id: "s_0907"
      - name: "inositol-P-ceramide B' (C24)"
      - compartment: "g"
      - formula: "C48H96NO12P"
      - charge: 0
      - smiles: "CCCCCCCCCCCCCCCCCCCCCCC(C(=O)NC(COP(=O)(O)OC1C(C(C(C(C1O)O)O)O)O)C(CCCCCCCCCCCCCCC)O)O"
      - annotation: !!omap
          - chebi: "CHEBI:60406"
          - metanetx.chemical: "MNXM57474"
          - sbo: "SBO:0000247"
      - deltaG: 471.12
    - !!omap
      - id: "s_0908"
      - name: "inositol-P-ceramide B' (C24)"
      - compartment: "m"
      - formula: "C48H96NO12P"
      - charge: 0
      - smiles: "CCCCCCCCCCCCCCCCCCCCCCC(C(=O)NC(COP(=O)(O)OC1C(C(C(C(C1O)O)O)O)O)C(CCCCCCCCCCCCCCC)O)O"
      - annotation: !!omap
          - chebi: "CHEBI:60406"
          - metanetx.chemical: "MNXM57474"
          - sbo: "SBO:0000247"
      - deltaG: 471.12
    - !!omap
      - id: "s_0909"
      - name: "inositol-P-ceramide B' (C26)"
      - compartment: "er"
      - formula: "C50H100NO12P"
      - charge: 0
      - smiles: "CCCCCCCCCCCCCCCCCCCCCCCCC(C(=O)NC(COP(=O)(O)OC1C(C(C(C(C1O)O)O)O)O)C(CCCCCCCCCCCCCCC)O)O"
      - annotation: !!omap
          - chebi: "CHEBI:60409"
          - metanetx.chemical: "MNXM57475"
          - sbo: "SBO:0000247"
      - deltaG: 513.63
    - !!omap
      - id: "s_0910"
      - name: "inositol-P-ceramide B' (C26)"
      - compartment: "g"
      - formula: "C50H100NO12P"
      - charge: 0
      - smiles: "CCCCCCCCCCCCCCCCCCCCCCCCC(C(=O)NC(COP(=O)(O)OC1C(C(C(C(C1O)O)O)O)O)C(CCCCCCCCCCCCCCC)O)O"
      - annotation: !!omap
          - chebi: "CHEBI:60409"
          - metanetx.chemical: "MNXM57475"
          - sbo: "SBO:0000247"
      - deltaG: 513.63
    - !!omap
      - id: "s_0911"
      - name: "inositol-P-ceramide B' (C26)"
      - compartment: "m"
      - formula: "C50H100NO12P"
      - charge: 0
      - smiles: "CCCCCCCCCCCCCCCCCCCCCCCCC(C(=O)NC(COP(=O)(O)OC1C(C(C(C(C1O)O)O)O)O)C(CCCCCCCCCCCCCCC)O)O"
      - annotation: !!omap
          - chebi: "CHEBI:60409"
          - metanetx.chemical: "MNXM57475"
          - sbo: "SBO:0000247"
      - deltaG: 513.63
    - !!omap
      - id: "s_0912"
      - name: "inositol-P-ceramide C (C24)"
      - compartment: "er"
      - formula: "C48H96NO13P"
      - charge: 0
      - smiles: "CCCCCCCCCCCCCCCCCCCCCCC(C(=O)NC(COP(=O)(O)OC1C(C(C(C(C1O)O)O)O)O)C(C(CCCCCCCCCCCCCC)O)O)O"
      - annotation: !!omap
          - chebi: "CHEBI:60413"
          - metanetx.chemical: "MNXM527428"
          - sbo: "SBO:0000247"
      - deltaG: 433.08
    - !!omap
      - id: "s_0913"
      - name: "inositol-P-ceramide C (C24)"
      - compartment: "g"
      - formula: "C48H96NO13P"
      - charge: 0
      - smiles: "CCCCCCCCCCCCCCCCCCCCCCC(C(=O)NC(COP(=O)(O)OC1C(C(C(C(C1O)O)O)O)O)C(C(CCCCCCCCCCCCCC)O)O)O"
      - annotation: !!omap
          - chebi: "CHEBI:60413"
          - metanetx.chemical: "MNXM527428"
          - sbo: "SBO:0000247"
      - deltaG: 433.08
    - !!omap
      - id: "s_0914"
      - name: "inositol-P-ceramide C (C24)"
      - compartment: "m"
      - formula: "C48H96NO13P"
      - charge: 0
      - smiles: "CCCCCCCCCCCCCCCCCCCCCCC(C(=O)NC(COP(=O)(O)OC1C(C(C(C(C1O)O)O)O)O)C(C(CCCCCCCCCCCCCC)O)O)O"
      - annotation: !!omap
          - chebi: "CHEBI:60413"
          - metanetx.chemical: "MNXM527428"
          - sbo: "SBO:0000247"
      - deltaG: 433.08
    - !!omap
      - id: "s_0915"
      - name: "inositol-P-ceramide C (C26)"
      - compartment: "er"
      - formula: "C50H100NO13P"
      - charge: 0
      - smiles: "CCCCCCCCCCCCCCCCCCCCCCCCC(C(=O)NC(COP(=O)(O)OC1C(C(C(C(C1O)O)O)O)O)C(C(CCCCCCCCCCCCCC)O)O)O"
      - annotation: !!omap
          - chebi: "CHEBI:53004"
          - metanetx.chemical: "MNXM57486"
          - sbo: "SBO:0000247"
      - deltaG: 475.59
    - !!omap
      - id: "s_0916"
      - name: "inositol-P-ceramide C (C26)"
      - compartment: "g"
      - formula: "C50H100NO13P"
      - charge: 0
      - smiles: "CCCCCCCCCCCCCCCCCCCCCCCCC(C(=O)NC(COP(=O)(O)OC1C(C(C(C(C1O)O)O)O)O)C(C(CCCCCCCCCCCCCC)O)O)O"
      - annotation: !!omap
          - chebi: "CHEBI:53004"
          - metanetx.chemical: "MNXM57486"
          - sbo: "SBO:0000247"
      - deltaG: 475.59
    - !!omap
      - id: "s_0917"
      - name: "inositol-P-ceramide C (C26)"
      - compartment: "m"
      - formula: "C50H100NO13P"
      - charge: 0
      - smiles: "CCCCCCCCCCCCCCCCCCCCCCCCC(C(=O)NC(COP(=O)(O)OC1C(C(C(C(C1O)O)O)O)O)C(C(CCCCCCCCCCCCCC)O)O)O"
      - annotation: !!omap
          - chebi: "CHEBI:53004"
          - metanetx.chemical: "MNXM57486"
          - sbo: "SBO:0000247"
      - deltaG: 475.59
    - !!omap
      - id: "s_0918"
      - name: "inositol-P-ceramide D (C24)"
      - compartment: "er"
      - formula: "C48H96NO14P"
      - charge: 0
      - smiles: "CCCCCCCCCCCCCCCCCCCCCC(C(C(=O)NC(COP(=O)([O-])OC1C(C(C(C(C1O)O)O)O)O)C(C(CCCCCCCCCCCCCC)O)O)O)O"
      - annotation: !!omap
          - chebi: "CHEBI:60414"
          - metanetx.chemical: "MNXM527426"
          - sbo: "SBO:0000247"
      - deltaG: 395.04
    - !!omap
      - id: "s_0919"
      - name: "inositol-P-ceramide D (C24)"
      - compartment: "g"
      - formula: "C48H96NO14P"
      - charge: 0
      - smiles: "CCCCCCCCCCCCCCCCCCCCCC(C(C(=O)NC(COP(=O)([O-])OC1C(C(C(C(C1O)O)O)O)O)C(C(CCCCCCCCCCCCCC)O)O)O)O"
      - annotation: !!omap
          - chebi: "CHEBI:60414"
          - metanetx.chemical: "MNXM527426"
          - sbo: "SBO:0000247"
      - deltaG: 395.04
    - !!omap
      - id: "s_0920"
      - name: "inositol-P-ceramide D (C24)"
      - compartment: "m"
      - formula: "C48H96NO14P"
      - charge: 0
      - smiles: "CCCCCCCCCCCCCCCCCCCCCC(C(C(=O)NC(COP(=O)([O-])OC1C(C(C(C(C1O)O)O)O)O)C(C(CCCCCCCCCCCCCC)O)O)O)O"
      - annotation: !!omap
          - chebi: "CHEBI:60414"
          - metanetx.chemical: "MNXM527426"
          - sbo: "SBO:0000247"
      - deltaG: 395.04
    - !!omap
      - id: "s_0921"
      - name: "inositol-P-ceramide D (C26)"
      - compartment: "er"
      - formula: "C50H100NO14P"
      - charge: 0
      - smiles: "CCCCCCCCCCCCCCCCCCCCCCCC(C(C(=O)NC(COP(=O)([O-])OC1C(C(C(C(C1O)O)O)O)O)C(C(CCCCCCCCCCCCCC)O)O)O)O"
      - annotation: !!omap
          - chebi: "CHEBI:60415"
          - metanetx.chemical: "MNXM527427"
          - sbo: "SBO:0000247"
      - deltaG: 437.55
    - !!omap
      - id: "s_0922"
      - name: "inositol-P-ceramide D (C26)"
      - compartment: "g"
      - formula: "C50H100NO14P"
      - charge: 0
      - smiles: "CCCCCCCCCCCCCCCCCCCCCCCC(C(C(=O)NC(COP(=O)([O-])OC1C(C(C(C(C1O)O)O)O)O)C(C(CCCCCCCCCCCCCC)O)O)O)O"
      - annotation: !!omap
          - chebi: "CHEBI:60415"
          - metanetx.chemical: "MNXM527427"
          - sbo: "SBO:0000247"
      - deltaG: 437.55
    - !!omap
      - id: "s_0923"
      - name: "inositol-P-ceramide D (C26)"
      - compartment: "m"
      - formula: "C50H100NO14P"
      - charge: 0
      - smiles: "CCCCCCCCCCCCCCCCCCCCCCCC(C(C(=O)NC(COP(=O)([O-])OC1C(C(C(C(C1O)O)O)O)O)C(C(CCCCCCCCCCCCCC)O)O)O)O"
      - annotation: !!omap
          - chebi: "CHEBI:60415"
          - metanetx.chemical: "MNXM527427"
          - sbo: "SBO:0000247"
      - deltaG: 437.55
    - !!omap
      - id: "s_0924"
      - name: "iron(2+)"
      - compartment: "c"
      - formula: "Fe"
      - charge: 2
      - smiles: "[Fe+2]"
      - annotation: !!omap
          - bigg.metabolite: "fe2"
          - chebi: "CHEBI:29033"
          - kegg.compound: "C14818"
          - metanetx.chemical: "MNXM111"
          - sbo: "SBO:0000247"
      - deltaG: -48.55
    - !!omap
      - id: "s_0925"
      - name: "iron(2+)"
      - compartment: "e"
      - formula: "Fe"
      - charge: 2
      - smiles: "[Fe+2]"
      - annotation: !!omap
          - bigg.metabolite: "fe2"
          - chebi: "CHEBI:29033"
          - kegg.compound: "C14818"
          - metanetx.chemical: "MNXM111"
          - sbo: "SBO:0000247"
      - deltaG: -48.55
    - !!omap
      - id: "s_0926"
      - name: "iron(2+)"
      - compartment: "m"
      - formula: "Fe"
      - charge: 2
      - smiles: "[Fe+2]"
      - annotation: !!omap
          - bigg.metabolite: "fe2"
          - chebi: "CHEBI:29033"
          - kegg.compound: "C14818"
          - metanetx.chemical: "MNXM111"
          - sbo: "SBO:0000247"
      - deltaG: -48.55
    - !!omap
      - id: "s_0927"
      - name: "isoamyl acetate"
      - compartment: "c"
      - formula: "C7H14O2"
      - charge: 0
      - smiles: "CC(C)CCOC(=O)C"
      - annotation: !!omap
          - bigg.metabolite: "iamac"
          - chebi: "CHEBI:31725"
          - kegg.compound: "C12296"
          - metanetx.chemical: "MNXM7451"
          - sbo: "SBO:0000247"
      - deltaG: 58.39
    - !!omap
      - id: "s_0928"
      - name: "isoamyl acetate"
      - compartment: "e"
      - formula: "C7H14O2"
      - charge: 0
      - smiles: "CC(C)CCOC(=O)C"
      - annotation: !!omap
          - bigg.metabolite: "iamac"
          - chebi: "CHEBI:31725"
          - kegg.compound: "C12296"
          - metanetx.chemical: "MNXM7451"
          - sbo: "SBO:0000247"
      - deltaG: 58.39
    - !!omap
      - id: "s_0929"
      - name: "isoamylol"
      - compartment: "c"
      - formula: "C5H12O"
      - charge: 0
      - smiles: "CC(C)CCO"
      - annotation: !!omap
          - bigg.metabolite: "iamoh"
          - chebi: "CHEBI:15837"
          - kegg.compound: "C07328"
          - metanetx.chemical: "MNXM1670"
          - sbo: "SBO:0000247"
      - deltaG: 76.36
    - !!omap
      - id: "s_0930"
      - name: "isoamylol"
      - compartment: "e"
      - formula: "C5H12O"
      - charge: 0
      - smiles: "CC(C)CCO"
      - annotation: !!omap
          - bigg.metabolite: "iamoh"
          - chebi: "CHEBI:15837"
          - kegg.compound: "C07328"
          - metanetx.chemical: "MNXM1670"
          - sbo: "SBO:0000247"
      - deltaG: 76.36
    - !!omap
      - id: "s_0931"
      - name: "isoamylol"
      - compartment: "m"
      - formula: "C5H12O"
      - charge: 0
      - smiles: "CC(C)CCO"
      - annotation: !!omap
          - bigg.metabolite: "iamoh"
          - chebi: "CHEBI:15837"
          - kegg.compound: "C07328"
          - metanetx.chemical: "MNXM1670"
          - sbo: "SBO:0000247"
      - deltaG: 76.36
    - !!omap
      - id: "s_0932"
      - name: "isobutanol"
      - compartment: "c"
      - formula: "C4H10O"
      - charge: 0
      - smiles: "CC(C)CO"
      - annotation: !!omap
          - bigg.metabolite: "ibutoh"
          - chebi: "CHEBI:46645"
          - kegg.compound: "C14710"
          - metanetx.chemical: "MNXM5188"
          - sbo: "SBO:0000247"
      - deltaG: 55.27
    - !!omap
      - id: "s_0933"
      - name: "isobutanol"
      - compartment: "e"
      - formula: "C4H10O"
      - charge: 0
      - smiles: "CC(C)CO"
      - annotation: !!omap
          - bigg.metabolite: "ibutoh"
          - chebi: "CHEBI:46645"
          - kegg.compound: "C14710"
          - metanetx.chemical: "MNXM5188"
          - sbo: "SBO:0000247"
      - deltaG: 55.27
    - !!omap
      - id: "s_0934"
      - name: "isobutanol"
      - compartment: "m"
      - formula: "C4H10O"
      - charge: 0
      - smiles: "CC(C)CO"
      - annotation: !!omap
          - bigg.metabolite: "ibutoh"
          - chebi: "CHEBI:46645"
          - kegg.compound: "C14710"
          - metanetx.chemical: "MNXM5188"
          - sbo: "SBO:0000247"
      - deltaG: 55.27
    - !!omap
      - id: "s_0935"
      - name: "isobutyl acetate"
      - compartment: "c"
      - formula: "C6H12O2"
      - charge: 0
      - smiles: "CC(C)COC(=O)C"
      - annotation: !!omap
          - bigg.metabolite: "ibutac"
          - chebi: "CHEBI:50569"
          - metanetx.chemical: "MNXM57835"
          - sbo: "SBO:0000247"
      - deltaG: 37.29
    - !!omap
      - id: "s_0936"
      - name: "isobutyl acetate"
      - compartment: "e"
      - formula: "C6H12O2"
      - charge: 0
      - smiles: "CC(C)COC(=O)C"
      - annotation: !!omap
          - bigg.metabolite: "ibutac"
          - chebi: "CHEBI:50569"
          - metanetx.chemical: "MNXM57835"
          - sbo: "SBO:0000247"
      - deltaG: 37.29
    - !!omap
      - id: "s_0937"
      - name: "isobutyraldehyde"
      - compartment: "c"
      - formula: "C4H8O"
      - charge: 0
      - smiles: "CC(C)C=O"
      - annotation: !!omap
          - bigg.metabolite: "2mppal"
          - chebi: "CHEBI:48943"
          - metanetx.chemical: "MNXM5189"
          - sbo: "SBO:0000247"
      - deltaG: 45.27
    - !!omap
      - id: "s_0938"
      - name: "isobutyraldehyde"
      - compartment: "e"
      - formula: "C4H8O"
      - charge: 0
      - smiles: "CC(C)C=O"
      - annotation: !!omap
          - bigg.metabolite: "2mppal"
          - chebi: "CHEBI:48943"
          - metanetx.chemical: "MNXM5189"
          - sbo: "SBO:0000247"
      - deltaG: 45.27
    - !!omap
      - id: "s_0939"
      - name: "isobutyraldehyde"
      - compartment: "m"
      - formula: "C4H8O"
      - charge: 0
      - smiles: "CC(C)C=O"
      - annotation: !!omap
          - bigg.metabolite: "2mppal"
          - chebi: "CHEBI:48943"
          - metanetx.chemical: "MNXM5189"
          - sbo: "SBO:0000247"
      - deltaG: 45.27
    - !!omap
      - id: "s_0940"
      - name: "isocitrate"
      - compartment: "c"
      - formula: "C6H5O7"
      - charge: -3
      - smiles: "C(C(C(C(=O)O)O)C(=O)O)C(=O)O"
      - annotation: !!omap
          - bigg.metabolite: "icit"
          - chebi: "CHEBI:16087"
          - kegg.compound: "C00311"
          - metanetx.chemical: "MNXM89661"
          - sbo: "SBO:0000247"
      - deltaG: -232.08
    - !!omap
      - id: "s_0941"
      - name: "isocitrate"
      - compartment: "m"
      - formula: "C6H5O7"
      - charge: -3
      - smiles: "C(C(C(C(=O)O)O)C(=O)O)C(=O)O"
      - annotation: !!omap
          - bigg.metabolite: "icit"
          - chebi: "CHEBI:16087"
          - kegg.compound: "C00311"
          - metanetx.chemical: "MNXM89661"
          - sbo: "SBO:0000247"
      - deltaG: -232.08
    - !!omap
      - id: "s_0942"
      - name: "isocitrate"
      - compartment: "p"
      - formula: "C6H5O7"
      - charge: -3
      - smiles: "C(C(C(C(=O)O)O)C(=O)O)C(=O)O"
      - annotation: !!omap
          - bigg.metabolite: "icit"
          - chebi: "CHEBI:16087"
          - kegg.compound: "C00311"
          - metanetx.chemical: "MNXM89661"
          - sbo: "SBO:0000247"
      - deltaG: -232.08
    - !!omap
      - id: "s_0943"
      - name: "isopentenyl diphosphate"
      - compartment: "c"
      - formula: "C5H9O7P2"
      - charge: -3
      - smiles: "CC(=C)CCOP(=O)(O)OP(=O)(O)O"
      - annotation: !!omap
          - bigg.metabolite: "ipdp"
          - chebi: "CHEBI:128769"
          - kegg.compound: "C00129"
          - metanetx.chemical: "MNXM83"
          - sbo: "SBO:0000247"
      - deltaG: -344.02
    - !!omap
      - id: "s_0944"
      - name: "isopentenyl diphosphate"
      - compartment: "lp"
      - formula: "C5H9O7P2"
      - charge: -3
      - smiles: "CC(=C)CCOP(=O)(O)OP(=O)(O)O"
      - annotation: !!omap
          - bigg.metabolite: "ipdp"
          - chebi: "CHEBI:128769"
          - kegg.compound: "C00129"
          - metanetx.chemical: "MNXM83"
          - sbo: "SBO:0000247"
      - deltaG: -344.02
    - !!omap
      - id: "s_0945"
      - name: "isopentenyl diphosphate"
      - compartment: "m"
      - formula: "C5H9O7P2"
      - charge: -3
      - smiles: "CC(=C)CCOP(=O)(O)OP(=O)(O)O"
      - annotation: !!omap
          - bigg.metabolite: "ipdp"
          - chebi: "CHEBI:128769"
          - kegg.compound: "C00129"
          - metanetx.chemical: "MNXM83"
          - sbo: "SBO:0000247"
      - deltaG: -344.02
    - !!omap
      - id: "s_0946"
      - name: "itaconate"
      - compartment: "m"
      - formula: "C5H4O4"
      - charge: -2
      - smiles: "C=C(CC(=O)O)C(=O)O"
      - annotation: !!omap
          - bigg.metabolite: "itacon"
          - chebi: "CHEBI:17240"
          - kegg.compound: "C00490"
          - metanetx.chemical: "MNXM1747"
          - sbo: "SBO:0000247"
      - deltaG: -102.66
    - !!omap
      - id: "s_0949"
      - name: "itaconyl-CoA"
      - compartment: "m"
      - formula: "C26H35N7O19P3S"
      - charge: -5
      - smiles: "CC(C)(COP(=O)(O)OP(=O)(O)OCC1C(C(C(O1)N2C=NC3=C(N=CN=C32)N)O)OP(=O)(O)O)C(C(=O)NCCC(=O)NCCSC(=O)CC(=C)C(=O)O)O"
      - annotation: !!omap
          - bigg.metabolite: "itaccoa"
          - chebi: "CHEBI:57381"
          - kegg.compound: "C00531"
          - metanetx.chemical: "MNXM1671"
          - sbo: "SBO:0000247"
      - deltaG: -493.53
    - !!omap
      - id: "s_0950"
      - name: "ITP"
      - compartment: "c"
      - formula: "C10H11N4O14P3"
      - charge: -4
      - smiles: "C1=NC2=C(C(=O)N1)N=CN2C3C(C(C(O3)COP(=O)(O)OP(=O)(O)OP(=O)(O)O)O)O"
      - annotation: !!omap
          - bigg.metabolite: "itp"
          - chebi: "CHEBI:61402"
          - kegg.compound: "C00081"
          - metanetx.chemical: "MNXM423"
          - sbo: "SBO:0000247"
      - deltaG: -633.42
    - !!omap
      - id: "s_0951"
      - name: "keto-phenylpyruvate"
      - compartment: "c"
      - formula: "C9H7O3"
      - charge: -1
      - smiles: "C1=CC=C(C=C1)CC(=O)C(=O)[O-]"
      - annotation: !!omap
          - bigg.metabolite: "phpyr"
          - chebi: "CHEBI:18005"
          - kegg.compound: "C00166"
          - metanetx.chemical: "MNXM162242"
          - sbo: "SBO:0000247"
      - deltaG: -7.86
    - !!omap
      - id: "s_0952"
      - name: "L-2-amino-3-oxobutanoate"
      - compartment: "c"
      - formula: "C4H7NO3"
      - charge: 0
      - smiles: "CC(=O)C(C(=O)O)N"
      - annotation: !!omap
          - bigg.metabolite: "2aobut"
          - chebi: "CHEBI:40673"
          - kegg.compound: "C03508"
          - metanetx.chemical: "MNXM114087"
          - sbo: "SBO:0000247"
      - deltaG: -48.16
    - !!omap
      - id: "s_0953"
      - name: "L-2-aminoadipate"
      - compartment: "c"
      - formula: "C6H10NO4"
      - charge: -1
      - smiles: "C(CC(C(=O)O)N)CC(=O)O"
      - annotation: !!omap
          - bigg.metabolite: "L2aadp"
          - chebi: "CHEBI:58672"
          - kegg.compound: "C00956"
          - metanetx.chemical: "MNXM268"
          - sbo: "SBO:0000247"
      - deltaG: -69.63
    - !!omap
      - id: "s_0954"
      - name: "L-4-hydroxyglutamic semialdehyde"
      - compartment: "m"
      - formula: "C5H9NO4"
      - charge: 0
      - smiles: "C(C(C=O)O)C(C(=O)O)N"
      - annotation: !!omap
          - bigg.metabolite: "4hglusa"
          - chebi: "CHEBI:27809"
          - kegg.compound: "C05938"
          - metanetx.chemical: "MNXM2687"
          - sbo: "SBO:0000247"
      - deltaG: -55.7
    - !!omap
      - id: "s_0955"
      - name: "L-alanine"
      - compartment: "c"
      - formula: "C3H7NO2"
      - charge: 0
      - smiles: "CC(C(=O)O)N"
      - annotation: !!omap
          - bigg.metabolite: "ala__L"
          - chebi: "CHEBI:16977"
          - kegg.compound: "C00041"
          - metanetx.chemical: "MNXM32"
          - sbo: "SBO:0000247"
      - deltaG: -19.76
    - !!omap
      - id: "s_0956"
      - name: "L-alanine"
      - compartment: "e"
      - formula: "C3H7NO2"
      - charge: 0
      - smiles: "CC(C(=O)O)N"
      - annotation: !!omap
          - bigg.metabolite: "ala__L"
          - chebi: "CHEBI:16977"
          - kegg.compound: "C00041"
          - metanetx.chemical: "MNXM32"
          - sbo: "SBO:0000247"
      - deltaG: -19.76
    - !!omap
      - id: "s_0957"
      - name: "L-alanine"
      - compartment: "m"
      - formula: "C3H7NO2"
      - charge: 0
      - smiles: "CC(C(=O)O)N"
      - annotation: !!omap
          - bigg.metabolite: "ala__L"
          - chebi: "CHEBI:16977"
          - kegg.compound: "C00041"
          - metanetx.chemical: "MNXM32"
          - sbo: "SBO:0000247"
      - deltaG: -19.76
    - !!omap
      - id: "s_0958"
      - name: "L-allothreonine"
      - compartment: "c"
      - formula: "C4H9NO3"
      - charge: 0
      - smiles: "CC(C(C(=O)O)N)O"
      - annotation: !!omap
          - bigg.metabolite: "athr__L"
          - chebi: "CHEBI:28718"
          - kegg.compound: "C05519"
          - metanetx.chemical: "MNXM2125"
          - sbo: "SBO:0000247"
      - deltaG: -36.71
    - !!omap
      - id: "s_0959"
      - name: "L-allysine"
      - compartment: "c"
      - formula: "C6H11NO3"
      - charge: 0
      - smiles: "C(CC=O)CC(C(=O)O)N"
      - annotation: !!omap
          - bigg.metabolite: "L2aadp6sa"
          - chebi: "CHEBI:17917"
          - kegg.compound: "C04076"
          - metanetx.chemical: "MNXM89905"
          - sbo: "SBO:0000247"
      - deltaG: -2.7
    - !!omap
      - id: "s_0960"
      - name: "L-alpha-formylglycine"
      - compartment: "c"
      - formula: "C3H5NO3"
      - charge: 0
      - smiles: "C(=O)C(C(=O)O)N"
      - annotation: !!omap
          - bigg.metabolite: "2amsa"
          - chebi: "CHEBI:58671"
          - kegg.compound: "C11822"
          - metanetx.chemical: "MNXM2124"
          - sbo: "SBO:0000247"
      - deltaG: -65.99
    - !!omap
      - id: "s_0961"
      - name: "L-arabinitol"
      - compartment: "c"
      - formula: "C5H12O5"
      - charge: 0
      - smiles: "C(C(C(C(CO)O)O)O)O"
      - annotation: !!omap
          - bigg.metabolite: "abt"
          - chebi: "CHEBI:18403"
          - kegg.compound: "C00532"
          - metanetx.chemical: "MNXM801"
          - sbo: "SBO:0000247"
      - deltaG: -72.18
    - !!omap
      - id: "s_0962"
      - name: "L-arabinitol"
      - compartment: "e"
      - formula: "C5H12O5"
      - charge: 0
      - smiles: "C(C(C(C(CO)O)O)O)O"
      - annotation: !!omap
          - bigg.metabolite: "abt"
          - chebi: "CHEBI:18403"
          - kegg.compound: "C00532"
          - metanetx.chemical: "MNXM801"
          - sbo: "SBO:0000247"
      - deltaG: -72.18
    - !!omap
      - id: "s_0963"
      - name: "L-arabinose"
      - compartment: "c"
      - formula: "C5H10O5"
      - charge: 0
      - smiles: "C1C(C(C(C(O1)O)O)O)O"
      - annotation: !!omap
          - bigg.metabolite: "arab__L"
          - chebi: "CHEBI:30849"
          - kegg.compound: "C00259"
          - metanetx.chemical: "MNXM461"
          - sbo: "SBO:0000247"
      - deltaG: -82.69
    - !!omap
      - id: "s_0964"
      - name: "L-arabinose"
      - compartment: "e"
      - formula: "C5H10O5"
      - charge: 0
      - smiles: "C1C(C(C(C(O1)O)O)O)O"
      - annotation: !!omap
          - bigg.metabolite: "arab__L"
          - chebi: "CHEBI:30849"
          - kegg.compound: "C00259"
          - metanetx.chemical: "MNXM461"
          - sbo: "SBO:0000247"
      - deltaG: -82.69
    - !!omap
      - id: "s_0965"
      - name: "L-arginine"
      - compartment: "c"
      - formula: "C6H15N4O2"
      - charge: 1
      - smiles: "C(CC(C(=O)O)N)CN=C(N)N"
      - annotation: !!omap
          - bigg.metabolite: "arg__L"
          - chebi: "CHEBI:32682"
          - kegg.compound: "C00062"
          - metanetx.chemical: "MNXM70"
          - sbo: "SBO:0000247"
      - deltaG: 72.09
    - !!omap
      - id: "s_0966"
      - name: "L-arginine"
      - compartment: "e"
      - formula: "C6H15N4O2"
      - charge: 1
      - smiles: "C(CC(C(=O)O)N)CN=C(N)N"
      - annotation: !!omap
          - bigg.metabolite: "arg__L"
          - chebi: "CHEBI:32682"
          - kegg.compound: "C00062"
          - metanetx.chemical: "MNXM70"
          - sbo: "SBO:0000247"
      - deltaG: 72.09
    - !!omap
      - id: "s_0967"
      - name: "L-arginine"
      - compartment: "m"
      - formula: "C6H15N4O2"
      - charge: 1
      - smiles: "C(CC(C(=O)O)N)CN=C(N)N"
      - annotation: !!omap
          - bigg.metabolite: "arg__L"
          - chebi: "CHEBI:32682"
          - kegg.compound: "C00062"
          - metanetx.chemical: "MNXM70"
          - sbo: "SBO:0000247"
      - deltaG: 72.09
    - !!omap
      - id: "s_0968"
      - name: "L-arginine"
      - compartment: "v"
      - formula: "C6H15N4O2"
      - charge: 1
      - smiles: "C(CC(C(=O)O)N)CN=C(N)N"
      - annotation: !!omap
          - bigg.metabolite: "arg__L"
          - chebi: "CHEBI:32682"
          - kegg.compound: "C00062"
          - metanetx.chemical: "MNXM70"
          - sbo: "SBO:0000247"
      - deltaG: 72.09
    - !!omap
      - id: "s_0969"
      - name: "L-asparagine"
      - compartment: "c"
      - formula: "C4H8N2O3"
      - charge: 0
      - smiles: "C(C(C(=O)O)N)C(=O)N"
      - annotation: !!omap
          - bigg.metabolite: "asn__L"
          - chebi: "CHEBI:17196"
          - kegg.compound: "C00152"
          - metanetx.chemical: "MNXM147"
          - sbo: "SBO:0000247"
      - deltaG: -45.42
    - !!omap
      - id: "s_0970"
      - name: "L-asparagine"
      - compartment: "e"
      - formula: "C4H8N2O3"
      - charge: 0
      - smiles: "C(C(C(=O)O)N)C(=O)N"
      - annotation: !!omap
          - bigg.metabolite: "asn__L"
          - chebi: "CHEBI:17196"
          - kegg.compound: "C00152"
          - metanetx.chemical: "MNXM147"
          - sbo: "SBO:0000247"
      - deltaG: -45.42
    - !!omap
      - id: "s_0971"
      - name: "L-asparagine"
      - compartment: "m"
      - formula: "C4H8N2O3"
      - charge: 0
      - smiles: "C(C(C(=O)O)N)C(=O)N"
      - annotation: !!omap
          - bigg.metabolite: "asn__L"
          - chebi: "CHEBI:17196"
          - kegg.compound: "C00152"
          - metanetx.chemical: "MNXM147"
          - sbo: "SBO:0000247"
      - deltaG: -45.42
    - !!omap
      - id: "s_0972"
      - name: "L-asparagine"
      - compartment: "v"
      - formula: "C4H8N2O3"
      - charge: 0
      - smiles: "C(C(C(=O)O)N)C(=O)N"
      - annotation: !!omap
          - bigg.metabolite: "asn__L"
          - chebi: "CHEBI:17196"
          - kegg.compound: "C00152"
          - metanetx.chemical: "MNXM147"
          - sbo: "SBO:0000247"
      - deltaG: -45.42
    - !!omap
      - id: "s_0973"
      - name: "L-aspartate"
      - compartment: "c"
      - formula: "C4H6NO4"
      - charge: -1
      - smiles: "C(C(C(=O)O)N)C(=O)O"
      - annotation: !!omap
          - bigg.metabolite: "asp__L"
          - chebi: "CHEBI:29991"
          - kegg.compound: "C00049"
          - metanetx.chemical: "MNXM42"
          - sbo: "SBO:0000247"
      - deltaG: -110.88
    - !!omap
      - id: "s_0974"
      - name: "L-aspartate"
      - compartment: "e"
      - formula: "C4H6NO4"
      - charge: -1
      - smiles: "C(C(C(=O)O)N)C(=O)O"
      - annotation: !!omap
          - bigg.metabolite: "asp__L"
          - chebi: "CHEBI:29991"
          - kegg.compound: "C00049"
          - metanetx.chemical: "MNXM42"
          - sbo: "SBO:0000247"
      - deltaG: -110.88
    - !!omap
      - id: "s_0975"
      - name: "L-aspartate"
      - compartment: "m"
      - formula: "C4H6NO4"
      - charge: -1
      - smiles: "C(C(C(=O)O)N)C(=O)O"
      - annotation: !!omap
          - bigg.metabolite: "asp__L"
          - chebi: "CHEBI:29991"
          - kegg.compound: "C00049"
          - metanetx.chemical: "MNXM42"
          - sbo: "SBO:0000247"
      - deltaG: -110.88
    - !!omap
      - id: "s_0976"
      - name: "L-aspartate"
      - compartment: "p"
      - formula: "C4H6NO4"
      - charge: -1
      - smiles: "C(C(C(=O)O)N)C(=O)O"
      - annotation: !!omap
          - bigg.metabolite: "asp__L"
          - chebi: "CHEBI:29991"
          - kegg.compound: "C00049"
          - metanetx.chemical: "MNXM42"
          - sbo: "SBO:0000247"
      - deltaG: -110.88
    - !!omap
      - id: "s_0977"
      - name: "L-aspartate"
      - compartment: "v"
      - formula: "C4H6NO4"
      - charge: -1
      - smiles: "C(C(C(=O)O)N)C(=O)O"
      - annotation: !!omap
          - bigg.metabolite: "asp__L"
          - chebi: "CHEBI:29991"
          - kegg.compound: "C00049"
          - metanetx.chemical: "MNXM42"
          - sbo: "SBO:0000247"
      - deltaG: -110.88
    - !!omap
      - id: "s_0978"
      - name: "L-aspartate 4-semialdehyde"
      - compartment: "c"
      - formula: "C4H7NO3"
      - charge: 0
      - smiles: "C(C=O)C(C(=O)[O-])[NH3+]"
      - annotation: !!omap
          - bigg.metabolite: "aspsa"
          - chebi: "CHEBI:18051"
          - kegg.compound: "C00441"
          - metanetx.chemical: "MNXM361"
          - sbo: "SBO:0000247"
      - deltaG: -44.89
    - !!omap
      - id: "s_0979"
      - name: "L-citrulline"
      - compartment: "c"
      - formula: "C6H13N3O3"
      - charge: 0
      - smiles: "C(CC(C(=O)O)N)CNC(=O)N"
      - annotation: !!omap
          - bigg.metabolite: "citr__L"
          - chebi: "CHEBI:16349"
          - kegg.compound: "C00327"
          - metanetx.chemical: "MNXM211"
          - sbo: "SBO:0000247"
      - deltaG: 2.71
    - !!omap
      - id: "s_0980"
      - name: "L-cystathionine"
      - compartment: "c"
      - formula: "C7H14N2O4S"
      - charge: 0
      - smiles: "C(CSCC(C(=O)O)N)C(C(=O)O)N"
      - annotation: !!omap
          - bigg.metabolite: "cyst__L"
          - chebi: "CHEBI:17482"
          - kegg.compound: "C02291"
          - metanetx.chemical: "MNXM319"
          - sbo: "SBO:0000247"
      - deltaG: -18.59
    - !!omap
      - id: "s_0981"
      - name: "L-cysteine"
      - compartment: "c"
      - formula: "C3H7NO2S"
      - charge: 0
      - smiles: "C(C(C(=O)O)N)S"
      - annotation: !!omap
          - bigg.metabolite: "cys__L"
          - chebi: "CHEBI:17561"
          - kegg.compound: "C00097"
          - metanetx.chemical: "MNXM55"
          - sbo: "SBO:0000247"
      - deltaG: -15.24
    - !!omap
      - id: "s_0982"
      - name: "L-cysteine"
      - compartment: "e"
      - formula: "C3H7NO2S"
      - charge: 0
      - smiles: "C(C(C(=O)O)N)S"
      - annotation: !!omap
          - bigg.metabolite: "cys__L"
          - chebi: "CHEBI:17561"
          - kegg.compound: "C00097"
          - metanetx.chemical: "MNXM55"
          - sbo: "SBO:0000247"
      - deltaG: -15.24
    - !!omap
      - id: "s_0983"
      - name: "L-cysteinylglycine"
      - compartment: "c"
      - formula: "C5H10N2O3S"
      - charge: 0
      - smiles: "C(C(C(=O)NCC(=O)O)N)S"
      - annotation: !!omap
          - bigg.metabolite: "cgly"
          - chebi: "CHEBI:4047"
          - kegg.compound: "C01419"
          - metanetx.chemical: "MNXM683"
          - sbo: "SBO:0000247"
      - deltaG: -21.63
    - !!omap
      - id: "s_0984"
      - name: "L-cystine"
      - compartment: "c"
      - formula: "C6H12N2O4S2"
      - charge: 0
      - smiles: "C(C(C(=O)O)N)SSCC(C(=O)O)N"
      - annotation: !!omap
          - bigg.metabolite: "cysi__L"
          - chebi: "CHEBI:16283"
          - kegg.compound: "C00491"
          - metanetx.chemical: "MNXM927"
          - sbo: "SBO:0000247"
      - deltaG: -42.77
    - !!omap
      - id: "s_0985"
      - name: "L-cystine"
      - compartment: "v"
      - formula: "C6H12N2O4S2"
      - charge: 0
      - smiles: "C(C(C(=O)O)N)SSCC(C(=O)O)N"
      - annotation: !!omap
          - bigg.metabolite: "cysi__L"
          - chebi: "CHEBI:16283"
          - kegg.compound: "C00491"
          - metanetx.chemical: "MNXM927"
          - sbo: "SBO:0000247"
      - deltaG: -42.77
    - !!omap
      - id: "s_0986"
      - name: "L-gamma-glutamyl phosphate"
      - compartment: "c"
      - formula: "C5H8NO7P"
      - charge: -2
      - smiles: "C(CC(=O)OP(=O)(O)O)C(C(=O)O)N"
      - annotation: !!omap
          - bigg.metabolite: "glu5p"
          - chebi: "CHEBI:58274"
          - kegg.compound: "C03287"
          - metanetx.chemical: "MNXM1280"
          - sbo: "SBO:0000247"
      - deltaG: -303.94
    - !!omap
      - id: "s_0987"
      - name: "L-gamma-glutamyl-L-alanine"
      - compartment: "c"
      - formula: "C8H13N2O5"
      - charge: -1
      - smiles: "CC(C(=O)O)NC(=O)CCC(C(=O)O)N"
      - annotation: !!omap
          - bigg.metabolite: "gluala"
          - chebi: "CHEBI:50619"
          - kegg.compound: "C03738"
          - metanetx.chemical: "MNXM59390"
          - sbo: "SBO:0000247"
      - deltaG: -73.81
    - !!omap
      - id: "s_0988"
      - name: "L-gamma-glutamyl-L-cysteine"
      - compartment: "c"
      - formula: "C8H13N2O5S"
      - charge: -1
      - smiles: "C(CC(=O)NC(CS)C(=O)O)C(C(=O)O)N"
      - annotation: !!omap
          - bigg.metabolite: "glucys"
          - chebi: "CHEBI:58173"
          - kegg.compound: "C00669"
          - metanetx.chemical: "MNXM412"
          - sbo: "SBO:0000247"
      - deltaG: -69.15
    - !!omap
      - id: "s_0989"
      - name: "L-glucitol"
      - compartment: "c"
      - formula: "C6H14O6"
      - charge: 0
      - smiles: "C(C(C(C(C(CO)O)O)O)O)O"
      - annotation: !!omap
          - bigg.metabolite: "sbt__L"
          - chebi: "CHEBI:28789"
          - kegg.compound: "C01722"
          - metanetx.chemical: "MNXM4638"
          - sbo: "SBO:0000247"
      - deltaG: -89.12
    - !!omap
      - id: "s_0990"
      - name: "L-glucitol"
      - compartment: "e"
      - formula: "C6H14O6"
      - charge: 0
      - smiles: "C(C(C(C(C(CO)O)O)O)O)O"
      - annotation: !!omap
          - bigg.metabolite: "sbt__L"
          - chebi: "CHEBI:28789"
          - kegg.compound: "C01722"
          - metanetx.chemical: "MNXM4638"
          - sbo: "SBO:0000247"
      - deltaG: -89.12
    - !!omap
      - id: "s_0991"
      - name: "L-glutamate"
      - compartment: "c"
      - formula: "C5H8NO4"
      - charge: -1
      - smiles: "C(CC(=O)O)C(C(=O)O)N"
      - annotation: !!omap
          - bigg.metabolite: "glu__L"
          - chebi: "CHEBI:29985"
          - kegg.compound: "C00025"
          - metanetx.chemical: "MNXM89557"
          - sbo: "SBO:0000247"
      - deltaG: -90.92
    - !!omap
      - id: "s_0992"
      - name: "L-glutamate"
      - compartment: "e"
      - formula: "C5H8NO4"
      - charge: -1
      - smiles: "C(CC(=O)O)C(C(=O)O)N"
      - annotation: !!omap
          - bigg.metabolite: "glu__L"
          - chebi: "CHEBI:29985"
          - kegg.compound: "C00025"
          - metanetx.chemical: "MNXM89557"
          - sbo: "SBO:0000247"
      - deltaG: -90.92
    - !!omap
      - id: "s_0993"
      - name: "L-glutamate"
      - compartment: "m"
      - formula: "C5H8NO4"
      - charge: -1
      - smiles: "C(CC(=O)O)C(C(=O)O)N"
      - annotation: !!omap
          - bigg.metabolite: "glu__L"
          - chebi: "CHEBI:29985"
          - kegg.compound: "C00025"
          - metanetx.chemical: "MNXM89557"
          - sbo: "SBO:0000247"
      - deltaG: -90.92
    - !!omap
      - id: "s_0994"
      - name: "L-glutamate"
      - compartment: "n"
      - formula: "C5H8NO4"
      - charge: -1
      - smiles: "C(CC(=O)O)C(C(=O)O)N"
      - annotation: !!omap
          - bigg.metabolite: "glu__L"
          - chebi: "CHEBI:29985"
          - kegg.compound: "C00025"
          - metanetx.chemical: "MNXM89557"
          - sbo: "SBO:0000247"
      - deltaG: -90.92
    - !!omap
      - id: "s_0995"
      - name: "L-glutamate"
      - compartment: "p"
      - formula: "C5H8NO4"
      - charge: -1
      - smiles: "C(CC(=O)O)C(C(=O)O)N"
      - annotation: !!omap
          - bigg.metabolite: "glu__L"
          - chebi: "CHEBI:29985"
          - kegg.compound: "C00025"
          - metanetx.chemical: "MNXM89557"
          - sbo: "SBO:0000247"
      - deltaG: -90.92
    - !!omap
      - id: "s_0996"
      - name: "L-glutamate"
      - compartment: "v"
      - formula: "C5H8NO4"
      - charge: -1
      - smiles: "C(CC(=O)O)C(C(=O)O)N"
      - annotation: !!omap
          - bigg.metabolite: "glu__L"
          - chebi: "CHEBI:29985"
          - kegg.compound: "C00025"
          - metanetx.chemical: "MNXM89557"
          - sbo: "SBO:0000247"
      - deltaG: -90.92
    - !!omap
      - id: "s_0997"
      - name: "L-glutamic 5-semialdehyde"
      - compartment: "c"
      - formula: "C5H9NO3"
      - charge: 0
      - smiles: "C(CC(C(=O)O)N)C=O"
      - annotation: !!omap
          - bigg.metabolite: "glu5sa"
          - chebi: "CHEBI:17232"
          - kegg.compound: "C01165"
          - metanetx.chemical: "MNXM245"
          - sbo: "SBO:0000247"
      - deltaG: -23.8
    - !!omap
      - id: "s_0999"
      - name: "L-glutamine"
      - compartment: "c"
      - formula: "C5H10N2O3"
      - charge: 0
      - smiles: "C(CC(=O)N)C(C(=O)O)N"
      - annotation: !!omap
          - bigg.metabolite: "gln__L"
          - chebi: "CHEBI:18050"
          - kegg.compound: "C00064"
          - metanetx.chemical: "MNXM37"
          - sbo: "SBO:0000247"
      - deltaG: -24.32
    - !!omap
      - id: "s_1000"
      - name: "L-glutamine"
      - compartment: "e"
      - formula: "C5H10N2O3"
      - charge: 0
      - smiles: "C(CC(=O)N)C(C(=O)O)N"
      - annotation: !!omap
          - bigg.metabolite: "gln__L"
          - chebi: "CHEBI:18050"
          - kegg.compound: "C00064"
          - metanetx.chemical: "MNXM37"
          - sbo: "SBO:0000247"
      - deltaG: -24.32
    - !!omap
      - id: "s_1001"
      - name: "L-glutamine"
      - compartment: "n"
      - formula: "C5H10N2O3"
      - charge: 0
      - smiles: "C(CC(=O)N)C(C(=O)O)N"
      - annotation: !!omap
          - bigg.metabolite: "gln__L"
          - chebi: "CHEBI:18050"
          - kegg.compound: "C00064"
          - metanetx.chemical: "MNXM37"
          - sbo: "SBO:0000247"
      - deltaG: -24.32
    - !!omap
      - id: "s_1002"
      - name: "L-glutamine"
      - compartment: "v"
      - formula: "C5H10N2O3"
      - charge: 0
      - smiles: "C(CC(=O)N)C(C(=O)O)N"
      - annotation: !!omap
          - bigg.metabolite: "gln__L"
          - chebi: "CHEBI:18050"
          - kegg.compound: "C00064"
          - metanetx.chemical: "MNXM37"
          - sbo: "SBO:0000247"
      - deltaG: -24.32
    - !!omap
      - id: "s_1003"
      - name: "L-glycine"
      - compartment: "c"
      - formula: "C2H5NO2"
      - charge: 0
      - smiles: "C(C(=O)O)N"
      - annotation: !!omap
          - bigg.metabolite: "gly"
          - chebi: "CHEBI:15428"
          - kegg.compound: "C00037"
          - metanetx.chemical: "MNXM29"
          - sbo: "SBO:0000247"
      - deltaG: -39.05
    - !!omap
      - id: "s_1004"
      - name: "L-glycine"
      - compartment: "e"
      - formula: "C2H5NO2"
      - charge: 0
      - smiles: "C(C(=O)O)N"
      - annotation: !!omap
          - bigg.metabolite: "gly"
          - chebi: "CHEBI:15428"
          - kegg.compound: "C00037"
          - metanetx.chemical: "MNXM29"
          - sbo: "SBO:0000247"
      - deltaG: -39.05
    - !!omap
      - id: "s_1005"
      - name: "L-glycine"
      - compartment: "m"
      - formula: "C2H5NO2"
      - charge: 0
      - smiles: "C(C(=O)O)N"
      - annotation: !!omap
          - bigg.metabolite: "gly"
          - chebi: "CHEBI:15428"
          - kegg.compound: "C00037"
          - metanetx.chemical: "MNXM29"
          - sbo: "SBO:0000247"
      - deltaG: -39.05
    - !!omap
      - id: "s_1006"
      - name: "L-histidine"
      - compartment: "c"
      - formula: "C6H9N3O2"
      - charge: 0
      - smiles: "C1=C(NC=N1)CC(C(=O)O)N"
      - annotation: !!omap
          - bigg.metabolite: "his__L"
          - chebi: "CHEBI:15971"
          - kegg.compound: "C00135"
          - metanetx.chemical: "MNXM134"
          - sbo: "SBO:0000247"
      - deltaG: 42
    - !!omap
      - id: "s_1007"
      - name: "L-histidine"
      - compartment: "e"
      - formula: "C6H9N3O2"
      - charge: 0
      - smiles: "C1=C(NC=N1)CC(C(=O)O)N"
      - annotation: !!omap
          - bigg.metabolite: "his__L"
          - chebi: "CHEBI:15971"
          - kegg.compound: "C00135"
          - metanetx.chemical: "MNXM134"
          - sbo: "SBO:0000247"
      - deltaG: 42
    - !!omap
      - id: "s_1008"
      - name: "L-histidine"
      - compartment: "m"
      - formula: "C6H9N3O2"
      - charge: 0
      - smiles: "C1=C(NC=N1)CC(C(=O)O)N"
      - annotation: !!omap
          - bigg.metabolite: "his__L"
          - chebi: "CHEBI:15971"
          - kegg.compound: "C00135"
          - metanetx.chemical: "MNXM134"
          - sbo: "SBO:0000247"
      - deltaG: 42
    - !!omap
      - id: "s_1009"
      - name: "L-histidine"
      - compartment: "v"
      - formula: "C6H9N3O2"
      - charge: 0
      - smiles: "C1=C(NC=N1)CC(C(=O)O)N"
      - annotation: !!omap
          - bigg.metabolite: "his__L"
          - chebi: "CHEBI:15971"
          - kegg.compound: "C00135"
          - metanetx.chemical: "MNXM134"
          - sbo: "SBO:0000247"
      - deltaG: 42
    - !!omap
      - id: "s_1010"
      - name: "L-histidinol"
      - compartment: "c"
      - formula: "C6H12N3O"
      - charge: 1
      - smiles: "C1=C(NC=N1)CC(CO)N"
      - annotation: !!omap
          - bigg.metabolite: "histd"
          - chebi: "CHEBI:57699"
          - kegg.compound: "C00860"
          - metanetx.chemical: "MNXM1281"
          - sbo: "SBO:0000247"
      - deltaG: 123.6
    - !!omap
      - id: "s_1011"
      - name: "L-histidinol phosphate"
      - compartment: "c"
      - formula: "C6H11N3O4P"
      - charge: -1
      - smiles: "C1=C(NC=N1)CC(COP(=O)(O)O)N"
      - annotation: !!omap
          - bigg.metabolite: "hisp"
          - chebi: "CHEBI:57980"
          - kegg.compound: "C01100"
          - metanetx.chemical: "MNXM1482"
          - sbo: "SBO:0000247"
      - deltaG: -109.22
    - !!omap
      - id: "s_1012"
      - name: "L-homocysteine"
      - compartment: "c"
      - formula: "C4H9NO2S"
      - charge: 0
      - smiles: "C(CS)C(C(=O)O)N"
      - annotation: !!omap
          - bigg.metabolite: "hcys__L"
          - chebi: "CHEBI:17588"
          - kegg.compound: "C00155"
          - metanetx.chemical: "MNXM123"
          - sbo: "SBO:0000247"
      - deltaG: 5.86
    - !!omap
      - id: "s_1014"
      - name: "L-homoserine"
      - compartment: "c"
      - formula: "C4H9NO3"
      - charge: 0
      - smiles: "C(CO)C(C(=O)O)N"
      - annotation: !!omap
          - bigg.metabolite: "hom__L"
          - chebi: "CHEBI:15699"
          - kegg.compound: "C00263"
          - metanetx.chemical: "MNXM353"
          - sbo: "SBO:0000247"
      - deltaG: -34.9
    - !!omap
      - id: "s_1015"
      - name: "L-homoserine"
      - compartment: "e"
      - formula: "C4H9NO3"
      - charge: 0
      - smiles: "C(CO)C(C(=O)O)N"
      - annotation: !!omap
          - bigg.metabolite: "hom__L"
          - chebi: "CHEBI:15699"
          - kegg.compound: "C00263"
          - metanetx.chemical: "MNXM353"
          - sbo: "SBO:0000247"
      - deltaG: -34.9
    - !!omap
      - id: "s_1016"
      - name: "L-isoleucine"
      - compartment: "c"
      - formula: "C6H13NO2"
      - charge: 0
      - smiles: "CCC(C)C(C(=O)O)N"
      - annotation: !!omap
          - bigg.metabolite: "ile__L"
          - chebi: "CHEBI:17191"
          - kegg.compound: "C00407"
          - metanetx.chemical: "MNXM231"
          - sbo: "SBO:0000247"
      - deltaG: 41.71
    - !!omap
      - id: "s_1017"
      - name: "L-isoleucine"
      - compartment: "e"
      - formula: "C6H13NO2"
      - charge: 0
      - smiles: "CCC(C)C(C(=O)O)N"
      - annotation: !!omap
          - bigg.metabolite: "ile__L"
          - chebi: "CHEBI:17191"
          - kegg.compound: "C00407"
          - metanetx.chemical: "MNXM231"
          - sbo: "SBO:0000247"
      - deltaG: 41.71
    - !!omap
      - id: "s_1018"
      - name: "L-isoleucine"
      - compartment: "m"
      - formula: "C6H13NO2"
      - charge: 0
      - smiles: "CCC(C)C(C(=O)O)N"
      - annotation: !!omap
          - bigg.metabolite: "ile__L"
          - chebi: "CHEBI:17191"
          - kegg.compound: "C00407"
          - metanetx.chemical: "MNXM231"
          - sbo: "SBO:0000247"
      - deltaG: 41.71
    - !!omap
      - id: "s_1019"
      - name: "L-isoleucine"
      - compartment: "v"
      - formula: "C6H13NO2"
      - charge: 0
      - smiles: "CCC(C)C(C(=O)O)N"
      - annotation: !!omap
          - bigg.metabolite: "ile__L"
          - chebi: "CHEBI:17191"
          - kegg.compound: "C00407"
          - metanetx.chemical: "MNXM231"
          - sbo: "SBO:0000247"
      - deltaG: 41.71
    - !!omap
      - id: "s_1020"
      - name: "L-kynurenine"
      - compartment: "c"
      - formula: "C10H12N2O3"
      - charge: 0
      - smiles: "C1=CC=C(C(=C1)C(=O)CC(C(=O)O)N)N"
      - annotation: !!omap
          - bigg.metabolite: "Lkynr"
          - chebi: "CHEBI:16946"
          - kegg.compound: "C00328"
          - metanetx.chemical: "MNXM260"
          - sbo: "SBO:0000247"
      - deltaG: 39.9
    - !!omap
      - id: "s_1021"
      - name: "L-leucine"
      - compartment: "c"
      - formula: "C6H13NO2"
      - charge: 0
      - smiles: "CC(C)CC(C(=O)O)N"
      - annotation: !!omap
          - bigg.metabolite: "leu__L"
          - chebi: "CHEBI:15603"
          - kegg.compound: "C00123"
          - metanetx.chemical: "MNXM140"
          - sbo: "SBO:0000247"
      - deltaG: 36.44
    - !!omap
      - id: "s_1022"
      - name: "L-leucine"
      - compartment: "e"
      - formula: "C6H13NO2"
      - charge: 0
      - smiles: "CC(C)CC(C(=O)O)N"
      - annotation: !!omap
          - bigg.metabolite: "leu__L"
          - chebi: "CHEBI:15603"
          - kegg.compound: "C00123"
          - metanetx.chemical: "MNXM140"
          - sbo: "SBO:0000247"
      - deltaG: 36.44
    - !!omap
      - id: "s_1023"
      - name: "L-leucine"
      - compartment: "m"
      - formula: "C6H13NO2"
      - charge: 0
      - smiles: "CC(C)CC(C(=O)O)N"
      - annotation: !!omap
          - bigg.metabolite: "leu__L"
          - chebi: "CHEBI:15603"
          - kegg.compound: "C00123"
          - metanetx.chemical: "MNXM140"
          - sbo: "SBO:0000247"
      - deltaG: 36.44
    - !!omap
      - id: "s_1024"
      - name: "L-leucine"
      - compartment: "v"
      - formula: "C6H13NO2"
      - charge: 0
      - smiles: "CC(C)CC(C(=O)O)N"
      - annotation: !!omap
          - bigg.metabolite: "leu__L"
          - chebi: "CHEBI:15603"
          - kegg.compound: "C00123"
          - metanetx.chemical: "MNXM140"
          - sbo: "SBO:0000247"
      - deltaG: 36.44
    - !!omap
      - id: "s_1025"
      - name: "L-lysine"
      - compartment: "c"
      - formula: "C6H15N2O2"
      - charge: 1
      - smiles: "C(CCN)CC(C(=O)O)N"
      - annotation: !!omap
          - bigg.metabolite: "lys__L"
          - chebi: "CHEBI:32551"
          - kegg.compound: "C00047"
          - metanetx.chemical: "MNXM78"
          - sbo: "SBO:0000247"
      - deltaG: 61.82
    - !!omap
      - id: "s_1026"
      - name: "L-lysine"
      - compartment: "e"
      - formula: "C6H15N2O2"
      - charge: 1
      - smiles: "C(CCN)CC(C(=O)O)N"
      - annotation: !!omap
          - bigg.metabolite: "lys__L"
          - chebi: "CHEBI:32551"
          - kegg.compound: "C00047"
          - metanetx.chemical: "MNXM78"
          - sbo: "SBO:0000247"
      - deltaG: 61.82
    - !!omap
      - id: "s_1027"
      - name: "L-lysine"
      - compartment: "m"
      - formula: "C6H15N2O2"
      - charge: 1
      - smiles: "C(CCN)CC(C(=O)O)N"
      - annotation: !!omap
          - bigg.metabolite: "lys__L"
          - chebi: "CHEBI:32551"
          - kegg.compound: "C00047"
          - metanetx.chemical: "MNXM78"
          - sbo: "SBO:0000247"
      - deltaG: 61.82
    - !!omap
      - id: "s_1028"
      - name: "L-lysine"
      - compartment: "v"
      - formula: "C6H15N2O2"
      - charge: 1
      - smiles: "C(CCN)CC(C(=O)O)N"
      - annotation: !!omap
          - bigg.metabolite: "lys__L"
          - chebi: "CHEBI:32551"
          - kegg.compound: "C00047"
          - metanetx.chemical: "MNXM78"
          - sbo: "SBO:0000247"
      - deltaG: 61.82
    - !!omap
      - id: "s_1029"
      - name: "L-methionine"
      - compartment: "c"
      - formula: "C5H11NO2S"
      - charge: 0
      - smiles: "CSCCC(C(=O)O)N"
      - annotation: !!omap
          - bigg.metabolite: "met__L"
          - chebi: "CHEBI:16643"
          - kegg.compound: "C00073"
          - metanetx.chemical: "MNXM61"
          - sbo: "SBO:0000247"
      - deltaG: 31.2
    - !!omap
      - id: "s_1030"
      - name: "L-methionine"
      - compartment: "e"
      - formula: "C5H11NO2S"
      - charge: 0
      - smiles: "CSCCC(C(=O)O)N"
      - annotation: !!omap
          - bigg.metabolite: "met__L"
          - chebi: "CHEBI:16643"
          - kegg.compound: "C00073"
          - metanetx.chemical: "MNXM61"
          - sbo: "SBO:0000247"
      - deltaG: 31.2
    - !!omap
      - id: "s_1031"
      - name: "L-methionine"
      - compartment: "m"
      - formula: "C5H11NO2S"
      - charge: 0
      - smiles: "CSCCC(C(=O)O)N"
      - annotation: !!omap
          - bigg.metabolite: "met__L"
          - chebi: "CHEBI:16643"
          - kegg.compound: "C00073"
          - metanetx.chemical: "MNXM61"
          - sbo: "SBO:0000247"
      - deltaG: 31.2
    - !!omap
      - id: "s_1032"
      - name: "L-phenylalanine"
      - compartment: "c"
      - formula: "C9H11NO2"
      - charge: 0
      - smiles: "C1=CC=C(C=C1)CC(C(=O)O)N"
      - annotation: !!omap
          - bigg.metabolite: "phe__L"
          - chebi: "CHEBI:17295"
          - kegg.compound: "C00079"
          - metanetx.chemical: "MNXM97"
          - sbo: "SBO:0000247"
      - deltaG: 56.06
    - !!omap
      - id: "s_1033"
      - name: "L-phenylalanine"
      - compartment: "e"
      - formula: "C9H11NO2"
      - charge: 0
      - smiles: "C1=CC=C(C=C1)CC(C(=O)O)N"
      - annotation: !!omap
          - bigg.metabolite: "phe__L"
          - chebi: "CHEBI:17295"
          - kegg.compound: "C00079"
          - metanetx.chemical: "MNXM97"
          - sbo: "SBO:0000247"
      - deltaG: 56.06
    - !!omap
      - id: "s_1034"
      - name: "L-phenylalanine"
      - compartment: "m"
      - formula: "C9H11NO2"
      - charge: 0
      - smiles: "C1=CC=C(C=C1)CC(C(=O)O)N"
      - annotation: !!omap
          - bigg.metabolite: "phe__L"
          - chebi: "CHEBI:17295"
          - kegg.compound: "C00079"
          - metanetx.chemical: "MNXM97"
          - sbo: "SBO:0000247"
      - deltaG: 56.06
    - !!omap
      - id: "s_1035"
      - name: "L-proline"
      - compartment: "c"
      - formula: "C5H9NO2"
      - charge: 0
      - smiles: "C1CC(NC1)C(=O)O"
      - annotation: !!omap
          - bigg.metabolite: "pro__L"
          - chebi: "CHEBI:17203"
          - kegg.compound: "C00148"
          - metanetx.chemical: "MNXM114"
          - sbo: "SBO:0000247"
      - deltaG: 10000000
    - !!omap
      - id: "s_1036"
      - name: "L-proline"
      - compartment: "e"
      - formula: "C5H9NO2"
      - charge: 0
      - smiles: "C1CC(NC1)C(=O)O"
      - annotation: !!omap
          - bigg.metabolite: "pro__L"
          - chebi: "CHEBI:17203"
          - kegg.compound: "C00148"
          - metanetx.chemical: "MNXM114"
          - sbo: "SBO:0000247"
      - deltaG: 10000000
    - !!omap
      - id: "s_1037"
      - name: "L-proline"
      - compartment: "m"
      - formula: "C5H9NO2"
      - charge: 0
      - smiles: "C1CC(NC1)C(=O)O"
      - annotation: !!omap
          - bigg.metabolite: "pro__L"
          - chebi: "CHEBI:17203"
          - kegg.compound: "C00148"
          - metanetx.chemical: "MNXM114"
          - sbo: "SBO:0000247"
      - deltaG: 10000000
    - !!omap
      - id: "s_1038"
      - name: "L-saccharopine"
      - compartment: "c"
      - formula: "C11H19N2O6"
      - charge: -1
      - smiles: "C(CCNC(CCC(=O)O)C(=O)O)CC(C(=O)O)N"
      - annotation: !!omap
          - bigg.metabolite: "saccrp__L"
          - chebi: "CHEBI:57951"
          - kegg.compound: "C00449"
          - metanetx.chemical: "MNXM384"
          - sbo: "SBO:0000247"
      - deltaG: -56.6
    - !!omap
      - id: "s_1039"
      - name: "L-serine"
      - compartment: "c"
      - formula: "C3H7NO3"
      - charge: 0
      - smiles: "C(C(C(=O)O)N)O"
      - annotation: !!omap
          - bigg.metabolite: "ser__L"
          - chebi: "CHEBI:17115"
          - kegg.compound: "C00065"
          - metanetx.chemical: "MNXM53"
          - sbo: "SBO:0000247"
      - deltaG: -55.99
    - !!omap
      - id: "s_1040"
      - name: "L-serine"
      - compartment: "er"
      - formula: "C3H7NO3"
      - charge: 0
      - smiles: "C(C(C(=O)O)N)O"
      - annotation: !!omap
          - bigg.metabolite: "ser__L"
          - chebi: "CHEBI:17115"
          - kegg.compound: "C00065"
          - metanetx.chemical: "MNXM53"
          - sbo: "SBO:0000247"
      - deltaG: -55.99
    - !!omap
      - id: "s_1041"
      - name: "L-serine"
      - compartment: "e"
      - formula: "C3H7NO3"
      - charge: 0
      - smiles: "C(C(C(=O)O)N)O"
      - annotation: !!omap
          - bigg.metabolite: "ser__L"
          - chebi: "CHEBI:17115"
          - kegg.compound: "C00065"
          - metanetx.chemical: "MNXM53"
          - sbo: "SBO:0000247"
      - deltaG: -55.99
    - !!omap
      - id: "s_1042"
      - name: "L-serine"
      - compartment: "m"
      - formula: "C3H7NO3"
      - charge: 0
      - smiles: "C(C(C(=O)O)N)O"
      - annotation: !!omap
          - bigg.metabolite: "ser__L"
          - chebi: "CHEBI:17115"
          - kegg.compound: "C00065"
          - metanetx.chemical: "MNXM53"
          - sbo: "SBO:0000247"
      - deltaG: -55.99
    - !!omap
      - id: "s_1043"
      - name: "L-sorbose"
      - compartment: "c"
      - formula: "C6H12O6"
      - charge: 0
      - smiles: "C1C(C(C(C(O1)(CO)O)O)O)O"
      - annotation: !!omap
          - bigg.metabolite: "srb__L"
          - chebi: "CHEBI:17266"
          - kegg.compound: "C00247"
          - metanetx.chemical: "MNXM588"
          - sbo: "SBO:0000247"
      - deltaG: -100.77
    - !!omap
      - id: "s_1044"
      - name: "L-sorbose"
      - compartment: "e"
      - formula: "C6H12O6"
      - charge: 0
      - smiles: "C1C(C(C(C(O1)(CO)O)O)O)O"
      - annotation: !!omap
          - bigg.metabolite: "srb__L"
          - chebi: "CHEBI:17266"
          - kegg.compound: "C00247"
          - metanetx.chemical: "MNXM588"
          - sbo: "SBO:0000247"
      - deltaG: -100.77
    - !!omap
      - id: "s_1045"
      - name: "L-threonine"
      - compartment: "c"
      - formula: "C4H9NO3"
      - charge: 0
      - smiles: "CC(C(C(=O)O)N)O"
      - annotation: !!omap
          - bigg.metabolite: "thr__L"
          - chebi: "CHEBI:16857"
          - kegg.compound: "C00188"
          - metanetx.chemical: "MNXM142"
          - sbo: "SBO:0000247"
      - deltaG: -36.71
    - !!omap
      - id: "s_1046"
      - name: "L-threonine"
      - compartment: "e"
      - formula: "C4H9NO3"
      - charge: 0
      - smiles: "CC(C(C(=O)O)N)O"
      - annotation: !!omap
          - bigg.metabolite: "thr__L"
          - chebi: "CHEBI:16857"
          - kegg.compound: "C00188"
          - metanetx.chemical: "MNXM142"
          - sbo: "SBO:0000247"
      - deltaG: -36.71
    - !!omap
      - id: "s_1047"
      - name: "L-threonine"
      - compartment: "m"
      - formula: "C4H9NO3"
      - charge: 0
      - smiles: "CC(C(C(=O)O)N)O"
      - annotation: !!omap
          - bigg.metabolite: "thr__L"
          - chebi: "CHEBI:16857"
          - kegg.compound: "C00188"
          - metanetx.chemical: "MNXM142"
          - sbo: "SBO:0000247"
      - deltaG: -36.71
    - !!omap
      - id: "s_1048"
      - name: "L-tryptophan"
      - compartment: "c"
      - formula: "C11H12N2O2"
      - charge: 0
      - smiles: "C1=CC=C2C(=C1)C(=CN2)CC(C(=O)O)N"
      - annotation: !!omap
          - bigg.metabolite: "trp__L"
          - chebi: "CHEBI:16828"
          - kegg.compound: "C00078"
          - metanetx.chemical: "MNXM94"
          - sbo: "SBO:0000247"
      - deltaG: 101.55
    - !!omap
      - id: "s_1049"
      - name: "L-tryptophan"
      - compartment: "e"
      - formula: "C11H12N2O2"
      - charge: 0
      - smiles: "C1=CC=C2C(=C1)C(=CN2)CC(C(=O)O)N"
      - annotation: !!omap
          - bigg.metabolite: "trp__L"
          - chebi: "CHEBI:16828"
          - kegg.compound: "C00078"
          - metanetx.chemical: "MNXM94"
          - sbo: "SBO:0000247"
      - deltaG: 101.55
    - !!omap
      - id: "s_1050"
      - name: "L-tryptophan"
      - compartment: "m"
      - formula: "C11H12N2O2"
      - charge: 0
      - smiles: "C1=CC=C2C(=C1)C(=CN2)CC(C(=O)O)N"
      - annotation: !!omap
          - bigg.metabolite: "trp__L"
          - chebi: "CHEBI:16828"
          - kegg.compound: "C00078"
          - metanetx.chemical: "MNXM94"
          - sbo: "SBO:0000247"
      - deltaG: 101.55
    - !!omap
      - id: "s_1051"
      - name: "L-tyrosine"
      - compartment: "c"
      - formula: "C9H11NO3"
      - charge: 0
      - smiles: "C1=CC(=CC=C1CC(C(=O)O)N)O"
      - annotation: !!omap
          - bigg.metabolite: "tyr__L"
          - chebi: "CHEBI:17895"
          - kegg.compound: "C00082"
          - metanetx.chemical: "MNXM76"
          - sbo: "SBO:0000247"
      - deltaG: 16.58
    - !!omap
      - id: "s_1052"
      - name: "L-tyrosine"
      - compartment: "e"
      - formula: "C9H11NO3"
      - charge: 0
      - smiles: "C1=CC(=CC=C1CC(C(=O)O)N)O"
      - annotation: !!omap
          - bigg.metabolite: "tyr__L"
          - chebi: "CHEBI:17895"
          - kegg.compound: "C00082"
          - metanetx.chemical: "MNXM76"
          - sbo: "SBO:0000247"
      - deltaG: 16.58
    - !!omap
      - id: "s_1053"
      - name: "L-tyrosine"
      - compartment: "m"
      - formula: "C9H11NO3"
      - charge: 0
      - smiles: "C1=CC(=CC=C1CC(C(=O)O)N)O"
      - annotation: !!omap
          - bigg.metabolite: "tyr__L"
          - chebi: "CHEBI:17895"
          - kegg.compound: "C00082"
          - metanetx.chemical: "MNXM76"
          - sbo: "SBO:0000247"
      - deltaG: 16.58
    - !!omap
      - id: "s_1054"
      - name: "L-tyrosine"
      - compartment: "p"
      - formula: "C9H11NO3"
      - charge: 0
      - smiles: "C1=CC(=CC=C1CC(C(=O)O)N)O"
      - annotation: !!omap
          - bigg.metabolite: "tyr__L"
          - chebi: "CHEBI:17895"
          - kegg.compound: "C00082"
          - metanetx.chemical: "MNXM76"
          - sbo: "SBO:0000247"
      - deltaG: 16.58
    - !!omap
      - id: "s_1055"
      - name: "L-tyrosine"
      - compartment: "v"
      - formula: "C9H11NO3"
      - charge: 0
      - smiles: "C1=CC(=CC=C1CC(C(=O)O)N)O"
      - annotation: !!omap
          - bigg.metabolite: "tyr__L"
          - chebi: "CHEBI:17895"
          - kegg.compound: "C00082"
          - metanetx.chemical: "MNXM76"
          - sbo: "SBO:0000247"
      - deltaG: 16.58
    - !!omap
      - id: "s_1056"
      - name: "L-valine"
      - compartment: "c"
      - formula: "C5H11NO2"
      - charge: 0
      - smiles: "CC(C)C(C(=O)O)N"
      - annotation: !!omap
          - bigg.metabolite: "val__L"
          - chebi: "CHEBI:16414"
          - kegg.compound: "C00183"
          - metanetx.chemical: "MNXM199"
          - sbo: "SBO:0000247"
      - deltaG: 20.62
    - !!omap
      - id: "s_1057"
      - name: "L-valine"
      - compartment: "e"
      - formula: "C5H11NO2"
      - charge: 0
      - smiles: "CC(C)C(C(=O)O)N"
      - annotation: !!omap
          - bigg.metabolite: "val__L"
          - chebi: "CHEBI:16414"
          - kegg.compound: "C00183"
          - metanetx.chemical: "MNXM199"
          - sbo: "SBO:0000247"
      - deltaG: 20.62
    - !!omap
      - id: "s_1058"
      - name: "L-valine"
      - compartment: "m"
      - formula: "C5H11NO2"
      - charge: 0
      - smiles: "CC(C)C(C(=O)O)N"
      - annotation: !!omap
          - bigg.metabolite: "val__L"
          - chebi: "CHEBI:16414"
          - kegg.compound: "C00183"
          - metanetx.chemical: "MNXM199"
          - sbo: "SBO:0000247"
      - deltaG: 20.62
    - !!omap
      - id: "s_1059"
      - name: "lanosterol"
      - compartment: "c"
      - formula: "C30H50O"
      - charge: 0
      - smiles: "CC(CCC=C(C)C)C1CCC2(C1(CCC3=C2CCC4C3(CCC(C4(C)C)O)C)C)C"
      - annotation: !!omap
          - bigg.metabolite: "lanost"
          - chebi: "CHEBI:16521"
          - kegg.compound: "C01724"
          - metanetx.chemical: "MNXM482"
          - sbo: "SBO:0000247"
      - deltaG: 519.58
    - !!omap
      - id: "s_1061"
      - name: "lanosterol"
      - compartment: "e"
      - formula: "C30H50O"
      - charge: 0
      - smiles: "CC(CCC=C(C)C)C1CCC2(C1(CCC3=C2CCC4C3(CCC(C4(C)C)O)C)C)C"
      - annotation: !!omap
          - bigg.metabolite: "lanost"
          - chebi: "CHEBI:16521"
          - kegg.compound: "C01724"
          - metanetx.chemical: "MNXM482"
          - sbo: "SBO:0000247"
      - deltaG: 519.58
    - !!omap
      - id: "s_1065"
      - name: "laurate"
      - compartment: "c"
      - formula: "C12H23O2"
      - charge: -1
      - smiles: "CCCCCCCCCCCC(=O)[O-]"
      - annotation: !!omap
          - bigg.metabolite: "ddca"
          - chebi: "CHEBI:18262"
          - kegg.compound: "C02679"
          - metanetx.chemical: "MNXM402"
          - sbo: "SBO:0000247"
      - deltaG: 153.2
    - !!omap
      - id: "s_1067"
      - name: "laurate"
      - compartment: "e"
      - formula: "C12H23O2"
      - charge: -1
      - smiles: "CCCCCCCCCCCC(=O)[O-]"
      - annotation: !!omap
          - bigg.metabolite: "ddca"
          - chebi: "CHEBI:18262"
          - kegg.compound: "C02679"
          - metanetx.chemical: "MNXM402"
          - sbo: "SBO:0000247"
      - deltaG: 153.2
    - !!omap
      - id: "s_1070"
      - name: "laurate"
      - compartment: "p"
      - formula: "C12H23O2"
      - charge: -1
      - smiles: "CCCCCCCCCCCC(=O)[O-]"
      - annotation: !!omap
          - bigg.metabolite: "ddca"
          - chebi: "CHEBI:18262"
          - kegg.compound: "C02679"
          - metanetx.chemical: "MNXM402"
          - sbo: "SBO:0000247"
      - deltaG: 153.2
    - !!omap
      - id: "s_1073"
      - name: "lauroyl-CoA"
      - compartment: "c"
      - formula: "C33H54N7O17P3S"
      - charge: -4
      - smiles: "CCCCCCCCCCCC(=O)SCCNC(=O)CCNC(=O)C(C(C)(C)COP(=O)(O)OP(=O)(O)OCC1C(C(C(O1)N2C=NC3=C(N=CN=C32)N)O)OP(=O)(O)O)O"
      - annotation: !!omap
          - bigg.metabolite: "ddcacoa"
          - chebi: "CHEBI:57375"
          - kegg.compound: "C01832"
          - metanetx.chemical: "MNXM363"
          - sbo: "SBO:0000247"
      - deltaG: -257.64
    - !!omap
      - id: "s_1076"
      - name: "lauroyl-CoA"
      - compartment: "p"
      - formula: "C33H54N7O17P3S"
      - charge: -4
      - smiles: "CCCCCCCCCCCC(=O)SCCNC(=O)CCNC(=O)C(C(C)(C)COP(=O)(O)OP(=O)(O)OCC1C(C(C(O1)N2C=NC3=C(N=CN=C32)N)O)OP(=O)(O)O)O"
      - annotation: !!omap
          - bigg.metabolite: "ddcacoa"
          - chebi: "CHEBI:57375"
          - kegg.compound: "C01832"
          - metanetx.chemical: "MNXM363"
          - sbo: "SBO:0000247"
      - deltaG: -257.64
    - !!omap
      - id: "s_1077"
      - name: "Leu-tRNA(Leu)"
      - compartment: "c"
      - formula: "C6H13NOR"
      - charge: 1
      - annotation: !!omap
          - bigg.metabolite: "leutrna"
          - chebi: "CHEBI:16624"
          - kegg.compound: "C02047"
          - metanetx.chemical: "MNXM697"
          - sbo: "SBO:0000247"
      - deltaG: 10000000
    - !!omap
      - id: "s_1078"
      - name: "Leu-tRNA(Leu)"
      - compartment: "m"
      - formula: "C6H13NOR"
      - charge: 1
      - annotation: !!omap
          - bigg.metabolite: "leutrna"
          - chebi: "CHEBI:16624"
          - kegg.compound: "C02047"
          - metanetx.chemical: "MNXM697"
          - sbo: "SBO:0000247"
      - deltaG: 10000000
    - !!omap
      - id: "s_1079"
      - name: "leukotriene A4"
      - compartment: "c"
      - formula: "C20H29O3"
      - charge: -1
      - smiles: "CCCCCC=CCC=CC=CC=CC1C(O1)CCCC(=O)O"
      - annotation: !!omap
          - bigg.metabolite: "leuktrA4"
          - chebi: "CHEBI:57463"
          - kegg.compound: "C00909"
          - metanetx.chemical: "MNXM462"
          - sbo: "SBO:0000247"
      - deltaG: 288.62
    - !!omap
      - id: "s_1080"
      - name: "leukotriene A4"
      - compartment: "n"
      - formula: "C20H29O3"
      - charge: -1
      - smiles: "CCCCCC=CCC=CC=CC=CC1C(O1)CCCC(=O)O"
      - annotation: !!omap
          - bigg.metabolite: "leuktrA4"
          - chebi: "CHEBI:57463"
          - kegg.compound: "C00909"
          - metanetx.chemical: "MNXM462"
          - sbo: "SBO:0000247"
      - deltaG: 288.62
    - !!omap
      - id: "s_1081"
      - name: "leukotriene B4"
      - compartment: "c"
      - formula: "C20H31O4"
      - charge: -1
      - smiles: "CCCCCC=CCC(C=CC=CC=CC(CCCC(=O)O)O)O"
      - annotation: !!omap
          - bigg.metabolite: "leuktrB4"
          - chebi: "CHEBI:57461"
          - kegg.compound: "C02165"
          - metanetx.chemical: "MNXM637"
          - sbo: "SBO:0000247"
      - deltaG: 247.78
    - !!omap
      - id: "s_1082"
      - name: "leukotriene B4"
      - compartment: "n"
      - formula: "C20H31O4"
      - charge: -1
      - smiles: "CCCCCC=CCC(C=CC=CC=CC(CCCC(=O)O)O)O"
      - annotation: !!omap
          - bigg.metabolite: "leuktrB4"
          - chebi: "CHEBI:57461"
          - kegg.compound: "C02165"
          - metanetx.chemical: "MNXM637"
          - sbo: "SBO:0000247"
      - deltaG: 247.78
    - !!omap
      - id: "s_1085"
      - name: "lignoceric acid"
      - compartment: "er"
      - formula: "C24H47O2"
      - charge: -1
      - smiles: "CCCCCCCCCCCCCCCCCCCCCCCC(=O)O"
      - annotation: !!omap
          - bigg.metabolite: "lgnc"
          - chebi: "CHEBI:31014"
          - kegg.compound: "C08320"
          - metanetx.chemical: "MNXM3297"
          - sbo: "SBO:0000247"
      - deltaG: 410.24
    - !!omap
      - id: "s_1096"
      - name: "lipid"
      - compartment: "c"
      - charge: 0
      - annotation: !!omap
          - chebi: "CHEBI:18059"
          - kegg.compound: "C01356"
          - metanetx.chemical: "MNXM12117"
          - sbo: "SBO:0000649"
      - deltaG: 10000000
    - !!omap
      - id: "s_1097"
      - name: "lipoamide"
      - compartment: "m"
      - formula: "C8H15NOS2"
      - charge: 0
      - smiles: "C1CSSC1CCCCC(=O)N"
      - annotation: !!omap
          - bigg.metabolite: "lpam"
          - chebi: "CHEBI:17460"
          - kegg.compound: "C00248"
          - metanetx.chemical: "MNXM1024"
          - sbo: "SBO:0000247"
      - deltaG: 134.74
    - !!omap
      - id: "s_1098"
      - name: "lipoylprotein"
      - compartment: "m"
      - formula: "C8H14NOS2R"
      - charge: 0
      - annotation: !!omap
          - bigg.metabolite: "lpro"
          - chebi: "CHEBI:15804"
          - kegg.compound: "C02051"
          - metanetx.chemical: "MNXM998"
          - sbo: "SBO:0000247"
      - deltaG: 10000000
    - !!omap
      - id: "s_1099"
      - name: "Lys-tRNA(Lys)"
      - compartment: "c"
      - formula: "C6H15N2OR"
      - charge: 2
      - annotation: !!omap
          - bigg.metabolite: "lystrna"
          - chebi: "CHEBI:16047"
          - kegg.compound: "C01931"
          - metanetx.chemical: "MNXM89922"
          - sbo: "SBO:0000247"
      - deltaG: 10000000
    - !!omap
      - id: "s_1100"
      - name: "Lys-tRNA(Lys)"
      - compartment: "m"
      - formula: "C6H15N2OR"
      - charge: 2
      - annotation: !!omap
          - bigg.metabolite: "lystrna"
          - chebi: "CHEBI:16047"
          - kegg.compound: "C01931"
          - metanetx.chemical: "MNXM89922"
          - sbo: "SBO:0000247"
      - deltaG: 10000000
    - !!omap
      - id: "s_1101"
      - name: "malonyl-CoA"
      - compartment: "c"
      - formula: "C24H33N7O19P3S"
      - charge: -5
      - smiles: "CC(C)(COP(=O)(O)OP(=O)(O)OCC1C(C(C(O1)N2C=NC3=C(N=CN=C32)N)O)OP(=O)(O)O)C(C(=O)NCCC(=O)NCCSC(=O)CC(=O)O)O"
      - annotation: !!omap
          - bigg.metabolite: "malcoa"
          - chebi: "CHEBI:57384"
          - kegg.compound: "C00083"
          - metanetx.chemical: "MNXM40"
          - sbo: "SBO:0000247"
      - deltaG: -557.88
    - !!omap
      - id: "s_1104"
      - name: "malonyl-CoA"
      - compartment: "m"
      - formula: "C24H33N7O19P3S"
      - charge: -5
      - smiles: "CC(C)(COP(=O)(O)OP(=O)(O)OCC1C(C(C(O1)N2C=NC3=C(N=CN=C32)N)O)OP(=O)(O)O)C(C(=O)NCCC(=O)NCCSC(=O)CC(=O)O)O"
      - annotation: !!omap
          - bigg.metabolite: "malcoa"
          - chebi: "CHEBI:57384"
          - kegg.compound: "C00083"
          - metanetx.chemical: "MNXM40"
          - sbo: "SBO:0000247"
      - deltaG: -557.88
    - !!omap
      - id: "s_1105"
      - name: "maltose"
      - compartment: "c"
      - formula: "C12H22O11"
      - charge: 0
      - smiles: "C(C1C(C(C(C(O1)OC2C(OC(C(C2O)O)O)CO)O)O)O)O"
      - annotation: !!omap
          - bigg.metabolite: "malt"
          - chebi: "CHEBI:17306"
          - kegg.compound: "C00208"
          - metanetx.chemical: "MNXM165"
          - sbo: "SBO:0000247"
      - deltaG: -162.55
    - !!omap
      - id: "s_1106"
      - name: "maltose"
      - compartment: "e"
      - formula: "C12H22O11"
      - charge: 0
      - smiles: "C(C1C(C(C(C(O1)OC2C(OC(C(C2O)O)O)CO)O)O)O)O"
      - annotation: !!omap
          - bigg.metabolite: "malt"
          - chebi: "CHEBI:17306"
          - kegg.compound: "C00208"
          - metanetx.chemical: "MNXM165"
          - sbo: "SBO:0000247"
      - deltaG: -162.55
    - !!omap
      - id: "s_1107"
      - name: "mannan"
      - compartment: "c"
      - formula: "C6H10O5"
      - charge: 0
      - smiles: "C(C1C(C(C(C(O1)OC2C(OC(C(C2O)O)OC3C(OC(C(C3O)O)OC4C(OC(C(C4O)O)O)CO)CO)CO)O)O)O)O"
      - annotation: !!omap
          - bigg.metabolite: "mannan"
          - chebi: "CHEBI:28808"
          - kegg.compound: "C00464"
          - metanetx.chemical: "MNXM2020"
          - sbo: "SBO:0000247"
      - deltaG: 10000000
    - !!omap
      - id: "s_1108"
      - name: "mannan"
      - compartment: "er"
      - formula: "C6H10O5"
      - charge: 0
      - smiles: "C(C1C(C(C(C(O1)OC2C(OC(C(C2O)O)OC3C(OC(C(C3O)O)OC4C(OC(C(C4O)O)O)CO)CO)CO)O)O)O)O"
      - annotation: !!omap
          - bigg.metabolite: "mannan"
          - chebi: "CHEBI:28808"
          - kegg.compound: "C00464"
          - metanetx.chemical: "MNXM2020"
          - sbo: "SBO:0000247"
      - deltaG: 10000000
    - !!omap
      - id: "s_1109"
      - name: "mannose-(1D-myo-inositol 1-phosphate)2"
      - compartment: "er"
      - formula: "C18H34O22P2"
      - charge: 0
      - smiles: "C(C1C(C(C(C(O1)OC2C(C(C(C(C2OP(=O)(O)O)O)O)O)O)O)O)O)OP(=O)(O)OC3C(C(C(C(C3O)O)O)O)O"
      - annotation: !!omap
          - bigg.metabolite: "man2mi1p__D"
          - chebi: "CHEBI:60449"
          - metanetx.chemical: "MNXM61128"
          - sbo: "SBO:0000247"
      - deltaG: -657.63
    - !!omap
      - id: "s_1110"
      - name: "mannose-(1D-myo-inositol 1-phosphate)2"
      - compartment: "m"
      - formula: "C18H34O22P2"
      - charge: 0
      - smiles: "C(C1C(C(C(C(O1)OC2C(C(C(C(C2OP(=O)(O)O)O)O)O)O)O)O)O)OP(=O)(O)OC3C(C(C(C(C3O)O)O)O)O"
      - annotation: !!omap
          - bigg.metabolite: "man2mi1p__D"
          - chebi: "CHEBI:60449"
          - metanetx.chemical: "MNXM61128"
          - sbo: "SBO:0000247"
      - deltaG: -657.63
    - !!omap
      - id: "s_1111"
      - name: "mannose-1D-myo-inositol 1-phosphate"
      - compartment: "er"
      - formula: "C12H23O14P"
      - charge: 0
      - smiles: "C(C1C(C(C(C(O1)OC2C(C(C(C(C2OP(=O)(O)O)O)O)O)O)O)O)O)O"
      - annotation: !!omap
          - bigg.metabolite: "manmi1p__D"
          - chebi: "CHEBI:60448"
          - metanetx.chemical: "MNXM61129"
          - sbo: "SBO:0000247"
      - deltaG: -384.83
    - !!omap
      - id: "s_1112"
      - name: "mannose-1D-myo-inositol 1-phosphate"
      - compartment: "m"
      - formula: "C12H23O14P"
      - charge: 0
      - smiles: "C(C1C(C(C(C(O1)OC2C(C(C(C(C2OP(=O)(O)O)O)O)O)O)O)O)O)O"
      - annotation: !!omap
          - bigg.metabolite: "manmi1p__D"
          - chebi: "CHEBI:60448"
          - metanetx.chemical: "MNXM61129"
          - sbo: "SBO:0000247"
      - deltaG: -384.83
    - !!omap
      - id: "s_1114"
      - name: "mannosylinositol phosphorylceramide backbone"
      - compartment: "g"
      - formula: "C12H22NO15P"
      - annotation: !!omap
          - chebi: "CHEBI:25168"
          - metanetx.chemical: "MNXM12216"
          - sbo: "SBO:0000649"
      - deltaG: 10000000
    - !!omap
      - id: "s_1116"
      - name: "mannosylinositol phosphorylceramide A (C24)"
      - compartment: "er"
      - formula: "C54H106NO16P"
      - charge: 0
      - smiles: "CCCCCCCCCCCCCCCCCCCCCCCC(=O)NC(COP(=O)(O)OC1C(C(C(C(C1OC2C(C(C(C(O2)CO)O)O)O)O)O)O)O)C(CCCCCCCCCCCCCCC)O"
      - annotation: !!omap
          - chebi: "CHEBI:60416"
          - metanetx.chemical: "MNXM61061"
          - sbo: "SBO:0000247"
      - deltaG: 448.84
    - !!omap
      - id: "s_1117"
      - name: "mannosylinositol phosphorylceramide A (C24)"
      - compartment: "g"
      - formula: "C54H106NO16P"
      - charge: 0
      - smiles: "CCCCCCCCCCCCCCCCCCCCCCCC(=O)NC(COP(=O)(O)OC1C(C(C(C(C1OC2C(C(C(C(O2)CO)O)O)O)O)O)O)O)C(CCCCCCCCCCCCCCC)O"
      - annotation: !!omap
          - chebi: "CHEBI:60416"
          - metanetx.chemical: "MNXM61061"
          - sbo: "SBO:0000247"
      - deltaG: 448.84
    - !!omap
      - id: "s_1118"
      - name: "mannosylinositol phosphorylceramide A (C24)"
      - compartment: "m"
      - formula: "C54H106NO16P"
      - charge: 0
      - smiles: "CCCCCCCCCCCCCCCCCCCCCCCC(=O)NC(COP(=O)(O)OC1C(C(C(C(C1OC2C(C(C(C(O2)CO)O)O)O)O)O)O)O)C(CCCCCCCCCCCCCCC)O"
      - annotation: !!omap
          - chebi: "CHEBI:60416"
          - metanetx.chemical: "MNXM61061"
          - sbo: "SBO:0000247"
      - deltaG: 448.84
    - !!omap
      - id: "s_1119"
      - name: "mannosylinositol phosphorylceramide A (C26)"
      - compartment: "er"
      - formula: "C56H110NO16P"
      - charge: 0
      - smiles: "CCCCCCCCCCCCCCCCCCCCCCCCCC(=O)NC(COP(=O)(O)OC1C(C(C(C(C1OC2C(C(C(C(O2)CO)O)O)O)O)O)O)O)C(CCCCCCCCCCCCCCCCC)O"
      - annotation: !!omap
          - chebi: "CHEBI:62706"
          - metanetx.chemical: "MNXM106432"
          - sbo: "SBO:0000247"
      - deltaG: 491.35
    - !!omap
      - id: "s_1120"
      - name: "mannosylinositol phosphorylceramide A (C26)"
      - compartment: "g"
      - formula: "C56H110NO16P"
      - charge: 0
      - smiles: "CCCCCCCCCCCCCCCCCCCCCCCCCC(=O)NC(COP(=O)(O)OC1C(C(C(C(C1OC2C(C(C(C(O2)CO)O)O)O)O)O)O)O)C(CCCCCCCCCCCCCCCCC)O"
      - annotation: !!omap
          - chebi: "CHEBI:62706"
          - metanetx.chemical: "MNXM106432"
          - sbo: "SBO:0000247"
      - deltaG: 491.35
    - !!omap
      - id: "s_1121"
      - name: "mannosylinositol phosphorylceramide A (C26)"
      - compartment: "m"
      - formula: "C56H110NO16P"
      - charge: 0
      - smiles: "CCCCCCCCCCCCCCCCCCCCCCCCCC(=O)NC(COP(=O)(O)OC1C(C(C(C(C1OC2C(C(C(C(O2)CO)O)O)O)O)O)O)O)C(CCCCCCCCCCCCCCCCC)O"
      - annotation: !!omap
          - chebi: "CHEBI:62706"
          - metanetx.chemical: "MNXM106432"
          - sbo: "SBO:0000247"
      - deltaG: 491.35
    - !!omap
      - id: "s_1122"
      - name: "mannosylinositol phosphorylceramide B (C24)"
      - compartment: "er"
      - formula: "C54H106NO17P"
      - charge: 0
      - smiles: "CCCCCCCCCCCCCCCCCCCCCCCC(=O)NC(COP(=O)(O)OC1C(C(C(C(C1OC2C(C(C(C(O2)CO)O)O)O)O)O)O)O)C(C(CCCCCCCCCCCCCC)O)O"
      - annotation: !!omap
          - chebi: "CHEBI:60420"
          - metanetx.chemical: "MNXM61066"
          - sbo: "SBO:0000247"
      - deltaG: 410.8
    - !!omap
      - id: "s_1123"
      - name: "mannosylinositol phosphorylceramide B (C24)"
      - compartment: "g"
      - formula: "C54H106NO17P"
      - charge: 0
      - smiles: "CCCCCCCCCCCCCCCCCCCCCCCC(=O)NC(COP(=O)(O)OC1C(C(C(C(C1OC2C(C(C(C(O2)CO)O)O)O)O)O)O)O)C(C(CCCCCCCCCCCCCC)O)O"
      - annotation: !!omap
          - chebi: "CHEBI:60420"
          - metanetx.chemical: "MNXM61066"
          - sbo: "SBO:0000247"
      - deltaG: 410.8
    - !!omap
      - id: "s_1124"
      - name: "mannosylinositol phosphorylceramide B (C24)"
      - compartment: "m"
      - formula: "C54H106NO17P"
      - charge: 0
      - smiles: "CCCCCCCCCCCCCCCCCCCCCCCC(=O)NC(COP(=O)(O)OC1C(C(C(C(C1OC2C(C(C(C(O2)CO)O)O)O)O)O)O)O)C(C(CCCCCCCCCCCCCC)O)O"
      - annotation: !!omap
          - chebi: "CHEBI:60420"
          - metanetx.chemical: "MNXM61066"
          - sbo: "SBO:0000247"
      - deltaG: 410.8
    - !!omap
      - id: "s_1125"
      - name: "mannosylinositol phosphorylceramide B (C26)"
      - compartment: "er"
      - formula: "C56H110NO17P"
      - charge: 0
      - smiles: "CCCCCCCCCCCCCCCCCCCCCCCCCC(=O)NC(COP(=O)(O)OC1C(C(C(C(C1OC2C(C(C(C(O2)CO)O)O)O)O)O)O)O)C(C(CCCCCCCCCCCCCCCC)O)O"
      - annotation: !!omap
          - chebi: "CHEBI:62710"
          - metanetx.chemical: "MNXM61056"
          - sbo: "SBO:0000247"
      - deltaG: 453.31
    - !!omap
      - id: "s_1126"
      - name: "mannosylinositol phosphorylceramide B (C26)"
      - compartment: "g"
      - formula: "C56H110NO17P"
      - charge: 0
      - smiles: "CCCCCCCCCCCCCCCCCCCCCCCCCC(=O)NC(COP(=O)(O)OC1C(C(C(C(C1OC2C(C(C(C(O2)CO)O)O)O)O)O)O)O)C(C(CCCCCCCCCCCCCCCC)O)O"
      - annotation: !!omap
          - chebi: "CHEBI:62710"
          - metanetx.chemical: "MNXM61056"
          - sbo: "SBO:0000247"
      - deltaG: 453.31
    - !!omap
      - id: "s_1127"
      - name: "mannosylinositol phosphorylceramide B (C26)"
      - compartment: "m"
      - formula: "C56H110NO17P"
      - charge: 0
      - smiles: "CCCCCCCCCCCCCCCCCCCCCCCCCC(=O)NC(COP(=O)(O)OC1C(C(C(C(C1OC2C(C(C(C(O2)CO)O)O)O)O)O)O)O)C(C(CCCCCCCCCCCCCCCC)O)O"
      - annotation: !!omap
          - chebi: "CHEBI:62710"
          - metanetx.chemical: "MNXM61056"
          - sbo: "SBO:0000247"
      - deltaG: 453.31
    - !!omap
      - id: "s_1128"
      - name: "mannosylinositol phosphorylceramide B' (C24)"
      - compartment: "er"
      - formula: "C54H106NO17P"
      - charge: 0
      - smiles: "CCCCCCCCCCCCCCCCCCCCCCC(C(=O)NC(COP(=O)(O)OC1C(C(C(C(C1OC2C(C(C(C(O2)CO)O)O)O)O)O)O)O)C(CCCCCCCCCCCCCCC)O)O"
      - annotation: !!omap
          - chebi: "CHEBI:60418"
          - metanetx.chemical: "MNXM61060"
          - sbo: "SBO:0000247"
      - deltaG: 410.8
    - !!omap
      - id: "s_1129"
      - name: "mannosylinositol phosphorylceramide B' (C24)"
      - compartment: "g"
      - formula: "C54H106NO17P"
      - charge: 0
      - smiles: "CCCCCCCCCCCCCCCCCCCCCCC(C(=O)NC(COP(=O)(O)OC1C(C(C(C(C1OC2C(C(C(C(O2)CO)O)O)O)O)O)O)O)C(CCCCCCCCCCCCCCC)O)O"
      - annotation: !!omap
          - chebi: "CHEBI:60418"
          - metanetx.chemical: "MNXM61060"
          - sbo: "SBO:0000247"
      - deltaG: 410.8
    - !!omap
      - id: "s_1130"
      - name: "mannosylinositol phosphorylceramide B' (C24)"
      - compartment: "m"
      - formula: "C54H106NO17P"
      - charge: 0
      - smiles: "CCCCCCCCCCCCCCCCCCCCCCC(C(=O)NC(COP(=O)(O)OC1C(C(C(C(C1OC2C(C(C(C(O2)CO)O)O)O)O)O)O)O)C(CCCCCCCCCCCCCCC)O)O"
      - annotation: !!omap
          - chebi: "CHEBI:60418"
          - metanetx.chemical: "MNXM61060"
          - sbo: "SBO:0000247"
      - deltaG: 410.8
    - !!omap
      - id: "s_1131"
      - name: "mannosylinositol phosphorylceramide B' (C26)"
      - compartment: "er"
      - formula: "C56H110NO17P"
      - charge: 0
      - smiles: "CCCCCCCCCCCCCCCCCCCCCCCCC(C(=O)NC(COP(=O)(O)OC1C(C(C(C(C1OC2C(C(C(C(O2)CO)O)O)O)O)O)O)O)C(CCCCCCCCCCCCCCCCC)O)O"
      - annotation: !!omap
          - chebi: "CHEBI:62708"
          - metanetx.chemical: "MNXM61051"
          - sbo: "SBO:0000247"
      - deltaG: 453.31
    - !!omap
      - id: "s_1132"
      - name: "mannosylinositol phosphorylceramide B' (C26)"
      - compartment: "g"
      - formula: "C56H110NO17P"
      - charge: 0
      - smiles: "CCCCCCCCCCCCCCCCCCCCCCCCC(C(=O)NC(COP(=O)(O)OC1C(C(C(C(C1OC2C(C(C(C(O2)CO)O)O)O)O)O)O)O)C(CCCCCCCCCCCCCCCCC)O)O"
      - annotation: !!omap
          - chebi: "CHEBI:62708"
          - metanetx.chemical: "MNXM61051"
          - sbo: "SBO:0000247"
      - deltaG: 453.31
    - !!omap
      - id: "s_1133"
      - name: "mannosylinositol phosphorylceramide B' (C26)"
      - compartment: "m"
      - formula: "C56H110NO17P"
      - charge: 0
      - smiles: "CCCCCCCCCCCCCCCCCCCCCCCCC(C(=O)NC(COP(=O)(O)OC1C(C(C(C(C1OC2C(C(C(C(O2)CO)O)O)O)O)O)O)O)C(CCCCCCCCCCCCCCCCC)O)O"
      - annotation: !!omap
          - chebi: "CHEBI:62708"
          - metanetx.chemical: "MNXM61051"
          - sbo: "SBO:0000247"
      - deltaG: 453.31
    - !!omap
      - id: "s_1134"
      - name: "mannosylinositol phosphorylceramide C (C24)"
      - compartment: "er"
      - formula: "C54H106NO18P"
      - charge: 0
      - smiles: "CCCCCCCCCCCCCCCCCCCCCCC(C(=O)NC(COP(=O)(O)OC1C(C(C(C(C1OC2C(C(C(C(O2)CO)O)O)O)O)O)O)O)C(C(CCCCCCCCCCCCCC)O)O)O"
      - annotation: !!omap
          - chebi: "CHEBI:60422"
          - metanetx.chemical: "MNXM61065"
          - sbo: "SBO:0000247"
      - deltaG: 372.76
    - !!omap
      - id: "s_1135"
      - name: "mannosylinositol phosphorylceramide C (C24)"
      - compartment: "g"
      - formula: "C54H106NO18P"
      - charge: 0
      - smiles: "CCCCCCCCCCCCCCCCCCCCCCC(C(=O)NC(COP(=O)(O)OC1C(C(C(C(C1OC2C(C(C(C(O2)CO)O)O)O)O)O)O)O)C(C(CCCCCCCCCCCCCC)O)O)O"
      - annotation: !!omap
          - chebi: "CHEBI:60422"
          - metanetx.chemical: "MNXM61065"
          - sbo: "SBO:0000247"
      - deltaG: 372.76
    - !!omap
      - id: "s_1136"
      - name: "mannosylinositol phosphorylceramide C (C24)"
      - compartment: "m"
      - formula: "C54H106NO18P"
      - charge: 0
      - smiles: "CCCCCCCCCCCCCCCCCCCCCCC(C(=O)NC(COP(=O)(O)OC1C(C(C(C(C1OC2C(C(C(C(O2)CO)O)O)O)O)O)O)O)C(C(CCCCCCCCCCCCCC)O)O)O"
      - annotation: !!omap
          - chebi: "CHEBI:60422"
          - metanetx.chemical: "MNXM61065"
          - sbo: "SBO:0000247"
      - deltaG: 372.76
    - !!omap
      - id: "s_1137"
      - name: "mannosylinositol phosphorylceramide C (C26)"
      - compartment: "er"
      - formula: "C56H110NO18P"
      - charge: 0
      - smiles: "CCCCCCCCCCCCCCCCCCCCCCCCC(C(=O)NC(COP(=O)(O)OC1C(C(C(C(C1OC2C(C(C(C(O2)CO)O)O)O)O)O)O)O)C(C(CCCCCCCCCCCCCCCC)O)O)O"
      - annotation: !!omap
          - chebi: "CHEBI:62712"
          - metanetx.chemical: "MNXM61055"
          - sbo: "SBO:0000247"
      - deltaG: 415.27
    - !!omap
      - id: "s_1138"
      - name: "mannosylinositol phosphorylceramide C (C26)"
      - compartment: "g"
      - formula: "C56H110NO18P"
      - charge: 0
      - smiles: "CCCCCCCCCCCCCCCCCCCCCCCCC(C(=O)NC(COP(=O)(O)OC1C(C(C(C(C1OC2C(C(C(C(O2)CO)O)O)O)O)O)O)O)C(C(CCCCCCCCCCCCCCCC)O)O)O"
      - annotation: !!omap
          - chebi: "CHEBI:62712"
          - metanetx.chemical: "MNXM61055"
          - sbo: "SBO:0000247"
      - deltaG: 415.27
    - !!omap
      - id: "s_1139"
      - name: "mannosylinositol phosphorylceramide C (C26)"
      - compartment: "m"
      - formula: "C56H110NO18P"
      - charge: 0
      - smiles: "CCCCCCCCCCCCCCCCCCCCCCCCC(C(=O)NC(COP(=O)(O)OC1C(C(C(C(C1OC2C(C(C(C(O2)CO)O)O)O)O)O)O)O)C(C(CCCCCCCCCCCCCCCC)O)O)O"
      - annotation: !!omap
          - chebi: "CHEBI:62712"
          - metanetx.chemical: "MNXM61055"
          - sbo: "SBO:0000247"
      - deltaG: 415.27
    - !!omap
      - id: "s_1140"
      - name: "mannosylinositol phosphorylceramide D (C24)"
      - compartment: "er"
      - formula: "C54H106NO19P"
      - charge: 0
      - smiles: "CCCCCCCCCCCCCCCCCCCCCC(C(C(=O)NC(COP(=O)(O)OC1C(C(C(C(C1OC2C(C(C(C(O2)CO)O)O)O)O)O)O)O)C(C(CCCCCCCCCCCCCC)O)O)O)O"
      - annotation: !!omap
          - chebi: "CHEBI:60435"
          - metanetx.chemical: "MNXM61064"
          - sbo: "SBO:0000247"
      - deltaG: 334.72
    - !!omap
      - id: "s_1141"
      - name: "mannosylinositol phosphorylceramide D (C24)"
      - compartment: "g"
      - formula: "C54H106NO19P"
      - charge: 0
      - smiles: "CCCCCCCCCCCCCCCCCCCCCC(C(C(=O)NC(COP(=O)(O)OC1C(C(C(C(C1OC2C(C(C(C(O2)CO)O)O)O)O)O)O)O)C(C(CCCCCCCCCCCCCC)O)O)O)O"
      - annotation: !!omap
          - chebi: "CHEBI:60435"
          - metanetx.chemical: "MNXM61064"
          - sbo: "SBO:0000247"
      - deltaG: 334.72
    - !!omap
      - id: "s_1142"
      - name: "mannosylinositol phosphorylceramide D (C24)"
      - compartment: "m"
      - formula: "C54H106NO19P"
      - charge: 0
      - smiles: "CCCCCCCCCCCCCCCCCCCCCC(C(C(=O)NC(COP(=O)(O)OC1C(C(C(C(C1OC2C(C(C(C(O2)CO)O)O)O)O)O)O)O)C(C(CCCCCCCCCCCCCC)O)O)O)O"
      - annotation: !!omap
          - chebi: "CHEBI:60435"
          - metanetx.chemical: "MNXM61064"
          - sbo: "SBO:0000247"
      - deltaG: 334.72
    - !!omap
      - id: "s_1143"
      - name: "mannosylinositol phosphorylceramide D (C26)"
      - compartment: "er"
      - formula: "C56H110NO19P"
      - charge: 0
      - smiles: "CCCCCCCCCCCCCCCCCCCCCCCC(C(C(=O)NC(COP(=O)(O)OC1C(C(C(C(C1OC2C(C(C(C(O2)CO)O)O)O)O)O)O)O)C(C(CCCCCCCCCCCCCCCC)O)O)O)O"
      - annotation: !!omap
          - chebi: "CHEBI:62683"
          - metanetx.chemical: "MNXM61054"
          - sbo: "SBO:0000247"
      - deltaG: 377.23
    - !!omap
      - id: "s_1144"
      - name: "mannosylinositol phosphorylceramide D (C26)"
      - compartment: "g"
      - formula: "C56H110NO19P"
      - charge: 0
      - smiles: "CCCCCCCCCCCCCCCCCCCCCCCC(C(C(=O)NC(COP(=O)(O)OC1C(C(C(C(C1OC2C(C(C(C(O2)CO)O)O)O)O)O)O)O)C(C(CCCCCCCCCCCCCCCC)O)O)O)O"
      - annotation: !!omap
          - chebi: "CHEBI:62683"
          - metanetx.chemical: "MNXM61054"
          - sbo: "SBO:0000247"
      - deltaG: 377.23
    - !!omap
      - id: "s_1145"
      - name: "mannosylinositol phosphorylceramide D (C26)"
      - compartment: "m"
      - formula: "C56H110NO19P"
      - charge: 0
      - smiles: "CCCCCCCCCCCCCCCCCCCCCCCC(C(C(=O)NC(COP(=O)(O)OC1C(C(C(C(C1OC2C(C(C(C(O2)CO)O)O)O)O)O)O)O)C(C(CCCCCCCCCCCCCCCC)O)O)O)O"
      - annotation: !!omap
          - chebi: "CHEBI:62683"
          - metanetx.chemical: "MNXM61054"
          - sbo: "SBO:0000247"
      - deltaG: 377.23
    - !!omap
      - id: "s_1148"
      - name: "Met-tRNA(Met)"
      - compartment: "c"
      - formula: "C5H11NOSR"
      - charge: 1
      - annotation: !!omap
          - bigg.metabolite: "mettrna"
          - chebi: "CHEBI:16635"
          - kegg.compound: "C02430"
          - metanetx.chemical: "MNXM90636"
          - sbo: "SBO:0000247"
      - deltaG: 10000000
    - !!omap
      - id: "s_1149"
      - name: "Met-tRNA(Met)"
      - compartment: "m"
      - formula: "C5H11NOSR"
      - charge: 1
      - annotation: !!omap
          - bigg.metabolite: "mettrna"
          - chebi: "CHEBI:16635"
          - kegg.compound: "C02430"
          - metanetx.chemical: "MNXM90636"
          - sbo: "SBO:0000247"
      - deltaG: 10000000
    - !!omap
      - id: "s_1150"
      - name: "methanethiol"
      - compartment: "c"
      - formula: "CH4S"
      - charge: 0
      - smiles: "CS"
      - annotation: !!omap
          - bigg.metabolite: "ch4s"
          - chebi: "CHEBI:16007"
          - kegg.compound: "C00409"
          - metanetx.chemical: "MNXM652"
          - sbo: "SBO:0000247"
      - deltaG: 34.55
    - !!omap
      - id: "s_1151"
      - name: "methylglyoxal"
      - compartment: "c"
      - formula: "C3H4O2"
      - charge: 0
      - smiles: "CC(=O)C=O"
      - annotation: !!omap
          - bigg.metabolite: "mthgxl"
          - chebi: "CHEBI:17158"
          - kegg.compound: "C00546"
          - metanetx.chemical: "MNXM310"
          - sbo: "SBO:0000247"
      - deltaG: -21.04
    - !!omap
      - id: "s_1153"
      - name: "myo-inositol"
      - compartment: "c"
      - formula: "C6H12O6"
      - charge: 0
      - smiles: "C1(C(C(C(C(C1O)O)O)O)O)O"
      - annotation: !!omap
          - bigg.metabolite: "inost"
          - chebi: "CHEBI:17268"
          - kegg.compound: "C00137"
          - metanetx.chemical: "MNXM127"
          - sbo: "SBO:0000247"
      - deltaG: -103.12
    - !!omap
      - id: "s_1154"
      - name: "myo-inositol"
      - compartment: "e"
      - formula: "C6H12O6"
      - charge: 0
      - smiles: "C1(C(C(C(C(C1O)O)O)O)O)O"
      - annotation: !!omap
          - bigg.metabolite: "inost"
          - chebi: "CHEBI:17268"
          - kegg.compound: "C00137"
          - metanetx.chemical: "MNXM127"
          - sbo: "SBO:0000247"
      - deltaG: -103.12
    - !!omap
      - id: "s_1156"
      - name: "myo-inositol 1,3,4,5,6-pentakisphosphate"
      - compartment: "c"
      - formula: "C6H7O21P5"
      - charge: -10
      - smiles: "C1(C(C(C(C(C1OP(=O)(O)O)OP(=O)(O)O)OP(=O)(O)O)OP(=O)(O)O)OP(=O)(O)O)O"
      - annotation: !!omap
          - bigg.metabolite: "inospp1"
          - chebi: "CHEBI:57733"
          - kegg.compound: "C01284"
          - metanetx.chemical: "MNXM89719"
          - sbo: "SBO:0000247"
      - deltaG: -1170.46
    - !!omap
      - id: "s_1157"
      - name: "myo-inositol 1,3,4,5,6-pentakisphosphate"
      - compartment: "n"
      - formula: "C6H7O21P5"
      - charge: -10
      - smiles: "C1(C(C(C(C(C1OP(=O)(O)O)OP(=O)(O)O)OP(=O)(O)O)OP(=O)(O)O)OP(=O)(O)O)O"
      - annotation: !!omap
          - bigg.metabolite: "inospp1"
          - chebi: "CHEBI:57733"
          - kegg.compound: "C01284"
          - metanetx.chemical: "MNXM89719"
          - sbo: "SBO:0000247"
      - deltaG: -1170.46
    - !!omap
      - id: "s_1158"
      - name: "myo-inositol hexakisphosphate"
      - compartment: "c"
      - formula: "C6H6O24P6"
      - charge: -12
      - smiles: "C1(C(C(C(C(C1OP(=O)(O)O)OP(=O)(O)O)OP(=O)(O)O)OP(=O)(O)O)OP(=O)(O)O)OP(=O)(O)O"
      - annotation: !!omap
          - bigg.metabolite: "minohp"
          - chebi: "CHEBI:58130"
          - kegg.compound: "C01204"
          - metanetx.chemical: "MNXM491"
          - sbo: "SBO:0000247"
      - deltaG: -1373.22
    - !!omap
      - id: "s_1159"
      - name: "myo-inositol hexakisphosphate"
      - compartment: "n"
      - formula: "C6H6O24P6"
      - charge: -12
      - smiles: "C1(C(C(C(C(C1OP(=O)(O)O)OP(=O)(O)O)OP(=O)(O)O)OP(=O)(O)O)OP(=O)(O)O)OP(=O)(O)O"
      - annotation: !!omap
          - bigg.metabolite: "minohp"
          - chebi: "CHEBI:58130"
          - kegg.compound: "C01204"
          - metanetx.chemical: "MNXM491"
          - sbo: "SBO:0000247"
      - deltaG: -1373.22
    - !!omap
      - id: "s_1161"
      - name: "myristate"
      - compartment: "c"
      - formula: "C14H27O2"
      - charge: -1
      - smiles: "CCCCCCCCCCCCCC(=O)O"
      - annotation: !!omap
          - bigg.metabolite: "ttdca"
          - chebi: "CHEBI:30807"
          - kegg.compound: "C06424"
          - metanetx.chemical: "MNXM314"
          - sbo: "SBO:0000247"
      - deltaG: 195.39
    - !!omap
      - id: "s_1163"
      - name: "myristate"
      - compartment: "e"
      - formula: "C14H27O2"
      - charge: -1
      - smiles: "CCCCCCCCCCCCCC(=O)O"
      - annotation: !!omap
          - bigg.metabolite: "ttdca"
          - chebi: "CHEBI:30807"
          - kegg.compound: "C06424"
          - metanetx.chemical: "MNXM314"
          - sbo: "SBO:0000247"
      - deltaG: 195.39
    - !!omap
      - id: "s_1166"
      - name: "myristate"
      - compartment: "p"
      - formula: "C14H27O2"
      - charge: -1
      - smiles: "CCCCCCCCCCCCCC(=O)O"
      - annotation: !!omap
          - bigg.metabolite: "ttdca"
          - chebi: "CHEBI:30807"
          - kegg.compound: "C06424"
          - metanetx.chemical: "MNXM314"
          - sbo: "SBO:0000247"
      - deltaG: 195.39
    - !!omap
      - id: "s_1176"
      - name: "myristoyl-CoA"
      - compartment: "c"
      - formula: "C35H58N7O17P3S"
      - charge: -4
      - smiles: "CCCCCCCCCCCCCC(=O)SCCNC(=O)CCNC(=O)C(C(C)(C)COP(=O)(O)OP(=O)(O)OCC1C(C(C(O1)N2C=NC3=C(N=CN=C32)N)O)OP(=O)(O)O)O"
      - annotation: !!omap
          - bigg.metabolite: "tdcoa"
          - chebi: "CHEBI:57385"
          - kegg.compound: "C02593"
          - metanetx.chemical: "MNXM224"
          - sbo: "SBO:0000247"
      - deltaG: -215.45
    - !!omap
      - id: "s_1179"
      - name: "myristoyl-CoA"
      - compartment: "p"
      - formula: "C35H58N7O17P3S"
      - charge: -4
      - smiles: "CCCCCCCCCCCCCC(=O)SCCNC(=O)CCNC(=O)C(C(C)(C)COP(=O)(O)OP(=O)(O)OCC1C(C(C(O1)N2C=NC3=C(N=CN=C32)N)O)OP(=O)(O)O)O"
      - annotation: !!omap
          - bigg.metabolite: "tdcoa"
          - chebi: "CHEBI:57385"
          - kegg.compound: "C02593"
          - metanetx.chemical: "MNXM224"
          - sbo: "SBO:0000247"
      - deltaG: -215.45
    - !!omap
      - id: "s_1180"
      - name: "N(1)-acetylspermidine"
      - compartment: "c"
      - formula: "C9H23N3O"
      - charge: 2
      - smiles: "CC(=O)NCCCNCCCCN"
      - annotation: !!omap
          - bigg.metabolite: "N1aspmd"
          - chebi: "CHEBI:58324"
          - kegg.compound: "C00612"
          - metanetx.chemical: "MNXM501"
          - sbo: "SBO:0000247"
      - deltaG: 198.36
    - !!omap
      - id: "s_1181"
      - name: "N(1)-acetylspermine"
      - compartment: "c"
      - formula: "C12H31N4O"
      - charge: 3
      - smiles: "CC(=O)NCCCNCCCCNCCCN"
      - annotation: !!omap
          - bigg.metabolite: "N1sprm"
          - chebi: "CHEBI:58101"
          - kegg.compound: "C02567"
          - metanetx.chemical: "MNXM600"
          - sbo: "SBO:0000247"
      - deltaG: 286.1
    - !!omap
      - id: "s_1182"
      - name: "N(2)-acetyl-L-ornithine"
      - compartment: "m"
      - formula: "C7H14N2O3"
      - charge: 0
      - smiles: "CC(=O)NC(CCCN)C(=O)O"
      - annotation: !!omap
          - bigg.metabolite: "acorn"
          - chebi: "CHEBI:16543"
          - kegg.compound: "C00437"
          - metanetx.chemical: "MNXM817"
          - sbo: "SBO:0000247"
      - deltaG: 25.2
    - !!omap
      - id: "s_1183"
      - name: "N(pros)-methyl-L-histidine"
      - compartment: "c"
      - formula: "C7H11N3O2"
      - charge: 0
      - smiles: "CN1C=NC=C1CC(C(=O)O)N"
      - annotation: !!omap
          - bigg.metabolite: "NPmehis"
          - chebi: "CHEBI:27596"
          - kegg.compound: "C01152"
          - metanetx.chemical: "MNXM722872"
          - sbo: "SBO:0000247"
      - deltaG: 69.78
    - !!omap
      - id: "s_1184"
      - name: "N,N'-diacetylchitobiosyldiphosphodolichol"
      - compartment: "c"
      - formula: "C36H62N2O17P2"
      - charge: -2
      - smiles: "CC(CCC=C(C)CCC=C(C)CCC=C(C)CCC=C(C)C)CCOP(=O)([O-])OP(=O)([O-])OC1C(C(C(C(O1)CO)OC2C(C(C(C(O2)CO)O)O)NC(=O)C)O)NC(=O)C"
      - annotation: !!omap
          - bigg.metabolite: "chito2pdol"
          - chebi: "CHEBI:12427"
          - kegg.compound: "C04537"
          - metanetx.chemical: "MNXM63078"
          - sbo: "SBO:0000247"
      - deltaG: 28.23
    - !!omap
      - id: "s_1185"
      - name: "N,N'-diformyldityrosine"
      - compartment: "c"
      - formula: "C20H22N2O8"
      - charge: -2
      - smiles: "C1=CC(=C(C=C1CC(C(=O)O)NC=O)C2=C(C=CC(=C2)CC(C(=O)O)NC=O)O)O"
      - annotation: !!omap
          - chebi: "CHEBI:50611"
          - metanetx.chemical: "MNXM147516"
          - sbo: "SBO:0000247"
      - deltaG: -58.51
    - !!omap
      - id: "s_1186"
      - name: "N,N'-diformyldityrosine"
      - compartment: "e"
      - formula: "C20H22N2O8"
      - charge: -2
      - smiles: "C1=CC(=C(C=C1CC(C(=O)O)NC=O)C2=C(C=CC(=C2)CC(C(=O)O)NC=O)O)O"
      - annotation: !!omap
          - chebi: "CHEBI:50611"
          - metanetx.chemical: "MNXM147516"
          - sbo: "SBO:0000247"
      - deltaG: -58.51
    - !!omap
      - id: "s_1187"
      - name: "N-(5-phospho-beta-D-ribosyl)anthranilate"
      - compartment: "c"
      - formula: "C12H13NO9P"
      - charge: -3
      - smiles: "C1=CC=C(C(=C1)C(=O)O)NC2C(C(C(O2)COP(=O)(O)O)O)O"
      - annotation: !!omap
          - bigg.metabolite: "pran"
          - chebi: "CHEBI:18277"
          - kegg.compound: "C04302"
          - metanetx.chemical: "MNXM1489"
          - sbo: "SBO:0000247"
      - deltaG: -268.33
    - !!omap
      - id: "s_1188"
      - name: "N-[(R)-4-phosphonopantothenoyl]-L-cysteine"
      - compartment: "c"
      - formula: "C12H20N2O9PS"
      - charge: -3
      - annotation: !!omap
          - bigg.metabolite: "4ppcys"
          - chebi: "CHEBI:59458"
          - kegg.compound: "C04352"
          - metanetx.chemical: "MNXM483"
          - sbo: "SBO:0000247"
      - deltaG: -234.34
    - !!omap
      - id: "s_1189"
      - name: "N-acetyl-alpha-D-glucosamine 1-phosphate"
      - compartment: "c"
      - formula: "C8H14NO9P"
      - charge: -2
      - smiles: "CC(=O)NC1C(C(C(OC1OP(=O)(O)O)CO)O)O"
      - annotation: !!omap
          - bigg.metabolite: "acgal1p"
          - chebi: "CHEBI:16446"
          - kegg.compound: "C04501"
          - metanetx.chemical: "MNXM340"
          - sbo: "SBO:0000247"
      - deltaG: -289.62
    - !!omap
      - id: "s_1190"
      - name: "N-acetyl-D-glucosamine 6-phosphate"
      - compartment: "c"
      - formula: "C8H14NO9P"
      - charge: -2
      - smiles: "CC(=O)NC1C(C(C(OC1O)COP(=O)(O)O)O)O"
      - annotation: !!omap
          - bigg.metabolite: "acgam6p"
          - chebi: "CHEBI:57513"
          - kegg.compound: "C00357"
          - metanetx.chemical: "MNXM63556"
          - sbo: "SBO:0000247"
      - deltaG: -289.62
    - !!omap
      - id: "s_1191"
      - name: "N-acetyl-L-gamma-glutamyl phosphate"
      - compartment: "m"
      - formula: "C7H9NO8P"
      - charge: -3
      - smiles: "CC(=O)NC(CCC(=O)OP(=O)(O)O)C(=O)O"
      - annotation: !!omap
          - bigg.metabolite: "acg5p"
          - chebi: "CHEBI:57936"
          - kegg.compound: "C04133"
          - metanetx.chemical: "MNXM1384"
          - sbo: "SBO:0000247"
      - deltaG: -316.61
    - !!omap
      - id: "s_1192"
      - name: "N-acetyl-L-glutamate"
      - compartment: "m"
      - formula: "C7H9NO5"
      - charge: -2
      - smiles: "CC(=O)NC(CCC(=O)O)C(=O)O"
      - annotation: !!omap
          - bigg.metabolite: "acglu"
          - chebi: "CHEBI:44337"
          - kegg.compound: "C00624"
          - metanetx.chemical: "MNXM730"
          - sbo: "SBO:0000247"
      - deltaG: -100.74
    - !!omap
      - id: "s_1193"
      - name: "N-acetylputrescine"
      - compartment: "c"
      - formula: "C6H15N2O"
      - charge: 1
      - smiles: "CC(=O)NCCCCN"
      - annotation: !!omap
          - bigg.metabolite: "aprut"
          - chebi: "CHEBI:58263"
          - kegg.compound: "C02714"
          - metanetx.chemical: "MNXM1153"
          - sbo: "SBO:0000247"
      - deltaG: 10000000
    - !!omap
      - id: "s_1194"
      - name: "N-carbamoyl-L-aspartate"
      - compartment: "c"
      - formula: "C5H6N2O5"
      - charge: -2
      - smiles: "C(C(C(=O)O)NC(=O)N)C(=O)O"
      - annotation: !!omap
          - bigg.metabolite: "cbasp"
          - chebi: "CHEBI:32814"
          - kegg.compound: "C00438"
          - metanetx.chemical: "MNXM465"
          - sbo: "SBO:0000247"
      - deltaG: -146.32
    - !!omap
      - id: "s_1195"
      - name: "N-formyl-L-kynurenine"
      - compartment: "c"
      - formula: "C11H12N2O4"
      - charge: 0
      - smiles: "C1=CC=C(C(=C1)C(=O)CC(C(=O)O)N)NC=O"
      - annotation: !!omap
          - bigg.metabolite: "Lfmkynr"
          - chebi: "CHEBI:30249"
          - kegg.compound: "C02700"
          - metanetx.chemical: "MNXM1288"
          - sbo: "SBO:0000247"
      - deltaG: 3.66
    - !!omap
      - id: "s_1196"
      - name: "N-formyl-L-tyrosine"
      - compartment: "c"
      - formula: "C10H10NO4"
      - charge: -1
      - smiles: "C1=CC(=CC=C1CC(C(=O)O)NC=O)O"
      - annotation: !!omap
          - bigg.metabolite: "Nfortyr"
          - chebi: "CHEBI:50603"
          - metanetx.chemical: "MNXM63755"
          - sbo: "SBO:0000247"
      - deltaG: -21.3
    - !!omap
      - id: "s_1198"
      - name: "NAD"
      - compartment: "c"
      - formula: "C21H26N7O14P2"
      - charge: -1
      - smiles: "C1=CC(=C[N+](=C1)C2C(C(C(O2)COP(=O)([O-])OP(=O)(O)OCC3C(C(C(O3)N4C=NC5=C(N=CN=C54)N)O)O)O)O)C(=O)N"
      - annotation: !!omap
          - bigg.metabolite: "nad"
          - chebi: "CHEBI:57540"
          - kegg.compound: "C00003"
          - metanetx.chemical: "MNXM8"
          - sbo: "SBO:0000247"
      - deltaG: -285.49
    - !!omap
      - id: "s_1199"
      - name: "NAD"
      - compartment: "er"
      - formula: "C21H26N7O14P2"
      - charge: -1
      - smiles: "C1=CC(=C[N+](=C1)C2C(C(C(O2)COP(=O)([O-])OP(=O)(O)OCC3C(C(C(O3)N4C=NC5=C(N=CN=C54)N)O)O)O)O)C(=O)N"
      - annotation: !!omap
          - bigg.metabolite: "nad"
          - chebi: "CHEBI:57540"
          - kegg.compound: "C00003"
          - metanetx.chemical: "MNXM8"
          - sbo: "SBO:0000247"
      - deltaG: -285.49
    - !!omap
      - id: "s_1200"
      - name: "NAD"
      - compartment: "m"
      - formula: "C21H26N7O14P2"
      - charge: -1
      - smiles: "C1=CC(=C[N+](=C1)C2C(C(C(O2)COP(=O)([O-])OP(=O)(O)OCC3C(C(C(O3)N4C=NC5=C(N=CN=C54)N)O)O)O)O)C(=O)N"
      - annotation: !!omap
          - bigg.metabolite: "nad"
          - chebi: "CHEBI:57540"
          - kegg.compound: "C00003"
          - metanetx.chemical: "MNXM8"
          - sbo: "SBO:0000247"
      - deltaG: -285.49
    - !!omap
      - id: "s_1201"
      - name: "NAD"
      - compartment: "n"
      - formula: "C21H26N7O14P2"
      - charge: -1
      - smiles: "C1=CC(=C[N+](=C1)C2C(C(C(O2)COP(=O)([O-])OP(=O)(O)OCC3C(C(C(O3)N4C=NC5=C(N=CN=C54)N)O)O)O)O)C(=O)N"
      - annotation: !!omap
          - bigg.metabolite: "nad"
          - chebi: "CHEBI:57540"
          - kegg.compound: "C00003"
          - metanetx.chemical: "MNXM8"
          - sbo: "SBO:0000247"
      - deltaG: -285.49
    - !!omap
      - id: "s_1202"
      - name: "NAD"
      - compartment: "p"
      - formula: "C21H26N7O14P2"
      - charge: -1
      - smiles: "C1=CC(=C[N+](=C1)C2C(C(C(O2)COP(=O)([O-])OP(=O)(O)OCC3C(C(C(O3)N4C=NC5=C(N=CN=C54)N)O)O)O)O)C(=O)N"
      - annotation: !!omap
          - bigg.metabolite: "nad"
          - chebi: "CHEBI:57540"
          - kegg.compound: "C00003"
          - metanetx.chemical: "MNXM8"
          - sbo: "SBO:0000247"
      - deltaG: -285.49
    - !!omap
      - id: "s_1203"
      - name: "NADH"
      - compartment: "c"
      - formula: "C21H27N7O14P2"
      - charge: -2
      - smiles: "C1C=CN(C=C1C(=O)N)C2C(C(C(O2)COP(=O)(O)OP(=O)(O)OCC3C(C(C(O3)N4C=NC5=C(N=CN=C54)N)O)O)O)O"
      - annotation: !!omap
          - bigg.metabolite: "nadh"
          - chebi: "CHEBI:57945"
          - kegg.compound: "C00004"
          - metanetx.chemical: "MNXM10"
          - sbo: "SBO:0000247"
      - deltaG: -270.68
    - !!omap
      - id: "s_1204"
      - name: "NADH"
      - compartment: "er"
      - formula: "C21H27N7O14P2"
      - charge: -2
```
